# Patient-derived organoids reveal hypoxia-driven plasticity and therapeutic vulnerabilities in pheochromocytomas and paragangliomas

**DOI:** 10.1101/2025.08.22.671868

**Authors:** Xiaojing Wang, Maite Calucho, Cynthia M. Estrada-Zuniga, Mehwish Noureen, Hector Gonzalez-Cantu, Samantha Mossuto, Jonathan N. Levi, Bethany Landry, Qianjin Guo, Kailee A. Rutherford, Zi-Ming Cheng, Andreanne Sannajust, Sapna Raghunathan, Sanskriti Balaji, Viviane Nascimento da Conceicao, Summer Norris, S. Yusra Munawar, Huyen Thi-Lam Nguyen, Jacob De Rosa, Gail Tomlinson, James X. Wu, Michael W. Yeh, Masha J. Livhits, Anand Vaidya, James Powers, Ron Lechan, Art Tischler, Miao Zhang, Paul Graham, Camilo Jimenez, Mirko Peitzsch, Nicole Bechmann, Graeme Eisenhofer, Yanli Ding, Mio Kitano, Alice Soragni, Patricia L M Dahia

**Affiliations:** Department of Population Sciences, Greehey Children’s Cancer Research Institute, University of Texas Health Science Center at San Antonio, San Antonio, TX, USA; Mays Cancer Center, University of Texas Health Science Center at San Antonio, San Antonio, TX, USA; Department of Orthopaedic Surgery, David Geffen School of Medicine, University of California Los Angeles, CA, USA; Division of Hematology and Medical Oncology, Dept of Medicine, University of Texas Health Science Center at San Antonio, San Antonio, TX, USA; Division of Endrocrinology, Dept of Medicine, University of Texas Health Science Center at San Antonio, San Antonio, TX, USA; Division of Pedriatrics Oncology, Dept Pediatrics, University of Texas Health Science Center at San Antonio, San Antonio, TX, USA; Section of Endocrine Surgery, Geffen School of Medicine, Division of University of California Los Angeles, CA; Center for Adrenal Disorders, Division of Endocrinology, Dept Medicine, Brigham and Women’s Hospital, Harvard Medical School, Boston, MA, USA; Dept Pathology and Laboratory Medicine, Tufts Medical Center, Boston, MA, USA; Dept Medicine, Tufts Medical Center, Boston, MA, USA; Department of Pathology, University of Texas MD Anderson Cancer Center, Houston, TX, USA; Department of Surgical Oncology, University of Texas MD Anderson Cancer Center, Houston, TX, USA; Department of Endocrine Neoplasia, University of Texas MD Anderson Cancer Center, Houston, TX, USA; Institute for Clinical Chemistry and Laboratory Medicine, University Hospital Carl Gustav Carus, Medical Faculty Carl Gustav Carus, Technische Universität Dresden, Fetscherstrasse 74, 01307 Dresden, Germany; Dept Pathology, University of Texas Health Science Center at San Antonio, San Antonio, TX, USA; Division of Surgical Oncology, University of Texas Health Science Center at San Antonio, San Antonio, TX, USA; Jonsson Comprehensive Cancer Center, University of California Los Angeles, CA; Eli and Edythe Broad Center of Regenerative Medicine and Stem Cell Research, University of California Los Angeles, CA

## Abstract

Pheochromocytomas and paragangliomas (PPGLs) are rare chromaffin cell-derived neuroendocrine tumors of sympathetic (catecholamine-producing) or parasympathetic (nonsecretory) origin, frequently driven by dysregulation of hypoxia-inducible factor (HIF) signaling, particularly HIF-2α. Although often benign, PPGLs can metastasize unpredictably, with limited therapeutic options once disseminated. Progress has been hindered by the lack of robust preclinical models, especially those that capture their molecular complexity and microenvironmental influences. To address this gap, we established patient-derived tumor organoids (PDOs) from 35 PPGLs, encompassing a broad spectrum of clinical and molecular phenotypes. The organoids retained key immunohistochemical, genomic, transcriptomic, and catecholamine-secretory features of their parental tumors. PPGL organoids cultured under hypoxic conditions generally exhibited enhanced viability, supporting hypoxia as a driver of cell survival. Hypoxia activated HIF-1α and expanded ASCL1^+^ cell populations, suggesting a lineage shift toward an immature chromaffin state. In contrast, long-term normoxic cultures activated hypoxia inducible factor 2α (HIF-2α) and acquired a hybrid sympathoblast–mesenchymal identity in subpopulations with upregulation of extracellular matrix and cell cycle markers, independent of genotype. These features resemble high-risk neuroblastoma subtypes and establish a molecular parallel suggestive of shared lineage plasticity and pathogenic programs, detectable in primary PPGLs. Drug screening across a library of up to 51 drugs and combinations revealed both shared and unique vulnerabilities, with response rates to approved therapies matching clinical observations. The CDK4/6 inhibitor abemaciclib, previously unexplored in PPGLs, elicited the strongest activity. Abemaciclib-responsive PDOs and their matched tumors, including a metastatic sample, exhibited epithelial mesenchyme transition enrichment, nominating potential biomarkers for patient stratification. Our results establish PDOs as a novel platform for modeling neuroendocrine tumor biology, reveal microenvironment-driven plasticity in PPGLs, with potential translational relevance, and identify actionable vulnerabilities in a disease with few effective systemic therapies.

**Main findings:**

1. PDOs can be successfully generated from PPGLs of various genetic backgrounds and reflect parental tumor properties
2. PDO cultures grown in hypoxia retain main molecular features of parental tumors, have increased viability and a more immature developmental/biosynthetic profile
3. Long term PDOs grown for 4 weeks in normoxia activate HIF2α, drift toward a hybrid sympathoblast-mesenchymal-like identity resembling relapsed/therapy resistant neuroblastomas, features that can be detected in primary tumors
4. A subset of PDOs respond to Abemaciclib, a drug class not previously used therapeutically in PPGLs

## Introduction

Pheochromocytomas and paragangliomas (PPGLs) are rare neuroendocrine tumors that originate from autonomic nervous system cells in the adrenal or extra adrenal paraganglia, respectively^1^. These tumors often produce and secrete catecholamines such as norepinephrine and epinephrine, or both, underlying their variable clinical presentations^2^. PPGLs are classified into three molecular groups: pseudo-hypoxic (cluster 1), kinase signaling (cluster 2), and Wnt alterations (cluster 3), each closely associated with their main genetic driver and reflective of their catecholamine profile^3^. Driver mutations in PPGLs occur in a mutually exclusive manner, with about one-third present in the germline^4^.

Alterations of hypoxia pathway genes are a major component of the molecular pathogenesis of the majority of PPGLs, including in many tumors that progress to metastases^1^. Mutations occur in genes that directly or indirectly cause constitutive activation of hypoxia inducible factor 2 alpha subunit (HIF2α), a key component of the physiological response to hypoxia^4,5^. HIF2α is required for the development of sympathoadrenal tissues^6,7^. This dependence on hypoxia has also been shown in rodent models, where sympathoadrenal cells are developmentally dependent on hypoxia and cell lines derived from rodent pheochromocytoma models preferentially grow under hypoxic conditions^8^. EPAS1, the gene encoding HIF2α, is itself a target of somatic mutations in PPGLs^9–12^. Intriguingly, systemic hypoxia related to certain congenital cardiopathies increases the risk for PPGLs carrying EPAS1 activating mutations, and the low relative O_2_ of high-altitude has also been linked to increase the risk of PPGLs^13–15^. Together, these observations support a close link between hypoxia, HIF2α and PPGLs, with underlying mechanisms and translational implications that remain to be fully elucidated in patient-relevant models.

While often benign, approximately 15% of pheochromocytomas and up to 50% of paragangliomas can metastasize, with 5-year survival rate lower than 50%^16,17^. Some PPGLs progress with multiple recurrences that may become inoperable and difficult to manage. Surgical histopathology cannot reliably distinguish between benign and malignant PPGLs, and the diagnosis of malignancy relies on metastatic spread to non-chromaffin tissue. The malignant transformation of PPGLs can therefore be unpredictable, and a diagnosis of malignancy relies on the presence of metastases, leading to late or retrospective detection with negative impact on patient outcomes^18^. In addition, some PPGLs progress with multiple recurrences that may become inoperable and challenging for clinical management^19^. There are few therapeutic options for metastatic and unresectable tumors, with modest survival benefits^17,19^. The inability to identify highly recurrent, aggressive PPGLs before they progress, coupled with the lack of effective therapies for metastatic or inoperable disease, represents a critical challenge^17,19^.

The full spectrum of drug-sensitivity in primary or metastatic PPGLs remains unknown. Clinical trials are few, further constrained by a small and molecularly diverse patient population, contributing to slow therapeutic advancements and a continued unmet need for effective therapies^17,20^. Advancing our understanding of PPGL biology is therefore critical to enable early detection of malignant, aggressive, and recurrent tumors, and to identify, validate, and devise novel targeted strategies. Yet, progress towards closing these gaps has been hindered by the lack of clinically relevant experimental models that faithfully recapitulate the molecular and microenvironmental complexity of these rare neoplasms.

Models based on cell lines or patient derived xenografts (PDXs) are scarce and fail to fully capture the genetic, biochemical, and microenvironmental heterogeneity of PPGLs^21–24^. Preclinical studies have largely used rodent lines such as PC12 or MPC/MTT, which differ from human tumors in catecholamine metabolism, growth characteristics, and differentiation status, while the only available immortalized human line, hPheo1, represents a single genetic background and a restricted differentiation state, retaining some but not all features of primary PPGLs^25^.

Patient-derived tumor organoids (PDOs) have proven capable of faithfully modeling the architecture, lineage differentiation, and functional dependencies of a wide range of epithelial malignancies^26,27^. Until recently, application of PDOs to non-epithelial tumors was limited largely to brain tumors, with few exceptions^27–30^. Building on this gap, we have established organoids from non-epithelial tumors, including benign neurofibromas and over twenty sarcoma subtypes within our mini-ring platform that preserves the original tumor cellular heterogeneity and enables rapid, high throughput drug screening alongside detailed mechanistic studies to identify and validate therapeutic strategies^31,32^.

Here we present a scalable platform for the establishment of PDOs generated from PPGLs of 35 patients. We validated PDOs from genetically diverse PPGL specimens grown under varying oxygen levels, developing both short- and long-term unpassaged models. These organoids reproduce the histochemical array, genomic alterations, transcriptomic profiles, and catecholamine secretion patterns of their source tumors. Under hypoxic conditions, PPGL PDOs exhibit enhanced viability and adopt a more immature biosynthetic profile. In contrast, long-term normoxic organoids activate HIF2α and transcriptionally converge on a sympathoblast-mesenchymal state akin to therapy resistant neuroblastomas. Lastly, we identify unique drug sensitivities, subsets of cases that may benefit from existing therapies as well as nominate abemaciclib as a potential new therapy for a subset of PPGLs. By deploying a human-based, robust, and scalable preclinical platform that captures PPGL heterogeneity and microenvironmental influences, we provide a new tool for probing disease mechanisms and accelerating targeted treatment and biomarker discovery in the rare disease setting.

## Results

### Establishment of genetically and clinically diverse PPGL organoids for short- and long-term modeling under normoxia and hypoxia

To establish the feasibility of developing PPGL-derived organoids, we procured tissue from 35 independent patients diagnosed with PPGL who enrolled in IRB-approved protocols at four participating centers (**Figure 1a-e, Supplementary Table 1**). The cohort included patients with a median age of 47.5 years (range 15-83) and 57% female in aggregate (70.8% for pheochromocytomas and 27% for paragangliomas), representing distinct clinical histories and a wide variety of racial and ethnic backgrounds, and genetic drivers (**Figure 1b-c**). Twenty-five tumors were pheochromocytomas and 11 were paragangliomas (**Figure 1b**), including one of head and neck (carotid body) location (UTH1314). Three of the pheochromocytomas were primary tumors or local recurrence derived from patients with synchronous or metachronous metastatic/recurrent disease (UTH1301, UTH1308 and UTH1328, **Supplementary Table 1**). An additional patient (UTH1212) developed a recurrence of the pheochromocytoma and development of a new paraganglioma within a year of the initial surgery. Three patients had received prior treatment before the surgical procedure from which samples were collected. One patient with a carotid body paraganglioma underwent tumor embolization prior to surgical resection, while two others with locally recurrent tumors, who developed metastases either concurrently or during follow up received radiation for bone lesions (**Supplementary Table 1**).

**Figure 1.**
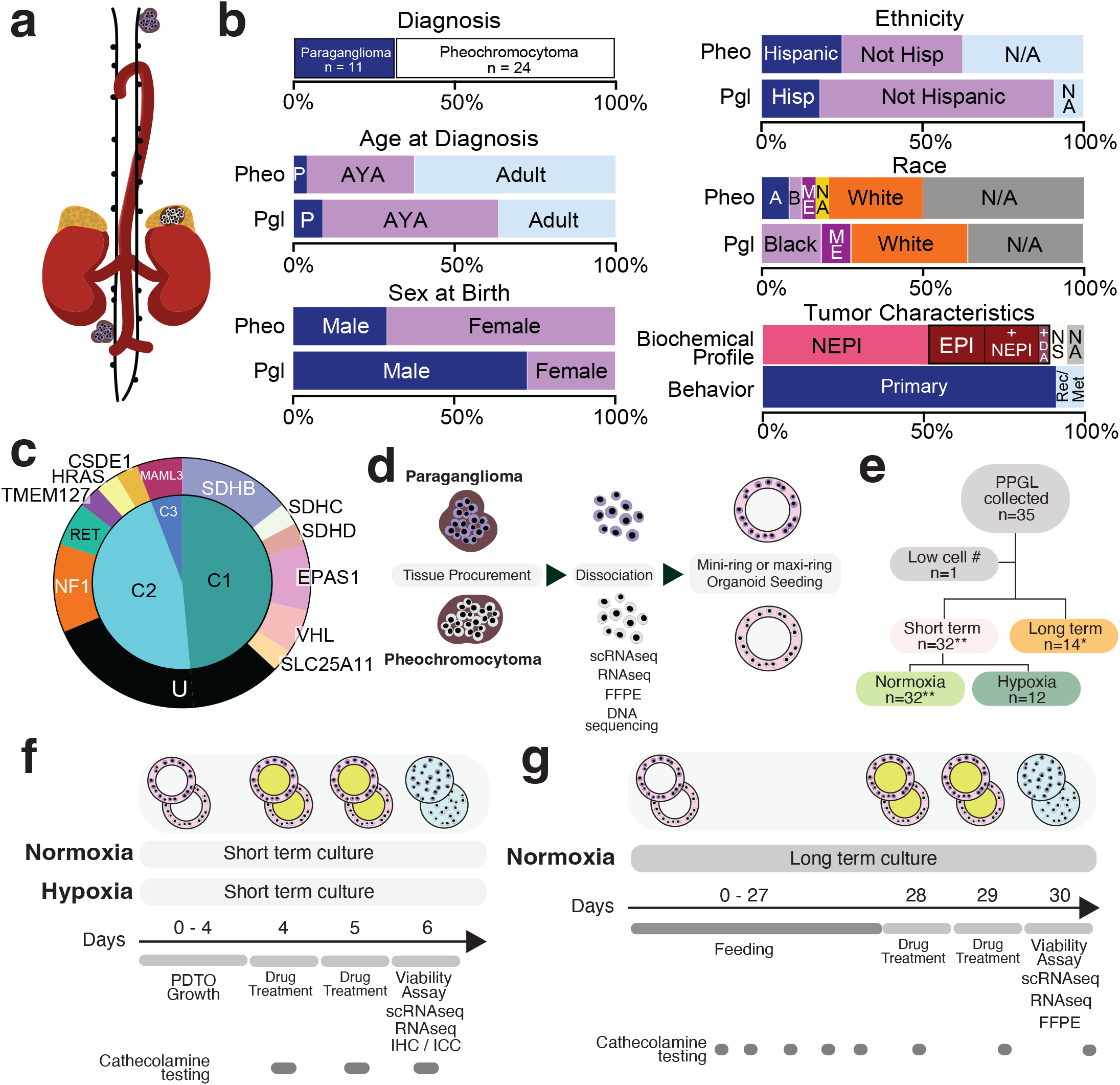
Study design and clinical, molecular, and experimental features of the PPGL PDO cohort. a) Sketch illustrating the location of pheochromocytomas (adrenal medulla) and paragangliomas of neck and abdominal/ pelvic area collected in this study; b) Summary of the study cohort based on tumor location (diagnosis), age at diagnosis, sex, race, ethnicity, biochemical profile and tumor behavior, P=pediatric, AYA=adolescents/young adults, A=Asian, M.E.=middle east, Hisp=Hispanic, B=Black, NA= Native American, NE-PI=norepinephrine, EPI=epinephrine, NS=non secreting, Rec=recurrent, Met=metastatic, N/A=not applicable, not known; c) Distribution of the genetic and molecular features of the cohort used in this study; d) Schematic of sample processing and seeding; e) Number of samples processed and used for generating organoids under three culture conditions; f) Schematic of short term cultures: conditions, treatment, collection and downstream experiments, PDTO (or PDO)=patient-derived tumor organoid, scRNAseq=single cell RNAseq, IHC=immunohistochemistry, ICC=immunocytochemistry; g) Schematic of long term cultures: conditions, treatment, collection and downstream experiments; FFPE=formalin-fixed, paraffin-embedded.

Of the 33 patients with preoperative catecholamines and/or metanephrine measurements, 10 had elevated epinephrine (EPI) and/or metanephrine (MN); most of these patients also had high levels of norepinephrine (NEPI) or normetanephrine (NMN) (**Supplementary Table 1**). Eighteen patients had isolated elevations of NEPI and/ or NMN and while three had preoperatory catecholamines and/or metanephrines levels within the normal range.

The main genetic driver event underlying the PPGL diagnosis was established in 25 patients (71.4%). Among 22 samples with matched normal DNA available for analysis, 12 (54%) carried a germline mutation and 10 (45%) had a detectable somatic variant (**Figure 1c, Supplementary Table 1**). The most common genotype involved a variant in one of the SDH subunit genes (n=7/35, 20%), including a rare somatic SDHB mutation. The three main molecular clusters, as defined by transcription profiling and/or genotyping, were represented in this cohort: pseudohypoxic (or also known as molecular cluster 1) in 17 samples, kinase signaling (cluster 2) in 16 samples, and WNT signaling (cluster 3) in two samples carrying UBT-F::MAML3 fusions. In general, our cohort includes a robust representation of patients with diverse demographic characteristics, genetic backgrounds, molecular clustering, biochemical profiles, and disease behaviors.

Of the 35 tumors collected, one did not produce sufficient cells to initiate culture (UTH1305, **Figure 1e**). All the remaining samples were seeded in a ring-shaped geometry in Matrigel (**Figure 1f-g**), either as maxi-rings in 24-well plates for hematoxylin and eosin staining (H&E) staining, immunohistochemistry, or molecular analyses (bulk or single-cell RNA-seq, catecholamine secretion) or as automation-compatible mini-rings in 96-well plates for immunofluorescence and drug screening, following our established pipelines^33,34^. Each assay was performed on subsets of samples, depending on tissue availability and experimental design. Three samples showed no visible viable cells on H&E of formalin-fixed paraffin-embedded samples (FFPE), likely due to extensive calcification (UTH1216, UTH1219, and UTH1226). Cultures were maintained under normoxia for six days as per our standard conditions (n=32, PDO-N; **Figure 1f**). We elected to grow a subset of 14 samples for four weeks without passaging (named “long-term” or PDO-L; **Figure 1g**) to preserve the original cellular composition and tumor heterogeneity, while also assessing viability, stability, and potential molecular or phenotypic evolution over time. In addition, to investigate the role of hypoxia in PPGL biology and drug responses, PDOs established from 16 patients were cultured in hypoxic conditions (1% O_2_, PDO-H) for six days (**Figure 1f**), enabling the evaluation of oxygen-dependent effects on viability, differentiation state, cell composition and catecholamine metabolism.

### PPGL-derived organoids recapitulate cellular composition and hypoxia-driven phenotypes of their tumors of origin

Short- and long-term normoxia cultures from pheochromocytomas and paragangliomas of diverse genetic backgrounds grew robustly in our platform, forming dense clusters and spindle-shaped structures visible in brightfield imaging (**Figure 2a**). PDOs from twenty-three PPGL patients were fixed and paraffin embedded for histopathological analysis. The morphology of shortterm cultures was independent of the media tested, so we proceeded with composition-controlled serum-free medium for all subsequent studies (**Supplementary Figure 1a** and **Methods**).

**Figure 2.**
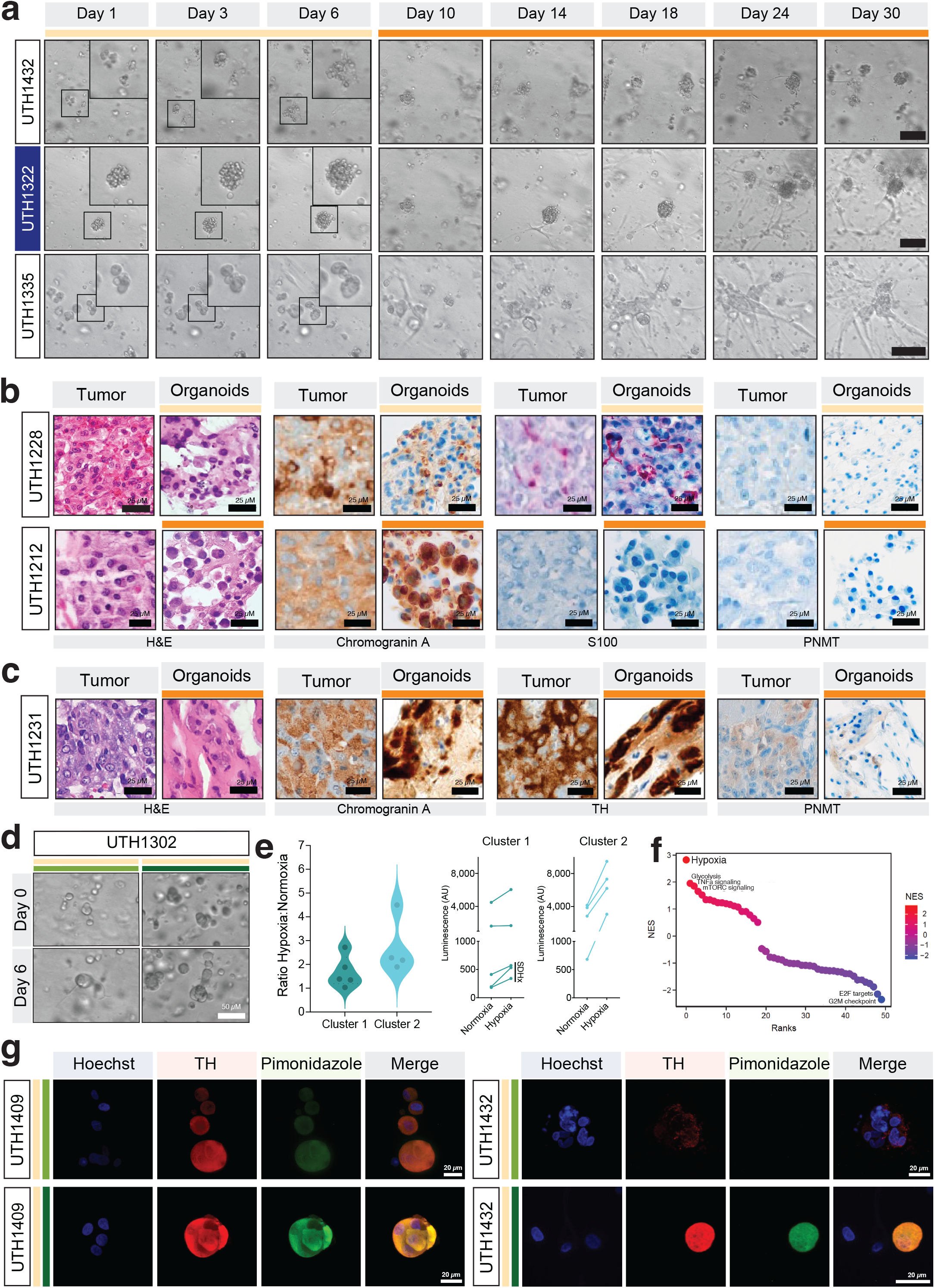
PPGL PDOs preserve parental tumor immunophenotype and display enhanced viability under hypoxia. a) Brightfield images of PDOs cultured under normoxia at different time points, derived from representative tumors of the three molecular clusters: Cluster 1/pseudohypoxia=UTH1322, Cluster 2/kinase signaling=UTH1335, and Cluster 3/Wnt signaling=UTH1432, inset shows magnification of the indicated boxed area of short term cultures, scale bar is 25µm; b) Images of the primary tumors and matched short term (top row) or long term (bottom row) PDO of two samples displaying: H&E staining, and immunohistochemistry (IHC) for chromogranin A, S100, and PNMT (top row is a short term culture, UTH1228), bottom row is a long term culture (UTH1212), scale bar is as indicated; c) IHC of chromogranin A, tyrosine hydroxylase (TH) and PNMT of the primary tumor and respective long term PDO (UTH1231), scale bar is as indicated; d) Brightfield images of a PDO cultured under normoxia and hypoxia at seeding day and at day 6 (UTH1302), scale bar is as indicated; e) Viability (ATPase) assay of PDOs of organoids performed at day 6 in matched normoxia/hypoxia cultures measured in 2-6 replicates per sample displaying ratio hypoxia/normoxia of each sample based on the molecular cluster of parent tumor (left) or average values per condition and cluster type (Cluster 2, p=0.03 ratio paired t-test; N=normoxia, H=hypoxia, SDH-mutant tumors are indicated); f) Gene set enrichment analysis (GSEA) of hypoxia and normoxia PDOs ordered by their normalized enrichment scores (NES), scale of positively enriched to negatively enriched NES is shown on the right; g) Immunofluorescence (IF) image of TH (red), the hypoxia sensor pimonidazole (green), Hoescht (blue, staining nuclei) and merged image of two paired normoxia (top)/hypoxia (bottom) PDO cultures, scale bar is as indicated.

PDOs generally did not form a characteristic nest-like ‘zellballen’ patterns found in many PPGLs (**Figure 2b**), an architecture critically dependent on fibrovascular septa^35,36^, which is not expected to emerge in avascular ex vivo culture systems. Despite the absence of this tissue-level feature, both short and long term PDOs contained cells expressing the neural crest marker chromogranin A (ChGA), and the sustentacular cell marker S100, confirming the presence of both tumor and supporting cells (**Figure 2b-c**). Among the PDOs analyzed, one lacked S100-positive cells, consistent with its parent PPGL (UTH1212, **Figure 2b**). This PDO was established from a patient who recurred within 12 months of surgery. S100-negative staining has been suggested to correlate with clinically aggressive tumors^37^. We also performed IHC of the endothelial vascular marker CD34. Interestingly, the sample with the strongest CD34 expression was a tumor with pseudohypoxic molecular profile (UTH1228; **Supplementary Figure 1b-c**), a group which is typically enriched for angiogenic signaling^38,39^.

Importantly, ChGA-positive tumor cells remained present at later stages of PDO culture (PDO-L, **Figure 2b-c**). In UTH1231, a cluster 2 pheochromocytoma harboring a CSDE1 mutation, long-term cultures expressed the sympathoadrenal lineage marker tyrosine hydroxylase (TH) and RET with similar distribution (**Figure 2c** and **Supplementary Figure 1d**). RET expression, typically enriched in kinase-type PPGLs^40,41^, supports the retention of sympathoadrenal properties throughout the culture timeframe (**Supplementary Figure 1d**). In this PDO-L we detected a subset of cells expressing phenyl ethanolamine N-methyltransferase (PNMT), the enzyme responsible for the conversion of norepinephrine to epinephrine, mirroring the parent tumor (**Figure 2c**). In contrast, PNMT expression was absent in two other PDOs and their corresponding parent PPGLs (UTH1212 and UTH1228, **Figure 2b**), further supporting the notion that PPGL PDOs preserve key features of the primary tumors. Overall, PPGL PDO cultures maintained the cell diversity and characteristics that closely reflect their parental tumors.

Consistent with observations in rodent PPGL models^42,43^ cultures maintained at 1% O_2_ (PDO-H) exhibited higher cell viability relative to their PDO-N counterparts as measured by ATP-release assay (~1.8 ±1.03 fold on average, p=0.06, **Figure 2d-e**), with the effect more pronounced in samples from the molecular kinase group (2.57 ±0.87 fold vs 1.13±0.96 for hypoxic, p=0.03, **Figure 2e**). Interestingly, the three samples with SDH mutations: two with SDHB (UTH1314 and UTH1322) and one with an SDHD mutation (UTH1306), showed viability levels in hypoxia comparable to those observed in normoxia (**Figure 2e**), potentially reflecting their pre-existing pseudohypoxia activation, so that further oxygen reduction does not confer any additional growth advantage. The UTH1314 PDO derived from a non-secreting paraganglioma that underwent embolization prior to surgical removal to reduce tumor volume (**Supplementary Table 1**), a procedure known to induce fibrosis and tissue scarring, which may have contributed to the reduced ATP levels of this sample. RNAseq and pathway enrichment analysis identified hypoxia as the most enriched pathway in these samples (**Figure 2f, Supplementary Tables 2 and 3**). Lastly, we confirmed the presence of hypoxic regions within PDO-H cultures by detecting pimonidazole adducts (**Figure 2g**). Pimonidazole is a well-established marker of intracellular hypoxia^44^ and was detected in tumor (TH-positive) cells (**Figure 2g**), further validating the hypoxic status of tumor cells in these cultures. Overall, these results provide evidence that PPGL-derived organoids faithfully preserve cellular diversity and molecular features of PPGLs of various genetic backgrounds and further indicate that hypoxia supports organoid viability in patient-derived models.

### PPGL PDOs Preserve Catecholamine Secretion Profiles, with Hypoxia Promoting Immature Phenotypes and Prolonged Culture Reducing Biosynthetic Capacity

A defining functional hallmark of PPGLs is their capacity to synthesize and secrete catecholamines, a process that underlies many of the clinical symptoms associated with these tumors^45^. To assess whether PDOs preserve this key property, we measured the levels of catecholamines (dopamine-DA, norepinephrine-NEPI and epinephrine-EPI) and their metabolites, 3-methoxytyramine (3-MTY), normetanephrine (NMN) and metanephrines (MN) in the supernatant of PDO cultures derived from 16 PPGL tumors (**Figure 3a-c**). Catecholamine levels were measured in the supernatant of 16 short-term PDO cultures at three time points. Nine long-term PDO cultures were assessed at five time points (**Figure 3a**). Catecholamine levels were also measured in matched parental tumors for comparison in a subset of cases (n=8; **Figure 3d** and **Supplementary Table 4**).

**Figure 3.**
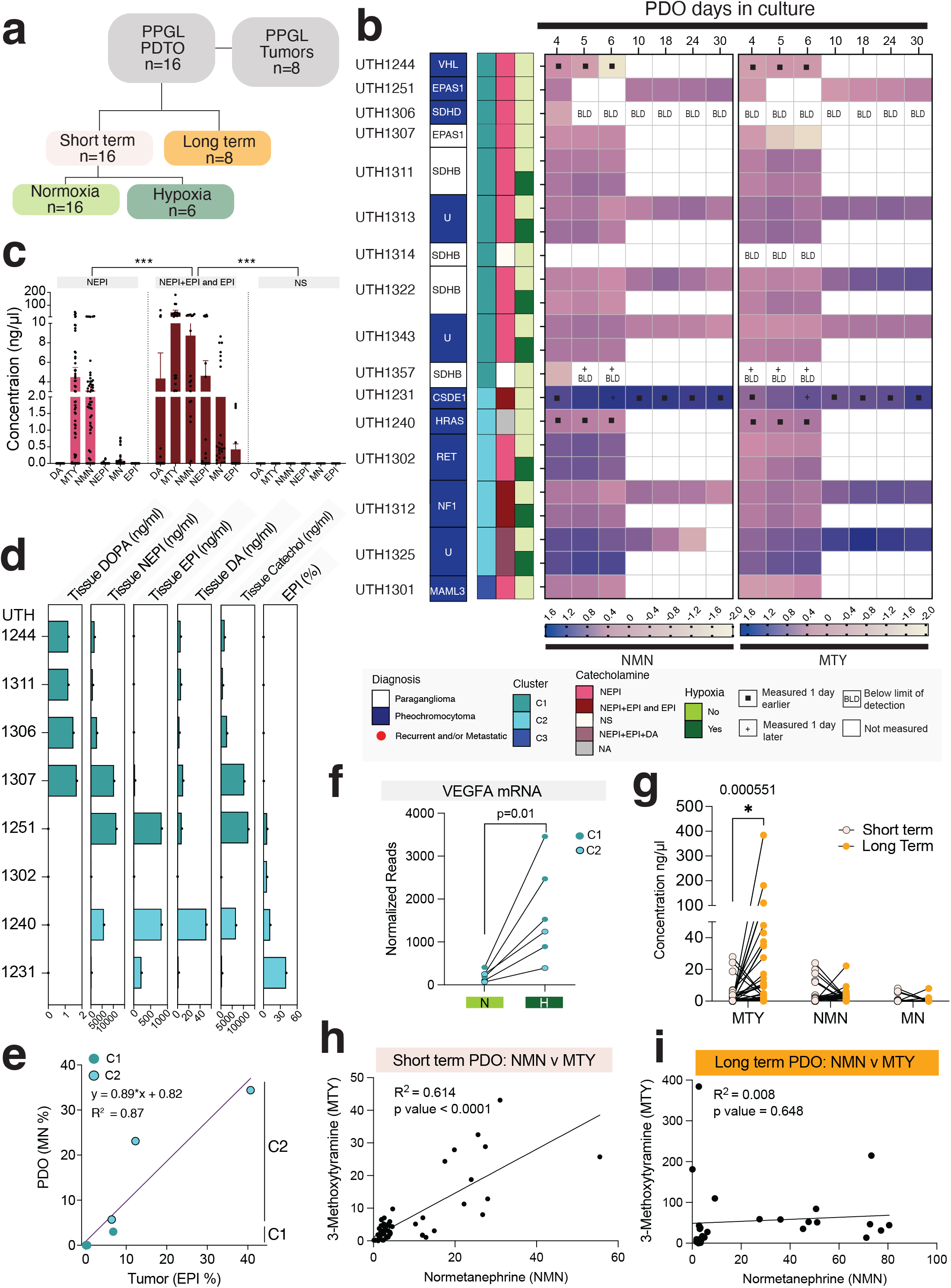
PPGL PDOs are functional and reproduce catecholamine secretion patterns of parental tumors. a) Number of PDO samples and parent tumors from which catecholamines and metanephrines were measured from supernatant or tissue, respectively. PDO culture conditions are indicated (related to Suppl Table 3); b) Log transformed Normetanephrine (NMN) and 3-Methoxythyramine (MTY) levels in supernatant PDO short term normoxic, hypoxic and long term cultures at the designated days; scale bar is shown, and features of the derived parental tumor are as indicated; c) Aggregate levels of DA-dopamine, NEPI-norepinephrine, EPI-epinephrine, NMN, MN-metanephrine and MTY in culture supernatant of samples split by the biochemical profile of tumors prior to surgery (NEPI-secreting, NEPI+EPI- or EPI-secreting, or Nonsecretory-NS; d) Amounts of DOPA, NEPI, EPI, DA, total catecholamines and % EPI from 8 tumors, colored by their molecular cluster (green=cluster 1, teal=cluster 2); e) Correlation between the % epinephrine (EPI) in tumor tissue and % metanephrine (MN) in corresponding PDO supernatant; f) VEGFA mRNA in paired normoxic (N) and hypoxic (H) PDO samples collected at day 6 of culture (p value=paired t-test); g) MTY, NMN and MN values in paired short-term/long term PDO supernatant (p value=paired t-test); h) Correlation between MTY and NMN levels in short-term cultures, r, CI, r^2^, p value are shown; i) Correlation between MTY and NMN levels in long-term cultures, r, CI, r2, p value are shown.

In PDOs, metanephrines were detectable at higher levels than catecholamines (**Figure 3b-c**), often exceeding the range observed in patients’ diagnostic preoperative samples. In some cultures, metanephrine concentrations were two orders of magnitude higher than typical plasma values, such as 0.005–0.5 ng/ml in plasma versus ~100 ng/ml in PDOs (**Figure 3b-c** and **Supplementary Table 1**)^46,47^. Metanephrines were measurable at the earliest timepoint tested, 3 to 4 days after organoid seeding (**Figure 3b**). There was overall concordance between metanephrines and catecholamine profiles detected in PDO supernatants and the dominant preoperative plasma and/or urine profile of the patients (**Figure 3b-c, Supplementary Tables 1 and 4**), as well as with the molecular cluster of the parental tumors (**Supplementary Figure 2a**). Metanephrines remained detectable until the end of culture in 14/16 short-term PDOs and 6/8 long-term PDOs, although levels generally declined over time (**Figure 3b, Supplementary Figure 2b-c** and **Supplementary Table 4**). No catecholamines or metanephrines were detected in PDOs derived from two clinically non-secretory tumors, corresponding to patients with normal preoperative catecholamine and metanephrine levels (UTH1314 and UTH1357; **Figure 3b-c**).

Kinase-type (molecular cluster 2) PDOs had higher overall levels of catecholamines and metanephrines than pseudohypoxia-related PDOs (**Figure 3b-c** and **Supplementary Figure 2a**), consistent with reported differences in tumor catecholamine content between these molecular groups^45^. PDOs derived from five kinase-type (C2) PPGLs showed detectable MN and NMN, those from eleven pseudohypoxic-type (C1) tumors had only NMN, and three pseudohypoxic-type along with the single WNT-signaling (C3) PDOs displayed predominant NMN with trace amounts of MN (**Figure 3b, Supplementary Figure 2a-c**, and **Supplementary Table 4**). We observed a remarkable concordance between the catecholamine profiles of PDOs and their matched tumors (R_2_ = 0.87; **Figure 3d-e**). For instance, PDOs derived from an EPAS1-mutant tumor (UTH1251) secreted both NMN and MN, mirroring the tumor’s catecholamine profile (**Figure 3d-e** and **Supplementary Figure 2b**), but differing from the patient’s preoperative plasma, which showed elevated NMN only (**Supplementary Table 1**). This suggests that PDO supernatants may more closely reflect the tumor’s intrinsic biochemical profile.

Hypoxic cultures (PDO-H) showed significantly higher NMN and MTY levels than their matched normoxic PDO-N (p<0.05; **Figure 3b** and **Supplementary Figure 2d**). These elevations were consistent with the hypoxia-inducible increase in dopamine and NEPI previously seen in rodent pheochromocytoma cells^43^. They were accompanied by transcriptional changes associated with hypoxia, including increased VEGF mRNA levels, regardless of the molecular type of the parent tumor (**Figure 3f**). However, the hormonal increase was much more attenuated for the mature catecholamine epinephrine (EPI) and its metabolite metanephrine (MN; **Supplementary Figure 2e**). This suggests that hypoxia impaired the terminal step of catecholamine biosynthesis, resulting in diminished catecholamine maturation. We next examined the impact of extended time in culture on catecholamine production. In PDO-L, 3-MTY levels increased (**Figure 3g** and **Supplementary Table 4**) and their correlation with NMN was markedly reduced compared with short-term cultures (r = 0.008 vs 0.78, p < 0.001; **Figures 3h-i**). A similar shift occurred for the respective catecholamines, with the DA-NE-PI correlation dropping from r = 0.96 in short-term cultures to 0.05 in long-term cultures (**Supplementary Figures 2f-g**). Furthermore, mRNA levels of catecholamine biosynthetic genes in eight matched short-term and long-term PDOs were lower at day 30 than in their short-term counterparts (**Supplementary Figure 2h**), consistent with the progressive decrease in NMN levels in the culture supernatant over time. These findings indicate that prolonged culture is associated with a decline in catecholamine biosynthetic capacity in PPGL PDOs. In conclusion, PPGL PDOs recapitulate the biochemical profiles of their parental tumors in short-term cultures, that hypoxia preferentially increases immature catecholamines, and that prolonged culture times diminish the tumor cells’ capacity to synthesize and store catecholamines.

### Hypoxia maintains transcriptional features in PPGL PDOs while long-term culture induces genotype independent pseudohypoxic profiles

To understand the dynamic molecular events of the PDOs in response to hypoxia or time in culture, we evaluated the transcription distribution of these samples using UMAP plotting of RNAseq primary and PDO samples, including 53 bulk RNAseq and 30 pseudobulk’ RNAseq from single cell/single nuclei (sc/snRNAseq), including pseudobulk data from 6 other PPGLs from which we had previously performed snRNAseq^41^ (**Figure 4a** and **Supplementary Figure 3a**). In addition, to enhance the clustering strength of the data, we integrated these samples with 212 additional primary PPGL samples comprising bulk RNAseq from a group of 28 additional UTHSA samples from our cohort^48^ and 183 samples from the PPGL TCGA cohort (PCPG)^49^. In total, 294 primary tumors and PDO PPGLs were used for UMAP plotting (**Supplementary Table 5**). This approach revealed the expected distribution of primary tumors into three main molecular clusters, pseudohypoxic (C1), kinase (C2) and WNT (C3) signaling (**Figure 4b**)^3,48,50^.

**Figure 4.**
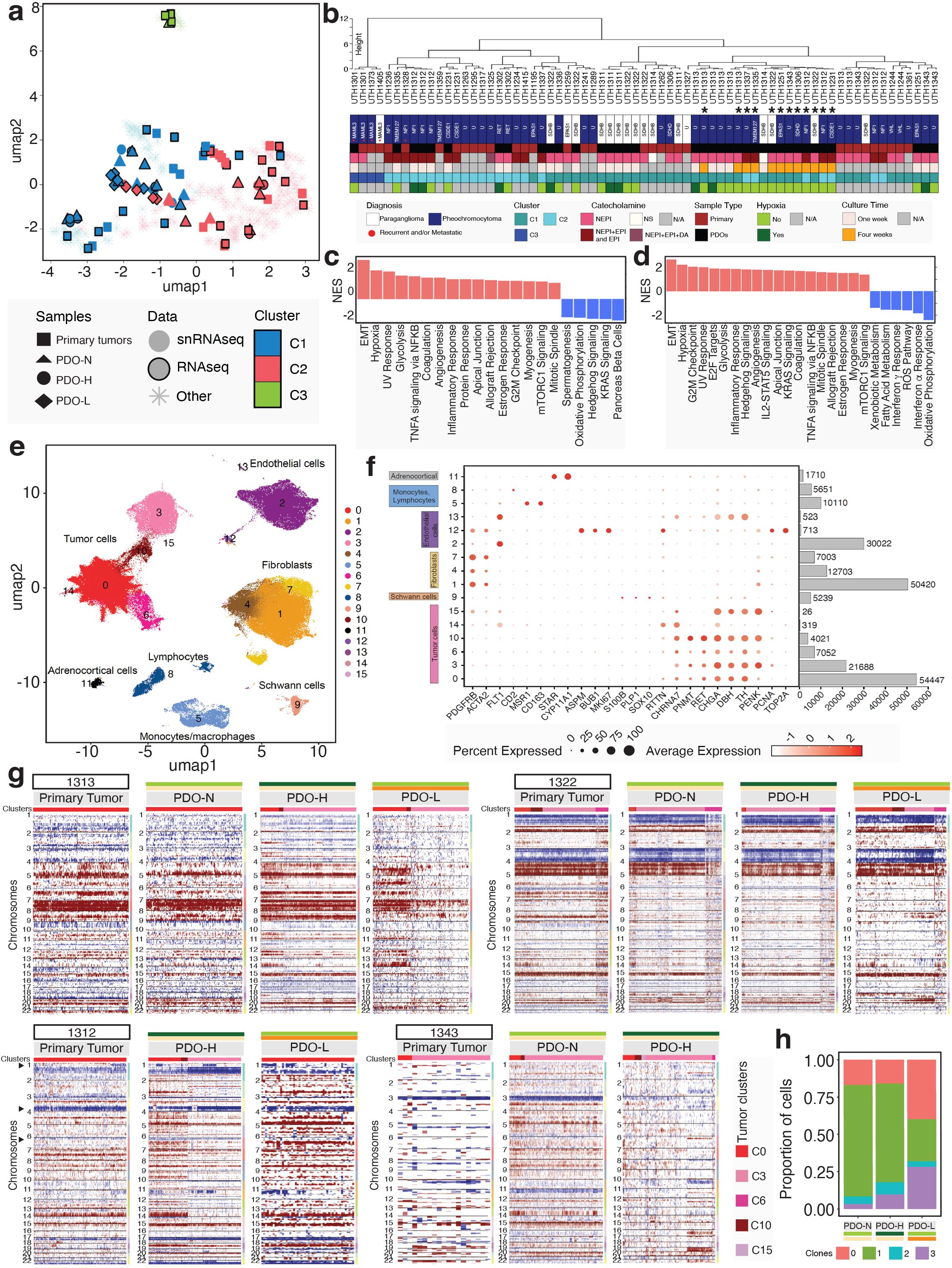
Short-term PDOs align with parental molecular states, while long-term PDOs acquire a pseudo-hypoxic profile. a) Uniform Manifold Approximation and Projection (UMAP) plot of bulk RNAseq and pseudobulk (all cells from single cell or single nuclei RNA-seq from 83 samples (41 primary tumors and 42 PDOs), their respective parental tumors, and other PPGLs from our dataset, and public databases (TCGA), sample type, molecular clusters and PDO conditions are labeled as indicated, Other represents publicly available samples shown in the clustering background); b) Dendrogram of samples shown in (a) depicting close alignment of short-term PDOs cultured at normoxia (PDO-N) and hypoxia (PDO-H) with their respective parental tumors, and a cluster formed by long term PDOs (PDO-L) of various genotypes (asterisks); c) Gene set enrichment analysis (GSEA) pathway enrichment analysis of the comparison between short term and long term PDO bulk RNAseq data ordered by significant NES scores; d) GSEA pathway enrichment analysis of the comparison between short term and long term PDO using the tumor cell matrix of deconvoluted RNAseq data ordered by significant NES scores; e) UMAP plot of sc/snRNAseq data from 30 samples including 13 PDOs and 17 parent tumors showing 16 cell clusters as indicated; f) Dot plot showing average expression and percentage of cells expressing representative markers of the 16 clusters shown in (e), bars indicate the number of cells in each cluster; g) inferCNV plots of four primary tumors and derived PDOs showing areas of predicted copy gain (red) or loss (blue) of tumor cells (columns) displaying their respective cluster (header) along their chromosomal location (rows), PDO culture conditions are shown: short-term normoxia (PDO-N), short-term hypoxia (PDO-H) and long-term normoxia (PDO-L); h) Proportion of cells in the four re-clustered tumor clusters in each PDO condition (PDO-N, H and L) as indicated (related to Suppl Fig 5e and 5f).

PDO-N generally aligned closely with their matched parent tumors, when available, or with tumors with related genotypes across the three main PPGL molecular groups, with few exceptions, in agreement with the cultures retaining the main molecular attributes of the primary tumors (**Figure 4b**). Despite a clear induction of a hypoxia transcription response in PDO-H cultures, as shown by the enrichment of hypoxia-related pathways (**Figure 1f** and **Supplementary Tables 2 and 3**), PDO-H samples maintained their original molecular group identity and similarity to their normoxic counterpart in all 6 normoxia-hypoxia pairs evaluated by RNAseq (**Figure 4b**). For example, normoxic/hypoxic PDO pairs from a RET-mutant (UTH1302) or NF1-mutant pheochromocytoma (UTH1312) retained their kinase-type identity on UMAP, and so did two normoxic/hypoxic PDOs derived from SDHB-(UTH1311 and UTH1322) related PPGLs. Two other pairs without a defined genotype (UTH1313 and UTH1343), also clustered with the expected profile (**Figure 4b**). One of the SDHB-derived PDOs, UTH1311, was also cultured at 5% O_2_ and this sample formed a tight cluster with the corresponding normoxic and 1% hypoxia-cultured PDOs (**Figure 4b**). These findings suggest that the exposure to low oxygen for one week preserves the main molecular identity and driver genotypes of PDOs.

In contrast to our findings on PDOs developed in hypoxia, all long-term cultures (PDO-L) samples formed a single cluster within the pseudohypoxic group, regardless of the molecular type of the parent tumor and despite being cultured under normoxic conditions (**Figures 4a-b**). For instance, PDO-L from NF1-(UTH1312) and TMEM127-derived (UTH1335) PPGLs clustered away from their matched short-term PDOs that aligned with the kinase cluster (**Figure 4b**). This distribution suggested that the PDO-L drifted transcriptionally over time in culture and acquired properties that were shared by other PDO-L samples independent of their genotype, and which resembled signatures typical of pseudohypoxic tumors. Accordingly, pathway analysis of PDO-L relative to short term cultures showed increased enrichment of epithelial mesenchymal transition and extracellular matrix (ECM) deposition (**Figure 4c** and **Supplementary Table 6-7**), a property previously reported in tumors with SDH mutation^51,52^ or SDH deficiency and inflammatory pathways^53^.

Activation of hypoxia, ECM and inflammatory signals collectively suggest a state of cellular stress, reminiscent of that reported in other long-term tumor PDOs^54,55^. Pseudohypoxic PPGLs characteristically are more angiogenic and have higher stromal component compared with other PPGLs^3,50,56^, so the alignment of long-term samples with this molecular group might be related to the dominance of nontumoral cells in these samples. To examine this feature, we performed in silico deconvolution of the bulk RNAseq with the MuSiC program^57^, and applied a previously published snRNAseq dataset of PPGLs as reference for PPGL cell subtypes^50^ to assess stromal content. Indeed, tumor cell proportions had significantly lower scores in PDO-L samples when compared to PDO-N or PDO-H, and conversely, the stromal scores and proportions were significantly higher in PDO-L relative to short-term cultures and the matched parental tumors (**Supplementary Figure 3b** and **Supplementary Table 8**).

To determine whether the transcriptional differences in PDO-L were driven by changes in tumor cell composition, we re-analyzed the data using an expression matrix restricted to tumor cells, excluding stromal and immune cell clusters (see **Methods**). The new UMAP alignment revealed similar results to the bulk RNAseq data (**Supplementary Figure 3c**). Importantly, pathway analysis of the tumor-only component (**Figure 4d**) produced results highly concordant with the bulk RNAseq (**Figure 4c**), with PDO-L cultured under normoxic conditions exhibiting significant enrichment of hypoxia, EMT and ECM-related pathways. These findings suggest that while short term PDOs strongly preserve molecular properties of the primary PPGLs, regardless of the levels of oxygen in the cultures, PDO-L diverge toward a pseudohypoxic profile strongly driven by the tumor cell compartment, with contribution from stromal cells.

### Single-cell profiling reveals preserved genomic structure and divergent hypoxia programs in PPGL PDOs

To gain further insight into the mechanisms underlying the dynamic changes of PPGL PDO in culture, we sequenced the transcriptome of 334,737 cells from 30 samples (average of 11,158 cells per sample, range 2,090-43,811; **Supplementary Table 9**). The dataset included 13 PDOs and 17 primary tumors, five of which we had previously reported^41^ (**Supplementary Figure 3d**). An initial batch of PDOs was processed as single nuclei to the handling of frozen primary tumors (**Supplementary Table 9**), but low yields prompted a switch to a single-cell protocol (see **Methods**). After we applied filters to remove mitochondrial and ribosomal RNA, doublets removal, and ambient RNA correction (see **Methods** and **Supplementary Figures 3e-f**), we retained a total of 211,647 transcriptomes for analysis (**Figure 4e**). We batch-corrected and integrated the data from all samples (**Supplementary Figure 3g**), performed UMAP clustering, and annotated the resulting cell clusters using various reference programs and manual annotation as described^41^ (**Supplementary Figure 4a-b**). We identified 16 cell clusters, including chromaffin marker-positive cells, Schwann-type cells, adrenocortical cells, fibroblasts, endothelial cells, lymphocytes and monocytes/macrophages (**Figure 4e-f** and **Supplementary Table 10**).

We confirmed the identity of the tumor cells using the copy number variation (CNV) inference with the inferCNV program, applying stromal and immune cells as the normal tissue reference, as we reported^41^. We identified the expected CNVs characteristic of parental PPGLs with established driver events, such as chromosome 1p loss in SDHB-mutant samples, or chromosome 3p/11p loss in a VHL-related PPGL (**Supplementary Figure 5a**) and verified the robustness of the outcome of inferCNV in tumor and derived PDOs regardless of the tissue source or dissociation protocol (**Supplementary Figure 5b**). Overall, we found that inferCNV analyses of the 13 PDOs derived from 5 PPGLs showed high concordance between the parental tumor and the PDO profile in all PDO conditions, including PDO-L (**Figure 4g**). We observed heterogeneity in primary tumor CNV profile from sample UTH1312, and this variability was maintained in both the hypoxia and long term PDOs (**Figure 4g**, arrow heads). This sample showed incongruent clustering in UMAP analyses at both bulk and pseudobulk levels (**Figure 4c** and **Supplementary Figure 3b**), suggesting that the observed transcriptional discrepancies may reflect underlying intratumoral heterogeneity. Overall, the data suggests that PDOs preserved the CNV profiles of their parental tumors across culture conditions, including intratumoral heterogeneity, implying that the transcriptional and biochemical changes observed in PDO-H and PDO-L are not driven by structural variation or clonal selection.

Focusing on the analysis of tumor cells, we observed that most samples contributed to the dominant clusters, and the smaller clusters were derived from a few samples (**Supplementary Figure 5c**). Differences in sample processing protocols introduced specific biases in cluster distribution, with cells arising from snRNAseq and scRNAseq enriching separate clusters (**Supplementary Figure 5c**). As this bias was present in primary tumors and in PDOs, suggesting that it was not an artifact of the ex-vivo cultures, we re-clustered the tumor cells, which yielded four tumor subclusters with contributions from both processing methods (**Figure 4h, Supplementary Figure 5d-e** and **Supplementary Table 11**). With this approach, the new tumor cluster 0 was predominantly composed of primary tumors both from frozen and dissociated tissue, whereas the new tumor cluster 1 was the most abundant in PDO-N and PDO-H (**Figure 4h** and **Supplementary Figures 5d-e**). The tumor cluster 3, small in PDO-N, expanded in PDO-H and more markedly in PDO-L (**Figure 4h** and **Supplementary Figures 5d-e**). Among nontumor cells, there was an expansion of fibroblast subpopulations in PDO-H and PDO-L (**Supplementary Figure 5f**). We used the re-clustered tumor cells for all subsequent analyses.

We previously found that both PDO-H and PDO-L had increased hypoxia signatures relative to PDO-N (**Supplementary Tables 3 and 7**). To identify potential distinctions between these two transcriptional profiles in our samples, we curated non-overlapping signatures of HIF1α (HIF1A)- and HIF2α (HIF2A)-specific target gene sets^58–60^ (**Supplementary Table 12**) and quantified hypoxia scores in the re-clustered tumor cells using AUCell^61^ (**Figure 5a-c**). We found that among primary tumors HIF2α emerged as the dominant signature, as expected based on reported PPGL data^3,60^, while PDOs expressed both HIF1α and HIF2α signatures (**Figures 5a-b**). When stratified by PDO condition, we observed that PDO-H had the highest HIF1α scores and HIF2α predominated in PDO-L (**Figure 5c**). We also detected pimonidazole expression in tumor cells (TH-positive) of a UBTF::MAML3 fusion-positive PDO-L culture (UTH1432), demonstrating intracellular hypoxia (**Figure 5d**). These findings validate the broad impact of environmental hypoxia in eliciting both HIF1α and HIF2α-driven transcription in tumor cells in the cultures maintained at 1% O_2_. HIF2α remains dominant in tumor cells grown over extended culture times (PDO-L), underscoring the importance of HIF2α for tumor cell viability ex vivo.

**Figure 5.**
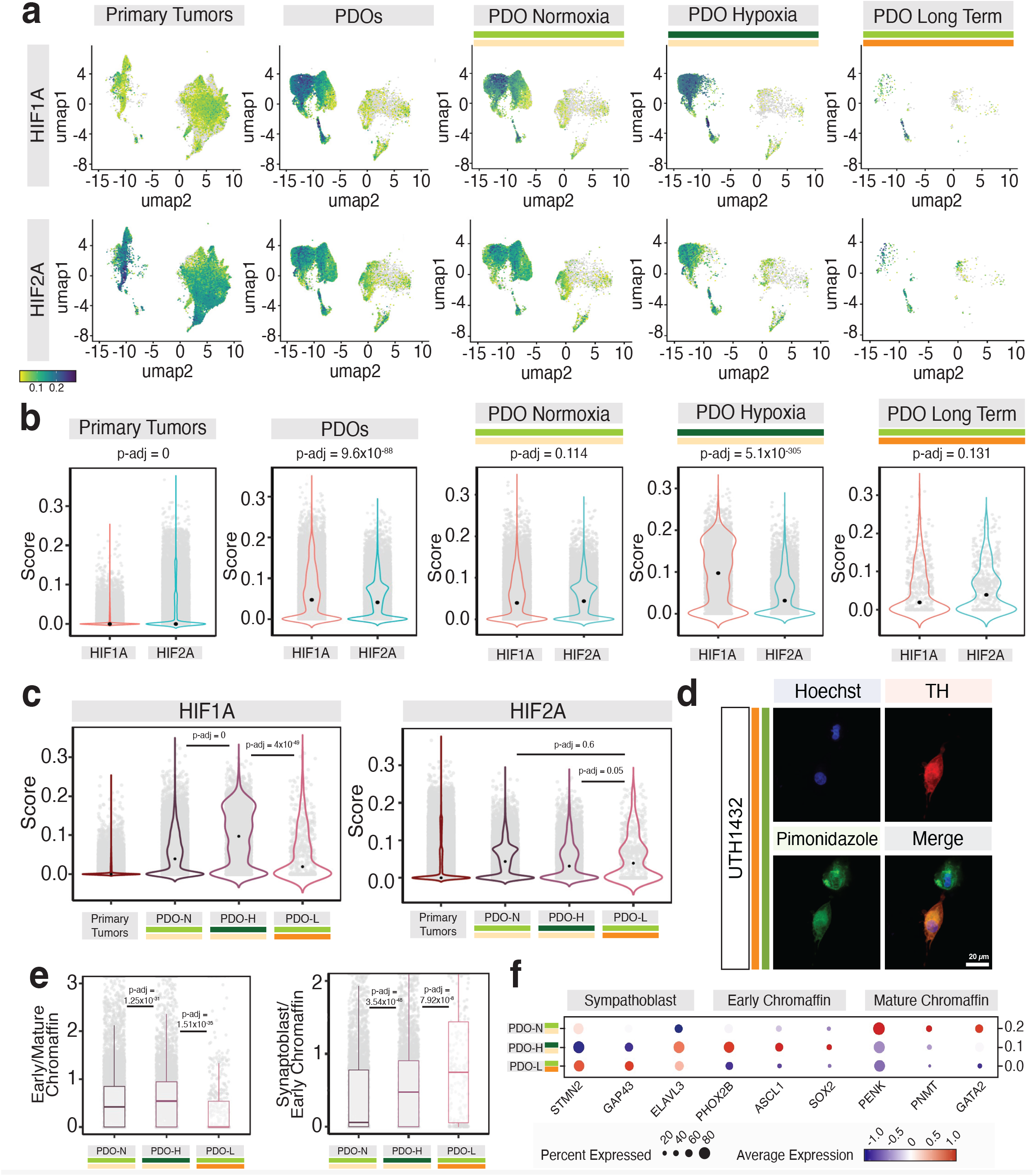
Single-cell analysis reveals differential engagement of hypoxia programs and chromaffin lineage states in PDOs. a) AUCell scores of HIF1α (top) versus HIF2α (bottom) signatures in primary tumors and PDOs (combined) or split in the three culture condition; b) Violin plots displaying comparisons between HIF1α and HIF2α for each of the samples shown in (a); c) AUCell scores of HIF1α and HIF2α signatures across primary tumors and PDO conditions; d) Immunofluorescence of TH (red) and pimonidazole (green) in a representative PDO-L sample, cultured at normoxia; e) Ratios of early/mature chromaffin markers (left) and ratio of sympathoblast/ early chromaffin marker (right) of AUCell scores of individual cells in PDO-N, PDO-H and PDO-L samples, p-values were calculated using Wilcoxon test in R (related to Suppl Fig 6a); f) Dot plot (average expression and percentage of cells expressing) of representative markers of sympathoblasts, early and late chromaffin cells in the three PDO conditions.

### Hypoxia and long-term culture shift PPGL PDOs toward less mature chromaffin states

To evaluate whether HIF1α and/or HIF2α activation influenced the tumor cell differentiation process, we generated consensus and non-overlapping signatures of chromaffin developmental subtypes based on multiple sources^62–67^ (**Supplementary Table 13**) and applied AUCell scoring to evaluate the primary tumors (both originating from frozen and dissociated samples) and PDOs at various conditions.

We found that the most abundant cell subtypes were chromaffin markers of early (less mature) and late (mature) stages (**Supplementary Figure 6a**). PDO-N also showed predominant mature markers, while PDO-H samples showed a higher proportion of immature cells, including ASCL1+ bridge cells and sympathoblasts relative to more mature chromaffin stages (**Figure 5e-f**). PDO-L had a more marked shift toward sympathoblast marker expression (**Figure 5e-f**). Most tumor clusters in primary tumors expressed markers of late chromaffin cells (**Supplementary Figure 6a**), regardless of their anatomic location since these samples originate from both pheochromocytomas (adrenal medulla) and paragangliomas (extra-adrenal paraganglia). Only a small fraction of cells in primary tumors cells expressed Schwann cell precursor (SCP) or bridge markers, whereas in PDOs these markers were detected at higher levels and in distinct subpopulations, primarily within tumor cell cluster 1 and most of tumor cell cluster 3 (**Supplementary Figure 6a**). The data indicates that PPGL organoids retain a degree of plasticity in chromaffin developmental programs, with perturbations such as hypoxia and prolonged culture associated with a shift toward earlier differentiation stages and enrichment for sympathoblast features, particularly in a subset of PDO-L tumor cells (**Figure 5f**).

### Developmental trajectory mapping reveals culture-dependent timing of energy-related and hypoxia-associated programs

PPGL PDOs derived from both pheochromocytomas (adrenal medulla) and paragangliomas (extra-adrenal paraganglia) exhibited heterogeneous tumor cell subpopulations spanning early chromaffin, mature chromaffin, and sympathoblast-like states (**Figure 5a** and **Supplementary Figure 6a**). This distribution aligns with a model in which sympathoblasts can develop from chromaffin cells^68^ and with evidence that early Schwann precursor cells are primed toward alternative fates in the Schwann and sympathoadrenal lineages, implying microheterogeneity along the differentiation continuum^62,68^. Our findings suggest that a subset of PPGL tumor cells retains plasticity to interconvert between subtypes at transition points within the chromaffin lineage, which is promoted by changes in culture conditions (hypoxia, longterm growth).

To gain additional insights into the differentiation dynamics in culture, we next estimated tumor cells’ developmental trajectories by determining cellular differentiation states and reconstructing cell differentiation lineages of the tumor clusters using Cytotrace^69^. The new (reclustered) tumor cluster 1 appeared as the earliest cell group in the differentiation trajectory relative to the other tumor subclusters both in primary samples (**Figure 6a**) and in PDOs (**Figure 6b**). Therefore, we designated cluster 1 as the ‘root’ cell population and reanalyzed the trajectories using Monocle 3^70^ to assess differentiation patterns across PDO conditions. We found conserved trajectories across the three conditions normoxia, hypoxia and longterm culture (**Supplementary Figures 6b-c**). We next generated a heatmap of the Monocle 3 analysis to examine the genes underlying the tumor cell trajectory and found extensive overlap in the gene programs that were enriched in the early tumor cells of both primary tumors and PDOs (**Supplementary Figure 6d**). Genes involved in mitochondrial respiration and protein translation, previously reported in primary PPGLs^38^, accounted for a large component of differentially expressed gene (DEG) sets in PDOs (**Supplementary Table 14**). Both the magnitude and temporal pattern of expression varied by culture condition, indicating that neoplastic chromaffin cells engage these energy-related programs in a context-dependent manner. In PDO-H, these signatures appeared earlier and persisted longer along the tumor cell trajectory, whereas in PDO-L, they emerged later (**Supplementary Figure 6d**). Other differentially expressed genes (DEGs) in these analyses were RGS4 and RGS5 (**Supplementary Table 14**), which are implicated in the hypoxia response and oxygen sensitivity of the adrenal medulla and carotid body^71^. In addition, DLK1^72^ and NTRK1^73^ (**Supplementary Figure 6d**), genes that are epigenetically regulated by EZH2 in neuroblastoma and have been associated with sympathoblast lineage development and neuroblastoma survival^73,74^ were also differentially represented along the PDOs trajectory.

**Figure 6.**
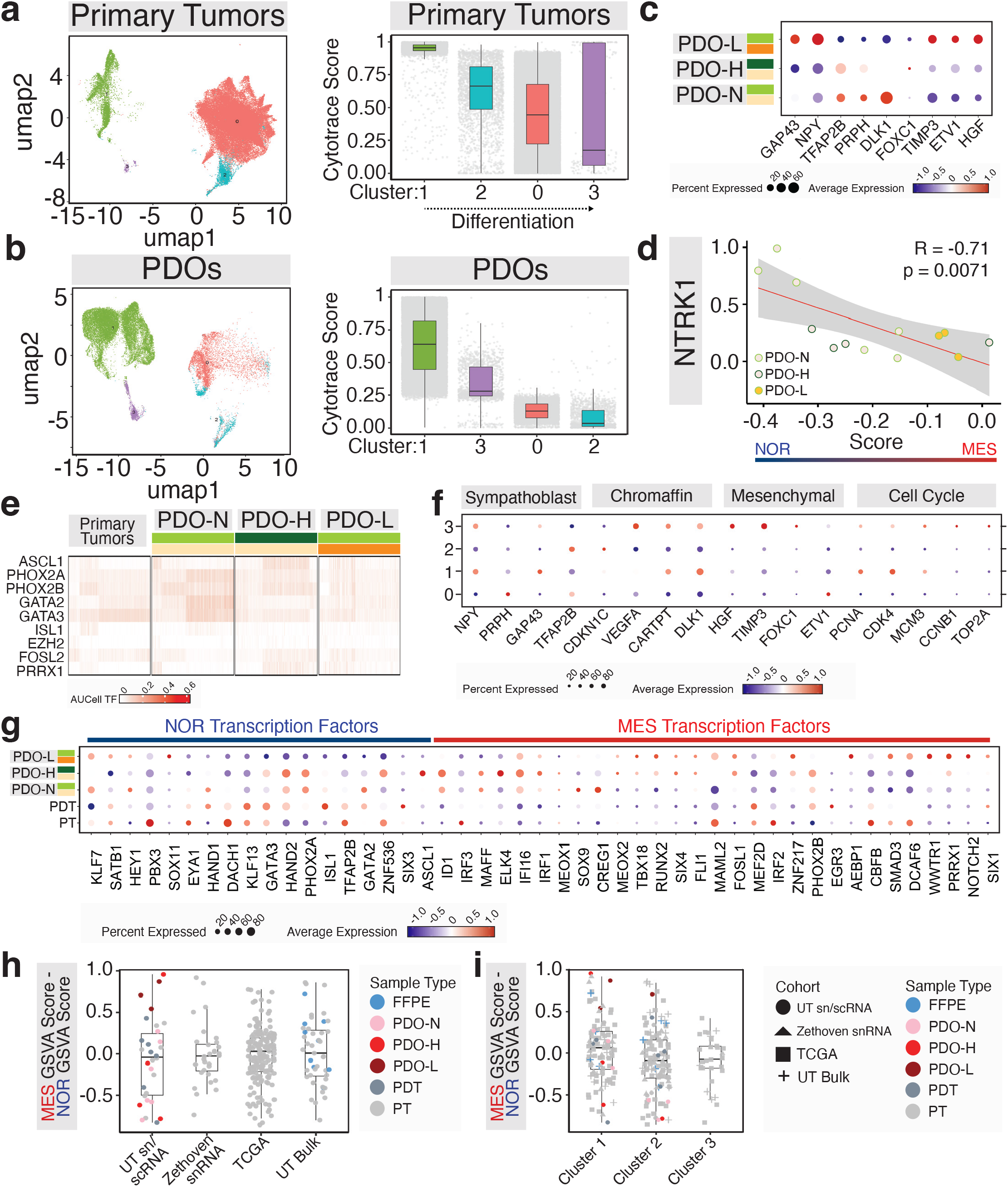
Long-term PPGL PDOs acquire a mesenchymal-like transcriptional state distinct from primary tumors. a) Estimated differentiation state of tumor cell clusters of primary tumor samples using Cytotrace plotted in the order of least to most differentiated cell cluster (right), corresponding tumor cluster UMAP is shown on the left; b) Estimated differentiation state of tumor cell clusters in PDOs using Cytotrace plotted in the order of least to most differentiated cell cluster (right), corresponding tumor cluster UMAP is shown on the left; c) Dot plot (average expression and percentage of cells expressing) of sympathoblasts and mesenchymal markers in the three PDO conditions, N, H and L; d) Correlation between the NOR-MES differential score (see Methods) and the NTRK1 gene expression in PDO N, H and L; e) pySCENIC analysis of selected NOR-MES transcription factors across primary tumors and the three PDO conditions, scale bar is shown below; f) Dot plot of the expression of selected markers of sympathoblast, chromaffin, mesenchymal and cell cycle in PDOs across individual tumor cell clusters 0-3 (Related to Suppl Fig 7b); g) Dot plot of the NOR-MES transcription factors that are associated with superenhancers defining NOR and MES distribution in neuroblastomas primary tumors (PT and PDT-frozen and dissociated tumor samples) and the three PDO conditions, N, H and L-all dot plot graphs display average expression and percentage of cells expressing the indicated markers (Related to Suppl Fig 7c); h) Plots of the MES-NOR differential gsva scores of four sample cohorts: primary tumors and PDOs based tumor cells from sc/sn RNAseq data of the present study (UT sc/snRNAseq, n=30), snRNAseq from a published PPGL cohort (Zethoven et al n=36), deconvoluted tumor matrix from bulk RNAseq from the TCGA PPGL cohort (n=183), and deconvoluted tumor matrix from bulk RNAseq from a separate cohort of UT PPGL samples (n=47), sample/processing type is indicated (FFPE-forma-lin-fixed paraffin-embedded); i) Plots of MES-NOR differential gsva scores of samples shown in (h) displayed by their molecular clusters

### Noradrenergic–mesenchymal plasticity in PPGL PDOs and primary tumors resembles neuroblastoma

We had previously found that epithelial mesenchymal transition (EMT) was the top-ranked pathway among PDO-L samples when compared with PDO-N (**Figure 4c-d** and **Supplementary Table 7**). This mesenchymal enrichment, together with the unsupervised trajectory analysis of PDOs and the observed maturation shifts toward earlier chromaffin states in PDO-H or sympathoblast-like states in PDO-L, suggested that the divergence of PDO-L from PDO-N resembles the heterogeneity seen in neuroblastoma, a tumor type that also recapitulates the adrenal developmental trajectory^63,65,66^. In neuroblastoma, superenhancer modulation underlies two main identities: a noradrenergic state (NOR, also referred to as ADRN-like) and a mesenchymal state (MES, also termed neural crest cell– or NCC-like)^75,76^. These neuroblastoma identities display substantial plasticity, with the capacity to interconvert between noradrenergic and mesenchymal phenotypes in response to tumor-intrinsic signals or microenvironmental cues^76–79^.

We thus assessed whether markers associated with NOR and MES states were also detected in the PDO samples and/or could be influenced by their culture conditions^75^. First, we quantified the expression of NOR-defining genes (TFAP2B, PRPH, GAP43, NPY) and MES-defining genes (FOXC1, TIMP3, ETV1, HGF, EZH2, SOX11) in tumor cells from our sc/snRNAseq cohort. PDO-N exhibited a NOR-enriched profile and PDO-L had a striking resemblance to the MES profile, while PDO-H showed generally an intermediate expression pattern (**Figure 6c**). We used a more comprehensive, previously curated gene list^76^ to estimate the MES-NOR score in these samples, and observed similar distribution^76^(**Figure 6d**). Because this analysis was restricted to tumor cell clusters, the observed profiles could not be attributed to stromal contamination. These results suggest that PDOs may undergo a noradrenergic-to-mesenchymal transition (NMT) under specific culture conditions.

To assess the regulatory network of PDOs and estimate potential overlaps between them and those underlying neuroblastoma superenhancer regulation, we applied pySCENIC^80^ to our samples. Sixty-one predicted regulons obtained with this unsupervised analysis included several transcription factors expected to be involved in PPGLs (**Supplementary Figure 7a** and **Supplementary Table 15**), such as those related to the chromaffin/noradrenergic lineage (ASCL1, GATA2/3, PHOX2A/PHOX2B, etc.) and others that we have previously identified in association with kinase-type PPGLs (e.g. EGR1)^41^. In addition, EPAS1, the gene coding for HIF2α, was among the significant transcription factors with highest scores in PDO-H and PDO-L tumor cells (**Supplementary Figure 7a**), in alignment with our observations above (**Figures 5a-c**), and underscoring its relevance for PPGLs.

Notably, several transcription factors linked to the neuroblastoma MES identity were also amongst the enriched regulons (**Figure 6e**). To explore these findings, we examined the expression levels of transcription factors previously identified to drive the NOR-MES identities^75,78,79^ in our samples and found enrichment of the NOR transcription factor profile in PDO-N and high expression of MES-related transcription factors in PDO-L, although some NOR transcription factors were also expressed in this group (**Supplementary Figure 7b**). PDO-H cultures exhibited an intermediate pattern between PDO-L and PDO-N (**Supplementary Figure 7b**), in agreement with our observation of selected NOR-MES markers shown in **Figure 6c**.

To establish which tumor cell subpopulations within these three conditions carried the features associated with the ‘NOR-MES’ transition, we evaluated the expression of these genes across tumor clusters. Tumor cluster 3 had the highest expression of MES markers (**Figure 6f**) and a subset of cells in this cluster also expressed proliferation markers (**Figure 6f**). We used additional stringent filtering methods to confirm that these cells preserved tumor identity and were not stromal cell- or clustering-derived artifacts (see **Methods**). These findings suggest that selected tumor subpopulations in PPGL PDOs possess both plasticity and proliferative capacity that may confer unique biological properties to tumor cells and render them amenable to modulation.

To place these findings in the context of primary samples, we extended the NOR/MES transcription factor analyses to tumor cells of our sc/ snRNAseq PPGL cohort relative to PDOs. NOR markers were predominantly expressed in the primary PPGLs, as expected (**Figure 6g**). However, mesenchymal markers were also expressed in these primary tumors, though to a lesser extent than in PDO-L samples (**Figure 6g**). Cluster-specific analysis of tumor cells revealed a distribution similar to that of PDOs, with MES-related markers most prominently expressed in tumor cluster 3 (**Supplementary Figure 7c**). These results suggest that transitional states between NOR and MES profiles are likely present in primary tumors and not only in PDOs. To extend this observation, we calculated MES-NOR differential scores in four PPGL datasets: our sc/snRNAseq cohort, the deconvoluted tumor expression matrix from bulk RNAseq samples included in this study, an snRNAseq cohort from Zethoven^39^, and the TCGA primary PPGL bulk RNAseq dataset^3^ (**Supplementary Table 16**). In these cohorts, we observed a broad range of scores across PPGLs of various genotypes and molecular clusters (**Figure 6h**), and a trend toward enrichment in tumors belonging to the molecular pseudohypoxic group, although it did not reach statistical significance (**Figure 6i**). These findings suggest that, similar to neuroblastomas, plasticity between NOR-like and MES-like phenotypes may also occur in PPGLs and contribute to their phenotypic heterogeneity.

### Drug Sensitivity Profiling of Normoxic, Hypoxic, and Long-Term PPGL PDOs

PPGLs have few systemic treatment options, and their combined clinical and molecular heterogeneity, along with largely uncharacterized therapeutic sensitivities, highlights the need for functional models that can uncover genotype- and state-specific vulnerabilities. To address this, we evaluated drug responses in baseline PPGL PDOs and in matched cultures maintained under hypoxia (PDO-H) or prolonged growth (PDO-L) to determine how these conditions influence sensitivity profiles.

We screened 50 PPGL patient-derived organoid (PDO) plates generated from 25 patients in our automated mini-ring platform^31–34^. Forty-seven plates passed quality control and were included in the analysis. Most represented short-term normoxic cultures (PDO-N, n=36; **Figure 7a**), a subset of which was also evaluated under hypoxic conditions (PDO-H, n=11; **Figure 7a**) or after prolonged culture (PDO-L, n=10). Across all conditions, the number of compounds and combinations tested per sample ranged from 8 to 51, with a mean of 30 and median of 31. Depending on sample availability, we screened either at a single concentration in discovery mode or at multiple concentrations spanning three orders of magnitude (**Figure 7a– b**).

**Figure 7.**
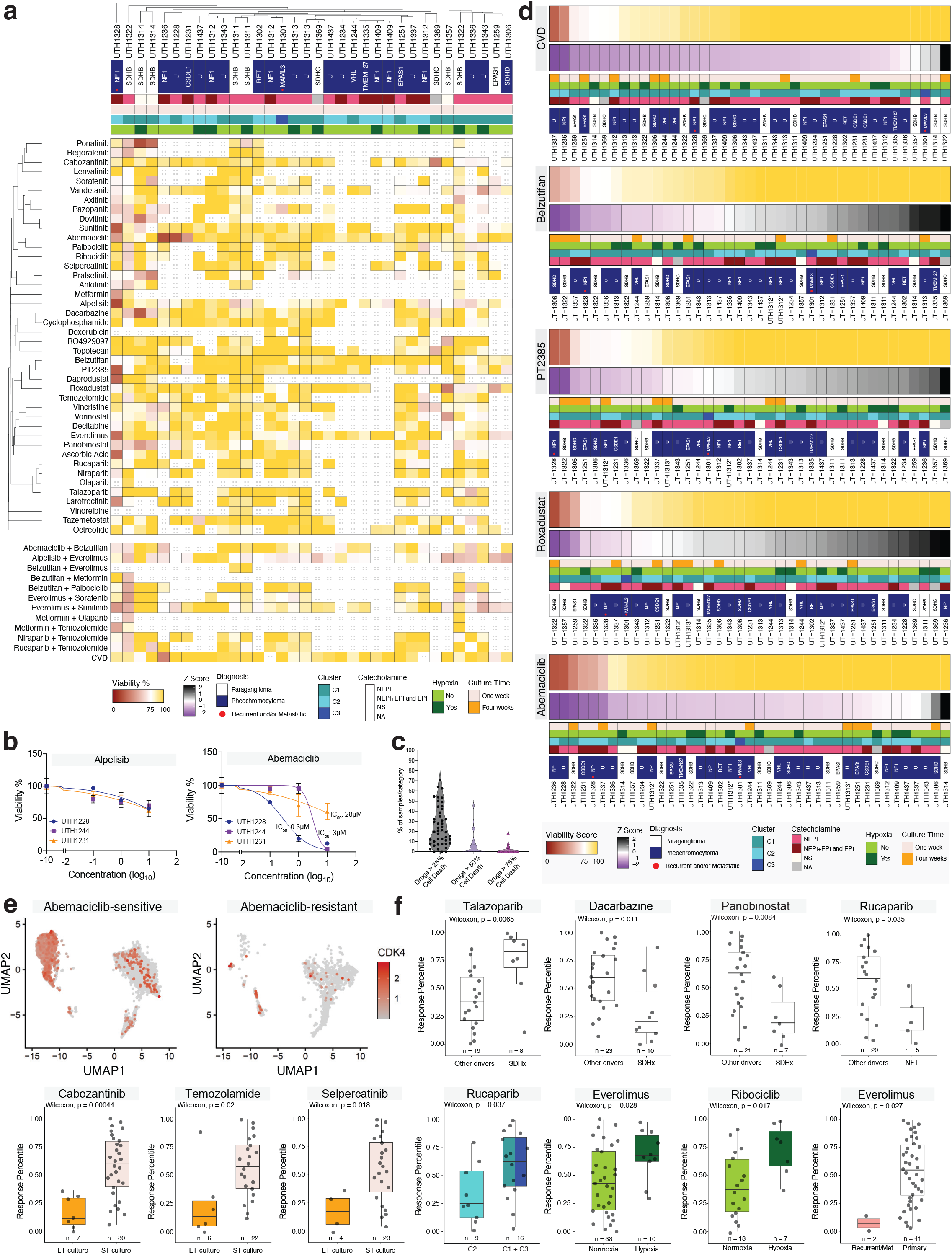
PPGL PDOs display genotype- and context-dependent drug sensitivities across therapeutic classes. a) Heatmaps of PDO viability to the indicated drugs (or drug combinations) at 1 µM. Each column is a unique PDO sample, red indicates high cell death in response to treatment. Colored bars underneath each heatmap represent the Z score, diagnosis, molecular cluster, PDO condition and catecholamine profile, Norepinephrine (NEPI), Epinephrine (EPI and EPI+NEPI), Nonsecreting NS; b) Dose-response curves of three exemplary PDOs to two distinct drugs, Alpelisib and Abemaciclib and respective half-maximal inhibitory concentration (IC50), data are presented as mean +/-SEM; c) Percentage of PDOs showing the indicated level of drug response based on % cell death (>25%, >50% and >75%), showing individual samples, median and the interquartile range; d) Heatmaps of PDO sensitivity to individual drugs or combinations as indicated, displayed as in (a). The PDO viability score represents that sample’s viability normalized to the mean response to treatment across all samples. Samples are ranked from low residual viability percentile (most responsive samples) to the highest residual viability percentile (least responsive samples); e) UMAP plot displaying CDK4 expression on tumor cells from responsive and resistant PDOs and respective primary tumor samples. f) Sensitivity rank plots comparing the response of the specified PDO groups to the indicated drugs: pheochromocytoma (pheo) vs paragangliomas (pgl), short vs long term culture, hypoxia vs normoxia, primary vs recurrent, SDH-mutant vs WT, NF1 mutant vs WT. Boxplots represent the interquartile range.

We used a customizable drug library including agents previously used in PPGLs (cyclophosphamide, vincristine, and dacarbazine-CVD-combination chemotherapy, sunitinib, selpercatinib, pralsetinib, and cabozantinib)^81,82^, additional multi-kinase inhibitors, and other classes of FDA-approved drugs, including PARP inhibitors, HDAC inhibitors, and CDK4/6 inhibitors (**Supplementary Table 17**). Given the relevance of the HIF2α pathway activation in PPGLs, particularly those with pseudohypoxic molecular features^5^, belzutifan, a HIF2α-selective inhibitor^83,84^ recently approved by the FDA for treatment of locally advanced, unresectable, or metastatic PPGLs^85^, was also included. We also evaluated an earlier generation HIF2α inhibitor, PT-2385^59,86^, and roxadustat^87^, a HIF prolyl hydroxylase inhibitor, which prevents HIF2α inactivation.

At the 1 µM screening dose, 3 PDOs (6.8%) showed no response to any tested drug, defined as <25% cell death for all compounds (**Figure 7c** and **Supplementary Table 18**). Nearly half of PDOs (40.9%) had at least one drug producing >50% cell death, and 15.9% had at least one drug producing >75% cell death. Two PDOs (4.5%) were hypersensitive, displaying ≥75% cell death in about a fifth of all drugs tested (**Figure 7c**). Notably, both hypersensitive PDOs were PDO-L, potentially reflecting higher proportions of rapidly cycling non-tumor cells. Non-responders were tested with fewer drugs (median 26 drugs) compared with responders (median 33 drugs), suggesting that limited screening may contribute to the inability to identify positive hits, in line with our previous observations in sarcoma^32^. Neither PDO-H nor PDO-L cultures showed statistically significant overrepresentation in any drug response category by Fisher’s exact test. PDO-N and PDO-H generally exhibited concordant drug responses (**Figure 7a**).

For each drug, we normalized the viability of a given sample to the mean viability observed across all PDOs treated with that same drug or combination to calculate viability scores^32^. This normalization mitigates bias from compounds with broad cytotoxic activity and pinpoints samples with distinctive sensitivity profiles to specific agents (**Figure 7d**). For instance, the CVD combination has been used as standard of care in metastatic PPGLs^88–90^, with reports of potentially improved outcomes in patients harboring SDHB mutations^49,91^. CVD was active on 17% of tested PPGL PDOs tested, a rate consistent with the proportion reported in patients (22%)^92^. In the present cohort, only one of the SDHB-derived PDOs showed a mild response to CVD (**Figure 7d**).

### Therapeutic Modulation of HIF2α Pathway in PPGL PDOs

Since mutation-driven, constitutive activation of HIF2α is a frequent pathogenic event in PPGL, closely linked to tumor development and progression^5^, and given the recent approval of belzutifan, a selective HIF2α inhibitor, for advanced, inoperable and/or metastatic PPGLs (LITESPARK15 trial, NCT04924075), we examined the effect of low doses of belzutifan in 41 PDO models established from n=24 patients (**Figure 7d**). We also tested the earlier generation HIF2α inhibitor, PT2385 (42 PDOs from n=23 patients, **Figure 7d**). Both compounds demonstrated activity in a subset of samples, with belzutifan yielding a viability score < 75 in 6/41 samples, including 3 SDH-related samples and one NF1-driven PPGL with metastatic disease (**Figure 7d**). Similarly, PT2385 showed comparable activity in 8/42 samples, including 3 SDH-related PDOs from two patients and 3 NF1-mutant samples from two patients, one with metastatic disease (**Figure 7d**). Lower level belzutifan activity was detected in PDOs with additional pseudohypoxic molecular types, including samples with activating mutations of EPAS1, the gene encoding HIF2α, a subgroup predicted to derive the greatest benefit from HIF2α inhibition^5,93^. Although modest in magnitude, the responses to belzutifan (15%) are relatively close to the reported overall response rate of 26% that supported FDA approval for advanced PPGLs ^85^. Notably, given that the 26% response rate was observed in advanced cases, and that one of two metastatic PDOs in our cohort exhibited measurable response, these findings highlight the potential of this platform to identify advanced cases more likely to benefit from HIF2α inhibition.

The combination of belzutifan with palbociclib, an FDA-approved CDK4/6 inhibitor, is under investigation in VHL-deficient renal cell carcinoma (NCT05468697)^94^. In renal cancers, VHL loss leads to elevated HIF2α, which upregulates CCND1, the cyclin partner of CDK4/6 that drives cell cycle progression^95^. Given that HIF2α is frequently elevated in PPGLs beyond those VHL-mutated, combining CDK4/6 inhibition with HIF2α targeting is mechanistically justified. We tested the combination of belzutifan with palbociclib in 23 PDOs, observing a modest increase in response rate (17%) compared with belzutifan alone (14%). The combination of belzutifan with another CDK4/6 inhibitor, abemaciclib, resulted in a statistically significant increase in activity in NF1-driven PPGLs compared to all other tested models (Wilcoxon, p = 0.036; **Supplementary Figure 8b**).

Interestingly, although considered a core feature of many PPGLs, genetic hyperactivation of the HIF2α beyond a certain threshold has been considered toxic to adrenomedullary cells^58^. To explore this in our model, we also treated PDO cultures with roxadustat, an agent that stabilizes both HIF1α and HIF2α by inhibiting HIF hydroxylases^58,87^, and found that the top responders all carried pseudohypoxic mutations (**Figure 7d**). These results suggest that in certain contexts, excessive HIF2α activity may impair cell viability. We therefore hypothesized that, conversely, partial inhibition of HIF2α might relieve this toxicity and facilitate growth in a subset of tumors. To test this possibility, we evaluated whether HIF inhibitors could promote PDO growth defined as viability scores >130, corresponding to >120% viability (**Supplementary Figure 8a**). For both drugs, 3/4 samples exhibiting growth were derived from pseudohypoxic tumors (**Supplementary Figure 8a**). Notably, the three samples showing pronounced cell death with roxadustat (UTH1259, UTH1236, UTH1357) also demonstrated the most robust growth with PT2385, suggesting complementary effects (**Figure 7d** and **Supplementary Figure 8a**). These findings indicate that further characterization of HIF2α modulation on PDO viability will be required to optimize the therapeutic potential of agents targeting this pathway.

### Sensitivity of a subset of PPGL PDOs to the CDK4/6 Inhibitor Abemaciclib

The CDK4/6 inhibitor abemaciclib produced one of the strongest responses across all drugs tested, with 6/44 samples exhibiting a viability score ≤ 55 (**Figure 7d**). In particular, a subset of PPGLs demonstrated strong responses to abemaciclib, even when compared to responses across n = 139 PDOs from 10 tumor types (**Supplementary Figure 8c**). Although CDK4/6 inhibitors have not previously been used in PPGL treatment, preclinical studies have reported their ability to promote differentiation and reduce proliferation of adrenomedullary cells^96,97^. Among the top responders were PDOs derived from two NF1-mutant cases, including a metastatic tumor (UTH1328); however, there was no statistically significant association between responsiveness and any molecular feature or genotype. In other cancers, response to CDK4/6 inhibitors has been reported to involve multiple mechanisms that may be both tumor type- and stromal tissue-dependent^95, 98, 99^. Therefore, identification of biomarkers of response and/or resistance has been challenging^95, 98, 99^. Notably, in our PPGL PDO drug screening, the abemaciclib response profile was not reproduced by other CDK4/6 inhibitors, palbociclib or ribociclib. Such variability in response among drugs within the same class has been reported previously in both clinical and pre-clinical contexts^32, 96, 100, 101^.

We next analyzed transcriptomics data to identify molecular features distinguishing the top responders from those at the bottom of the response curve (**Supplementary Figure 8c**). We compared the expression profiles of 4 responsive PDOs with those of 7 non-responsive PDOs, and we also evaluated the transcriptomes of the corresponding primary tumors (n = 4 responders vs n = 4 non-responders). Pathway analysis revealed that responsive PDOs (and their respective parental tumors) were enriched for signatures associated with cell proliferation, epithelial-to-mesenchymal transition and glycolysis, whereas resistant PDOs were enriched for pathways related to cytoskeletal organization and cell adhesion (**Supplementary Figure 8c** and **Supplementary Tables 19 and 20**). To further investigate the potential cellular targets of abemaciclib, we assessed CDK4 expression in our single-cell transcriptomic dataset (**Figure 7e**). We found that CDK4 showed higher expression in tumor cluster 1 in the responsive PDO/primary sample group, and in tumor cluster 3 of the resistant samples. Both of these clusters had exhibited the highest proliferative activity among PDO tumor cells (**Supplementary Figure 7c**), and cluster 3 demonstrated a combined sympathoblast and mesenchymal transcriptional program (**Figure 6f**), suggesting biological features potentially linked to drug sensitivity. Collectively, these findings identify a subset of PDOs sensitive to an FDA-approved agent not yet tested in this disease context, abemaciclib, with actionable therapeutic potential.

### Associations Between Molecular and Microenvironmental Features and Drug Response in PPGL PDOs

We next examined correlations between drug sensitivity and underlying molecular or microenvironmental features across the PDO cohort, focusing on both established clinical regimens and single agents. We identified several statistically significant trends within the current dataset. For example, although no genotype-specific association was observed for the CVD regimen, dacarbazine alone showed greater activity in SDH-mutated PPGLs compared to tumors with other drivers (Wilcoxon p = 0.011, **Figure 7f**). Dacarbazine is thought to be the active agent within CVD for neuroendocrine tumors, including PPGL^91,102^.

Panobinostat showed higher activity in SDH-mutated PPGL PDOs than in other genotypes (Wilcoxon p = 0.0084; **Figure 7f**). This aligns with mechanistic data showing that HDAC inhibition stabilizes mutant SDHB by limiting proteostasis-mediated degradation, supporting the concept of proteostatic vulnerability in some SDH mutant tumors^103^.

Everolimus showed higher activity in normoxia than in hypoxia across PPGL PDOs (Wilcoxon p = 0.028, **Figure 7f**). This pattern is consistent with hypoxia-driven suppression of mTORC1 activity due to HIF-induced REDD1 and TSC1/2, which has been linked to relative resistance to mTORC1 inhibitors in hypoxic tumor regions^104^. Clinical experience with everolimus in PPGL is limited but includes a phase 2 single agent study reporting modest activity, supporting its evaluation in this disease context in association with PDO-driven patient selection^105^.

Long-term PDOs converge on pseudo-hypoxic and mesenchymal programs with higher angiogenic signaling and proliferative activity, which can heighten dependence on receptor tyrosine kinases and vascular pathways. This is consistent with the observed stronger activity on PDO-L of cabozantinib (Wilcoxon p = 0.00044; **Figure 7f**), a multi-kinase inhibitor of VEGFR, c-MET, RET and c-KIT with documented clinical efficacy in progressive metastatic PPGLs^106^. We showed that PDO-L also preserve sympathoadrenal features, including RET expression, for the duration of the culture (**Supplementary Figure 1d**). This could justify activity of a highly selective RET inhibitor like selpercatinib in mechanistically distinct modes of RET activation across PDO-L (Wilcoxon p = 0.018; **Figure 7f**). These findings indicate that drug sensitivity in PPGL PDOs is influenced by molecular- and culture-associated phenotypes, revealing potentially targetable vulnerabilities in subsets of patients.

## Discussion

We developed a new, scalable model of PPGL PDOs that recapitulates multiple properties of primary PPGLs, including protein expression, biochemical/hormonal profile and transcription clustering. Organoids preserved cellular complexity and subpopulations at single-cell resolution, and they maintained copy number architecture consistent with matched tumors across conditions, indicating genomic stability during culture in different conditions.

Culturing in hypoxia preserved the tumor cell states detected in primary samples and favored HIF1α program engagement in tumor cells while maintaining HIF-2α activity at levels similar to primary tumors. Their similarity to the molecular profile of the primary tumors and higher viability suggest that PDOs cultured under hypoxia could recreate an environment that closely resembles primary tumors, albeit with a shift toward earlier chromaffin cells and catecholamine biosynthesis maturation. These results align with developmental and clinical observations, including HIF-2α dependence in sympathoadrenal development, enrichment of EPAS1 mutations in PPGL from patients with cyanotic congenital heart disease, and altitude-associated PPGL risk^6,15^. In line with hypoxia-responsive behavior reported in rodent pheochromocytoma cell lines^42^, these results directly anchor PPGL pathogenesis and growth to low-oxygen adaptation in human models.

In contrast, long term cultures retain fewer tumor cells that engage pseudohypoxic pathways with predominant HIF2α activation, regardless of their genotype. Our analyses suggest both tumor-intrinsic signals and extrinsic contribution from nontumor cells influence this shift of the long term cultures. In addition, PDO-L tumor cells increase the proportion of sympathoblast markers and contain subpopulations that carry properties associated with mesenchymal features reminiscent of the MES phenotype of neuroblastomas, contrasting with the predominant NOR (noradrenergic)-like profile of short term, normoxia-cultured organoids (PDO-N). Whether these changes are related to activation of HIF, and/or reflect responses to other environmental cues remains unknown. In neuroblastomas, plasticity between NOR-predominant or MES-predominant identities has been observed, and suggested to be an epigenetic regulated process, dependent on both tumor intrinsic and microenvironmental signals^75,76,78^. For example, certain growth factors, including EGF, TNFα or PRRX1 promoted an MES phenotype in vitro, while in vivo growth of xenografts tended to direct MES tumor cells toward NOR in neuroblastomas^78^. Tumor cell subpopulations which co-express noradrenergic and mesenchymal markers have been detected in primary neuroblastomas, suggesting that tumor cells exist in a transitional state between the two identities^75,76,78,79^. Our data suggest that the microenvironment of PPGL PDO culture, in alignment with previous observations of in vitro co-cultures^107^, supports plasticity of tumor cells. Importantly, we also detected cells co-expressing NOR/MES genes in primary PPGLs, suggesting that these transitional cell states may also occur in vivo and co-exist within individual tumors. MES phenotype has been associated with more aggressive tumor features and treatment resistance in neuroblastomas^78, 79^. Further work will be needed to define these cells, determine genetic and epigenetic drivers of these cell states and assess their functional properties, as well as the ability to redirect their states with defined perturbagens. Importantly, it will be critical to establish whether cells with potential capability to switch between chromaffin and mesenchymal identity within primary tumors reflect pre-existing subpopulation(s) with distinctive biological aggressiveness, spatially defined distribution, or other properties which may predict their progression and/or treatment response.

Our PPGL platform recapitulated responses to agents used clinically or under active evaluation. Belzutifan and PT2385, both targeting HIF2α, produced activity in subsets of models. Responses to belzutifan were within a range (>20%) similar to the reported clinical response that led to the recent approval of this drug in metastatic PPGLs. Lack or responses to belzutifan have been documented in prior spheroid PPGL culture models^82^, highlighting an advantage of organoids for modeling hypoxia-dependent biology. Extended evaluation of belzutifan in larger PDO cohorts and conditions may offer additional insights into properties of response and/or primary resistance.

One of the most striking single-agent drug responses in our cohort was observed against the CDK4/CDK6 inhibitor abemaciclib, which has not been previously explored for the treatment of metastatic or recurrent PPGLs. We did not observe clear association between the responsive or resistant tumors and specific genotypes; however, a PDO derived from an NF1 tumor that evolved with metastases was among the top responders, suggesting that this CDK4/6 inhibition deserves additional examination in aggressive PPGLs. Responsive PDOs showed differential expression of genes in the ECM pathway, which should be evaluated as markers in future, larger studies to independently validate this observation. Interestingly, in samples with scRNAseq data, expression of CDK4 was more abundant in tumor cell clusters containing ‘less mature’ markers, suggesting that that these subpopulations may be the cell targets of abemaciclib. Importantly, as we observed plasticity in tumor cells in response to culture conditions, it will be important to characterize in future studies how tumor and the tumor microenvironment (TME) might respond and/or adapt to chronic therapies. Of interest, combination of a PI3K inhibitor and the CDK4/6 inhibitor ribociclib showed anti-tumor activity in PPGL-derived xenografts^108^. In addition, preclinical studies in renal cell carcinomas, cancers that are uniquely dependent on HIF2α, association of a HIF2α inhibitor with CDK4/6 inhibitor showed enhanced tumor regression^95^. These data suggest that similar combinations may warrant future investigation in PPGLs.

Additional trends followed clinical observations while adding precision and longitudinal context. Cabozantinib activity was stronger in longterm cultures, consistent with increased reliance on angiogenic and receptor tyrosine kinase signaling in the pseudo-hypoxic, extracellular matrix-rich state^106^. Everolimus activity was reduced under hypoxia relative to normoxia, consistent with hypoxia-mediated suppression of mTORC1 signaling^104^. Histone deacetylase inhibitors, including panobinostat, preferentially acted on SDH-mutant organoids in this dataset. HDAC inhibitors can attenuate hypoxia signaling by suppressing HIF transcriptional activity and VEGF expression, in part through HIF-1α destabilization, which provides a mechanistic link to the angiogenic, pseudohypoxic state of SDH deficient tumors^109^. HDAC inhibition has also been reported to increase the half-life of mutant SDHB protein, suggesting an additional SDH-contextual vulnerability to this drug class^103^. The capacity of panobinostat to counter HIF-dependent programs could explain its higher activity in SDH PDOs.

Organoids are increasingly recognized as essential tools for studying tumor biology across diverse cancer types^27,110,111^. Here we show how, in PPGL, they address challenges posed by genomic heterogeneity, the benign nature of many tumors that limits xenograft and cell line generation, and the secretory activity and hypoxia dependence that define the disease. PDOs enable mechanistic studies of lineage plasticity, drug persistence and resistance as well as evolutionary trajectories. In this context, we observed transitions in long-term culture that may parallel aggressive states. Similarly to other slow growing, benign or recalcitrant tumors^32,112^, PPGL PDOs can be used for drug screening within our platform. In this setting, they demonstrate concordance with clinical responses observed in patients, while also enabling more precise tailoring of standard-of-care therapies through biomarker-driven stratification and providing a framework for drug repurposing. With regulatory agencies placing increasing emphasis on patient-derived models and the first ex vivo-only validated therapies entering clinical use^113^, PDOs have the potential to transform PPGL research and therapeutic development.

## Conclusions

PPGL organoids reproduce key tumor features, preserve cellular complexity and subpopulations, and enable condition-dependent discovery. Hypoxia maintains native tumor identity and catecholamine biosynthesis, whereas long-term normoxia unmasks a pseudohypoxic, extracellular matrix-rich state with mesenchymal features. Drug profiling captures clinically observed activity, including belzutifan responses that have been challenging to model in two-dimensional PPGL cultures, and identifies abemaciclib sensitivity in a subset of organoids enriched for proliferative, less mature tumor states.

## Limitations and future directions

The cohort size and condition space are limited, particularly at single-cell resolution. The integration of single-nucleus RNAseq and single-cell RNAseq may bias representation of specific clusters, although joint and stratified analyses mitigated this concern. Drug assays measured bulk viability; development of high-content, cell type–resolved pharmacology is a priority. Future studies should expand oxygen tension range and culture duration systematically, add chronic drug exposure paradigms to study adaptive responses in tumor and stromal compartments, and incorporate genetic perturbations to define causal drivers of noradrenergic to mesenchymal transitions and of HIF pathway selectivity. We have not generated cultures from metastatic tissues, in part due to the limited availability of these tissue types in clinical practice. These critically important samples should be a key focus of investigation in follow-up studies that leverage advances in technology for analyses under more constrained conditions and limited cell numbers.

## Acknowledgments

We thank the patients for donating their samples for this study. We acknowledge the UTHSA/ GCCRI Genome Sequencing Facility, which is supported by UT Health San Antonio, NIH-NCI P30 CA054174 (Cancer Center at UT Health San Antonio) and NIH Shared Instrument grant S10OD030311 (S10 grant to NovaSeq 6000 System), and CPRIT Core Facility Award (RP220662) for processing bulk RNAseq and single nuclei/ single cell RNAseq data. Additional bulk RNAseq sample preparation and sequencing was performed at the Technology Center for Genomics & Bioinformatics (TCGB) at UCLA. We also acknowledge the assistance of the South Texas Research Laboratory (STRL) UTHSCSA and of the translational Pathology Core Laboratory (TPCL) at UCLA for histology and immunohistochemical services. We are grateful to Dr. Carolina Solis-Herrera, MD, and fellows of the Endocrine and Surgical Oncology divisions for their support of patient enrollment at UTHSA, Dr. David Gius, PhD, Rolando Trevino Jr, William Cole Corbett and Daniel Garcia for assistance with staining and slide scanning, and to Karen Bohannon-Johnson for assistance with tissue procurement, and to Nhat Truong, Melissa Heredia, Nasrin Tavanaie and all members of the Soragni and Dahia lab for their technical assistance.

## Author Contributions

S.R., M.K., J.W., M.W.Y., M.L., A.V., R.L., G.T., P.G., and C.J. consented the patients and facilitated the collection of surgical specimens. C.E.Z., C.J., A.S. and P.L.M.D. curated the clinical database. H.G.C., Z-M.C., C.E.Z., B.L., J.P. processed samples and/or genotyping. Y.D., A.T. and M.Z. performed pathology assessments. M.C., S.M., K.A.R., A.S., H.T.L.N., performed organoid establishment, characterization, and drug screening experiments. A.S., J.N.L. performed organoid drug response data analyses. M.P., N.B. and G.E. performed biochemical measurements. M.N., Q.G., C.E.Z. and X.W. processed and analyzed bulk RNAseq, single cell and single nuclei RNAseq analyses. X.W. supervised the bioinformatics analysis. C.E.Z., A.S. and P.L.M.D. wrote the paper with contributions from all authors.

## Materials and Methods

### Patient samples collection

This study was conducted under a protocol approved by the Institutional Review Board (IRB) of the University of Texas Health Science Center at San Antonio (UTHSA). Patients were with diagnosis of pheochromocytoma or paraganglioma (PPGL) provided written informed consent to participate in the repository study under protocol #HSC06-069H (NCT03160274). Tissue from the University of California, Los Angeles (UCLA) patients was collected under IRB protocol #10-001857; University of Texas MD Anderson Cancer Center (MDACC), Houston-TX, under protocol # LAB09-0654; Tufts University in Boston, Massachusetts collected under IRB protocol #8945, and Brigham and Women’s Hospital (BWH), Harvard Medical School, Boston-MA under protocol #2013P000564. UTHSA, BWH and MDACC are members of the American-Australian-Asian-Adrenal-Alliance (A5). Tumor specimens from 39 patients obtained from discarded material during a surgical procedure (and later histologically confirmed as pheochromocytoma or paraganglioma) were included in this study, along with blood or saliva samples were collected.

### Tumor Tissue Processing for Organoid Generation

PPGL tissue fragments were collected from surgery and placed in RPMI or Hypothermosol within 30-60 min of resection. Fragments were enzymatically dissociated into a single-cell suspension following the protocol reported by Nguyen et al. (2020)^33^. Briefly, tissue was mechanically minced into 1-3 mm^3^ fragments in a 200U/mL Collagenase IV (Gibco 17104019) solution diluted in PBS. The tissue suspension was then incubated at 37C and vortexed periodically. Once tissue suspension appeared turbid, the undigested fragments were allowed to sediment, and the cell-enriched supernatant was centrifuged at 500g for 5 minutes. Fresh collagenase IV solution was added to the remaining tissue fragments, and incubation and subsequent steps were repeated until the tissue was completely digested. If red blood cells (RBC) were present in the resulting cell pellet, these were lysed using an ammonium chloride solution (Stem Cell Technology 07850) as previously described^33^. Following RBC lysis, cells were resuspended in serum-free RPMI and counted and analyzed for viability with AO/PI (Nexcellom) using a Cellometer K2 Fluorescent Cell Counter (Nexcellom). Samples collected from UTHSA and UCLA were processed as described above. Samples collected at BWH and MDACC were shipped overnight in Hypothermosol and processed at UTHSA upon arrival, as described above. All samples were frozen in Cryostor CS10 (Stem Cell Technology). Samples were preserved in liquid nitrogen until the initiation of organoid seeding at UCLA.

Samples collected at Tufts University were processed as follows: tumor tissue was minced into 2 to 3mm fragments in 10 mL calcium and magnesium-free Hank’s (CMF HBSS Gibco cat # 12444-014), centrifuged at 500g X 5 min and perform RBC cleanup as above. Pellet was re-suspended in culture medium with 15% FBS and allowed to decant for 30 min. Cell fraction was digested with collagenase B (1.5 mg/mL, Boehringer Manheim, cat # 1088 823) or Collagenase Type (3mg/mL, Worthington Cat# LS004176) in HBSS for 45 min at 37°C. DNAse 1 was added to 50 ug/ml. Digest was allowed to decant and digestion supernatant was removed and washed in CMF HBSS and mechanically disrupted with a plastic pipette tip. Cells were frozen in freezing medium (Gibco cat# 11101-011), stored in liquid nitrogen.

### Organoid Generation: Seeding Mini-Rings and Maxi-Rings Cultures

Primary cells from human PPGL tumors were seeded in a 3:4 mixture of complete Mammocult medium (Stem Cell Technology, supplemented with 2.5ml of hydrocortisone and 1ml of heparin as per manufacturer’s instructions) and Matrigel basement membrane (Corning), maintained at low temperature until plating. For drug screening and immunofluorescence, 10 µl of the mixture were seeded in a ring shape per well of a 96-well plate at 5,000 cells per well. For histological analysis, catecholamine profiling and molecular analyses, 70 µl rings were seeded in 24-well plates at 100,000 cells per well. Rings were solidified at 37 °C for 30 min before adding 100 µl Mammocult (96-well) or 1 ml (24-well). Cultures were maintained at 37 °C, 5% CO_2_, under either normoxia (~21% O_2_) or hypoxia (1% O_2_) for short-term (6 days) or long-term (30 days) timelines. Brightfield imaging was performed daily for normoxic cultures during the first 6 days and every other day thereafter; hypoxic cultures were imaged every other day. For drug treatments, samples were cultured for 3 days (or 4 days if thawed) before replacing media with drug-containing Mammocult (1% DMSO final). Long-term cultures were treated at day 28, with two consecutive days of drug exposure delivered using a Microlab NIMBUS automated liquid handler (Hamilton).

#### Histology and Immunohistochemistry (IHC)

Histopathology analysis was performed on the primary tumor and the PDOs. Organoid cells were processed for histopathology on days 7 and 30 from maxi-ring wells following the following steps: without disturbing the rings, all the supernatant was aspirated, and then the maxi-ring cultures were washed with a pre-warmed PBS (1ml/well). Next, 500µl of 10% buffered formalin was added to each well, incubated at 37°C for 5 min, transferred to for 30 min and kept at 37°C for 24 hours. The next day, all the organoids were transferred using a pipette tip to a Falcon Tube and the supernatant was removed without disturbing the pellet. 4µl of Histogel were added to the pellet. Five ul of Histogel were added to a new cassette allowing it to solidify; with a spatula, the mix of cells and Histogel were placed into the cassette Histogel bed, and 5µl of Histogel were added to the top of the cells. Once solidified, the cassette was closed, wrapped in parafilm and placed on ice for 2-3 minutes, and then cassette was transferred to a beaker containing 70% EtOH, and then processed for embedding, sectioning and staining33. Standard hematoxylin and eosin (HE) staining were performed in primary tumors and PDOs. Immunohistochemistry was performed using the antibodies against Chromogranin A, clone DAK-A3 (DAKO, M0869), dilution 1:200; Tyrosine Hydroxylase (Immunostar, 22941), dilution 1:4000; S100 (DAKO, IR504), ‘ready to use’; CD34 (Ventana, 790-2927), ‘ready to use’; PNMT Polyclonal antibody (Invitrogen, PA5-62289), dilution 1:100; and Ki67 (Ventana, 790-4286), ready to use’. Following deparaffinization and rehydration, heat-mediated antigen retrieval was performed using Citrate at pH 6.0 for PNMT and TH and EDTA at pH 8.0 for Chromogranin A, S100, CD34, and Ki67. All staining was performed using Benchmark Ultra IHC stainer. Slides were scanned using a Leica Aperio Versa 200 scanner and an Olympus VS200 Slide Scanner. The images were acquired and analyzed using QuPath v0.5.1. Slides were evaluated by a pathologist (Y.D.), who was blinded to clinical, histopathological details and sequencing data.

### Immunofluorescence (IF)

At the endpoint of short- or long-term culture, cells in 96-well plates were incubated with 400 µM pimonidazole (MedChemExpress HY-105129A) for 2 h, then fixed in 50 µL 4% PFA after PBS washes. Cells were permeabilized in 200 µL 0.5% Triton X-100 (Sigma-Aldrich T8787) for 1 h, blocked for 2 h in 200 µL 10% FBS (Thermo Fisher Scientific A5670801) and 0.2% Triton X-100, then incubated overnight at 4 °C with primary antibodies diluted in 2% FBS/0.2% Triton X-100: anti-tyrosine hydroxylase (1:4000, Immunostar 22941) and anti-pimonidazole (1:50, Hypoxyprobe HP1-Mab-1). Following PBS washes, cells were incubated with secondary antibodies (1:250 anti-rat, Thermo Fisher A21247; 1:250 anti-mouse, Thermo Fisher A11004) and 1:1000 Hoechst (Thermo Fisher H3570) for nuclear counterstain, washed, and stored at 4 °C. Imaging was performed using the Revolve and Confocal Microscope System (Echo Laboratories).

### Catecholamines and Metanephrines measurement

For catecholamine analysis, media were collected from 24-well cultures every other day (1 mL replaced with fresh Mammocult) and from 96-well cultures every 9 days, with 50 µL media replenished every 3 days between collections. Supernatants were stored individually in 1.5 mL Eppendorf tubes or 96-well plates (Axygen P-96-450V-C-S; Greiner 786261) at −20 °C until analysis. For organoid culture supernatant, norepinephrine, epinephrine and dopamine as well as their respective O-methylated metabolites normetanephrine, metanephrine and 3-methoxytyramine in cell culture supernatants were measured by liquid chromatography tandem mass spectrometry (LC-MS/ MS) using an approach earlier described for the analysis of urinary catecholamine metabolites114. In brief, analytes of interest were extracted from 100µL of cell culture supernatants using the formerly explained solid phase extraction protocol. For detection of metabolites after chromatographic separation, a QTRAP 6500+ (Sciex) applying positive electrospray ionization including the ion source parameters curtain gas (35psi), ESI voltage (5500V), source temperature (450°C), gas 1 (70psi) and gas 2 (60psi) was used. Primary tumor samples were processed and catecholamines (norepinephrine, epinephrine and dopamine), and the catecholamine precursor dihydroxyphenylalanine (DOPA) levels were determined by liquid chromatography with electrochemical detection (LC-ECD) as previously described^115^.

### DNA extraction and genotyping

DNA was extracted from germline and/or frozen tumors utilizing the standard method by Qiagen Genomic-Tip (Qiagen, Valencia, CA). Germline material was available from 32 samples and was processed simultaneously with the tumor DNA. All samples underwent genetic screening for pheochromocytoma/paraganglioma susceptibility genes using standard protocols based on Sanger sequencing and/or target panel sequencing^48^.

### RNA extraction and bulk RNAseq

For primary tumors, RNA was extracted from frozen tumors using the Qiagen RNeasy Mini Kit (Qiagen, Hilden, DEU). For RNAseq purposes, the quality of Total RNAs was ensured by Agilent Tape Station (Agilent Technologies, Santa Clara, CA), and RIN was recorded. Nineteen samples were processed at the Genome Sequencing Facility from Greehey Children’s Cancer Research Institute, utilizing NEBNext rRNA depletion kit v2 (New England Biolabs, Ipswich, MA) and sequenced using Illumina NovaSeq.

For organoid cultures, cells from one maxi-ring were released from Matrigel on average after 6 days or 4 weeks, by incubating with 1 mL of 5 mg/ mL dispase (Life Technologies #17105-041) at 37 °C for 20 min. Organoids were transferred to a 15 mL Falcon tube, neutralized with 9 mL cold RPMI, centrifuged at 1000 g for 5 min, and snap frozen after removing all supernatant. RNA was extracted using the ABclonal mRNA-seq Stranded kit, and quality control was performed as above. Index-tagged, rRNA-depleted libraries are prepared and sequenced at the UCLA Technology Center for Genomics & Bioinformatics (TCGB) using a NovaSeq X Plus.

Raw FASTQ files were aligned to the human reference genome (GRCh38/hg38) using STAR-Fusion v0.13.0, which integrates the STAR aligner with fusion transcript detection capabilities, as we reported^48^. The raw gene count matrix generated above was normalized using the Trimmed Mean of M-values (TMM) method implemented in the edgeR package. When integrating multiple datasets, batch effects were corrected using the ComBat_ seq function from the sva R package. Following batch correction, the data were normalized and transformed to log-transformed counts per million (logCPM) using edgeR, to facilitate downstream clustering and visualization. Additionally, we included publicly available genotype and expression datasets from normalized data from our earlier cohort^48^ and from the TCGA PCPG cohort^3^ (downloaded from the NCI Genomic Data Commons - GDC-Website).

### Clustering and differential gene expression

RNA-seq datasets from multiple sources (UTHSA, UCLA, and publicly available datasets and TCGA^3^ were merged, followed by batch effect correction using ComBat_seq. Expression values were normalized and transformed to logCPM using edgeR. Uniform Manifold Approximation and Projection (UMAP) visualizations were generated using the top genes ranked by median absolute deviation (MAD) across all samples. For the sc/snRNA-seq datasets, pseudobulk expression matrices were created by aggregating all cells within each sample (see below). A UMAP plot was constructed using the top 1,700 MAD-ranked genes.

To generate tumor cell-specific UMAP plots, we derived tumor expression matrices from bulk RNA-seq data by performing BayesPrism deconvolution on raw counts, using tumor profiles from our dataset and the Zethoven et al^50^ sample set as references. For the sc/snRNA-seq data, we aggregated only tumor cells to create pseudobulk tumor matrices. These tumor-specific matrices were then merged, batch-corrected with ComBat_seq, normalized and logCPM-transformed with edgeR, and visualized using UMAP based on the top 1,000 MAD-ranked genes.

Differential expression analysis was performed using Bioconductor R package edgeR^116^. Genes with log2FC >2 and FDR <0.05 were identified as upregulated whereas, genes with log2FC < −2 and FDR <0.05 were identified as downregulated. These comparisons were applied for the following groups: 1) Seven hypoxia samples were compared against thirteen short-term normoxia samples. In total, 26 genes were identified as differentially expressed, with 25 upregulated and 1 downregulated; 2) Nine long-term normoxia samples were compared against thirteen short-term normoxia samples. In total, 630 genes were identified as differentially expressed, with 290 upregulated and 340 downregulated; 3) Samples that showed differential drug response (abemaciclib). PDOs and primary tumor samples were evaluated separately. Samples with the most extreme drug response (based on their viability score-see below). The four most responsive samples in both groups were compared against seven and four resistant samples in the PDOs and primary tumors analysis, respectively. In total, 281 (upregulated: 129, downregulated: 152) and 514 (upregulated: 322, downregulated: 192) genes were identified as differentially expressed in PDO and primary tumors, respectively. Gene expression was compared between response and resistant samples using Wilcoxon signed-rank test.

### Gene set enrichment analysis (GSEA)

Analysis was conducted using Bioconductor R package clusterProfiler^117^. Gene lists sorted based on the log2FC calculated using edgeR were used as input to perform the GSEA for the hallmarks, C2, and C5 gene sets from the MSigDB^118^. Pathways with FDR < 0.05 were considered as significant.

### In silico deconvolution

Bulk RNA sequencing data of PDO and primary tumors were deconvoluted using R packages estimate^119^ and MuSiC^57^. A publicly available cohort of 32 single nuclei PPGLs^50^ were used as a single cell reference for deconvolution using MuSiC. Various scores (stromal, tumor, ESTIMATE) in different PDOs groups based on culture condition and time were compared using Wilcoxon signedrank test. We performed deconvolution of the raw counts of the bulk RNA-seq datasets using PPGL single nuclei RNAseq dataset from Zethoven et al^50^ as cell type reference, and applied BayesPrism to generate a tumor-only expression matrix.

#### Single-cell (sc) and single nuclei (sn) RNASeq

Organoids from five maxi-rings were released from Matrigel on average after 6 days or 4 weeks by incubation in 1 mL/well of 5 mg/mL dispase (Life Technologies #17105-041) at 37 °C for 20 min. Organoids were transferred to a 15 mL Falcon tube, neutralized with 9 mL cold RPMI, centrifuged at 600 g for 5 min, resuspended in 1 mL CryoStor CS10 (Stem Cell Technology #100-1061), cryopreserved at −80 °C until sequenced.

Single-cell suspensions from previously dissociated PPGLs or organoid cultures were prepared for single-cell RNA sequencing following the guidelines provided by 10X Genomics. Briefly, thawed cells were resuspended in 9 mL of serum-free RPMI to remove the freezing reagent and then centrifuged at 500g for 5 minutes. The resulting cell pellet was resuspended in a volume appropriate to its size (200 µL to 1 mL, depending on the pellet size) and assessed for cell count and viability. If the cell viability was ≥70%, the cells were centrifuged again at 500g for 5 minutes, resuspended in 100 µL, and then processed for library construction. In cases where cell viability was <70%, cells were stained with 7-amino-actinomycin D (7-AAD) (Invitrogen 00699350), and live cells were selectively enriched using FACS. These sorted cells were then processed for library construction.

Nuclei isolation was performed as reported^41^. Briefly, a ~50-100 mg frozen PPGLs tumor fragment was minced on ice with 1mL of TST buffer (146mM NaCL, 10mM Tris-HCL pH 7.5, 21mM MgCl2, 1mM CaCl2, 0.03% Tween 20, 0.01% BSA in nuclease-free water) for ten minutes. The homogenized tissue lysate was then strained (70uM strainer), and the strainer was washed with an additional 1mL of TST buffer. The volume of the tissue lysate was adjusted to 5mL and centrifuged at 500g for 5 minutes at 4C. The resulting nuclei pellet was resuspended in ST buffer (similar formulation as TST buffer excluding Tween 20) and strained twice (35uM strainer). The resuspended nuclei solution was stained with 4’9,6 – diamidino – 2-phenylindole (DAPI), and nuclei were enriched by fluorescence-activated cell (nuclei) sorting using a FACS Aria II instrument. Sorted nuclei were resuspended in NSB buffer (1X PBS (without Mg2+ and Ca2+), 1% BSA, and 0.2U/µL RNAse inhibitor) and processed for library construction. For nuclei isolated from dissociated organoid (cells), cells were first pelleted by centrifugation at 500g for 5 minutes, and cells were mechanically lysed by mildly pipetting up-and-down for 10 minutes in 1mL of TST buffer and subsequently prepared following the same methodology as tissue samples.

### sn/scRNAseq library generation and data preprocessing

snRNA-seq was performed with the 10x Genomics 3’ gene expression kit (v.3.1) according to standard protocol. Libraries were sequenced on an Illumina NovaSeq 6000. Cell Ranger v.5.0.0 (10x Genomics)75 was used to align the sequencing reads to the hg38 human reference genome. By quality control removing mitochondrial RNA, ribosomal RNA, and doublets, 123090 of cells were discarded. 211,647 cells were used for further analysis, with a median of 2,536 genes sequenced per nucleus/cell. With a total of 12,754,545,095 reads, an average of 83.08% of reads in cells, 89.12% of reads confidently mapped to the genome, and an average of 74.44% of reads mapped to a pre-mRNA reference transcriptome. For each sample, the filtered matrix from CellRanger’s pipeline was analyzed using Seurat (version 5.0.3)^120^. Doublets were identified, followed by ambient RNA correction in each sample using Bioconductor R packages scDblFinder^121^ (default parameters) and decontX^122^ (default parameters), respectively. As the data was processed using either the single-nuclei or single-cell sequencing with respect to the sample type (frozen tumor, organoids, dissociated tumors), therefore considering these factors, we applied different thresholds to filter the mitochondrial and ribosomal reads. For frozen tumors, nuclei with mitochondrial and ribosomal reads less than 15 and 5 percent, respectively; for organoid samples with single cell and single nuclei, cells and nuclei with mitochondrial reads less than 15 and 20 percent, and ribosomal reads less than 15 and 10 percent, respectively were used for downstream analysis. For the dissociated tumors and organoids sequenced using single-cell sequencing, cells with mitochondrial and ribosomal reads less than 15 were used for downstream analysis. In addition to this, all nuclei/cells considered for the downstream analysis were identified as “singlet” by scDblFinder, having more than 200 genes with UMI greater than 500. Quality control parameters are displayed in Supplementary Table 9.

### sc/snRNAseq analysis

After filtering, data was merged and log-normalized using scaling factor of 10000. Then, the top 3000 variable features were identified, and normalized data was scaled. Dimension reduction was performed on the top variable features using principal component analysis (PCA). To determine the number of PCs used for the downstream analysis, Elbow plot was used that ranks the PCs based on the percentage of variance. To remove the batch effect, batch correction algorithm Harmony^20^ was applied using sample identifier as the grouping variable. Seurat was used to identify the clusters using the FindNeighbors and FindClusters functions. Various resolutions were tested to identify the best possible number of clusters, leading to the selection of resolution 0.3 where we identified 16 clusters. To identify the differentially expressed genes between the clusters FindAllMarkers function in Seurat was used with the following parameters: logfc.threshold = 0.25, min.pct = 0.3, only. pos = TRUE

### Cell type annotation

Various tools were used to annotate the cells. Methods using the cluster’s markers such as ACT^123^ and an R package GPTCelltype^124^ were used to identify the cluster labels. Major cell types identified in the sc/snRNA sequencing dataset were tumor, fibroblasts, endothelial and immune. Clusters were annotated using the tools ACT, GPT, Celltype, Garnett and the cells markers from various published studies^15,62,63,65–68,126–129^. The input used for ACT webserver included top 100 markers (avg_log2FC>1) for all clusters identified using Seurat FindAllMarkers function, species as Human and tissue as Adrenal medulla were used to obtain the annotation for each cluster. To obtain the annotation using GPTCelltype, markers identified using Seurat FindAllMarkers function with avg_log2FC>1 for all clusters were used as input using gpt-3.5-turbo model to identify the cell types. Additionally, other methods which use the expression matrix to annotate the cells such as Garnett^130^, scType^131^, scMatch^132^ and SingleR^133^ were also used. Besides, the expression counts matrix these methods also require the reference gene sets for annotation. The reference sets used for each method were as follows: a) Garnett: a pre-trained classifier provided by Garnett named “adrenal_classifier’’ was used to classify the cells (this classifier is trained on human fetal adrenal tissue^125^); b) scType: a database file provided by scType using the tissue type as adrenal while preparing the gene sets; c) scMatch: FANTOM5 dataset provided within scMatch was used as a reference; d) SingleR: Built-in references Human-PrimaryCellAtlasData and BlueprintEncodeData in an R package SingleR were used to annotate the cells; e) AUCell: markers identified by two previous studies^65,125^ for cell types in pheochromocytoma and paraganglioma were used as gene sets to calculate the scores for each cell and identify the cell type with the highest score. Final cluster labels were defined based on the consensus from the different methods.

### Pseudobulk

Pseudobulk counts for the samples were generated using AggregateExpression function in Seurat.

### Copy number variation analysis

R package inferCNV^134^ with cutoff=0.1 and other parameters as default was used for the copy number variation analysis. Cells which were identified as tumor cells were used as input for copy number variation (CNV) inference while using other cells as reference. This analysis was performed for each sample separately.

### AUCell

We used AUCell to estimate scores of gene sets, which calculates the area under the curve(AUC) score among specified ranked genes. We applied AUCell for the following analyses:

For hypoxia pathway analysis, a curated gene list for HIF1α and HIF2α genes was selected based on published references^60,135^. To assess gene expression across the samples, we evaluated individual expression levels in our tumor cell clusters from sc/snRNA seq data. We excluded genes that showed no expression in over 40% of the samples, those with invariable expression across the tumor clusters, and genes that overlapped between the two signatures. AUCell scores were obtained from the resulting nonoverlapping HIF1α and HIF2α signatures (Supplementary Table 12). P-values were calculated using Wilcoxon test in R.

For analysis of chromaffin cell developmental lineages, a curated gene list representing various developmental stages of chromaffin cells was generated for Schwann cell precursor (SCP) cells, bridge stage, early chromaffin cells, mature chromaffin cells, and sympathoblasts. The genes were annotated based on reported markers^62,63,65–68,126,127^, using similar criteria as described above and listed on Supplementary Table 13. For cell type proportions, we calculated ratios of AUCell score of the specified signatures for each individual cell. For comparison, we performed 1000 iterations of random selection of an equal number of genes from the specified gene sets and calculated the average AUCell score of all the iterations. P-values were calculated using Wilcoxon test in R.

### Reclustering of tumor cell types

To assess the robustness of the trends that we observed in the tumor clusters we further reevaluated the tumor cells. We analyzed the copy number variation calls to identify the cells with very low variations. To make sure that the trends we observed were not affected by the cells expressing very few genes, we applied various thresholds to remove those cells. We used three different approaches: a) We filtered cells based on the nFeatureRNA, mitochondrial and ribosomal percentages. We excluded cells with less than 500 genes and less than 10 and 5 percent and less than 15 and 10 percent of mitochondrial and ribosomal content for the snRNA and scRNA sequenced samples respectively; b) We filtered cells based on the nCountRNA, nFeatureRNA, mitochondrial and ribosomal percentages. We excluded cells with less than 1000 nCountRNA, 500 genes and less than 10 and 5 percent and less than 15 and 10 percent of mitochondrial and ribosomal content for the snRNA and scRNA sequenced samples respectively, and c) We filtered cells based on the 3 median absolute deviations for the nCountRNA, nFeatureRNA, mitochondrial and ribosomal percentages, for each sample separately. After removing the cells based on these cutoffs, we performed the clustering again by normalizing and scaling the data. We plotted CNVs of individual samples in the new clusters and observed consistent results with the original analyses, confirming the tumor identity of the tumor cell clusters.

### Trajectory analysis

Cells differentiation potential and trajectories were identified using Cytotrace^69^ and Monocle3^70^ using default parameters. This analysis was performed in primary tumors and PDOs using the tumor clusters only as input. Genes that are differentially expressed along the trajectory were identified using graph test function. The output is a table listing the Moran’s l value, p value and q value (p-adjusted value). Genes with Moran’s l value >0.25 and q value <0.05 were selected.

### Python-based SCENIC (pySCENIC)

To identify key ‘regulons’ (transcription factors -TFs- and TF-regulatory proteins) and estimate their activity in scRNA-seq data from our cohort, including tumor cell clusters from primary tumors, PDO-N, PDO-H and PDO-L samples, we used the pySCENIC workflow described previously^80^ and the pySCENIC TF database. The top 5000 variable gene expression matrix was used as an input for pySCENIC. First, co-expression modules were identified using GRNBoost. Next, motif enrichment was assessed using the cisTarget algorithm, which ranks motif occurrences across genomic regions and computes a Normalized Enrichment Score (NES). A threshold of NES > 3.0 was applied to identify significantly enriched motifs: Sixty-one regulons were identified. The mean AUCell score of each regulon in different sample types is shown in Supplementary Table 15.

### Noradrenergic (NOR) and mesenchymal (MES) features comparison

We used the NOR and MES transcription factors as previously reported^76^. We used gene set variation analysis to calculate NOR and MES features scores in the following cohorts: TCGA^3^, UT bulk samples^48^ (and current study), UT previous snRNAseq cohort^41^, Zethoven et al PPGL snRNA cohort^50^ and the sn/scRNA data generated in this study. For the bulk RNAseq samples (TCGA and UT cohorts), tumor-only matrix obtained after deconvolution for the bulk samples. For the sc/ snRNA datasets, tumor-only pseudobulk matrix were used. The datasets were log-transformed to counts per million (logCPM) using edgeR in R. Each data set was analyzed separately to calculate the NOR and MES scores (displayed in Supplementary Table 20). The difference of the two scores (MES gsva – NOR gsva) was used for comparison. Samples from all cohorts were also plotted by their molecular cluster type. P-values were calculated using Wilcoxon test in R.

### Data Analysis for Drug Screening

Organoids were incubated at 37°C and 5% CO_2_ throughout drug treatment at either 21% (normoxia) or 1% O_2_ (hypoxia). Drug screen analyses and visualizations were conducted in R using the tidyverse package (for data wrangling and preprocessing^136^. Following drug treatment, ATP luminescence values were normalized to the DMSO control to determine viability^32^ (see below). Viability scores were calculated by normalizing each viability measurement to the mean viability of all samples treated with the respective drug. Heatmaps were generated using ggplot2^136^, patchwork^137^, gridExtra, and scales^138^. Drugs with shared protein targets were clustered using the Jaccard distance via the philentropy package^139^. Samples were clustered based on drug responses using the continuous Euclidean distance. Dendrograms were generated with the pheatmap package^140^. All final figures were assembled in Adobe Illustrator.

### Viability Assay

For 96-well drug-treated cultures, endpoint viability was measured 24h after the final treatment. Wells were washed with PBS, and organoids were released from Matrigel using 50 µl of 5 mg/mL dispase at 37 °C for 25 min, with shaking to ensure complete dissolution. Luminescence was measured after adding 75 µl of ATP reagent (CellTiter-Glo, Promega) using a SpectraMax iD3 plate reader (Molecular Devices), and values were normalized to cell viability. Plates with Z-factor and/or Z-robust factor > 0.2 were considered adequate for further analyses^32^. The viability of each well was calculated by normalizing the luminescent signal to the average signal from the manually seeded control wells. An unpaired t-test with Welch’s correction was performed in Prism 9 (GraphPad). p-values less than 0.05 were deemed significant.

