## Supplemental Tables + Figures for "Patient-derived organoids reveal hypoxia-driven plasticity and therapeutic vulnerabilities in pheochromocytomas and paragangliomas"

### Supplementary Tables

All tables will be available at Synapse repository (synapse.org/PPGLorganoids).

| | UTH ID | Diagnosis | Race | Ethnicity | Sex at Birth | Age at diagnosis | Multiple PPGLs | Metastatic or Recurrent | Tumor Size (cm) | Biochemical Profile | Fold increase of catecholamine (above upper reference limit) | Treatment beside surgery | Familial History | Follow-up time (y) | Genetic Driver Event | Tissue of origin | Driver Gene | Exon | Nucleotide Change | Protein Change | Mutation Classification | Molecular Cluster |
|---|---|---|---|---|---|---|---|---|---|---|---|---|---|---|---|---|---|---|---|---|---|---|
| 1 | **1212** | Pheo | White | Nonhispanic | F | 27 | No | Yes | 5.9 | NEPI+EPI | >20x, 1.2x | | No | 1 | Unknown | | | | | | | C2 |
| 2 | **1216** | Pgl | N/A | Hispanic | F | 27 | No | No | N/A | NEPI | 2.2x | | Yes | 5 | Detected | Germline | SDHB | 2 | c.166_170het_delCCT CA | p.P56Yfs*5 | Pathogenic | C1 |
| 3 | **1219** | Pgl | N/A | Hispanic | M | 59 | No | No | 4.5 | NEPI | 12x | | No | 1.1 | Unknown | | | | | | | C2 |
| 4 | **1226** | Pheo | White | Nonhispanic | F | 47 | No | No | 3.6 | NEPI+EPI | 13x, 12x | | No | 2 | Detected | Germline | RET | 12 | c.2225C>T | p.T742M | VUS | C2 |
| 5 | **1228** | Pheo | N/A | Hispanic | M | 42 | No | No | 4 | NEPI | 3x | | No | 4 | Unknown | | | | | | | C1 |
| 6 | **1231** | Pheo | N/A | Hispanic | F | 77 | No | No | 3.5 | NEPI+EPI | 10x, 10x | | No | 4 | Detected | Somatic | CSDE1, PIK3CA | 15 (CSDE1); 21 (PIK3CA) | CSDE1 c.1611het_delT; | 537Lfs*19, PIK3CA | Pathogenic | C2 |
| 7 | **1234** | Pheo | N/A | Nonhispanic | F | 66 | No | No | N/A | EPI | N/A | | N/A | 3 | Unknown | | | | | | | C2 |
| 8 | **1236** | Pheo | N/A | Hispanic | F | 39 | No | No | N/A | EPI | N/A | | No | 3 | Detected | Somatic | NF1 | 2 | c.96_120het_del, c.121G>GT | p.K33Nfs*3 | Pathogenic | C2 |
| 9 | **1240** | Pheo | White | N/A | F | 51 | No | No | N/A | N/A | N/A | | No | 3 | Detected | Somatic | HRAS | 3 | c.181C>A | p.Q61K | Pathogenic/Likely Pathogenic | C2 |
| 10 | **1244** | Pheo | White | Hispanic | F | 15 | No | No | 4 | NEPI | 5x | | N/A | 3 | Detected | Germline | VHL | 3 | c.499C>T | p.R167W | Pathogenic | C1 |
| 11 | **1251** | Pheo | White | Hispanic | F | 37 | No | No | 5 | NEPI | 20x | | No | 2 | Detected | Somatic | EPAS1 | 12 | c.1589C>CT | p.A530V | Pathogenic | C1 |
| 12 | **1259** | Pgl | Middle Eastern | Nonhispanic | M | 70 | No | No | 5 | NEPI | N/A | | No | 2 | Detected | Somatic | EPAS1 | 12 | c.1615G>GA | p.D539N | Likely Pathogenic | C1 |
| 13 | **1301** | Pheo | N/A | N/A | F | 28 | Yes | Yes | 2.4 | NE | 5x | | No | 8 | Detected | Somatic | UBTF::MAML3 | e15(UBTF):: e2(MAML3) | UBTF c.1626::MAML3 c.469 | isoform 1 | Pathogenic/Likely Pathogenic | C3 |
| 14 | **1302** | Pheo | N/A | N/A | M | 19 | No | No | 5.5 | NEPI | 2.5x, 1.2x | | No | 1 | Detected | Germline | RET | 11 | c.1900T>TG | p.C634G | Pathogenic | C2 |
| 15 | **1305** | Pgl | White | Nonhispanic | M | 19 | Yes | No | 1.8 | NS | 1 | | N/A | 1 | Detected | Germline | VHL | 1 | c.250G>GA | p.V84M | Pathogenic/Likely Pathogenic | C1 |
| 16 | **1306** | Pheo | African-American | Nonhispanic | F | 61 | Yes | No | 3 | NEPI | 5.4 | | Yes | 1 | Detected | Germline | SDHD | 3 | c.198G>A | p.W66* | Pathogenic | C1 |
| 17 | **1307** | Pgl | African-American | Nonhispanic | F | 48 | Yes | No | 2.8 | NEPI | 1.9 | | N/A | 1 | Detected | Somatic | EPAS1 | 12 | c.1592C>CG | p.P531R | Pathogenic | C1 |
| 18 | **1308** | Pgl | White | Nonhispanic | M | 50 | Yes | Yes | 5.5 | NEPI | 1.5 | Radiation treatment for bone mets | Yes | 1 | Detected | Germline | SLC25A11 | 1 | c.95+2het_dupA | splice | | C1 |
| 19 | **1311** | Pgl | White | Nonhispanic | M | 25 | Yes | No | 6 and | NEPI | 12 | | Yes | 1 | Detected | Germline | SDHB | 6 | c.623G>GA | p.G208E | Pathogenic/Likely Pathogenic | C1 |
| 20 | **1312** | Pheo | Asian | Nonhispanic | M | 63 | No | No | N/A | NEPI+EPI | 6x, 10x | | N/A | 1 | Detected | Somatic | NF1 | 15 | c.1692_8het_delTGCT CCT | p.D564Efs*2 | Likely Pathogenic | C2 |
| 21 | **1313** | Pheo | N/A | N/A | M | 54 | No | No | N/A | NEPI | 5x | | N/A | 1 | Unknown | | | | | | | C1 |
| 22 | **1314** | H&N Pgl | N/A | Nonhispanic | M | 33 | No | No | 5.2 | NS | N/A | Embolization before surgery | No | 1 | Detected | Somatic | SDHB | 7 | c.746_765+21het_del | splice | Likely Pathogenic | C1 |
| 23 | **1322** | Pgl | White | Nonhispanic | M | 34 | No | No | 3.3 | NEPI | 2 | | N/A | 1 | Detected | Germline | SDHB | 6 | c.575G>GC | p.C192S | Likely Pathogenic/VUS | C1 |
| 24 | **1325** | Pheo | Native-American | Hispanic | F | 31 | No | No | >5 | NEPI+EPI+DA | 4x, 2x, 4X | | No | 1 | Unknown | | | | | | | C2 |
| 25 | **1328** | Pheo | White | Nonhispanic | M | 60 | Yes | Yes | 11 | NEPI+EPI | 17.2 | Radiation treatment for bone mets | N/A | 1 | Detected | Germline | NF1 | 51 | c.7549C>T | p.R2517* | Pathogenic | C2 |
| 26 | **1335** | Pheo | N/A | N/A | F | 51 | No | No | N/A | EPI | N/A | | N/A | 10 | Detected | Undetermined | TMEM127 | 3 | c.329C>A | p.A110D | Pathogenic/Likely Pathogenic | C2 |
| 27 | **1336** | Pheo | N/A | N/A | F | 50 | No | No | N/A | NEPI | N/A | | N/A | 10 | Unknown | | | | | | | C1 |
| 28 | **1337** | Pheo | N/A | N/A | F | 83 | No | No | N/A | EPI | N/A | | N/A | 11 | Unknown | | | | | | | C2 |
| 29 | **1343** | Pheo | Middle-Eastern | Nonhispanic | F | 39 | No | No | N/A | NEPI | 8x | | N/A | 1 | Unknown | | | | | | | C1 |
| 30 | **1357** | Pgl | African-American | Nonhispanic | F | 15 | No | No | N/A | NS | N/A | | N/A | 0.6 | Detected | Undetermined | SDHB | 6 | c.607G>GT | p.G203* | Pathogenic | C1 |
| 31 | **1369** | Pgl | N/A | N/A | M | 34 | Yes | No | 5.8 | NEPI | 5x | | | 13 | Detected | Germline | SDHC | 5 | c.397C>T | p.R133* | Pathogenic | C1 |
| 32 | **1409** | Pheo | White | Nonhispanic | F | 47 | Yes | No | 7.8 | EPI | | | | 0.6 | Detected | Germline | NF1 | 13 | c.1466A>G | p.Y489C | Pathogenic | C2 |
| 33 | **1415** | Pheo | White | Nonhispanic | F | 54 | No | No | 3.5 | NEPI+EPI | 3x | | N/A | 0.6 | Unknown | | | | | | | C2 |
| 34 | **1432** | Pheo | N/A | N/A | M | 52 | No | No | 14 | NEPI+EPI | 3.5x | | N/A | 0.3 | Detected | Somatic | UBTF::MAML3 | e17(UBTF):: e2(MAML3) | UBTF c.1905::MAML3 c.469 | isoform 2 | Pathogenic/Likely Pathogenic | C3 |
| 35 | **1437** | Pheo | Asian | Nonhispanic | M | 30 | No | No | 2.5 | NE | 2x | | No | 0.3 | Unknown | | | | | | | C2 |

**Supplementary Table 1.** Clinical, molecular characteristics of each sample in the cohort. Abbreviations: Molecular Cluster: C1 - Pseudo-hypoxic Cluster, C2 - Kinase Signaling Cluster, C3 - WNT Cluster

| Gene | log2FC | PValue | FDR | diffexpressed |
|---|---|---|---|---|
| MIR210HG | 4.840412455 | 1.29E-10 | 2.20E-06 | UP |
| DARS1-AS1 | 2.282141851 | 1.50E-09 | 1.28E-05 | UP |
| NDRG1 | 2.668441871 | 7.77E-09 | 3.50E-05 | UP |
| PDE4C | 4.522643357 | 8.18E-09 | 3.50E-05 | UP |
| PDK1-AS1 | 2.325715445 | 2.67E-08 | 9.15E-05 | UP |
| HILPDA | 2.59125045 | 4.01E-08 | 0.0001145954 | UP |
| ZNF395 | 2.572726628 | 1.03E-07 | 0.0002121711 | UP |
| AK4 | 2.651931669 | 1.05E-07 | 0.0002121711 | UP |
| SLC2A1 | 2.218280632 | 1.11E-07 | 0.0002121711 | UP |
| P4HA1 | 1.856870059 | 3.50E-07 | 0.0005996029 | NO |
| PERM1 | 3.871561306 | 3.87E-07 | 0.0006027680 | UP |
| ANKRD37 | 2.39315265 | 5.02E-07 | 0.0006830540 | UP |
| AK4P1 | 2.524016678 | 5.32E-07 | 0.0006830540 | UP |
| VEGFA | 3.203540939 | 5.58E-07 | 0.0006830540 | UP |
| NARF | 0.7501766346 | 1.58E-06 | 0.0018088991 | NO |
| TREM1 | 3.849648698 | 2.02E-06 | 0.0021674288 | UP |
| MT1G | 2.756612964 | 3.17E-06 | 0.0031949025 | UP |
| DARS1 | 0.8340402776 | 3.82E-06 | 0.0034405831 | NO |
| LDHA | 1.439080727 | 3.85E-06 | 0.0034405831 | NO |
| ADM | 2.159068868 | 4.01E-06 | 0.0034405831 | UP |
| CABP1 | 1.766727944 | 6.02E-06 | 0.0049141416 | NO |
| RAB17 | 1.726644132 | 7.03E-06 | 0.0054806641 | NO |
| PDK1 | 1.593888506 | 8.07E-06 | 0.0058595715 | NO |
| SLC16A3 | 2.095937533 | 8.20E-06 | 0.0058595715 | UP |
| ZNF292 | 0.8382152785 | 8.66E-06 | 0.0059401480 | NO |
| PPP1R3G | 2.026480458 | 1.04E-05 | 0.0068391580 | UP |
| TPI1P1 | 1.280854668 | 1.20E-05 | 0.0076291315 | NO |
| BNIP3 | 1.840484656 | 1.32E-05 | 0.0080612861 | NO |
| ARRDC3 | 1.684882538 | 1.45E-05 | 0.0080934632 | NO |
| DDIT4 | 1.801437178 | 1.45E-05 | 0.0080934632 | NO |
| ICOSLG | 1.825539562 | 1.46E-05 | 0.0080934632 | NO |
| INHA | 4.054788761 | 1.51E-05 | 0.0081137718 | UP |
| KISS1R | 4.565562023 | 2.07E-05 | 0.0107279287 | UP |
| LBP | -4.964882876 | 2.76E-05 | 0.0139197872 | DOWN |
| PCCB | -0.8452731492 | 2.92E-05 | 0.0142953377 | NO |
| BNIP3P1 | 1.895344519 | 3.11E-05 | 0.0147910732 | NO |
| ADSS1 | 2.549808448 | 3.55E-05 | 0.0164263366 | UP |
| LUCAT1 | 1.981828208 | 4.61E-05 | 0.02078135 | NO |
| PPP1R13L | 1.750956709 | 5.15E-05 | 0.0222501870 | NO |
| BHLHE40 | 1.428923264 | 5.19E-05 | 0.0222501870 | NO |
| ALDOC | 1.661645706 | 5.68E-05 | 0.0236250605 | NO |
| TAP2 | -0.6042049576 | 5.79E-05 | 0.0236250605 | NO |
| MT1F | 1.575424899 | 5.99E-05 | 0.0238932469 | NO |
| BNIP3L | 1.059041012 | 6.28E-05 | 0.0241066490 | NO |
| C4orf47 | 1.802970467 | 6.33E-05 | 0.0241066490 | NO |
| C4orf3 | 0.8762011077 | 7.02E-05 | 0.0261712621 | NO |
| TPI1 | 1.360060891 | 7.71E-05 | 0.0275861176 | NO |
| CCN5 | 2.133680703 | 7.72E-05 | 0.0275861176 | UP |
| STC1 | 1.871019903 | 8.93E-05 | 0.0312277314 | NO |
| LRRFIP1 | 0.8572524596 | 9.43E-05 | 0.0323468122 | NO |
| NKX3-1 | 1.881194756 | 0.0001021930 | 0.0343488955 | NO |
| PGK1 | 1.479948688 | 0.0001080438 | 0.0356170864 | NO |
| USP13 | -0.5394523244 | 0.0001237335 | 0.0400196273 | NO |
| PFKL | 0.6080986127 | 0.0001281401 | 0.0406773704 | NO |
| CA12 | 2.36245558 | 0.0001307815 | 0.0407610454 | UP |
| HIF1A-AS3 | 1.610825367 | 0.0001554717 | 0.0475910256 | NO |
| MAP2K1 | 0.789220042 | 0.0001586214 | 0.0477033061 | NO |
| STAT1 | -0.6474661494 | 0.0001643148 | 0.0484203641 | NO |
| FUT11 | 0.9809183889 | 0.0001717340 | 0.0484203641 | NO |
| CLEC3B | 2.255440434 | 0.0001722320 | 0.0484203641 | UP |
| ICAM5 | 2.182681127 | 0.0001723044 | 0.0484203641 | UP |

**Supplementary Table 2.**

Differentially expressed genes between short term normoxia (PDO-N) and short term hypoxia (PDO-H) bulk RNAseq, only displaying genes with FDR less than or equal to 0.05.

UP: (log2FC>2 & FDR <0.05) upregulated genes in hypoxia, DOWN: (log2FC<-2 & FDR <0.05) downregulated in hypoxia, NO: not differentially expressed

| Description | set size | enrichment score | NES | pvalue | p.adjust | core_enrichment |
|---|---|---|---|---|---|---|
| HALLMARK_HYPOXIA | 190 | 0.692571655 | 2.821442444 | 1.00E-10 | 1.67E-09 | INHA/CDKN1C/VEGFA/NDRG1/AK4/CA12/SLC2A1/ADM/CCN5/STC1/SLC2A5/P4HA1/DDIT4/ERRFI1/PPFIA4/ALDOC/PDK1/ERO1A/IL6/PGK1/LDHA/BHLHE40/SERPINE1/HK2/SAP30/ANGPTL4/TPI1/PLIN2/MT2A/ENO2/STC2/SLC2A3/EFNA3/MT1E/BTG1/BNIP3L/MAFF/KDM3A/PPP1R3C/IER3/GBE1/GAPDH/VLDLR/TMEM45A/GPC4/PFKP/DUSP1/NCAN/ZNF292/ALDOA/FOS/MIF/ENO1/TGFBI/GCK/S100A4/DPYSL4/IGFBP3/MXI1/SDC4/PHKG1/CCN1/PLAUR/PDGFB/AKAP12/GPI/UGP2/P4HA2/PFKL/TGFB3/ZFP36/ATF3/GYS1/SCARB1/TIPARP/IRS2/WSB1/PGM1/PPP1R15A/KLF7/STBD1/NEDD4L/CAV1/JUN/SLC6A6/RORA/PFKFB3/RBPJ/BGN/CAVIN1/JMJD6/PNRC1/PKLR/NR3C1/CCNG2/NFIL3 |
| HALLMARK_G2M_CHECKPOINT | 198 | -0.624783654 | -2.351458667 | 1.00E-10 | 1.67E-09 | HIF1A/STMN1/MAD2L1/CENPE/GINS2/ORC6/E2F1/NDC80/PLK4/STIL/KIF15/RAD54L/CCNA2/CDC45/KIF11/PRC1/KIF2C/CCNB2/TACC3/CDC6/UBE2C/TOP2A/KNL1/KIF4A/NUSAP1/HMMR/PLK1/E2F2/CDK1/MYBL2/CENPF/AURKB/TTK/BIRC5/BUB1/TPX2/MKI67/POLQ/KIF23/CENPA/NEK2/TROAP/PBK/CDC20/EXO1 |
| HALLMARK_E2F_TARGETS | 199 | -0.57075822 | -2.148968458 | 1.00E-10 | 1.67E-09 | SHMT1/MCM6/MMS22L/NAA38/POLA2/DONSON/TIMELESS/SPAG5/GINS4/CDKN2A/MCM2/EZH2/CTPS1/PTTG1/BRCA2/HMGB3/CDC25A/HELLS/LMNB1/BARD1/CKS2/RFC3/TCF19/GINS1/TFRC/CCNE1/CDCA3/STMN1/MAD2L1/DSCC1/UBE2T/CENPE/ORC6/ASF1B/PLK4/CENPM/SPC24/RAD51AP1/CDCA8/KIF2C/TRIP13/TK1/CCNB2/TACC3/SPC25/TOP2A/KIF4A/HMMR/PLK1/CDK1/MELK/MYBL2/AURKB/BIRC5/RRM2/KIF18B/MKI67/DIAPH3/DEPDC1/BUB1B/DLGAP5/CDC20 |
| HALLMARK_GLYCOLYSIS | 185 | 0.478073803 | 1.951127065 | 4.44E-07 | 5.55E-06 | VEGFA/AK4/EGLN3/SLC16A3/STC1/P4HA1/DDIT4/PPFIA4/SPAG4/ERO1A/PGK1/LDHA/HK2/SAP30/ANGPTL4/TPI1/ENO2/STC2/EFNA3/PLOD2/NOL3/IER3/VLDLR/GPC4/PFKP/ZNF292/ALDOA/MIF/ENO1/TGFBI/DPYSL4/IGFBP3/MXI1/HS6ST2/PLOD1/PGAM1/UGP2/P4HA2/MPI/GYS1/IRS2/PKM |
| HALLMARK_TNFA_SIGNALING_VIA_NFKB | 192 | 0.451583048 | 1.849780778 | 4.38E-06 | 4.38E-05 | VEGFA/CCL20/ICOSLG/IL6/LIF/BHLHE40/SERPINE1/CXCL3/PTGS2/SLC2A3/EGR1/EGR2/INHBA/TNFAIP8/BTG1/CEBPD/MAFF/IER3/IL1B/DUSP1/FOS/SIK1/TRAF1/CXCL1/SLC16A6/EGR3/RHOB/FOSB/SDC4/KLF4/NR4A2/CCN1/PLAUR/BIRC3/TGIF1/JUNB/CXCL6/ZFP36/ATF3/HES1/TIPARP/IRS2/IL23A/PER1/CEBPB/DUSP4/PPP1R15A/JUN/FOSL1/PFKFB3/TUBB2A/PANX1/IER2/ZC3H12A/PNRC1/TNIP1/GADD45A/KLF2/NFIL3 |
| HALLMARK_BILE_ACID_METABOLISM | 90 | -0.557005286 | -1.882714903 | 3.94E-05 | 0.00032827 | ABCA1/PAOX/PEX7/NR3C2/PEX26/SLC27A2/ABCA6/CAT/ALDH9A1/PEX11G/SLC22A18/HACL1/CYP46A1/PFKM/CYP7B1/DIO2/SCP2/AMACR/GCLM/ABCD1/EPHX2/HSD17B4/FDXR/DHCR24/RETSAT/ALDH1A1/GSTK1/CYP39A1/BCAR3/CYP27A1/ACSL5/CH25H/RXRG/LCK/ABCD2/TFCP2L1/ABCG4/ABCA9/RBP1/ABCA4/BMP6/ABCA8 |
| HALLMARK_MITOTIC_SPINDLE | 199 | -0.453623909 | -1.707944692 | 5.38E-05 | 0.00038398 | PXN/CDK5RAP2/KIF3C/FGD6/BRCA2/FGD4/SSH2/ARHGAP27/LMNB1/FSCN1/DOCK4/FLNB/ECT2/MID1/SHROOM2/CENPE/NDC80/KIF15/KIF11/PRC1/KIF2C/CCNB2/TOP2A/KIF4A/NUSAP1/PLK1/CDK1/ANLN/CENPF/TTK/BIRC5/BUB1/TPX2/KIF23/NEK2/DLGAP5 |
| HALLMARK_MTORC1_SIGNALING | 200 | 0.40117347 | 1.65700969 | 7.77E-05 | 0.00048592 | AK4/SLC2A1/EGLN3/STC1/P4HA1/DDIT4/PDK1/ERO1A/PGK1/LDHA/BHLHE40/HK2/TPI1/SLC2A3/PLOD2/GBE1/GAPDH/VLDLR/ALDOA/ENO1/SLC7A5/GPI/PFKL/CD9/PGM1/PPP1R15A/SLC6A6/AURKA/NUPR1/NFIL3/ITGB2/SHMT2 |
| HALLMARK_INTERFERON_ALPHA_RESPONSE | 95 | -0.51685071 | -1.756494153 | 0.000335891 | 0.001679454 | TRIM5/PLSCR1/C1S/MOV10/IL7/CD74/SAMD9/NCOA7/GBP2/IFI44L/LAP3/DHX58/EPSTI1/CD47/IFITM2/SLC25A28/LGALS3BP/PNPT1/OASL/PARP12/HLA-C/IFIH1/GMPR/CASP8/TMEM140/TRIM25/PARP9/ISG15/CMPK2/TRIM14/HERC6/RSAD2/MX1/SELL/GBP4/IFI27/PROCR/CASP1/SAMD9L/BATF2/OAS1/CCRL2/RTP4/IFIT3/USP18/CSF1/IFIT2 |
| HALLMARK_XENOBIOTIC_METABOLISM | 170 | -0.441370145 | -1.627761601 | 0.000323047 | 0.001679454 | GABARAPL1/ACOX1/IRF8/CAT/ALDH9A1/HMOX1/GART/TPST1/PINK1/CYP2E1/AHCY/HACL1/ELOVL5/TKFC/SLC22A1/ALAS1/BCAT1/ACOX2/MCCC2/GCKR/DHRS1/SSR3/MAN1A1/CDA/SLC1A5/NINJ1/ASL/APOE/ACO2/ACOX3/RETSAT/VNN1/GCNT2/AKR1C2/ENTPD5/PTGES/AKR1C3/ESR1/IGFBP4/ABCC2/CDO1/SLC6A12/PEMT/GSTM4/PROS1/PTGR1/CYP27A1/EPHA2/SPINT2/ATOH8/MAOA/JUP/PGD/NQO1/AOX1/RBP4/TGFB2/ABCD2/ITIH1/EPHX1/FBLN1/CD36/HSD11B1/CYP26A1/PTGDS/PDK4/HSD17B2/KYNU/CES1/ADH1C/TDO2 |
| HALLMARK_SPERMATOGENESIS | 86 | -0.486299675 | -1.632954832 | 0.005455014 | 0.024795519 | RFC4/PCSK1N/ZC2HC1C/EZH2/OAZ3/SPATA6/TOPBP1/POMC/MTOR/DDX25/CCNA1/IDE/SCG5/PACRG/SHE/ACE/AKAP4/CCT6B/ACRBP/CNIH2/KIF2C/CCNB2/ADCYAP1/NCAPH/CDK1/TTK/BUB1/NEK2 |
| HALLMARK_INTERFERON_GAMMA_RESPONSE | 189 | -0.3727663 | -1.396656323 | 0.010505502 | 0.043772923 | VCAM1/DHX58/EPSTI1/IFITM2/SLC25A28/NUP93/LGALS3BP/PNPT1/OASL/PARP12/PDE4B/IFIH1/IRF4/CASP8/TNFAIP2/OAS2/DDX58/TRIM25/IL15RA/CIITA/IDO1/ISG15/CMPK2/TRIM14/ST3GAL5/HERC6/RSAD2/CCL5/IRF5/MX1/GBP4/IFI27/TNFSF10/APOL6/HLA-DMA/STAT1/ICAM1/CASP1/VAMP8/HIF1A/SAMD9L/BATF2/CCL7/IL2RB/IFIT1/RTP4/IFIT3/USP18/SECTM1/STAT4/CCL2/METTL7B/IFIT2/CFH/TNFAIP6/BANK1/P2RY14/CD38 |
| HALLMARK_KRAS_SIGNALING_UP | 180 | 0.329910626 | 1.345779301 | 0.016961713 | 0.065237358 | CCL20/IGF2/SPP1/TMEM158/ERO1A/LIF/ANGPTL4/PTGS2/TPH1/CPE/MMP9/INHBA/GALNT3/TRIB2/IL1B/ADAM8/TRAF1/IGFBP3/TFPI/MYCN/KLF4/MMP10/FGF9/PLAUR/BIRC3/AKAP12/ETV4/APOD/F13A1/PPP1R15A/PTPRR/GYPC |
| HALLMARK_TGF_BETA_SIGNALING | 52 | 0.464931539 | 1.523464542 | 0.018563266 | 0.066297379 | CDKN1C/NOG/SERPINE1/TGIF1/SMAD7/JUNB/PPP1R15A/SKI/TGFBR1/ARID4B/FKBP1A/RAB31/NCOR2 |
| HALLMARK_IL6_JAK_STAT3_SIGNALING | 76 | -0.449368332 | -1.48511598 | 0.02215058 | 0.073835267 | CCR1/IL18R1/STAT1/EBI3/TNFRSF12A/IL12RB1/CCL7/TNFRSF21/LTB/CSF1/CD36/CD38/CNTFR/PLA2G2A |
| HALLMARK_UNFOLDED_PROTEIN_RESPONSE | 113 | 0.354343013 | 1.340685941 | 0.03857394 | 0.113607675 | VEGFA/DDIT4/ERO1A/STC2/CHAC1/SLC7A5/IMP3/ATF3/CEBPB/RPS14/TUBB2A/DNAJB9/IGFBP1/NFYA/EIF2AK3/ASNS/SPCS3/ATF4/NPM1/DCP2 |
| HALLMARK_APICAL_JUNCTION | 179 | -0.356040727 | -1.320798249 | 0.038626609 | 0.113607675 | YWHAH/ACTG1/CERCAM/WNK4/NF2/VCAM1/ZYX/MYH10/EGFR/CADM2/ITGA3/SHC1/ITGB4/ICAM4/PIK3CB/LIMA1/MAP4K2/ACTG2/LAMA3/NECTIN2/THY1/ADAM9/FYB1/FBN1/THBS3/PECAM1/ADAM15/SDC3/SLIT2/FSCN1/NEGR1/CD34/MAPK11/CLDN5/ICAM2/ICAM1/SHROOM2/DMP1/MMP2/JUP/VCAN/CX3CL1/MPZL2/NECTIN3/SLC30A3/CLDN4/CADM3/TNFRSF11B/LAMC2/MDK/ADAMTS5/CDH4/LAMB3/CLDN11 |
| HALLMARK_FATTY_ACID_METABOLISM | 142 | -0.357072724 | -1.289671147 | 0.057437408 | 0.147694524 | AOC3/NBN/BPHL/ACADVL/GLUL/AADAT/CRAT/MCEE/HIBCH/D2HGDH/UROD/GCDH/CPT2/EHHADH/ACAA1/YWHAH/GABARAPL1/ACOX1/ACADM/ALDH9A1/HSDL2/HSD17B10/ME1/ACSL4/ACSM3/ELOVL5/ACOT2/ENO3/ACADS/PTPRG/DECR1/UROS/REEP6/TP53INP2/BMPR1B/ACO2/FH/ALDH3A2/FASN/HSD17B4/HMGCS1/DHCR24/RETSAT/VNN1/ALDH1A1/ACSS1/ACSL5/GPD1/MAOA/CA4/G0S2/AQP7/EPHX1/CD36/ADH1C/TDO2/XIST |
| HALLMARK_INFLAMMATORY_RESPONSE | 174 | -0.34377598 | -1.272757095 | 0.05907781 | 0.147694524 | CCL5/SELL/SGMS2/NMUR1/IL7R/TNFSF10/IL18R1/HBEGF/ICAM1/EBI3/HIF1A/ITGB8/CCL7/DCBLD2/GPC3/EREG/CCRL2/LCK/IL2RB/SCN1B/RTP4/CX3CL1/CHST2/TLR3/IL1A/PTGER4/HRH1/CCL2/CSF1/HAS2/VIP/TNFAIP6/SELE/TNFSF15/PCDH7/PROK2 |
| HALLMARK_IL2_STAT5_SIGNALING | 185 | 0.302559546 | 1.234813779 | 0.054264437 | 0.147694524 | CDKN1C/NDRG1/ITIH5/SPP1/P4HA1/LIF/BHLHE40/HK2/PLIN2/SLC2A3/MAFF/IL1RL1/TRAF1/RHOB/ETV4/AHNAK/SOCS1/ARL4A/CTSZ/RORA/AGER/MUC1/NFKBIZ/NFIL3/RHOH/IL4R/CD86 |
| HALLMARK_APICAL_SURFACE | 37 | -0.479368745 | -1.39259308 | 0.071678322 | 0.155822438 | SHROOM2/DCBLD2/IL2RB/CX3CL1/NTNG1/CRYBG1/SLC2A4/RTN4RL1/PCSK9 |
| HALLMARK_KRAS_SIGNALING_DN | 120 | -0.366055063 | -1.293071302 | 0.068965517 | 0.155822438 | SELENOP/TFAP2B/ENTPD7/ZC2HC1C/GPR3/SGK1/P2RX6/SPTBN2/BMPR1B/CCNA1/IDUA/BARD1/CCDC106/VPS50/SNN/ITIH3/DTNB/RSAD2/CYP39A1/KCNN1/MX1/CPB1/SLC38A3/CHST2/PKP1/SLC30A3/AKR1B10/TGFB2/SERPINB2/TFCP2L1/ABCG4/FGF16/KLHDC8A/NGB/ARHGDIG/NRIP2/CNTFR/PNMT |
| HALLMARK_EPITHELIAL_MESENCHYMAL_TRANSITION | 195 | -0.334208481 | -1.254497787 | 0.070503597 | 0.155822438 | MYLK/AREG/CALU/LAMA3/THY1/WNT5A/MSX1/FBN1/DPYSL3/FBLN5/GPC1/SLIT3/GJA1/PFN2/ELN/IGFBP4/CXCL8/SERPINE2/SLIT2/CDH2/COMP/GREM1/SDC1/ECM1/MATN2/LAMA1/TNFRSF12A/MMP2/LOXL1/NID2/VCAN/IL32/PTX3/FZD8/FAP/MEST/ADAM12/ECM2/MMP1/SGCD/VEGFC/TNFRSF11B/MATN3/LAMC2/DKK1/TFPI2/FBLN1/LAMA2/CXCL12/FBLN2/POSTN/ABI3BP/LUM |
| HALLMARK_P53_PATHWAY | 193 | 0.299645971 | 1.228327029 | 0.090207587 | 0.182849937 | NDRG1/DDIT4/LIF/BTG1/IER3/TXNIP/FOS/ALOX15B/S100A4/ZFP36L1/KLF4/ST14/PERP/CDH13/SOCS1/TP63/ATF3/ZBTB16/TM4SF1/PPP1R15A/JUN/SFN/H2AW/FGF13/UPP1/KRT17/GADD45A/STEAP3/NUPR1/RPL18/RAD9A/HRAS/RPL36/PDGFA/OSGIN1/S100A10/FAM162A/RAB40C/FBXW7/PRMT2/SERTAD3/RRAD/POLH/RHBDF2/PLK3/CD81/HINT1 |
| HALLMARK_UV_RESPONSE_UP | 150 | 0.302467722 | 1.198306076 | 0.091772152 | 0.182849937 | CDKN1C/CCK/IL6/ENO2/NPTX2/BTG1/NTRK3/IGFBP2/ALDOA/FOS/RHOB/FOSB/PDLIM3/JUNB/KCNH2/SLC25A4/ATF3/E2F5/HSPA13/SLC6A8/SELENOW/POLG2/BTG3/CYP1A1/AGO2/ASNS/RAB27A/RRAD/MRPL23/OLFM1/CHRNA5/DNAJB1/CTSV/PRKCD/SHOX2/CDC34/HSPA2 |
| HALLMARK_MYOGENESIS | 166 | 0.289640719 | 1.16832538 | 0.095081967 | 0.182849937 | PTGIS/PPFIA4/BHLHE40/CKMT2/STC2/MB/MYBPH/PPP1R3C/CTF1/CFD/LDB3/IGFBP3/EPHB3/MYOZ1/KCNH2/CDH13/APOD/ITGB5/SOD3/MEF2A/AEBP1/KCNH1/APLNR/OCEL1/MYH7/SMTN/IGF1/FXYD1/ACTN2/ERBB3 |
| HALLMARK_PANCREAS_BETA_CELLS | 29 | -0.481492942 | -1.332879742 | 0.111900533 | 0.207223209 | CHGA/ABCC8/LMO2/DPP4/SCGN/SST/SYT13 |
| HALLMARK_ANGIOGENESIS | 32 | 0.438647149 | 1.294716452 | 0.116389549 | 0.20783848 | VEGFA/STC1/SPP1/PRG2/KCNJ8/S100A4/CXCL6/SERPINA5/FGFR1 |
| HALLMARK_COAGULATION | 103 | -0.354864706 | -1.223186989 | 0.13458529 | 0.232043603 | CSRP1/CAPN5/ACOX2/S100A1/PLEK/CTSH/RAPGEF3/ADAM9/FBN1/F12/PECAM1/PROS1/COMP/APOC1/RGN/PLAU/F2RL2/MMP2/PLAT/SERPINB2/C2/MMP1/GDA/ITIH1/DPP4/TFPI2/CFH/CFI |
| HALLMARK_PEROXISOME | 97 | -0.329595727 | -1.123130946 | 0.248407643 | 0.405730315 | EHHADH/ACAA1/SLC27A2/YWHAH/ACOX1/CAT/ALDH9A1/ACSL4/ELOVL5/ABCB1/ABCC5/SCP2/FDPS/ABCD1/CTPS1/EPHX2/HSD17B4/DHCR24/RETSAT/IDE/ALDH1A1/ABCC8/GSTK1/ACSL5/RXRG/FABP6/SEMA3C/CRABP1/ABCD2/CRABP2/TOP2A/DHRS3 |
| HALLMARK_UV_RESPONSE_DN | 143 | 0.27371593 | 1.072955873 | 0.251552795 | 0.405730315 | BDNF/BHLHE40/SERPINE1/MT1E/VLDLR/RGS4/DUSP1/TFPI/CCN1/SMAD7/LTBP1/CELF2/PIK3CD/RUNX1/CAV1/DBP/CACNA1A/PMP22/NR3C1/ATRX/MGLL |
| HALLMARK_HEDGEHOG_SIGNALING | 34 | 0.372195853 | 1.113809847 | 0.30071599 | 0.442229398 | VEGFA/VLDLR/NKX6-1/HEY1/DPYSL2 |
| HALLMARK_CHOLESTEROL_HOMEOSTASIS | 73 | -0.330410883 | -1.086212308 | 0.293159609 | 0.442229398 | ACTG1/ANTXR2/CPEB2/ACSS2/JAG1/TRIB3/TP53INP1/FABP5/FDPS/FASN/NIBAN1/HMGCS1/GPX8/ATF5/PPARG/TNFRSF12A/FBXO6/LPL/AVPR1A/GLDC/MAL2/ADH4 |
| HALLMARK_ESTROGEN_RESPONSE_LATE | 179 | -0.291441886 | -1.081157023 | 0.284692418 | 0.442229398 | CHPT1/ASS1/GFUS/LTF/SLC27A2/FKBP5/HSPB8/IL17RB/S100A9/CISH/ELOVL5/FAM102A/RPS6KA2/SNX10/ACOX2/FABP5/TH/RABEP1/PDZK1/SLC9A3R1/WFS1/SGK1/AREG/RET/ALDH3A2/CACNA2D2/GALE/CCNA1/TOB1/PTGES/SERPINA3/IGFBP4/TST/HR/FLNB/CYP4F11/NPY1R/SLC24A3/TPD52L1/GINS2/PLK4/STIL/MEST/LAMC2/MDK/CDC6/TFPI2/TOP2A/CXCL12/IGSF1/GAL/KIF20A/CDC20/CXCL14/DHRS2 |
| HALLMARK_WNT_BETA_CATENIN_SIGNALING | 40 | -0.375343935 | -1.098669217 | 0.318890815 | 0.455558307 | NOTCH4/TP53/JAG1/KAT2A/JAG2/FRAT1/AXIN2/PSEN2/FZD8/DKK1/WNT6 |
| HALLMARK_COMPLEMENT | 177 | -0.27673222 | -1.02581984 | 0.395982783 | 0.549976088 | KCNIP3/PLEK/CDA/F5/CTSS/CTSH/SH2B3/ADAM9/TIMP2/PCLO/DOCK9/PHEX/ERAP2/DOCK4/CCL5/APOC1/LIPA/PRSS3/CASP10/CASP1/F7/PLA2G7/LCK/CDK5R1/PLAT/SERPINB2/C2/ITIH1/DPP4/TFPI2/CFH/CTSC/CD36/KYNU/PCSK9 |
| HALLMARK_ALLOGRAFT_REJECTION | 166 | -0.277247271 | -1.023624081 | 0.413199426 | 0.558377603 | CDKN2A/CAPG/CTSS/MAP4K1/THY1/CCR5/TLR1/FYB1/FGR/MRPL3/EIF4G3/NCF4/CD28/CD4/CD247/HLA-DOA/CCL5/ITGAL/CCR1/TAP2/ZAP70/HLA-DMA/STAT1/ICAM1/IL27RA/HIF1A/CD3D/IL12RB1/CCL7/EREG/LCK/GCNT1/IL2RB/HLA-DMB/TLR3/CD2/TGFB2/STAT4/CCL2/IL12A/C2/LTB/NOS2/CSF1/IL11 |
| HALLMARK_ADIPOGENESIS | 190 | -0.26893712 | -1.008056537 | 0.435277383 | 0.572733398 | SAMM50/ACLY/IDH3A/HIBCH/SLC25A10/CPT2/PTGER3/ACOX1/HSPB8/DLAT/MTARC2/CAT/ACADM/ME1/BCL2L13/TKT/ACADS/DECR1/SCP2/ABCB8/REEP6/ESRRA/SLC1A5/CYP4B1/LAMA4/MYLK/CHCHD10/APOE/AIFM1/MIGA2/EPHX2/STOM/ACO2/FZD4/CS/LPCAT3/ITGA7/RETSAT/TOB1/QDPR/TST/SPARCL1/PEMT/STAT5A/PPARG/COQ3/LPL/GPX3/NABP1/COL15A1/UCP2/CAVIN2/CD36/ENPP2/ANGPT1 |
| HALLMARK_ESTROGEN_RESPONSE_EARLY | 183 | -0.260552129 | -0.971425767 | 0.532577904 | 0.682792184 | MINDY1/RRP12/PDZK1/SLC9A3R1/ZNF185/MAST4/WFS1/RETREG1/AKAP1/AREG/RET/FASN/MYBBP1A/MYBL1/SYNGR1/FHL2/TOB1/P2RY2/PTGES/GJA1/IGFBP4/RHOD/KCNK15/SH3BP5/REEP1/NAV2/HR/ABLIM1/KAZN/CBFA2T3/WWC1/FLNB/NPY1R/RHOBTB3/MREG/SLC24A3/TPD52L1/GREB1/INPP5F/BCL11B/DHRS3/CXCL12/MLPH/AQP3/DHRS2 |
| HALLMARK_ANDROGEN_RESPONSE | 95 | 0.245197308 | 0.90686833 | 0.681697613 | 0.852122016 | NDRG1/NKX3-1/TNFAIP8/MAK/HPGD/AZGP1/AKAP12/SMS/H1-0/SLC26A2/DNAJB9/MAF/ACTN1/KRT8/IDI1/SPCS3/ZBTB10/TPD52/NGLY1/ARID5B/UBE2I/SCD/PMEPA1/DBI/PDLIM5/GNAI3 |
| HALLMARK_MYC_TARGETS_V1 | 199 | 0.220781961 | 0.909996589 | 0.752442997 | 0.895765472 | PGK1/LDHA/RPL34/RPL14/NAP1L1/SMARCC1/HNRNPA1/PRDX4/AP3S1/RPL18/RPL22/RPS3/VDAC1/DDX18/DEK/EIF3J/POLD2/U2AF1/EIF3D/FBL/PTGES3/RPS2/MRPL23/RSL1D1/PPIA/HNRNPA3/RPS6/RPS10/HDGF/NPM1/RACK1/RPS5/YWHAQ/ERH/RPL6/IMPDH2/HNRNPC/LSM2/SNRPD3/EIF2S2/EIF4H/PCBP1/SNRPD2/HSPE1/PSMA2/EIF4G2/SSB/USP1/SNRPA1/CANX/HPRT1/RPLP0/HNRNPD/PSMA6/NDUFAB1/HSP90AB1/RNPS1/SYNCRIP/PSMC6/TARDBP/XRCC6/CDK4/TCP1/SNRPG/HSPD1/CNBP/HDDC2/LSM7/EEF1B2/EIF2S1/PABPC4/SET/CCT4/PRPS2/PSMD7/PABPC1/TRIM28/CUL1 |
| HALLMARK_APOPTOSIS | 151 | -0.242208829 | -0.880014645 | 0.750364964 | 0.895765472 | TIMP2/FDXR/CCNA1/RETSAT/PSEN2/TNFSF10/CASP1/TNFRSF12A/MMP2/EREG/GPX3/CD2/IL1A/AVPR1A/TGFB2/PLAT/PLPPR4/TOP2A/CD38/LUM |
| HALLMARK_HEME_METABOLISM | 173 | -0.230007159 | -0.852004407 | 0.81061693 | 0.932400932 | SLC7A11/SNCA/OSBP2/LMO2/TMCC2/TFRC/SLC6A9/UCP2/SLC22A4/ACKR1/ABCG2/SELENBP1/ACP5/E2F2/AQP3/SPTA1 |
| HALLMARK_NOTCH_SIGNALING | 31 | 0.266456862 | 0.782656921 | 0.820512821 | 0.932400932 | SAP30/HES1/TCF7L2/FZD7/FZD5/HEYL/DTX4/DTX2 |
| HALLMARK_MYC_TARGETS_V2 | 57 | -0.249387916 | -0.781636632 | 0.842715232 | 0.936350258 | RRP9/NOC4L/PRMT3/NOP56/DCTPP1/DUSP2/NIP7/BYSL/AIMP2/PUS1/MCM4/RABEPK/UTP20/MYC/TMEM97/TFB2M/NOP2/MCM5/RRP12/SRM/SLC29A2/MYBBP1A/PLK4/PLK1 |
| HALLMARK_OXIDATIVE_PHOSPHORYLATION | 200 | -0.214657175 | -0.808297832 | 0.921852388 | 1 | ATP5MC3/SURF1/TIMM10/IDH3B/BDH2/NDUFS2/TCIRG1/ATP5F1C/MDH2/MDH1/MRPS22/ACAT1/SDHC/DLD/NDUFA4/SLC25A6/PDP1/SDHA/HCCS/NDUFB4/MGST3/ATP1B1/IDH3G/MRPS11/IDH1/NDUFA8/NDUFA6/CYC1/SUCLG1/IDH2/ETFB/OXA1L/NDUFV1/SDHB/UQCRC1/SLC25A11/HADHA/ACADVL/ISCU/ATP6V1H/COX15/TOMM70/NDUFA3/PMPCA/IDH3A/FDX1/ATP5F1B/NNT/MTRR/ACAA1/DLAT/GOT2/ATP5MC1/ACADM/HSD17B10/MFN2/NDUFB5/COX10/NDUFS1/ALAS1/ATP5F1A/DECR1/LDHB/UQCRC2/NDUFS8/MRPS15/AIFM1/FXN/OPA1/ACO2/FH/CS/OGDH/SLC25A5/COX17/RETSAT/SLC25A12/SLC25A20/MAOB/PDK4 |
| HALLMARK_PI3K_AKT_MTOR_SIGNALING | 99 | 0.175000125 | 0.650291551 | 0.994550409 | 1 | SLC2A1/PDK1/FGF17/PRKCB/SFN/NGF/PRKAR2A/HRAS/ADCY2/PRKAA2 |
| HALLMARK_REACTIVE_OXYGEN_SPECIES_PATHWAY | 47 | -0.189109408 | -0.571566569 | 0.993162393 | 1 | LSP1/MBP/HMOX2/ABCC1/CAT/GCLM/ERCC2/G6PD/GPX3/NQO1/MGST1 |
| HALLMARK_PROTEIN_SECRETION | 94 | 0.13776433 | 0.506606194 | 1 | 1 | BNIP3/GNAS/AP3S1/STX16/ATP6V1B1/GLA/RAB22A/CAV2/TPD52/SSPN/GALC/TMED10/SOD1 |
| HALLMARK_DNA_REPAIR | 149 | -0.12907176 | -0.468376143 | 1 | 1 | POLA2/RFC4/PDE4B/AK1/TYMS/CDA/ERCC2/AK3/COX17/CMPK2/ADA/CCNO/RFC3/ZWINT/RAD51 |

**Supplementary Table 3.** Gene Set Enrichment Analysis (GSEA): pathway enrichment of the comparison between PDO N (short-term normoxia) and PDO H (short term hypoxia). Key: NES=normalized enrichment score (positive values are enriched in the Hypoxia cohort), Core enrichment: the subset of genes that contributes most to the enrichment result.

| Sample ID | Culture day | Molecular Cluster | Genotype | Number of days in media | PDO supernatant, Average Corrected Values (ng/ml) | | | | | | Tumor tissue (ng/mg) | | | | | Tumor % EPI |
|---|---|---|---|---|---|---|---|---|---|---|---|---|---|---|---|---|
| | | | | | NMN | MN | MTY | NEPI | EPI | DA | DOPA | NEPI | EPI | DA | Total catechol. | |
| 1231T | N/A | N/A | CSDE1+PIK3CA | N/A | N/A | N/A | N/A | N/A | N/A | N/A | n.d. | 338.51 | 233.7 | 2.25 | 574.46 | 40.68 |
| 1231-N | 3 | C2 | CSDE1+PIK3CA | 3 | 41.02 | 21.47 | 18.96 | 6.13 | 2.21 | 0.12 | N/A | N/A | N/A | N/A | N/A | N/A |
| 1231-N | 5 | C2 | CSDE1+PIK3CA | 2 | 71.00 | 44.35 | 35.85 | 1.16 | 1.09 | 0.08 | N/A | N/A | N/A | N/A | N/A | N/A |
| 1231-N | 7 | C2 | CSDE1+PIK3CA | 2 | 43.37 | 27.78 | 22.70 | BLD | BLD | BLD | N/A | N/A | N/A | N/A | N/A | N/A |
| 1231-L | 3 | C2 | CSDE1 + PIK3CA | 3 | 26.79 | 11.72 | 8.01 | 0.01 | BLD | BLD | N/A | N/A | N/A | N/A | N/A | N/A |
| 1231-L | 9 | C2 | CSDE1 + PIK3CA | 2 | 71.19 | 53.78 | 13.55 | BLD | BLD | BLD | N/A | N/A | N/A | N/A | N/A | N/A |
| 1231-L | 17 | C2 | CSDE1 + PIK3CA | 2 | 77.17 | 65.99 | 31.14 | 0.02 | BLD | BLD | N/A | N/A | N/A | N/A | N/A | N/A |
| 1231-L | 23 | C2 | CSDE1 + PIK3CA | 2 | 47.60 | 25.28 | 52.12 | 0.01 | BLD | BLD | N/A | N/A | N/A | N/A | N/A | N/A |
| 1231-L | 29 | C2 | CSDE1 + PIK3CA | 1 | 50.63 | 17.67 | 84.20 | 0.02 | BLD | BLD | N/A | N/A | N/A | N/A | N/A | N/A |
| 1231-L | 9 | C2 | CSDE1+PIK3CA | 2 | 30.93 | BLD | BLD | BLD | BLD | BLD | N/A | N/A | N/A | N/A | N/A | N/A |
| 1231-L | 17 | C2 | CSDE1+PIK3CA | 3 | 27.29 | BLD | BLD | BLD | BLD | BLD | N/A | N/A | N/A | N/A | N/A | N/A |
| 1231-L | 28 | C2 | CSDE1+PIK3CA | 1 | BLD | BLD | BLD | BLD | BLD | BLD | N/A | N/A | N/A | N/A | N/A | N/A |
| 1231-L | 30 | C2 | CSDE1+PIK3CA | 1 | 41.55 | 13.69 | 129.76 | 28.15 | 5.66 | 4.32 | N/A | N/A | N/A | N/A | N/A | N/A |
| 1240T | N/A | N/A | HRAS | N/A | N/A | N/A | N/A | N/A | N/A | N/A | n.d. | 5867.01 | 829.74 | 51.54 | 6748.29 | 12.3 |
| 1240-N | 3 | C2 | HRAS | 3 | 2.63 | 0.79 | 3.13 | BLD | BLD | BLD | N/A | N/A | N/A | N/A | N/A | N/A |
| 1240-N | 4 | C2 | HRAS | 1 | 6.92 | 0.66 | 2.16 | BLD | BLD | BLD | N/A | N/A | N/A | N/A | N/A | N/A |
| 1240-N | 5 | C2 | HRAS | 1 | 0.80 | BLD | BLD | BLD | BLD | BLD | N/A | N/A | N/A | N/A | N/A | N/A |
| 1244T | N/A | N/A | VHL | N/A | N/A | N/A | N/A | N/A | N/A | N/A | n.d. | 1700.41 | 1.52 | 5.79 | 1707.72 | 0.09 |
| 1244-N | 3 | C1 | VHL | 3 | 28.31 | BLD | BLD | BLD | BLD | BLD | N/A | N/A | N/A | N/A | N/A | N/A |
| 1244-N | 5 | C1 | VHL | 2 | 2.65 | BLD | BLD | BLD | BLD | BLD | N/A | N/A | N/A | N/A | N/A | N/A |
| 1244-N | 7 | C1 | VHL | 2 | 2.94 | BLD | BLD | BLD | BLD | BLD | N/A | N/A | N/A | N/A | N/A | N/A |
| 1244-N | 3 | C1 | VHL | 3 | 0.36 | BLD | 0.18 | BLD | BLD | BLD | N/A | N/A | N/A | N/A | N/A | N/A |
| 1244-N | 4 | C1 | VHL | 1 | 1.11 | BLD | 0.95 | BLD | BLD | BLD | N/A | N/A | N/A | N/A | N/A | N/A |
| 1244-N | 5 | C1 | VHL | 1 | 0.74 | BLD | 1.30 | BLD | BLD | BLD | N/A | N/A | N/A | N/A | N/A | N/A |
| 1251T | N/A | N/A | EPAS1 | N/A | N/A | N/A | N/A | N/A | N/A | N/A | 1.70 | 11396.00 | 831.30 | 7.60 | 12227.20 | 6.8 |
| 1251-L | 4 | C1 | EPAS1 | 4 | 2.23 | 0.07 | 0.24 | BLD | BLD | BLD | N/A | N/A | N/A | N/A | N/A | N/A |
| 1251-L | 10 | C1 | EPAS1 | 2 | 2.51 | 0.09 | 0.21 | BLD | BLD | BLD | N/A | N/A | N/A | N/A | N/A | N/A |
| 1251-L | 18 | C1 | EPAS1 | 2 | 2.78 | 0.12 | 0.92 | BLD | BLD | BLD | N/A | N/A | N/A | N/A | N/A | N/A |
| 1251-L | 24 | C1 | EPAS1 | 2 | 3.41 | 0.20 | 1.28 | BLD | BLD | BLD | N/A | N/A | N/A | N/A | N/A | N/A |
| 1251-L | 30 | C1 | EPAS1 | 2 | 4.37 | 0.24 | 2.07 | BLD | BLD | BLD | N/A | N/A | N/A | N/A | N/A | N/A |
| 1301-N | 4 | C3 | UBTF-MAML3 | 4 | 1.44 | 0.03 | 0.13 | BLD | BLD | BLD | N/A | N/A | N/A | N/A | N/A | N/A |
| 1301-N | 5 | C3 | UBTF-MAML3 | 1 | 1.83 | BLD | 1.02 | BLD | BLD | BLD | N/A | N/A | N/A | N/A | N/A | N/A |
| 1301-N | 6 | C3 | UBTF-MAML3 | 1 | 1.31 | BLD | 0.87 | BLD | BLD | BLD | N/A | N/A | N/A | N/A | N/A | N/A |
| 1302T | N/A | N/A | RET | N/A | N/A | N/A | N/A | N/A | N/A | N/A | n.d. | 134.60 | 9.20 | 0.10 | 143.80 | 6.43 |
| 1302-N | 4 | C2 | RET | 4 | 12.87 | 0.77 | 1.06 | BLD | BLD | BLD | N/A | N/A | N/A | N/A | N/A | N/A |
| 1302-N | 5 | C2 | RET | 1 | 12.02 | 0.68 | 1.68 | BLD | BLD | BLD | N/A | N/A | N/A | N/A | N/A | N/A |
| 1302-N | 6 | C2 | RET | 1 | 5.89 | BLD | 1.90 | BLD | BLD | BLD | N/A | N/A | N/A | N/A | N/A | N/A |
| 1302-H | 4 | C2 | RET | 4 | 8.74 | BLD | 2.78 | BLD | BLD | BLD | N/A | N/A | N/A | N/A | N/A | N/A |
| 1302-H | 5 | C2 | RET | 1 | 8.67 | BLD | 4.14 | BLD | BLD | BLD | N/A | N/A | N/A | N/A | N/A | N/A |
| 1302-H | 6 | C2 | RET | 1 | 5.70 | BLD | 5.83 | BLD | BLD | BLD | N/A | N/A | N/A | N/A | N/A | N/A |
| 1306T | N/A | N/A | SDHD | N/A | N/A | N/A | N/A | N/A | N/A | N/A | 1.20 | 2750.70 | 5.00 | 3.60 | 2755.70 | 0.18 |
| 1306-N | 4 | C1 | SDHD | 4 | BLD | BLD | BLD | BLD | BLD | BLD | N/A | N/A | N/A | N/A | N/A | N/A |
| 1306-N | 5 | C1 | SDHD | 1 | BLD | BLD | BLD | BLD | BLD | BLD | N/A | N/A | N/A | N/A | N/A | N/A |
| 1306-N | 6 | C1 | SDHD | 1 | BLD | BLD | BLD | BLD | BLD | BLD | N/A | N/A | N/A | N/A | N/A | N/A |
| 1306-L | 4 | C1 | SDHD | 4 | BLD | BLD | BLD | BLD | BLD | BLD | N/A | N/A | N/A | N/A | N/A | N/A |
| 1306-L | 10 | C1 | SDHD | 2 | BLD | BLD | BLD | BLD | BLD | BLD | N/A | N/A | N/A | N/A | N/A | N/A |
| 1306-L | 18 | C1 | SDHD | 2 | BLD | BLD | BLD | BLD | BLD | BLD | N/A | N/A | N/A | N/A | N/A | N/A |
| 1306-L | 24 | C1 | SDHD | 2 | BLD | BLD | BLD | BLD | BLD | BLD | N/A | N/A | N/A | N/A | N/A | N/A |
| 1306-L | 30 | C1 | SDHD | 2 | BLD | BLD | BLD | BLD | BLD | BLD | N/A | N/A | N/A | N/A | N/A | N/A |
| 1307T | N/A | N/A | EPAS1 | N/A | N/A | N/A | N/A | N/A | N/A | N/A | 1.50 | 10394.00 | 43.30 | 10.20 | 10437.20 | 0.41 |
| 1307-N | 4 | C1 | EPAS1 | 4 | 1.02 | BLD | | BLD | BLD | BLD | N/A | N/A | N/A | N/A | N/A | N/A |
| 1307-N | 5 | C1 | EPAS1 | 1 | 1.14 | BLD | 0.01 | BLD | BLD | BLD | N/A | N/A | N/A | N/A | N/A | N/A |
| 1307-N | 6 | C1 | EPAS1 | 1 | 2.07 | BLD | 1.26 | BLD | BLD | BLD | N/A | N/A | N/A | N/A | N/A | N/A |
| 1311T | N/A | N/A | SDHB | N/A | N/A | N/A | N/A | N/A | N/A | N/A | 1.20 | 929.50 | 1.00 | 6.50 | 930.50 | 0.11 |
| 1311-N | 4 | C1 | SDHB | 4 | 2.18 | BLD | 2.51 | BLD | BLD | BLD | N/A | N/A | N/A | N/A | N/A | N/A |
| 1311-N | 5 | C1 | SDHB | 1 | 1.78 | BLD | 2.72 | BLD | BLD | BLD | N/A | N/A | N/A | N/A | N/A | N/A |
| 1311-N | 6 | C1 | SDHB | 1 | 2.13 | BLD | 2.24 | BLD | BLD | BLD | N/A | N/A | N/A | N/A | N/A | N/A |
| 1311-H | 4 | C1 | SDHB | 4 | 3.66 | BLD | 4.32 | BLD | BLD | BLD | N/A | N/A | N/A | N/A | N/A | N/A |
| 1311-H | 5 | C1 | SDHB | 1 | 4.32 | BLD | 4.61 | BLD | BLD | BLD | N/A | N/A | N/A | N/A | N/A | N/A |
| 1311-H | 6 | C1 | SDHB | 1 | 3.24 | BLD | 3.88 | BLD | BLD | BLD | N/A | N/A | N/A | N/A | N/A | N/A |
| 1312-N | 4 | C2 | NF1 | 4 | BLD | BLD | BLD | BLD | BLD | BLD | N/A | N/A | N/A | N/A | N/A | N/A |
| 1312-N | 5 | C2 | NF1 | 1 | BLD | BLD | BLD | BLD | BLD | BLD | N/A | N/A | N/A | N/A | N/A | N/A |
| 1312-N | 6 | C2 | NF1 | 1 | 0.60 | BLD | BLD | BLD | BLD | BLD | N/A | N/A | N/A | N/A | N/A | N/A |
| 1312-H | 4 | C2 | NF1 | 4 | BLD | BLD | BLD | BLD | BLD | BLD | N/A | N/A | N/A | N/A | N/A | N/A |
| 1312-H | 5 | C2 | NF1 | 1 | BLD | BLD | BLD | BLD | BLD | BLD | N/A | N/A | N/A | N/A | N/A | N/A |
| 1312-H | 6 | C2 | NF1 | 1 | BLD | BLD | BLD | BLD | BLD | BLD | N/A | N/A | N/A | N/A | N/A | N/A |
| 1312-L | 4 | C2 | NF1 | 4 | BLD | BLD | BLD | BLD | BLD | BLD | N/A | N/A | N/A | N/A | N/A | N/A |
| 1312-L | 10 | C2 | NF1 | 2 | 2.53 | 0.09 | 8.65 | BLD | BLD | BLD | N/A | N/A | N/A | N/A | N/A | N/A |
| 1312-L | 18 | C2 | NF1 | 2 | 2.03 | BLD | 17.05 | BLD | BLD | BLD | N/A | N/A | N/A | N/A | N/A | N/A |
| 1312-L | 24 | C2 | NF1 | 2 | 1.77 | BLD | BLD | BLD | BLD | BLD | N/A | N/A | N/A | N/A | N/A | N/A |
| 1312-L | 30 | C2 | NF1 | 2 | 6.09 | 0.37 | 18.01 | BLD | BLD | BLD | N/A | N/A | N/A | N/A | N/A | N/A |
| 1313-N | 4 | C1 | Unknown | 4 | 2.71 | BLD | 7.49 | BLD | BLD | BLD | N/A | N/A | N/A | N/A | N/A | N/A |
| 1313-N | 5 | C1 | Unknown | 1 | 4.76 | BLD | 18.45 | BLD | BLD | BLD | N/A | N/A | N/A | N/A | N/A | N/A |
| 1313-N | 6 | C1 | Unknown | 1 | 2.38 | BLD | 22.11 | BLD | BLD | BLD | N/A | N/A | N/A | N/A | N/A | N/A |
| 1313-H | 4 | C1 | Unknown | 4 | 2.82 | BLD | 26.16 | BLD | BLD | BLD | N/A | N/A | N/A | N/A | N/A | N/A |
| 1313-H | 5 | C1 | Unknown | 1 | 3.26 | BLD | 8.14 | BLD | BLD | BLD | N/A | N/A | N/A | N/A | N/A | N/A |
| 1313-H | 6 | C1 | Unknown | 1 | 2.24 | BLD | 5.80 | BLD | BLD | BLD | N/A | N/A | N/A | N/A | N/A | N/A |
| 1313-L | 4 | C1 | Unknown | 4 | 2.03 | BLD | 6.56 | BLD | BLD | BLD | N/A | N/A | N/A | N/A | N/A | N/A |
| 1313-L | 10 | C1 | Unknown | 2 | 2.34 | BLD | 7.89 | BLD | BLD | BLD | N/A | N/A | N/A | N/A | N/A | N/A |
| 1313-L | 18 | C1 | Unknown | 2 | 4.27 | BLD | 11.01 | BLD | BLD | BLD | N/A | N/A | N/A | N/A | N/A | N/A |
| 1313-L | 24 | C1 | Unknown | 2 | 13.71 | BLD | 16.18 | BLD | BLD | BLD | N/A | N/A | N/A | N/A | N/A | N/A |
| 1313-L | 30 | C1 | Unknown | 2 | 12.15 | BLD | 11.69 | BLD | BLD | BLD | N/A | N/A | N/A | N/A | N/A | N/A |
| 1314-N | 4 | C1 | SDHB | 4 | BLD | BLD | BLD | BLD | BLD | BLD | N/A | N/A | N/A | N/A | N/A | N/A |
| 1314-N | 5 | C1 | SDHB | 1 | BLD | BLD | BLD | BLD | BLD | BLD | N/A | N/A | N/A | N/A | N/A | N/A |
| 1314-N | 6 | C1 | SDHB | 1 | BLD | BLD | BLD | BLD | BLD | BLD | N/A | N/A | N/A | N/A | N/A | N/A |
| 1322-N | 4 | C1 | SDHB | 4 | 1.42 | BLD | 17.71 | BLD | BLD | BLD | N/A | N/A | N/A | N/A | N/A | N/A |
| 1322-N | 5 | C1 | SDHB | 1 | 1.08 | BLD | 1.86 | BLD | BLD | BLD | N/A | N/A | N/A | N/A | N/A | N/A |
| 1322-N | 6 | C1 | SDHB | 1 | 1.65 | BLD | 1.85 | BLD | BLD | BLD | N/A | N/A | N/A | N/A | N/A | N/A |
| 1322-H | 4 | C1 | SDHB | 4 | 3.01 | BLD | 3.51 | BLD | BLD | BLD | N/A | N/A | N/A | N/A | N/A | N/A |
| 1322-H | 5 | C1 | SDHB | 1 | 3.72 | BLD | 5.43 | BLD | BLD | BLD | N/A | N/A | N/A | N/A | N/A | N/A |
| 1322-H | 6 | C1 | SDHB | 1 | 3.52 | BLD | 7.04 | BLD | BLD | BLD | N/A | N/A | N/A | N/A | N/A | N/A |
| 1322-L | 4 | C1 | SDHB | 4 | 1.76 | BLD | 3.80 | BLD | BLD | BLD | N/A | N/A | N/A | N/A | N/A | N/A |
| 1322-L | 10 | C1 | SDHB | 2 | 2.66 | BLD | 7.29 | BLD | BLD | BLD | N/A | N/A | N/A | N/A | N/A | N/A |
| 1322-L | 18 | C1 | SDHB | 2 | 3.92 | BLD | 18.76 | BLD | BLD | BLD | N/A | N/A | N/A | N/A | N/A | N/A |
| 1322-L | 24 | C1 | SDHB | 2 | 3.65 | BLD | 23.03 | BLD | BLD | BLD | N/A | N/A | N/A | N/A | N/A | N/A |
| 1322-L | 30 | C1 | SDHB | 2 | 3.20 | BLD | 31.67 | BLD | BLD | BLD | N/A | N/A | N/A | N/A | N/A | N/A |
| 1325-N | 4 | C2 | Unknown | 4 | 14.98 | 5.02 | 57.57 | 13.52 | BLD | 5.72 | N/A | N/A | N/A | N/A | N/A | N/A |
| 1325-N | 5 | C2 | Unknown | 1 | 14.61 | 4.24 | 187.27 | 3.35 | BLD | BLD | N/A | N/A | N/A | N/A | N/A | N/A |
| 1325-N | 6 | C2 | Unknown | 1 | 6.53 | 1.59 | 101.28 | 0.23 | BLD | BLD | N/A | N/A | N/A | N/A | N/A | N/A |
| 1325-H | 4 | C2 | Unknown | 4 | BLD | BLD | 60.43 | 12.65 | BLD | 6.01 | N/A | N/A | N/A | N/A | N/A | N/A |
| 1325-H | 5 | C2 | Unknown | 1 | 31.06 | 6.66 | 43.11 | 16.18 | 2.01 | BLD | N/A | N/A | N/A | N/A | N/A | N/A |
| 1325-H | 6 | C2 | Unknown | 1 | 26.51 | 6.84 | 29.69 | 12.17 | 1.39 | BLD | N/A | N/A | N/A | N/A | N/A | N/A |
| 1325-L | 4 | C2 | Unknown | 4 | 16.73 | 5.37 | 64.35 | 18.54 | 1.92 | 5.35 | N/A | N/A | N/A | N/A | N/A | N/A |
| 1325-L | 10 | C2 | Unknown | 2 | 6.34 | 1.46 | 268.00 | 12.76 | 1.24 | 39.57 | N/A | N/A | N/A | N/A | N/A | N/A |
| 1325-L | 18 | C2 | Unknown | 2 | 1.52 | 0.32 | 304.53 | 3.34 | BLD | 45.41 | N/A | N/A | N/A | N/A | N/A | N/A |
| 1325-L | 24 | C2 | Unknown | 2 | BLD | BLD | 154.03 | 0.17 | BLD | 19.24 | N/A | N/A | N/A | N/A | N/A | N/A |
| 1325-L | 30 | C2 | Unknown | 2 | BLD | BLD | 59.74 | 0.04 | BLD | BLD | N/A | N/A | N/A | N/A | N/A | N/A |
| 1343-N | 4 | C1 | Unknown | 4 | 2.11 | BLD | 1.78 | BLD | BLD | BLD | N/A | N/A | N/A | N/A | N/A | N/A |
| 1343-N | 5 | C1 | Unknown | 1 | 2.80 | BLD | 2.40 | BLD | BLD | BLD | N/A | N/A | N/A | N/A | N/A | N/A |
| 1343-N | 6 | C1 | Unknown | 1 | 2.49 | BLD | 2.56 | BLD | BLD | BLD | N/A | N/A | N/A | N/A | N/A | N/A |
| 1343-H | 4 | C1 | Unknown | 4 | 2.44 | BLD | 2.78 | BLD | BLD | BLD | N/A | N/A | N/A | N/A | N/A | N/A |
| 1343-H | 5 | C1 | Unknown | 1 | 2.17 | BLD | BLD | BLD | BLD | BLD | N/A | N/A | N/A | N/A | N/A | N/A |
| 1343-H | 6 | C1 | Unknown | 1 | BLD | BLD | BLD | BLD | BLD | BLD | N/A | N/A | N/A | N/A | N/A | N/A |
| 1343-L | 4 | C1 | Unknown | 4 | BLD | BLD | BLD | BLD | BLD | BLD | N/A | N/A | N/A | N/A | N/A | N/A |
| 1343-L | 10 | C1 | Unknown | 2 | 26.45 | BLD | 104.95 | BLD | BLD | BLD | N/A | N/A | N/A | N/A | N/A | N/A |
| 1343-L | 18 | C1 | Unknown | 2 | 54.84 | BLD | 307.79 | BLD | BLD | BLD | N/A | N/A | N/A | N/A | N/A | N/A |
| 1343-L | 24 | C1 | Unknown | 2 | 52.87 | BLD | 277.72 | BLD | BLD | BLD | N/A | N/A | N/A | N/A | N/A | N/A |
| 1343-L | 30 | C1 | Unknown | 2 | 5.50 | BLD | 80.25 | BLD | BLD | BLD | N/A | N/A | N/A | N/A | N/A | N/A |
| 1357-N | 5 | C1 | Unknown | 5 | BLD | BLD | BLD | BLD | BLD | BLD | N/A | N/A | N/A | N/A | N/A | N/A |
| 1357-N | 6 | C1 | Unknown | 1 | BLD | BLD | BLD | BLD | BLD | BLD | N/A | N/A | N/A | N/A | N/A | N/A |
| 1357-N | 7 | C1 | Unknown | 1 | BLD | BLD | BLD | 14.22 | BLD | BLD | N/A | N/A | N/A | N/A | N/A | N/A |

**Supplementary Table 4.** Catecholamines and Metanephrines in PDO supernatant (average corrected: average of duplicate values and divided by the number of days exposed to media at the time of collection), Catecholamines measured in tumor tissue. Abbreviations: T-tumor tissue; N-organoid supernatant, normoxia; H-organoid supernatant, hypoxia; L-organoid supernatant, long-term & normoxia; NMN - Normetanephrine; MN - Metanephrine; MTY - 3-Methoxytyramine; NEPI - Norepinephrine; EPI - Epinephrine; DA - Dopamine; DOPA- Dihydroxyphenylalanine,BLD - below limit of detectio; Molecular Cluster: C1 - Pseudohypoxic Cluster; C2 - Kinase Signaling Cluster; C3 - WNT Signaling Cluster

| Sample ID | UMAP1 | UMAP2 | Sample type | Molecular Group | RNA source | Source | Reference |
|---|---|---|---|---|---|---|---|
| 1343-PDO-N | -2.10364192 | 1.000568106 | PDO | C1 | scRNAseq | scRNA | this study |
| 1343-PDO-H | -2.060702922 | 1.572842057 | PDO | C1 | scRNAseq | scRNA | this study |
| 1343T | 0.258554212 | 1.712540614 | primary, PDT | C1 | scRNAseq | scRNA | this study |
| 1195T | -0.450913491 | 0.469062583 | primary, PF | C1 | snRNAseq | snRNA | this study |
| 1244T | -1.096423172 | 2.293408927 | primary, PF | C1 | snRNAseq | snRNA | this study |
| 1259T | -0.632528357 | 0.712609744 | primary, PF | C1 | snRNAseq | snRNA | this study |
| 1314T | -2.856777225 | -2.763378833 | primary, PF | C1 | snRNAseq | snRNA | this study |
| 1313T | 0.184240892 | 1.297705898 | primary, PDT | C1 | snRNAseq | snRNA | this study |
| 1313-PDO-H | -1.154415918 | -0.2262855 | PDO | C1 | snRNAseq | snRNA | this study |
| 1313-PDO-N | -1.627485123 | -0.023460978 | PDO | C1 | snRNAseq | snRNA | this study |
| 1313-PDO-L | -1.204979342 | -0.231714152 | PDO | C1 | snRNAseq | snRNA | this study |
| 1322T | 0.387054643 | 1.574208828 | primary, PDT | C1 | snRNAseq | snRNA | this study |
| 1322-PDO-H | -0.663927925 | -0.823224629 | PDO | C1 | snRNAseq | snRNA | this study |
| 1322-PDO-N | -0.343368809 | -0.518399654 | PDO | C1 | snRNAseq | snRNA | this study |
| 1322-PDO-L | -2.146141927 | 0.122379077 | PDO | C1 | snRNAseq | snRNA | this study |
| 1322T | -2.444944696 | -1.907365965 | primary, PF | C1 | snRNAseq | snRNA | this study |
| 1311-PDO-N | -3.542104898 | -1.603634875 | PDO | C1 | bulk RNAseq | ucla_bulk | this study |
| 1322-PDO-H | -3.268370584 | -1.529129419 | PDO | C1 | bulk RNAseq | ucla_bulk | this study |
| 1313-PDO-H | -1.134049461 | -0.106187589 | PDO | C1 | bulk RNAseq | ucla_bulk | this study |
| 1322-PDO-N | -3.288049394 | -1.4580157 | PDO | C1 | bulk RNAseq | ucla_bulk | this study |
| 1313-PDO-L | -1.447840866 | -0.181409945 | PDO | C1 | bulk RNAseq | ucla_bulk | this study |
| 1311-PDO-H | -3.472018293 | -1.690457318 | PDO | C1 | bulk RNAseq | ucla_bulk | this study |
| 1322-PDO-L | -2.255527758 | -0.026404899 | PDO | C1 | bulk RNAseq | ucla_bulk | this study |
| 1343-PDO-L | -2.253445567 | 0.496869154 | PDO | C1 | bulk RNAseq | ucla_bulk | this study |
| 1343-PDO-H | -1.786373236 | 1.479651642 | PDO | C1 | bulk RNAseq | ucla_bulk | this study |
| 1343-PDO-N | -1.762814666 | 1.43765873 | PDO | C1 | bulk RNAseq | ucla_bulk | this study |
| 1314-PDO-N | -2.136452778 | 0.105646351 | PDO | C1 | bulk RNAseq | ucla_bulk | this study |
| 1336-PDO-N | -0.437861029 | -0.95866465 | PDO | C1 | bulk RNAseq | ucla_bulk | this study |
| 1313-PDO-N | -1.089187555 | -0.095585881 | PDO | C1 | bulk RNAseq | ucla_bulk | this study |
| 1251-PDO-L | -2.164623844 | 0.188823482 | PDO | C1 | bulk RNAseq | ucla_bulk | this study |
| 1313T | 0.238130793 | 1.474297984 | primary, PDT | C1 | bulk RNAseq | ucla_bulk | this study |
| 1251-PDO-N | -1.418869985 | 0.863968206 | PDO | C1 | bulk RNAseq | ucla_bulk | this study |
| 1244T | -0.602844555 | 2.126189306 | primary, PDT | C1 | bulk RNAseq | ucla_bulk | this study |
| 1343T | 0.247434778 | 1.676623446 | primary, PDT | C1 | bulk RNAseq | ucla_bulk | this study |
| 1306-PDO-L | -2.349866249 | 0.416630668 | PDO | C1 | bulk RNAseq | ucla_bulk | this study |
| 1311-PDO-H5 | -3.478131853 | -1.54143013 | PDO | C1 | bulk RNAseq | ucla_bulk | this study |
| 1241T | 1.172631194 | 1.311101801 | primary, FFPE | C1 | bulk RNAseq | ut_bulk_new | this study |
| 1262T | -2.979324137 | -2.868232142 | primary, FFPE | C1 | bulk RNAseq | ut_bulk_new | this study |
| 1289T | -0.874095404 | -1.738585037 | primary, FFPE | C1 | bulk RNAseq | ut_bulk_new | this study |
| 1306T | -2.687589189 | -2.114128734 | primary, PF | C1 | bulk RNAseq | ut_bulk_new | this study |
| 1311T | -3.547555529 | -2.312252017 | primary, PF | C1 | bulk RNAseq | ut_bulk_new | this study |
| 1317T | 2.30837634 | 1.291818138 | primary, PF | C1 | bulk RNAseq | ut_bulk_new | this study |
| 1327T | -2.017177452 | -1.200998095 | primary, PF | C1 | bulk RNAseq | ut_bulk_new | this study |
| 1361T | -1.368558175 | 2.459038301 | primary, PF | C1 | bulk RNAseq | ut_bulk_new | this study |
| 1415-PDO-N(1) | 0.375273519 | -1.575381972 | PDO | C2 | scRNAseq | scRNA | this study |
| 1415-PDO-N(2) | -0.750579267 | -0.691174046 | PDO | C2 | scRNAseq | scRNA | this study |
| 1415-PDO-N(3) | -0.734848702 | -0.775648724 | PDO | C2 | scRNAseq | scRNA | this study |
| 1415T-D1 | 0.794504185 | 0.981811829 | primary, PDT | C2 | bulk RNAseq | scRNA | this study |
| 1415T-D2 | 0.63290046 | 1.326210019 | primary, PDT | C2 | bulk RNAseq | scRNA | this study |
| 1415T-D3 | 0.697417367 | 1.206143554 | primary, PDT | C2 | bulk RNAseq | scRNA | this study |
| 1119T | 0.765957418 | -2.752174927 | primary, PF | C2 | bulk RNAseq | snRNA | this study |
| 1182T | 1.964387697 | -0.660532447 | primary, PF | C2 | bulk RNAseq | snRNA | this study |
| 1196T | 0.192894923 | -1.913117482 | primary, PF | C2 | bulk RNAseq | snRNA | this study |
| 393T | 2.91135541 | -0.402995473 | primary, PF | C2 | bulk RNAseq | snRNA | this study |
| 802T | 2.017900738 | 1.738417456 | primary, PF | C2 | bulk RNAseq | snRNA | this study |
| 1312T | 0.479970982 | 1.404335 | primary, PDT | C2 | bulk RNAseq | snRNA | this study |
| 1312-PDO-L | -1.974336529 | -0.048845705 | PDO | C2 | scRNAseq | snRNA | this study |
| 1312-PDO-H | 1.692663277 | -0.038636813 | PDO | C2 | scRNAseq | snRNA | this study |
| 1231-PDO-N | 1.146387581 | 0.029807627 | PDO | C2 | bulk RNAseq | ucla_bulk | this study |
| 1231T | 0.830050589 | 0.880097911 | primary, PDT | C2 | bulk RNAseq | ucla_bulk | this study |
| 1335-PDO-N | 1.74550984 | -0.486911183 | PDO | C2 | bulk RNAseq | ucla_bulk | this study |
| 1231-PDO-L | -1.931552308 | -0.105388389 | PDO | C2 | bulk RNAseq | ucla_bulk | this study |
| 1312-PDO-H | 1.852713831 | -0.012340334 | PDO | C2 | bulk RNAseq | ucla_bulk | this study |
| 1302-PDO-H | 1.456999262 | -2.216556518 | PDO | C2 | bulk RNAseq | ucla_bulk | this study |
| 1312-PDO-L | -1.640328001 | -0.252532076 | PDO | C2 | bulk RNAseq | ucla_bulk | this study |
| 1337-PDO-L | -1.085497667 | -0.658165317 | PDO | C2 | bulk RNAseq | ucla_bulk | this study |
| 1337-PDO-N | -0.295797111 | -1.708948428 | PDO | C2 | bulk RNAseq | ucla_bulk | this study |
| 1312-PDO-N | 1.728697486 | 0.045200854 | PDO | C2 | bulk RNAseq | ucla_bulk | this study |
| 1335-PDO-L | -1.192866334 | -0.600594163 | PDO | C2 | bulk RNAseq | ucla_bulk | this study |
| 1312T | 0.381821261 | 1.500915913 | primary, PDT | C2 | bulk RNAseq | ucla_bulk | this study |
| 1302-PDO-N | 1.472105279 | -2.095130502 | PDO | C2 | bulk RNAseq | ucla_bulk | this study |
| 1234T | 1.430374195 | -2.05923408 | primary, PF | C2 | bulk RNAseq | ut_bulk_new | this study |
| 1236T | 2.943217602 | -0.922949738 | primary, PF | C2 | bulk RNAseq | ut_bulk_new | this study |
| 1263T | 1.807573361 | 1.096744254 | primary, FFPE | C2 | bulk RNAseq | ut_bulk_new | this study |
| 1295T | 2.275627088 | 1.713664131 | primary, FFPE | C2 | bulk RNAseq | ut_bulk_new | this study |
| 1325T | 0.797473447 | -2.66235138 | primary, PF | C2 | bulk RNAseq | ut_bulk_new | this study |
| 1328T | 2.442823706 | 0.126658692 | primary, PF | C2 | bulk RNAseq | ut_bulk_new | this study |
| 1359T | 2.082685295 | 0.566685965 | primary, FFPE | C2 | bulk RNAseq | ut_bulk_new | this study |
| 1415T | 0.657017116 | -1.85743028 | primary, PF | C2 | bulk RNAseq | ut_bulk_new | this study |
| 1301-PDO-N | -0.748148794 | 7.221017649 | PDO | C3 | bulk RNAseq | ucla_bulk | this study |
| 1301T | -0.685827154 | 7.31718396 | primary, PF | C3 | bulk RNAseq | ut_bulk_new | this study |
| 1373T | -0.705285344 | 7.660149319 | primary, PF | C3 | bulk RNAseq | ut_bulk_new | this study |
| 1405T | -0.857716943 | 7.587963142 | primary, PF | C3 | bulk RNAseq | ut_bulk_new | this study |
| TCGA-S7-A7WQ-01A | -1.838257165 | 2.646854438 | primary, PF | C1 | bulk RNAseq | TCGA | Fishbein et al |
| TCGA-SR-A6MX-05A | -3.445957809 | -2.601060586 | primary, PF | C1 | bulk RNAseq | TCGA | Fishbein et al |
| TCGA-P8-A6RX-01A | -1.58383689 | 2.400261858 | primary, PF | C1 | bulk RNAseq | TCGA | Fishbein et al |
| TCGA-QR-A70Q-01A | -3.09725516 | -2.741866032 | primary, PF | C1 | bulk RNAseq | TCGA | Fishbein et al |
| TCGA-SR-A6MV-01A | -0.904711662 | 0.736827428 | primary, PF | C1 | bulk RNAseq | TCGA | Fishbein et al |
| TCGA-QR-A703-01A | -1.566919146 | 2.429742848 | primary, PF | C1 | bulk RNAseq | TCGA | Fishbein et al |
| TCGA-QT-A5XL-01A | -1.443021496 | 2.173938011 | primary, PF | C1 | bulk RNAseq | TCGA | Fishbein et al |
| TCGA-SR-A6MX-01A | -3.62460782 | -2.508442793 | primary, PF | C1 | bulk RNAseq | TCGA | Fishbein et al |
| TCGA-RW-A680-01A | -3.317536951 | -2.147318264 | primary, PF | C1 | bulk RNAseq | TCGA | Fishbein et al |
| TCGA-QR-A7IP-01A | -3.050530706 | -2.220716027 | primary, PF | C1 | bulk RNAseq | TCGA | Fishbein et al |
| TCGA-QR-A70I-01A | -3.466277114 | -2.184031233 | primary, PF | C1 | bulk RNAseq | TCGA | Fishbein et al |
| TCGA-QR-A6GX-01A | -3.236344934 | -2.082108008 | primary, PF | C1 | bulk RNAseq | TCGA | Fishbein et al |
| TCGA-RW-A68C-01A | -1.570895507 | 2.256592499 | primary, PF | C1 | bulk RNAseq | TCGA | Fishbein et al |
| TCGA-RW-A7CZ-01A | -3.262289515 | -2.5129776 | primary, PF | C1 | bulk RNAseq | TCGA | Fishbein et al |
| TCGA-WB-A81G-01A | -3.456342819 | -2.260133543 | primary, PF | C1 | bulk RNAseq | TCGA | Fishbein et al |
| TCGA-QR-A6H0-01A | -3.100924559 | -2.864378797 | primary, PF | C1 | bulk RNAseq | TCGA | Fishbein et al |
| TCGA-QR-A6GT-01A | -3.940089058 | -2.114589363 | primary, PF | C1 | bulk RNAseq | TCGA | Fishbein et al |
| TCGA-QR-A6GU-01A | -3.595949332 | -2.00610186 | primary, PF | C1 | bulk RNAseq | TCGA | Fishbein et al |
| TCGA-SR-A6MX-06A | -3.592879799 | -2.540520719 | primary, PF | C1 | bulk RNAseq | TCGA | Fishbein et al |
| TCGA-QR-A705-01A | -2.829659349 | -2.72166094 | primary, PF | C1 | bulk RNAseq | TCGA | Fishbein et al |
| TCGA-QR-A70P-01A | -3.258827845 | -2.135227453 | primary, PF | C1 | bulk RNAseq | TCGA | Fishbein et al |
| TCGA-QR-A6H3-01A | -3.118918865 | -2.800624722 | primary, PF | C1 | bulk RNAseq | TCGA | Fishbein et al |
| TCGA-QR-A70J-01A | -1.79012693 | 2.5175267 | primary, PF | C1 | bulk RNAseq | TCGA | Fishbein et al |
| TCGA-W2-A7HH-01A | -2.199193164 | -1.958421795 | primary, PF | C1 | bulk RNAseq | TCGA | Fishbein et al |
| TCGA-WB-A81E-01A | -2.51041395 | -2.441350078 | primary, PF | C1 | bulk RNAseq | TCGA | Fishbein et al |
| TCGA-W2-A7H7-01A | -3.594169918 | -2.376676982 | primary, PF | C1 | bulk RNAseq | TCGA | Fishbein et al |
| TCGA-QT-A7U0-01A | -1.123345524 | 1.21064284 | primary, PF | C1 | bulk RNAseq | TCGA | Fishbein et al |
| TCGA-WB-A80P-01A | -3.267698012 | -2.167645335 | primary, PF | C1 | bulk RNAseq | TCGA | Fishbein et al |
| TCGA-SP-A6QD-01A | -3.647778 | -1.975960482 | primary, PF | C1 | bulk RNAseq | TCGA | Fishbein et al |
| TCGA-TT-A6YK-01A | -0.9835118 | 0.786319052 | primary, PF | C1 | bulk RNAseq | TCGA | Fishbein et al |
| TCGA-QR-A70N-01A | -1.693742378 | 2.601289653 | primary, PF | C1 | bulk RNAseq | TCGA | Fishbein et al |
| TCGA-SR-A6MR-01A | -1.478738328 | -1.10428342 | primary, PF | C1 | bulk RNAseq | TCGA | Fishbein et al |
| TCGA-TT-A6YP-01A | -1.653272229 | 2.448519025 | primary, PF | C1 | bulk RNAseq | TCGA | Fishbein et al |
| TCGA-S7-A7X0-01A | -1.117001855 | 1.201047836 | primary, PF | C1 | bulk RNAseq | TCGA | Fishbein et al |
| TCGA-RW-A67X-01A | -1.831798035 | 2.388758927 | primary, PF | C1 | bulk RNAseq | TCGA | Fishbein et al |
| TCGA-QR-A6ZZ-01A | -1.365222736 | 2.401934843 | primary, PF | C1 | bulk RNAseq | TCGA | Fishbein et al |
| TCGA-WB-A822-01A | -3.847348731 | -2.09912083 | primary, PF | C1 | bulk RNAseq | TCGA | Fishbein et al |
| TCGA-S7-A7WV-01A | 2.308409665 | 1.328983229 | primary, PF | C1 | bulk RNAseq | TCGA | Fishbein et al |
| TCGA-QT-A5XJ-01A | -2.400974259 | -2.525060383 | primary, PF | C1 | bulk RNAseq | TCGA | Fishbein et al |
| TCGA-W2-A7UY-01A | -1.806318487 | 2.576333012 | primary, PF | C1 | bulk RNAseq | TCGA | Fishbein et al |
| TCGA-SP-A6QH-01A | -1.411253295 | 1.641733364 | primary, PF | C1 | bulk RNAseq | TCGA | Fishbein et al |
| TCGA-QR-A7IN-01A | -0.950879748 | 0.573790296 | primary, PF | C1 | bulk RNAseq | TCGA | Fishbein et al |
| TCGA-QR-A6H5-01A | -1.517102248 | 2.188675732 | primary, PF | C1 | bulk RNAseq | TCGA | Fishbein et al |
| TCGA-RM-A68W-01A | -2.853716603 | -2.860607452 | primary, PF | C1 | bulk RNAseq | TCGA | Fishbein et al |
| TCGA-QR-A6H6-01A | -3.461666281 | -2.390293074 | primary, PF | C1 | bulk RNAseq | TCGA | Fishbein et al |
| TCGA-S7-A7WP-01A | -1.360414781 | 1.971470199 | primary, PF | C1 | bulk RNAseq | TCGA | Fishbein et al |
| TCGA-RM-A68T-01A | -3.05865185 | -2.977702289 | primary, PF | C1 | bulk RNAseq | TCGA | Fishbein et al |
| TCGA-S7-A7X1-01A | -1.659670159 | 2.4762778 | primary, PF | C1 | bulk RNAseq | TCGA | Fishbein et al |
| TCGA-QR-A6GZ-05A | -1.460694747 | 2.624608041 | primary, PF | C1 | bulk RNAseq | TCGA | Fishbein et al |
| TCGA-WB-A814-01A | -1.315946767 | 1.552973363 | primary, PF | C1 | bulk RNAseq | TCGA | Fishbein et al |
| TCGA-QR-A70D-01A | -3.306238043 | -2.172934172 | primary, PF | C1 | bulk RNAseq | TCGA | Fishbein et al |
| TCGA-WB-A821-01A | -1.260369933 | 1.220255617 | primary, PF | C1 | bulk RNAseq | TCGA | Fishbein et al |
| TCGA-SP-A6QG-01A | -2.733738875 | -2.179024349 | primary, PF | C1 | bulk RNAseq | TCGA | Fishbein et al |
| TCGA-QR-A706-01A | -1.284764346 | 1.720567545 | primary, PF | C1 | bulk RNAseq | TCGA | Fishbein et al |
| TCGA-RW-A688-01A | 0.758722288 | -2.786690037 | primary, PF | C2 | bulk RNAseq | TCGA | Fishbein et al |
| TCGA-QR-A6GR-01A | 2.169203818 | 1.034775543 | primary, PF | C2 | bulk RNAseq | TCGA | Fishbein et al |
| TCGA-SR-A6MP-01A | 2.292020488 | -0.079272664 | primary, PF | C2 | bulk RNAseq | TCGA | Fishbein et al |
| TCGA-S7-A7WW-01A | 1.546365845 | 1.49851766 | primary, PF | C2 | bulk RNAseq | TCGA | Fishbein et al |
| TCGA-QT-A5XK-01A | 2.558380498 | -0.777504592 | primary, PF | C2 | bulk RNAseq | TCGA | Fishbein et al |
| TCGA-WB-A80L-01A | 0.15083453 | -0.114997278 | primary, PF | C2 | bulk RNAseq | TCGA | Fishbein et al |
| TCGA-WB-A81Q-01A | 1.759930056 | -1.244256443 | primary, PF | C2 | bulk RNAseq | TCGA | Fishbein et al |
| TCGA-RW-A684-01A | 1.287798582 | -1.022291075 | primary, PF | C2 | bulk RNAseq | TCGA | Fishbein et al |
| TCGA-SR-A6N0-01A | 1.98162369 | 1.650595434 | primary, PF | C2 | bulk RNAseq | TCGA | Fishbein et al |
| TCGA-WB-A81T-01A | 2.116216636 | 0.223465536 | primary, PF | C2 | bulk RNAseq | TCGA | Fishbein et al |
| TCGA-QR-A70M-01A | 1.466185636 | -1.15349609 | primary, PF | C2 | bulk RNAseq | TCGA | Fishbein et al |
| TCGA-QT-A5XP-01A | 2.791616953 | -1.056953229 | primary, PF | C2 | bulk RNAseq | TCGA | Fishbein et al |
| TCGA-SR-A6MQ-01A | 0.936865577 | -1.591436477 | primary, PF | C2 | bulk RNAseq | TCGA | Fishbein et al |
| TCGA-WB-A80M-01A | 0.815885694 | -0.780588033 | primary, PF | C2 | bulk RNAseq | TCGA | Fishbein et al |
| TCGA-QR-A70W-01A | 1.256280741 | -1.73177948 | primary, PF | C2 | bulk RNAseq | TCGA | Fishbein et al |
| TCGA-RT-A6Y9-01A | -1.356355975 | -1.845781661 | primary, PF | C2 | bulk RNAseq | TCGA | Fishbein et al |
| TCGA-QR-A70T-01A | -1.479275642 | -1.859833572 | primary, PF | C2 | bulk RNAseq | TCGA | Fishbein et al |
| TCGA-QR-A702-01A | 0.357736255 | -2.519859922 | primary, PF | C2 | bulk RNAseq | TCGA | Fishbein et al |
| TCGA-SQ-A6I6-01A | 0.900524918 | -2.260317222 | primary, PF | C2 | bulk RNAseq | TCGA | Fishbein et al |
| TCGA-WB-A820-01A | 2.335706939 | -0.293639735 | primary, PF | C2 | bulk RNAseq | TCGA | Fishbein et al |
| TCGA-TT-A6YO-01A | 2.203324353 | 1.743167243 | primary, PF | C2 | bulk RNAseq | TCGA | Fishbein et al |
| TCGA-WB-A80Y-01A | 0.308124496 | -0.356479012 | primary, PF | C2 | bulk RNAseq | TCGA | Fishbein et al |
| TCGA-WB-A81J-01A | 1.939392959 | -1.827903622 | primary, PF | C2 | bulk RNAseq | TCGA | Fishbein et al |
| TCGA-PR-A5PG-01A | 1.65499968 | -1.644613108 | primary, PF | C2 | bulk RNAseq | TCGA | Fishbein et al |
| TCGA-QR-A70A-01A | -1.485797508 | -1.578764852 | primary, PF | C2 | bulk RNAseq | TCGA | Fishbein et al |
| TCGA-P7-A5NY-05A | 2.414217514 | -0.953525361 | primary, PF | C2 | bulk RNAseq | TCGA | Fishbein et al |
| TCGA-QR-A6GS-01A | 2.866720683 | -1.017042448 | primary, PF | C2 | bulk RNAseq | TCGA | Fishbein et al |
| TCGA-SQ-A6I4-01A | 2.352798828 | 0.333854261 | primary, PF | C2 | bulk RNAseq | TCGA | Fishbein et al |
| TCGA-QR-A707-01A | 0.644111647 | -2.460122936 | primary, PF | C2 | bulk RNAseq | TCGA | Fishbein et al |
| TCGA-WB-A80O-01A | 2.898380207 | 0.278071647 | primary, PF | C2 | bulk RNAseq | TCGA | Fishbein et al |
| TCGA-QR-A70K-01A | 1.705423939 | -1.605376422 | primary, PF | C2 | bulk RNAseq | TCGA | Fishbein et al |
| TCGA-S7-A7WU-01A | -1.128874597 | -1.713576704 | primary, PF | C2 | bulk RNAseq | TCGA | Fishbein et al |
| TCGA-WB-A81P-01A | -0.605422643 | -1.702137503 | primary, PF | C2 | bulk RNAseq | TCGA | Fishbein et al |
| TCGA-QR-A70R-01A | 0.720988826 | -0.667288912 | primary, PF | C2 | bulk RNAseq | TCGA | Fishbein et al |
| TCGA-WB-A81I-01A | -0.977294892 | -1.005939021 | primary, PF | C2 | bulk RNAseq | TCGA | Fishbein et al |
| TCGA-QR-A70O-01A | 0.969805689 | -1.488054694 | primary, PF | C2 | bulk RNAseq | TCGA | Fishbein et al |
| TCGA-W2-A7HD-01A | 0.935202704 | -2.031090062 | primary, PF | C2 | bulk RNAseq | TCGA | Fishbein et al |
| TCGA-WB-A81N-01A | 2.11238045 | -1.960534929 | primary, PF | C2 | bulk RNAseq | TCGA | Fishbein et al |
| TCGA-QR-A6H4-01A | 2.925896029 | -0.616951277 | primary, PF | C2 | bulk RNAseq | TCGA | Fishbein et al |
| TCGA-S7-A7WX-01A | 2.507158144 | -0.234400411 | primary, PF | C2 | bulk RNAseq | TCGA | Fishbein et al |
| TCGA-QR-A708-01A | -0.352759331 | -0.765083047 | primary, PF | C2 | bulk RNAseq | TCGA | Fishbein et al |
| TCGA-SA-A6C2-01A | -2.181173744 | -1.431659374 | primary, PF | C2 | bulk RNAseq | TCGA | Fishbein et al |
| TCGA-S7-A7WM-01A | 2.643547809 | 0.488764298 | primary, PF | C2 | bulk RNAseq | TCGA | Fishbein et al |
| TCGA-SR-A6MY-01A | 2.854540737 | -0.500974501 | primary, PF | C2 | bulk RNAseq | TCGA | Fishbein et al |
| TCGA-RW-A68A-01A | 1.485828062 | -1.766232751 | primary, PF | C2 | bulk RNAseq | TCGA | Fishbein et al |
| TCGA-RW-A689-01A | 2.729846552 | -0.798890072 | primary, PF | C2 | bulk RNAseq | TCGA | Fishbein et al |
| TCGA-RW-A67V-01A | 1.016100159 | -0.95390831 | primary, PF | C2 | bulk RNAseq | TCGA | Fishbein et al |
| TCGA-S7-A7WL-01A | 2.533943813 | -1.3050477 | primary, PF | C2 | bulk RNAseq | TCGA | Fishbein et al |
| TCGA-W2-A7HF-01A | 1.083444797 | -2.363989091 | primary, PF | C2 | bulk RNAseq | TCGA | Fishbein et al |
| TCGA-WB-A819-01A | 2.696371406 | -0.185843087 | primary, PF | C2 | bulk RNAseq | TCGA | Fishbein et al |
| TCGA-WB-A80V-01A | 1.027531507 | -2.401693122 | primary, PF | C2 | bulk RNAseq | TCGA | Fishbein et al |
| TCGA-WB-A81W-01A | 1.922908754 | -1.609710096 | primary, PF | C2 | bulk RNAseq | TCGA | Fishbein et al |
| TCGA-S7-A7WN-01A | 2.079782316 | -1.093262521 | primary, PF | C2 | bulk RNAseq | TCGA | Fishbein et al |
| TCGA-P8-A5KD-01A | 2.161764517 | -0.863764875 | primary, PF | C2 | bulk RNAseq | TCGA | Fishbein et al |
| TCGA-QR-A6GW-01A | 2.181681069 | -1.76816782 | primary, PF | C2 | bulk RNAseq | TCGA | Fishbein et al |
| TCGA-RW-A68G-01A | 2.032044806 | 1.347610548 | primary, PF | C2 | bulk RNAseq | TCGA | Fishbein et al |
| TCGA-RW-A68F-01A | -3.157924179 | -1.99658287 | primary, PF | C2 | bulk RNAseq | TCGA | Fishbein et al |
| TCGA-RW-A67W-01A | 2.027760291 | -0.728229555 | primary, PF | C2 | bulk RNAseq | TCGA | Fishbein et al |
| TCGA-RW-A68B-01A | 1.463184817 | -1.569914497 | primary, PF | C2 | bulk RNAseq | TCGA | Fishbein et al |
| TCGA-QR-A6H1-01A | 1.568334953 | -1.156270314 | primary, PF | C2 | bulk RNAseq | TCGA | Fishbein et al |
| TCGA-SR-A6MS-01A | 0.107706361 | -0.368359528 | primary, PF | C2 | bulk RNAseq | TCGA | Fishbein et al |
| TCGA-RW-A7D0-01A | 2.27359986 | -0.723941899 | primary, PF | C2 | bulk RNAseq | TCGA | Fishbein et al |
| TCGA-TT-A6YJ-01A | 0.713933202 | -2.692693463 | primary, PF | C2 | bulk RNAseq | TCGA | Fishbein et al |
| TCGA-P8-A6RY-01A | -1.133557266 | -1.178255832 | primary, PF | C2 | bulk RNAseq | TCGA | Fishbein et al |
| TCGA-RW-A68D-01A | -1.994600656 | -2.242221403 | primary, PF | C2 | bulk RNAseq | TCGA | Fishbein et al |
| TCGA-QR-A6GO-01A | 2.251723475 | -0.888953362 | primary, PF | C2 | bulk RNAseq | TCGA | Fishbein et al |
| TCGA-RX-A8JQ-01A | 1.945146499 | -1.319746255 | primary, PF | C2 | bulk RNAseq | TCGA | Fishbein et al |
| TCGA-W2-A7H5-01B | 1.483314076 | -1.896308227 | primary, PF | C2 | bulk RNAseq | TCGA | Fishbein et al |
| TCGA-SR-A6MT-01A | -0.841678492 | -1.749736879 | primary, PF | C2 | bulk RNAseq | TCGA | Fishbein et al |
| TCGA-S7-A7WT-01A | 1.153190004 | -0.649885181 | primary, PF | C2 | bulk RNAseq | TCGA | Fishbein et al |
| TCGA-WB-A81R-01A | 1.075431883 | -2.191841759 | primary, PF | C2 | bulk RNAseq | TCGA | Fishbein et al |
| TCGA-QT-A5XN-01A | 1.050818486 | -2.121493013 | primary, PF | C2 | bulk RNAseq | TCGA | Fishbein et al |
| TCGA-WB-A81A-01A | -0.715537594 | -1.561264099 | primary, PF | C2 | bulk RNAseq | TCGA | Fishbein et al |
| TCGA-WB-A816-01A | 2.097303401 | 1.482997929 | primary, PF | C2 | bulk RNAseq | TCGA | Fishbein et al |
| TCGA-QR-A70G-01B | 2.441723342 | 1.332747778 | primary, PF | C2 | bulk RNAseq | TCGA | Fishbein et al |
| TCGA-P7-A5NY-01A | 1.121353257 | -1.900760262 | primary, PF | C2 | bulk RNAseq | TCGA | Fishbein et al |
| TCGA-SP-A6QF-01A | 2.519902689 | -0.42686416 | primary, PF | C2 | bulk RNAseq | TCGA | Fishbein et al |
| TCGA-SP-A6QJ-01A | 2.017201384 | -0.549748381 | primary, PF | C2 | bulk RNAseq | TCGA | Fishbein et al |
| TCGA-RT-A6YA-01A | 3.09369255 | -1.073130537 | primary, PF | C2 | bulk RNAseq | TCGA | Fishbein et al |
| TCGA-WB-A815-01A | -0.195301021 | -0.061247849 | primary, PF | C2 | bulk RNAseq | TCGA | Fishbein et al |
| TCGA-SP-A6QC-01A | 2.767345412 | -0.047281909 | primary, PF | C2 | bulk RNAseq | TCGA | Fishbein et al |
| TCGA-QR-A6GY-01A | 0.431711596 | -0.261689299 | primary, PF | C2 | bulk RNAseq | TCGA | Fishbein et al |
| TCGA-QT-A69Q-01A | 0.683888426 | -0.17839551 | primary, PF | C2 | bulk RNAseq | TCGA | Fishbein et al |
| TCGA-WB-A817-01A | 1.670831812 | 1.413472292 | primary, PF | C2 | bulk RNAseq | TCGA | Fishbein et al |
| TCGA-QR-A70E-01A | -1.906392092 | -2.167564396 | primary, PF | C2 | bulk RNAseq | TCGA | Fishbein et al |
| TCGA-SP-A6QI-01A | 1.833465493 | -1.306459719 | primary, PF | C2 | bulk RNAseq | TCGA | Fishbein et al |
| TCGA-W2-A7HB-01A | 1.076151164 | -2.468244107 | primary, PF | C2 | bulk RNAseq | TCGA | Fishbein et al |
| TCGA-W2-A7HC-01A | 2.699756149 | -0.01870598 | primary, PF | C2 | bulk RNAseq | TCGA | Fishbein et al |
| TCGA-QR-A70U-01A | 1.027308766 | -2.17387594 | primary, PF | C2 | bulk RNAseq | TCGA | Fishbein et al |
| TCGA-WB-A81K-01A | 1.755578055 | 0.97334744 | primary, PF | C2 | bulk RNAseq | TCGA | Fishbein et al |
| TCGA-QR-A700-01A | -0.107998032 | -0.704058756 | primary, PF | C2 | bulk RNAseq | TCGA | Fishbein et al |
| TCGA-QR-A70X-01A | 2.456653168 | 0.954478926 | primary, PF | C2 | bulk RNAseq | TCGA | Fishbein et al |
| TCGA-RT-A6YC-01A | 2.660684725 | 0.310773769 | primary, PF | C2 | bulk RNAseq | TCGA | Fishbein et al |
| TCGA-SR-A6MU-01A | 2.078520688 | 1.216601231 | primary, PF | C2 | bulk RNAseq | TCGA | Fishbein et al |
| TCGA-P7-A5NX-01A | -1.468789062 | -1.898052361 | primary, PF | C2 | bulk RNAseq | TCGA | Fishbein et al |
| TCGA-S7-A7X2-01A | 2.653003651 | -0.692070107 | primary, PF | C2 | bulk RNAseq | TCGA | Fishbein et al |
| TCGA-WB-A81H-01A | 2.499458461 | 1.317122072 | primary, PF | C2 | bulk RNAseq | TCGA | Fishbein et al |
| TCGA-RW-A67Y-01A | 0.874367831 | -2.579626634 | primary, PF | C2 | bulk RNAseq | TCGA | Fishbein et al |
| TCGA-SR-A6MZ-01A | 2.347164391 | 1.453651239 | primary, PF | C2 | bulk RNAseq | TCGA | Fishbein et al |
| TCGA-WB-A818-01A | 2.258612497 | -0.509501556 | primary, PF | C2 | bulk RNAseq | TCGA | Fishbein et al |
| TCGA-W2-A7HA-01B | 2.200839471 | -1.069901297 | primary, PF | C2 | bulk RNAseq | TCGA | Fishbein et al |
| TCGA-XG-A823-01A | 1.882076898 | 1.816677706 | primary, PF | C2 | bulk RNAseq | TCGA | Fishbein et al |
| TCGA-RW-A685-01A | 2.516988759 | 0.727726438 | primary, PF | C2 | bulk RNAseq | TCGA | Fishbein et al |
| TCGA-W2-A7HE-01A | 2.714823367 | 1.22091745 | primary, PF | C2 | bulk RNAseq | TCGA | Fishbein et al |
| TCGA-WB-A81M-01A | 0.810334466 | -1.017546815 | primary, PF | C2 | bulk RNAseq | TCGA | Fishbein et al |
| TCGA-WB-A80N-01A | 2.507341425 | 0.117848319 | primary, PF | C2 | bulk RNAseq | TCGA | Fishbein et al |
| TCGA-QT-A5XM-01A | 2.516139242 | -0.867455804 | primary, PF | C2 | bulk RNAseq | TCGA | Fishbein et al |
| TCGA-QR-A6H2-01A | -0.164315321 | -0.71393808 | primary, PF | C2 | bulk RNAseq | TCGA | Fishbein et al |
| TCGA-WB-A81D-01A | 2.756674481 | -0.357811019 | primary, PF | C2 | bulk RNAseq | TCGA | Fishbein et al |
| TCGA-QT-A5XO-01A | 1.848318826 | 0.596385308 | primary, PF | C2 | bulk RNAseq | TCGA | Fishbein et al |
| TCGA-TT-A6YN-01A | 1.211214433 | -1.476649342 | primary, PF | C2 | bulk RNAseq | TCGA | Fishbein et al |
| TCGA-WB-A81V-01A | 1.611908954 | -0.364495279 | primary, PF | C2 | bulk RNAseq | TCGA | Fishbein et al |
| TCGA-RW-A681-01A | 0.320291315 | -2.543469151 | primary, PF | C2 | bulk RNAseq | TCGA | Fishbein et al |
| TCGA-SP-A6QK-01A | 2.578550249 | 1.254219729 | primary, PF | C2 | bulk RNAseq | TCGA | Fishbein et al |
| TCGA-QR-A70V-01A | 0.528251457 | -2.665162046 | primary, PF | C2 | bulk RNAseq | TCGA | Fishbein et al |
| TCGA-WB-A81F-01A | 1.788225362 | -0.555045487 | primary, PF | C2 | bulk RNAseq | TCGA | Fishbein et al |
| TCGA-S7-A7WR-01A | 3.072912602 | -0.734929246 | primary, PF | C2 | bulk RNAseq | TCGA | Fishbein et al |
| TCGA-WB-A81S-01A | 2.721737167 | -1.096009366 | primary, PF | C2 | bulk RNAseq | TCGA | Fishbein et al |
| TCGA-S7-A7WO-01A | 2.15767032 | 1.665805549 | primary, PF | C2 | bulk RNAseq | TCGA | Fishbein et al |
| TCGA-RW-A686-06A | -0.446400755 | 7.232758473 | primary, PF | C3 | bulk RNAseq | TCGA | Fishbein et al |
| TCGA-QR-A70H-01A | -0.631605038 | 7.410913611 | primary, PF | C3 | bulk RNAseq | TCGA | Fishbein et al |
| TCGA-P8-A5KC-01A | -0.533519021 | 7.34052669 | primary, PF | C3 | bulk RNAseq | TCGA | Fishbein et al |
| TCGA-RW-A8AZ-01A | -0.806742081 | 7.442935479 | primary, PF | C3 | bulk RNAseq | TCGA | Fishbein et al |
| TCGA-RW-A686-01A | -0.614706056 | 7.247725484 | primary, PF | C3 | bulk RNAseq | TCGA | Fishbein et al |
| TCGA-WB-A80K-01A | -0.702035853 | 7.605747264 | primary, PF | C3 | bulk RNAseq | TCGA | Fishbein et al |
| TCGA-QR-A70C-01A | -0.557651209 | 7.384886977 | primary, PF | C3 | bulk RNAseq | TCGA | Fishbein et al |
| TCGA-PR-A5PH-01A | -0.420018925 | 7.113614311 | primary, PF | C3 | bulk RNAseq | TCGA | Fishbein et al |
| TCGA-PR-A5PF-01A | -0.836554988 | 7.470519166 | primary, PF | C3 | bulk RNAseq | TCGA | Fishbein et al |
| TCGA-WB-A80Q-01A | -0.642301906 | 7.370000219 | primary, PF | C3 | bulk RNAseq | TCGA | Fishbein et al |
| 128T | -1.764799196 | 2.202342446 | primary, PF | C1 | bulk RNAseq | UT_BULK | Estrada-Zuniga et al |
| 493_6T | -1.536805857 | 2.115056473 | primary, PF | C1 | bulk RNAseq | UT_BULK | Estrada-Zuniga et al |
| 563T | -1.932762786 | 2.721668036 | primary, PF | C1 | bulk RNAseq | UT_BULK | Estrada-Zuniga et al |
| 1104T | -1.697802047 | 2.407516297 | primary, PF | C1 | bulk RNAseq | UT_BULK | Estrada-Zuniga et al |
| 1130T | -0.011706288 | -0.345439126 | primary, PF | C1 | bulk RNAseq | UT_BULK | Estrada-Zuniga et al |
| 1147T | -3.410401142 | -2.250135439 | primary, PF | C1 | bulk RNAseq | UT_BULK | Estrada-Zuniga et al |
| 1183T | -0.334375273 | -0.446090998 | primary, PF | C1 | bulk RNAseq | UT_BULK | Estrada-Zuniga et al |
| 1195T | -0.500374083 | 0.542616831 | primary, PF | C1 | bulk RNAseq | UT_BULK | Estrada-Zuniga et al |
| 1228T | -1.064119822 | 1.955286443 | primary, PF | C1 | bulk RNAseq | UT_BULK | Estrada-Zuniga et al |
| 1098T | 2.590420223 | 0.149352441 | primary, PF | C2 | bulk RNAseq | UT_BULK | Estrada-Zuniga et al |
| 1100T | 1.918617086 | 1.480328809 | primary, PF | C2 | bulk RNAseq | UT_BULK | Estrada-Zuniga et al |
| 1108T | 0.42035283 | -1.791595101 | primary, PF | C2 | bulk RNAseq | UT_BULK | Estrada-Zuniga et al |
| 1111T | 1.818942697 | -2.297331245 | primary, PF | C2 | bulk RNAseq | UT_BULK | Estrada-Zuniga et al |
| 1119T | 0.8148191 | -2.826255743 | primary, PF | C2 | bulk RNAseq | UT_BULK | Estrada-Zuniga et al |
| 1143T | 1.068679145 | -1.994342762 | primary, PF | C2 | bulk RNAseq | UT_BULK | Estrada-Zuniga et al |
| 1145T | 2.036415114 | 1.309165244 | primary, PF | C2 | bulk RNAseq | UT_BULK | Estrada-Zuniga et al |
| 1157T | 2.085681079 | -1.663134675 | primary, PF | C2 | bulk RNAseq | UT_BULK | Estrada-Zuniga et al |
| 1164T | 2.959470319 | -0.27936662 | primary, PF | C2 | bulk RNAseq | UT_BULK | Estrada-Zuniga et al |
| 1169T | 2.398754978 | -0.493967115 | primary, PF | C2 | bulk RNAseq | UT_BULK | Estrada-Zuniga et al |
| 1194T | 0.613783586 | -0.254567626 | primary, PF | C2 | bulk RNAseq | UT_BULK | Estrada-Zuniga et al |
| 1204T | 2.863260087 | 1.072972557 | primary, PF | C2 | bulk RNAseq | UT_BULK | Estrada-Zuniga et al |
| 1212T | -0.833170586 | -1.891848098 | primary, PF | C2 | bulk RNAseq | UT_BULK | Estrada-Zuniga et al |
| 1219T | 0.514488694 | -0.54297555 | primary, PF | C2 | bulk RNAseq | UT_BULK | Estrada-Zuniga et al |
| 1226T | 0.657568104 | -0.390297328 | primary, PF | C2 | bulk RNAseq | UT_BULK | Estrada-Zuniga et al |
| 1231T | 1.141324404 | -0.219034835 | primary, PF | C2 | bulk RNAseq | UT_BULK | Estrada-Zuniga et al |
| 166T | -0.503885664 | 7.324162325 | primary, PF | C3 | bulk RNAseq | UT_BULK | Estrada-Zuniga et al |
| 1125T | -1.002192054 | 7.494476798 | primary, PF | C3 | bulk RNAseq | UT_BULK | Estrada-Zuniga et al |
| 1190T | -0.561454451 | 7.668578446 | primary, PF | C3 | bulk RNAseq | UT_BULK | Estrada-Zuniga et al |

**Supplementary Table 5.** UMAP of bulk RNAseq and pseudobulk RNAseq of primary pheochromocytomas/paragangliomas and derived organoids: C1=molecular cluster 1, C2= molecular cluster 2, C3=molecular cluster 3, PDO-N= short-term normoxia, PDO-H=short term hypoxia, PDO-L=long term normoxia, one of the samples was prepared in triplicate using three dissociation protocols (D1, D2, D3) and each produced its respective PDO.

**Supplementary Table 6.**

Table provided in synapse.org/PPGLorganoids.

Differentially expressed genes between PDO N (short term normoxia) and PDO L (long term normoxia). UP: (log2FC>2 & FDR <0.05) upregulated genes in long-term, DOWN: (log2FC<-2 & FDR <0.05) downregulated in long-term, NO: not differentially expressed

| Description | setSize | enrichmentScore | NES | pvalue | p.adjust | core_enrichment |
|---|---|---|---|---|---|---|
| HALLMARK_EPITHELIAL_MESENCHYMAL_TRANSITION | 197 | 0.75886379 | 3.179 | 1.00E-10 | 2.50E-09 | GREM1/AREG/TNFRSF11B/SFRP4/POSTN/SFRP1/WNT5A/LAMA1/COL1A1/FBN2/SERPINE2/CXCL6/IL6/PTX3/COL5A1/FBLN1/THBS2/ADAM12/OXTR/VCAN/COL3A1/PLOD2/NTM/COL1A2/COL5A2/SGCD/COL7A1/THBS1/FBLN2/COL12A1/LOX/DKK1/FBN1/FAP/TGFBI/THY1/TFPI2/FSTL1/CTHRC1/LOXL2/LAMA2/INHBA/MMP2/TGM2/BDNF/CCN2/VCAM1/CCN1/LOXL1/NID2/FMOD/EFEMP2/SERPINE1/IGFBP2/LUM/TNC/NT5E/GLIPR1/SPP1/GAS1/IGFBP3/NNMT/COL6A3/TPM2/EDIL3/FN1/COL16A1/LRP1/ACTA2/CALU/CALD1/P3H1/BMP1/BASP1/SPARC/FBLN5/COL6A2/SERPINH1/TAGLN/SPOCK1/LAMC1/MMP3/PCOLCE/CXCL12/ITGB5/FGF2/PCOLCE2/COL4A1/IGFBP4/ANPEP/ABI3BP/QSOX1/VEGFA/VEGFC/GPX7/PTHLH/SLIT2/SAT1/MYL9/PLOD1/TIMP1/FSTL3/ECM2/NOTCH2/COL4A2/FAS/DAB2/ITGA2/MMP14/MFAP5/SNAI2 |
| HALLMARK_HYPOXIA | 187 | 0.55996765 | 2.3213 | 1.00E-10 | 2.50E-09 | CCN5/STC2/HAS1/IL6/COL5A1/PPFIA4/SLC2A5/AK4/LOX/GPC3/TGFBI/ERRFI1/KDELR3/TGM2/P4HA1/CCN2/CA12/STC1/CCN1/FOS/ADORA2B/SERPINE1/IGFBP1/SLC6A6/IGFBP3/PRKCA/SLC2A1/SLC2A3/TPBG/PDK1/HK2/EGFR/ADM/PPP1R3C/EXT1/CHST2/TMEM45A/TGFB3/SRPX/PGM1/BNIP3L/VEGFA/CITED2/DDIT4/HDLBP/ANGPTL4/CAVIN1/PIM1/LXN/ERO1A/WSB1/P4HA2/GYS1/PGK1/DTNA/PLAUR/AKAP12/ENO3/GBE1/DUSP1/TPST2/LDHA/TES/NDRG1/UGP2/CAV1/BHLHE40/ANKZF1/FAM162A/PFKL/XPNPEP1/GLRX/ENO1/HEXA |
| HALLMARK_UV_RESPONSE_DN | 143 | 0.55220781 | 2.2323 | 1.67E-10 | 2.78E-09 | EFEMP1/COL1A1/GCNT1/COL3A1/COL1A2/COL5A2/FHL2/HAS2/BDNF/FZD2/CCN1/ADORA2B/SERPINE1/DDAH1/TFPI/PTGFR/PRKCA/FBLN5/IGFBP5/RUNX1/LPAR1/LAMC1/ABCC1/ADD3/MYC/NEK7/LTBP1/ERBB2/CITED2/RND3/NOTCH2/DAB2/IRS1/SNAI2/CELF2/SMAD3/TGFBR2/GJA1/DUSP1/NRP1/PDGFRB/CAV1/SYNJ2/APBB2/BHLHE40/INSIG1/RBPMS/SLC22A18/ATRN/RASA2 |
| HALLMARK_GLYCOLYSIS | 183 | 0.45898642 | 1.9022 | 2.11E-07 | 2.64E-06 | STC2/COL5A1/VCAN/PPFIA4/PLOD2/AK4/GPC3/TGFBI/SLC16A3/KDELR3/P4HA1/PYGL/STC1/ADORA2B/B4GALT1/NT5E/CHST6/CTH/IGFBP3/TPBG/SPAG4/HK2/EGFR/EXT1/KIF20A/CHST2/DEPDC1/IDUA/QSOX1/VEGFA/CITED2/DDIT4/HDLBP/PLOD1/ANGPTL4/ERO1A/MPI/P4HA2/GYS1/PGK1/PC/XYLT2/CDK1/B4GALT4/PMM2/HMMR/LDHA/UGP2/EGLN3/CD44/ANKZF1/FAM162A/FUT8/HS2ST1/GLRX/ENO1/GLCE/COPB2/CHPF2/B4GALT7/DCN/HOMER1/PGM2 |
| HALLMARK_TNFA_SIGNALING_VIA_NFKB | 190 | 0.43897275 | 1.8361 | 1.11E-06 | 1.11E-05 | AREG/CXCL6/IL6/PTX3/KYNU/ABCA1/INHBA/CCND1/F2RL1/CCL2/CCN1/FOSB/FJX1/SGK1/FOS/B4GALT1/SERPINE1/TNC/DNAJB4/NR4A2/GFPT2/SLC2A3/DRAM1/RCAN1/PLAU/BCL2A1/EGR1/MYC/TNFRSF9/VEGFA/SPHK1/SAT1/PDE4B/EGR2/SMAD3/CLCF1/PLAUR/PHLDA1/EGR3/KLF4/DUSP1/RIPK2/TGIF1/IL6ST/CD44/BHLHE40/ZFP36/LITAF/PTGS2/SLC16A6/CCNL1/TNFSF9/BMP2/PANX1/MCL1/IER2/BIRC2/NFAT5/SOCS3/PTGER4/NR4A3/PTPRE/PHLDA2/SDC4/FUT4 |
| HALLMARK_APICAL_JUNCTION | 178 | 0.38567868 | 1.5807 | 0.00018302 | 0.001525153 | ICAM5/TNFRSF11B/VCAN/FLNC/WNK4/FBN1/TGFBI/THY1/MMP2/VCAM1/ALOX15B/ACTG2/B4GALT1/NEGR1/CERCAM/NEXN/RAC2/ICAM4/ADAM9/COL16A1/CNN2/EGFR/BMP1/CADM3/LIMA1/CLDN11/CLDN7/PARVA/EPB41L2/SLIT2/MYL9/ITGA2/IRS1/CD86/LDLRAP1 |
| HALLMARK_INFLAMMATORY_RESPONSE | 169 | 0.39156649 | 1.611 | 0.00030725 | 0.002194622 | EREG/CCL7/CXCL6/IL6/PDPN/GPC3/ABCA1/INHBA/HAS2/CCL2/AQP9/ADORA2B/HIF1A/SERPINE1/BDKRB1/PCDH7/VIP/FPR1/OSMR/ICAM4/TPBG/IL18R1/ADM/SGMS2/LPAR1/CSF3/CHST2/IFITM1/MYC/DCBLD2/TNFRSF9/SPHK1/AHR/TIMP1/PDE4B/SLC1A2/IL1R1/TLR3/CMKLR1/CD82/PTGER2/MMP14/GP1BA |
| HALLMARK_COAGULATION | 101 | 0.44228168 | 1.7284 | 0.00051661 | 0.003228809 | THBS1/CFH/FBN1/DPP4/TFPI2/MMP2/SERPINE1/F2RL2/PLAT/ADAM9/FN1/LRP1/CTSK/BMP1/SPARC/PLAU/PRSS23/MMP3/CFD/ANXA1/PF4/MST1/P2RY1/RAPGEF3/TIMP1/ITGA2/MMP14/SERPING1/GP1BA/CTSB/HTRA1/CFI/CSRP1 |
| HALLMARK_G2M_CHECKPOINT | 198 | 0.35004233 | 1.4686 | 0.00092948 | 0.005163797 | CCND1/HIF1A/CENPF/DMD/MKI67/ESPL1/H2BC12/NEK2/KIF23/BUB1/MYC/NUSAP1/NDC80/TPX2/TTK/TOP2A/ABL1/TROAP/NOTCH2/CCNB2/KIF11/KNL1/PML/SMAD3/POLQ/PBK/KIF15/CCNA2/CDK1/TFDP1/STIL/CENPE/HMMR/CHEK1/BRCA2/BIRC5/PRC1/PLK1/KIF2C/ORC6/KMT5A/SLC7A1/SMARCC1/E2F3/PTTG1/SLC7A5/HMGA1/GSPT1/MAD2L1/PRIM2/UCK2/CHAF1A/TACC3/PLK4/KPNB1/EXO1 |
| HALLMARK_KRAS_SIGNALING_DN | 117 | -0.46650436 | -1.593 | 0.00120664 | 0.005237729 | DTNB/EPHA5/EFHD1/CAMK1D/P2RX6/DLK2/CPEB3/SNCB/YPEL1/SLC38A3/BARD1/HTR1B/KCND1/CLSTN3/BTG2/GPR3/ITIH3/ACTC1/NR6A1/RYR2/MFSD6/MACROH2A2/MAGIX/CELSR2/SIDT1/GPRC5C/SPTBN2/FGFR3/BMPR1B/RSAD2/MX1/ZC2HC1C/GDNF/TENT5C/NRIP2/PDE6B/MYOT/RYR1/CNTFR/KCNQ2/GPR19/WNT16/SOX10/KCNN1/KLHDC8A/NGB/TFAP2B/PNMT |
| HALLMARK_ALLOGRAFT_REJECTION | 153 | 0.37643556 | 1.524 | 0.00136181 | 0.005237729 | CCL11/EREG/CCL7/IL6/GCNT1/CCL13/GALNT1/THY1/ELANE/INHBA/CCL2/F2R/HIF1A/BCAT1/EGFR/CD7/TGFB2/PF4/IL11/SRGN/STAT1/TIMP1/FAS/TLR3/GPR65/CD86/HLA-G/CD1D/SOCS5/TLR6/IL16/CCND2/FCGR2B/RIPK2/LY75/IL15/FLNA |
| HALLMARK_OXIDATIVE_PHOSPHORYLATION | 200 | -0.40929746 | -1.496 | 0.00118327 | 0.005237729 | NQO2/NDUFS2/SDHB/UQCR10/IDH3B/ATP5MC3/ACAA2/SDHC/COX7A2L/NDUFS4/NDUFA4/NDUFA8/ATP6V1C1/ATP6V1F/ECHS1/POR/SLC25A11/PDHA1/SLC25A5/SLC25A12/HCCS/SLC25A20/ISCA1/ATP5MC1/ATP5MC2/NDUFB1/GPX4/UQCR11/CYC1/NDUFS3/UQCRB/MTX2/ETFB/NDUFS8/ATP6V0E1/NDUFA2/ATP6V1H/POLR2F/VDAC2/COX4I1/PHYH/ATP6V1E1/ATP5PD/HTRA2/NDUFV2/COX5A/UQCRFS1/ATP5PF/SUCLG1/COX7C/ACADSB/ALAS1/SURF1/OGDH/ECH1/ATP5F1D/ACAT1/ISCU/ATP5MG/COX5B/NDUFA6/UQCRQ/CYCS/NDUFB4/ATP6V1D/COX6B1/GOT2/MRPL15/NDUFA3/ECI1/ATP6V1G1/PDP1/NDUFS7/NDUFB3/ATP5ME/IDH3A/NDUFB2/NDUFA9/MRPL34/NDUFS6/TIMM10/NDUFB6/COX7B/NDUFA5/HSD17B10/NDUFC1/MPC1/COX7A2/ATP6V0B/NDUFA7/NDUFB7/NDUFA1/NDUFAB1/ATP6AP1/NDUFC2/COX6C/NDUFB8/COX6A1/ATP6V0C/ATP5F1E/COX8A/SLC25A4/ATP1B1/COX17/MAOB |
| HALLMARK_ESTROGEN_RESPONSE_EARLY | 178 | 0.35921676 | 1.4722 | 0.00130165 | 0.005237729 | CCN5/AREG/STC2/NPY1R/SLC24A3/FOXC1/FHL2/MUC1/TGM2/CCND1/CA12/RAB17/MYOF/FOS/B4GALT1/PAPSS2/GREB1/SLC2A1/TPBG/TSKU/PMAIP1/SLC1A1/PTGES/SLC16A1/RHOBTB3/PRSS23/CXCL12/ADD3/ALDH3B1/IGFBP4/RPS6KA2/MYC/RHOD/CLDN7/HR/OLFML3/OPN3/DHRS3/SLC19A2/MICB/KAZN/WWC1/GJA1/KRT8/EGR3/NBL1/ELOVL5/KLF4/DEPTOR/THSD4/KRT18/IL6ST/CD44/BHLHE40/AFF1/CISH/NADSYN1/SEMA3B/FASN/MYBBP1A |
| HALLMARK_MTORC1_SIGNALING | 199 | 0.33846596 | 1.4252 | 0.00155454 | 0.005551931 | PLOD2/AK4/P4HA1/STC1/SLC6A6/CTH/SLC2A1/SLC2A3/BCAT1/PDK1/HK2/PSAT1/SERPINH1/IGFBP5/PHGDH/ADD3/SCD/BUB1/CCNG1/PGM1/SLC1A5/DDIT4/SYTL2/CTSC/FGL2/RRM2/ERO1A/NUPR1/ELOVL6/PGK1/GBE1/ELOVL5/LDHA/TES/EGLN3/ACACA/BHLHE40/PLK1/USO1/PFKL/INSIG1/GLRX/FADS2/ENO1/UFM1/ACSL3/SERP1/SKAP2/SORD/TFRC/SLC7A5/LTA4H/FADS1/POLR3G/STARD4/PSPH/SHMT2 |
| HALLMARK_PANCREAS_BETA_CELLS | 25 | -0.6477656 | -1.721 | 0.00180014 | 0.006000479 | DCX/ABCC8/LMO2/INSM1/PAK3/PCSK2/CHGA/ISL1/SST |
| HALLMARK_UNFOLDED_PROTEIN_RESPONSE | 113 | 0.40094657 | 1.5984 | 0.00334847 | 0.010463977 | STC2/KDELR3/CCL2/NABP1/IGFBP1/FKBP14/PSAT1/WIPI1/ALDH18A1/VEGFA/DDIT4/PDIA5/TTC37/ERO1A/CHAC1/IARS1/ARFGAP1/XPOT/SEC31A/EEF2 |
| HALLMARK_MYOGENESIS | 165 | 0.35087167 | 1.4385 | 0.00417317 | 0.012274039 | STC2/COL1A1/IGF1/PTGIS/ADAM12/COL3A1/PPFIA4/BDKRB2/SGCD/COL15A1/LAMA2/AEBP1/IGFBP3/COL6A3/TPM2/PPP1R3C/SPARC/COL6A2/DMD/TAGLN/CDH13/IGFBP7/SSPN/FST/ITGB5/CTF1/FGF2/SCD/CFD/NOS1/MYH3/SPHK1/COL4A2/PC/DTNA/ENO3/CNN3/MYH11/DMPK |
| HALLMARK_ANGIOGENESIS | 32 | 0.56264718 | 1.7258 | 0.00475612 | 0.013211452 | POSTN/CXCL6/VCAN/COL3A1/COL5A2/FSTL1/STC1/LUM/SPP1/PF4/VEGFA/TNFRSF21/TIMP1 |
| HALLMARK_SPERMATOGENESIS | 87 | -0.44294759 | -1.456 | 0.00889456 | 0.0222364 | CLPB/STRBP/CHRM4/ACRBP/ACE/CLVS1/ZC2HC1C/ADCYAP1/MAP7/PCSK1N/SHE/CAMK4/CLGN/SCG3/SNAP91/PACRG/SCG5/NEFH/DDX25/IL13RA2/CNIH2 |
| HALLMARK_MITOTIC_SPINDLE | 199 | 0.31242752 | 1.3156 | 0.00847989 | 0.0222364 | CENPF/ESPL1/ANLN/NEK2/DLGAP5/KIF23/ECT2/BUB1/NUSAP1/NDC80/TPX2/PIF1/EPB41L2/TTK/TOP2A/ABL1/NOTCH2/CCNB2/CDK5RAP2/ITSN1/KIF11/KIF15/CDK1/MYO1E/SHROOM2/CENPE/ARFIP2/NEDD9/BRCA2/ARHGAP5/BIRC5/KNTC1/FLNA/PRC1/PLK1/PKD2/TRIO/PALLD/KIF2C/VCL/RASA2/ACTN4/MYH10/CEP192/KIF3B/SOS1/FLNB/DST/PLEKHG2/FGD4/CTTN/ARHGEF2/CDC42EP1/CLIP2/LLGL1/CAPZB/STAU1/MYO9B/MYH9/FARP1/CLIP1/CEP250/PDLIM5/ARHGAP10/GEMIN4/CDC27 |
| HALLMARK_HEDGEHOG_SIGNALING | 32 | -0.56134243 | -1.577 | 0.01455462 | 0.034653856 | HEY2/CDK5R1/GLI1/NRCAM/HEY1/UNC5C/SCG2/ADGRG1/ACHE/CRMP1/RTN1/AMOT/L1CAM/CNTFR/CELSR1 |
| HALLMARK_APOPTOSIS | 150 | 0.32547788 | 1.3198 | 0.03084271 | 0.07009707 | EREG/IL6/MMP2/CCND1/F2R/LUM/CTH/HGF/PLAT/TIMP2/IGFBP6/PMAIP1/TGFB2/CASP4/SLC20A1/ANXA1/CASP1/BNIP3L/ERBB2/ANKH/TOP2A/SAT1/TIMP1/FAS/IFITM3/EGR3/DPYD/BID/PDGFRB/CCND2/TXNIP/NEDD9/CAV1/KRT18/CD44/PLPPR4 |
| HALLMARK_UV_RESPONSE_UP | 147 | -0.38027049 | -1.344 | 0.03452685 | 0.075058379 | HYAL2/CHKA/PRKACA/DNAJB1/SQSTM1/CDKN1C/E2F5/CA2/CDC34/CEBPG/AQP3/FKBP4/SPR/BCL2L11/DGAT1/RRAD/ENO2/HLA-F/EPHX1/TUBA4A/NFKBIA/BTG2/SOD2/SLC25A4/NPTXR/MSX1/RASGRP1/KCNH2/PLCL1/GPX3/ABCB1/GAL/HMOX1/OLFM1/CHRNA5/LYN/NPTX2/RET/MAPK8IP2/GCH1/EPCAM/CYP1A1 |
| HALLMARK_IL2_STAT5_SIGNALING | 180 | 0.29827653 | 1.2259 | 0.0409732 | 0.085360826 | IL1RL1/MUC1/TGM2/P4HA1/NT5E/COL6A1/SPP1/F2RL2/SLC2A3/WLS/IL18R1/HK2/PLAGL1/TNFRSF18/GALM/MYC/TNFRSF9/SLC1A5/IKZF2/TNFRSF21/DHRS3/AHR/TTC39B/CCNE1/FGL2/PIM1/GPR65/CKAP4/PTGER2/CD86/TWSG1/ETV4/IFITM3/ENO3/MYO1E/PHLDA1/HIPK2/NRP1/CCND2/NCOA3/PLSCR1/NDRG1/PHTF2/CD44/BHLHE40/SLC39A8/ADAM19 |
| HALLMARK_E2F_TARGETS | 200 | 0.290065 | 1.2188 | 0.0430124 | 0.086024794 | DIAPH3/LUC7L3/MKI67/ESPL1/E2F8/MELK/CHEK2/DEPDC1/DLGAP5/MYC/BUB1B/TOP2A/CCNE1/CCNB2/RRM2/CDK1/SPAG5/CENPE/TK1/KIF18B/HMMR/CHEK1/BRCA2/BIRC5/PLK1/WDR90/SHMT1/KIF2C/CTPS1/ORC6/GINS4/SPC24/ATAD2/PTTG1/PRDX4/TFRC/GINS1/HMGA1/PSMC3IP/PAICS/WEE1/GSPT1/MAD2L1/IPO7/PRIM2/PRKDC/TACC3/TRIP13/PLK4/DONSON/DSCC1/PNN/NCAPD2/BRCA1/NAP1L1 |
| HALLMARK_XENOBIOTIC_METABOLISM | 165 | 0.28613769 | 1.1731 | 0.11792453 | 0.226777939 | PTGDS/AOX1/FBLN1/IGF1/KYNU/FMO3/HSD11B1/AQP9/SERPINE1/PAPSS2/IGFBP1/SLC6A6/CDA/BCAT1/PTGES/PYCR1/TGFB2/ADH1C/IGFBP4/VNN1/SMOX/SLC1A5/LCAT/IL1R1/PDK4/FAS/SSR3/PC/SLC35D1/ELOVL5/MAN1A1/CSAD |
| HALLMARK_COMPLEMENT | 170 | 0.27486045 | 1.1301 | 0.16267943 | 0.301258196 | IL6/KYNU/PCSK9/CFH/DPP4/TFPI2/SERPINE1/PLAT/ADAM9/FN1/LRP1/TIMP2/CDA/CDH13/CASP4/CASP1/TIMP1/CTSC/PHEX/COL4A2/PIM1/MMP14/SERPING1/GP1BA/GNB4/CPM/CTSB/PLAUR/ZEB1/CSRP1/PLA2G4A |
| HALLMARK_TGF_BETA_SIGNALING | 51 | 0.33358423 | 1.1511 | 0.2191358 | 0.353444843 | THBS1/SERPINE1/LTBP2/SLC20A1/SMAD3/HIPK2/ENG/TGIF1/SMURF2/SKI/MAP3K7/BMP2 |
| HALLMARK_KRAS_SIGNALING_UP | 179 | -0.31309605 | -1.133 | 0.20797011 | 0.353444843 | PTPRR/FLT4/ADAM8/VWA5A/MMP11/CTSS/MPZL2/SOX9/PECAM1/GUCY1A1/CFB/ID2/ANO1/TMEM100/PEG3/TSPAN1/SNAP25/TMEM176A/BTBD3/FUCA1/ST6GAL1/CXCR4/ADAMDEC1/ACE/IL10RA/RELN/HDAC9/EPB41L3/TMEM176B/ITGB2/LAT2/ABCB1/GPRC5B/MAP7/PCSK1N/MMP9/KIF5C/SCG3/MMP10/EPHB2/TSPAN13/SNAP91/SCG5/IGF2/CSF2RA/IL2RG/BIRC3/MYCN/TSPAN7/NAP1L2/IKZF1/PLVAP/FGF9 |
| HALLMARK_HEME_METABOLISM | 172 | -0.31541534 | -1.131 | 0.21799747 | 0.353444843 | UCP2/CCND3/RAD23A/FBXO7/DCAF11/BLVRB/CLIC2/SNCA/VEZF1/CA2/KAT2B/EPB41/AQP3/H1-0/SLC7A11/IGSF3/BTG2/HAGH/ABCG2/RBM38/PRDX2/ADD2/ATP6V0A1/OSBP2/BCAM/FN3K/TENT5C/ACP5/RAP1GAP/SLC22A4/TAL1/ACSL6/XK/TRIM58/LMO2/SPTB/TMCC2/ANK1/C3/DMTN |
| HALLMARK_PROTEIN_SECRETION | 94 | 0.28884321 | 1.1161 | 0.21455939 | 0.353444843 | ABCA1/EGFR/SSPN/LMAN1/BNIP3/SEC22B/CTSC/CAV2/BET1/KRT18/SEC31A/ADAM10/USO1/SEC24D/ATP6V1B1/COPB2/COPB1 |
| HALLMARK_IL6_JAK_STAT3_SIGNALING | 76 | 0.30298086 | 1.1089 | 0.23843416 | 0.372553381 | CCL7/IL6/PDGFC/OSMR/IL18R1/INHBE/PF4/STAT1/TNFRSF21/ITGA4/IL1R1/FAS/PIM1 |
| HALLMARK_NOTCH_SIGNALING | 31 | 0.36707309 | 1.1199 | 0.27988338 | 0.42406573 | WNT5A/CCND1/PRKCA/NOTCH2/PPARD/MAML2/KAT2A |
| HALLMARK_WNT_BETA_CATENIN_SIGNALING | 40 | -0.38378754 | -1.107 | 0.30630631 | 0.424593778 | HDAC5/JAG1/HDAC11/FZD8/HEY2/NOTCH4/LEF1/JAG2/HEY1/PSEN2/NKD1/TCF7/WNT6 |
| HALLMARK_APICAL_SURFACE | 37 | -0.38569093 | -1.1 | 0.3141994 | 0.424593778 | IL2RB/CX3CL1/MDGA1/SLC2A4/EFNA5/NTNG1/MAL/LYN/IL2RG/SCUBE1/GATA3 |
| HALLMARK_ESTROGEN_RESPONSE_LATE | 172 | -0.30290961 | -1.086 | 0.29024081 | 0.424593778 | CA2/DCXR/CYP26B1/SLC2A8/HPRT1/PLXNB1/FKBP4/ST14/PLAAT3/FAM102A/TRIM29/PGR/SLC9A3R1/PTGER3/SLC1A4/COX6C/ID2/BAG1/LSR/SLC29A1/FABP5/HOMER2/CD9/BCL2/ABCA3/CELSR2/CACNA2D2/PTPN6/SNX10/ITPK1/PRLR/ISG20/FGFR3/GAL/MDK/OLFM1/BATF/TSPAN13/LLGL2/MAPT/CKB/TIAM1/SFN/ST6GALNAC2/DHRS2/RET/SERPINA5/TH/CHST8/KRT19/ASCL1/PDZK1 |
| HALLMARK_BILE_ACID_METABOLISM | 90 | 0.28192826 | 1.0729 | 0.30078125 | 0.424593778 | DIO2/ABCA9/ABCA1/ABCA8/AQP9/CH25H/ABCA6/HSD17B6 |
| HALLMARK_INTERFERON_GAMMA_RESPONSE | 186 | -0.27912627 | -1.016 | 0.42521632 | 0.559495153 | PNP/PSME1/ISG15/IRF9/IRF7/STAT4/HLA-A/HLA-DMA/SOCS1/BATF2/ARL4A/CD274/PSME2/AUTS2/IFIT3/TDRD7/IL2RB/SECTM1/HLA-B/HLA-DQA1/NFKBIA/CFB/SOD2/IRF5/VAMP8/EIF4E3/IL10RA/PTPN6/OASL/CMPK2/BANK1/ISG20/LYSMD2/IFI30/CD74/RSAD2/HLA-DRB1/MX1/IL18BP/GBP4/IDO1/CSF2RB/CIITA/TRIM14/IFI27/P2RY14/GCH1/SLAMF7 |
| HALLMARK_INTERFERON_ALPHA_RESPONSE | 95 | -0.29252942 | -0.97 | 0.54374159 | 0.6971046 | PSME1/ISG15/IRF9/IRF7/LAMP3/BATF2/PSME2/IFIT3/TDRD7/HLA-C/TENT5A/OASL/CMPK2/ISG20/IFI30/CD74/RSAD2/MX1/GBP4/CCRL2/TRIM14/IFI27 |
| HALLMARK_ANDROGEN_RESPONSE | 93 | 0.24660047 | 0.9453 | 0.578125 | 0.72265625 | CCND1/SGK1/B4GALT1/SCD/LMAN1/ANKH/SAT1/PGM3/ARID5B/CENPN/AKAP12/KRT8/ELOVL5/SLC38A2/UAP1/NDRG1/INSIG1/SEC24D/MYL12A/ACTN1/ACSL3/SORD/ADAMTS1/ABCC4/PLPP1 |
| HALLMARK_PI3K_AKT_MTOR_SIGNALING | 97 | -0.28083045 | -0.935 | 0.61898396 | 0.754858484 | SQSTM1/CAB39L/ARHGDIA/VAV3/ECSIT/PIN1/CDKN1B/HRAS/RPS6KA1/CXCR4/ADCY2/CAMK4/PRKCB/IL2RG/TIAM1/SFN/LCK |
| HALLMARK_P53_PATHWAY | 193 | 0.22232359 | 0.9331 | 0.67326733 | 0.801508722 | F2R/ALOX15B/FOS/KRT17/ZMAT3/DRAM1/EPS8L2/CDH13/CASP1/PTPN14/CCNG1/POLH/CDKN2B/SPHK1/DDIT4/SAT1/FAS/CD82/ERCC5/PIDD1/NUPR1/SLC19A2/TRAF4/ADA/VDR/SLC35D1/KLF4/APAF1/CCND2/TXNIP/TM4SF1/MDM2/PLXNB2/NDRG1/FAM162A/DGKA/LDHB/DDB2/TRIAP1/TNFSF9/CCNK/RACK1/BMP2/SEC61A1 |
| HALLMARK_FATTY_ACID_METABOLISM | 141 | -0.25748706 | -0.905 | 0.69857697 | 0.812298806 | SERINC1/SUCLG1/H2AZ1/HSD17B11/GSTZ1/HSD17B7/ECH1/GCDH/CBR3/SLC22A5/ACAT2/ECI1/GLUL/PSME1/NCAPH2/CA2/ACSS1/HSP90AA1/HSD17B10/CEL/ACSL5/AOC3/BPHL/ENO2/PTS/EPHX1/S100A10/YWHAH/PCBD1/REEP6/BMPR1B/ODC1/IL4I1/TDO2/CA4/CYP1A1 |
| HALLMARK_MYC_TARGETS_V2 | 57 | 0.2441521 | 0.8539 | 0.75709779 | 0.8603384 | HK2/MYC/PLK1/MYBBP1A/SORD/TFB2M/GNL3/WDR43/PLK4/PRMT3/DUSP2/IPO4/NPM1/SLC19A1/EXOSC5/SRM/NOLC1 |
| HALLMARK_REACTIVE_OXYGEN_SPECIES_PATHWAY | 47 | -0.25332508 | -0.748 | 0.86806597 | 0.964517741 | NDUFA6/NDUFB4/MBP/SOD1/SRXN1/MSRA/NQO1/SOD2/PRDX2/CDKN2D/GPX3/HHEX |
| HALLMARK_CHOLESTEROL_HOMEOSTASIS | 73 | -0.23341174 | -0.747 | 0.88981869 | 0.967194227 | ALCAM/MVK/EBP/FABP5/CD9/PMVK/ATF5/CLU/LPL/AVPR1A/MAL2/TM7SF2 |
| HALLMARK_PEROXISOME | 94 | -0.21882503 | -0.724 | 0.9365722 | 0.996353404 | ECH1/SLC23A2/ESR2/SLC25A19/PEX11B/SOD1/CRABP1/CLN6/FIS1/PRDX5/CEL/ACSL5/HRAS/CACNA1B/SOD2/SLC25A4/YWHAH/ABCC8/ABCB1/RXRG/EPHX2/ABCD2/CADM1 |
| HALLMARK_DNA_REPAIR | 149 | -0.17552518 | -0.621 | 0.99744246 | 1 | RAE1/GPX4/APRT/RFC2/GTF2A2/ADRM1/NT5C/POLR2F/POLR3GL/SUPT4H1/VPS28/SURF1/NT5C3A/CETN2/POLB/POLR2K/RBX1/RFC3/NUDT9/EIF1B/MPC2/NELFE/DGUOK/PNP/STX3/HCLS1/PRIM1/HPRT1/SAC3D1/ALYREF/ARL6IP1/AK1/POLR2I/RPA3/USP11/BCAP31/NELFCD/BOLA2/CMPK2/BCAM/CCNO/COX17 |
| HALLMARK_MYC_TARGETS_V1 | 199 | 0.145594 | 0.6131 | 1 | 1 | MYC/PABPC1/PABPC4/PGK1/CCNA2/IARS1/TFDP1/TYMS/SERBP1/LDHA/XPOT/CTPS1/NCBP2/RPS6/SMARCC1/IMPDH2/PRDX4/RACK1/SF3B3/GSPT1/CAD/SRPK1/MAD2L1/GNL3/RPL6/EPRS1/KPNB1/NAP1L1/RSL1D1/RPL22/EIF4A1/ABCE1/RPL34/RPLP0/FAM120A/RPS10/EIF3D/NPM1/ETF1/RPS2/AP3S1/RPS5/SSB |
| HALLMARK_ADIPOGENESIS | 191 | -0.15961707 | -0.581 | 1 | 1 | CHCHD10/GHITM/UCP2/UQCRQ/MRPL15/ALDH2/SOD1/HSPB8/REEP5/PEMT/SPARCL1/IDH3A/ESYT1/DGAT1/ARL4A/COX7B/NDUFA5/SORBS1/ENPP2/SLC25A10/NKIRAS1/DHRS7/NDUFB7/COQ3/NDUFAB1/PTGER3/COX6A1/PIM3/COX8A/REEP6/CYP4B1/LPL/GPX3/EPHX2/C3/QDPR |

**Supplementary Table 7.** Gene Set Enrichment Analysis (GSEA): pathway enrichment of the comparison between PDO N (short term) and PDO L (long term). Key: NES=normalized enrichment score, positive values are enriched in the ‘Long term’ group; Core enrichment: the subset of genes that contributes most to the enrichment result.

| Deconvolution using ESTIMATE | | | |
|---|---|---|---|
| Sample_ID | StromalScore | ImmuneScore | ESTIMATEScore |
| 1302_H | -1532.342973 | -1665.372714 | -3197.715687 |
| 1302_N | -1285.255801 | -1567.479444 | -2852.735245 |
| 1337_N | -1153.000911 | -1130.178172 | -2283.179082 |
| 1336_N | -704.9626824 | -1353.748401 | -2058.711084 |
| 1311_H | -439.3413855 | -1137.646936 | -1576.988322 |
| 1311_N | -339.7756671 | -1128.457003 | -1468.23267 |
| 1311_H5 | -178.809499 | -1046.085088 | -1224.894587 |
| 1313_H | -52.29400621 | -1098.591347 | -1150.885353 |
| 1313_N | -74.28104227 | -976.8924762 | -1051.173518 |
| 1322_H | -256.1760399 | -790.7500985 | -1046.926138 |
| 1312_L | 152.9413422 | -1146.620158 | -993.6788154 |
| 1312_H | -275.5543804 | -656.4738868 | -932.0282672 |
| 1231_N | 11.53287659 | -917.9457998 | -906.4129232 |
| 1322_N | -130.1809553 | -766.2600766 | -896.4410319 |
| 1313_L | 304.0458818 | -1188.06617 | -884.0202881 |
| 1337_L | 249.8745388 | -1117.41885 | -867.5443114 |
| 1335_N | -233.0475408 | -632.3652227 | -865.4127635 |
| 1322_L | 250.7810977 | -1094.294469 | -843.5133716 |
| 1301_N | 147.1814942 | -976.7155741 | -829.5340799 |
| 1335_L | 358.3478186 | -1173.047011 | -814.6991927 |
| 1231_L | 320.3516467 | -1113.319792 | -792.9681451 |
| 1231_D | -377.5952966 | -351.3045746 | -728.8998712 |
| 1251_N | -22.53356679 | -652.6464465 | -675.1800133 |
| 1314_N | 377.2322588 | -1041.176172 | -663.9439129 |
| 1251_L | 369.4743563 | -947.4817076 | -578.0073513 |
| 1312_N | 79.61833471 | -290.4820584 | -210.8637237 |
| 1306_L | 116.556487 | -218.0472588 | -101.4907718 |
| 1343_L | 588.4773123 | -635.1863366 | -46.70902439 |
| 1313_D | 458.8307274 | -418.1241226 | 40.70660476 |
| 1312_D | 319.0585073 | -253.3619964 | 65.69651088 |
| 1343_H | 199.2809289 | -49.51388737 | 149.7670415 |
| 1343_N | 243.3604279 | 112.1428735 | 355.5033014 |
| 1343_D | 877.158018 | -2.884667575 | 874.2733504 |
| 1244_D | 741.9924346 | 371.3620394 | 1113.354474 |

| Deconvolution using MuSiC | | | | | | | | | | | | | |
|---|---|---|---|---|---|---|---|---|---|---|---|---|---|
| Sample ID | c1:Tumor | c0:Tumor | c4:Tumor | c3:Macropha | c2:Endotheli | c5:SCPs | c9:Unknown | c11:Endothel | c8:Immune | c6:Adrenoco | c7:Fibroblas | c10:Tumor | c12:Endothelial |
| 1302_H | 0.4473265306 | 0 | 0 | 0 | 0 | 0 | 0 | 0.0331007551 | 0.3588511293 | 0.0510696769 | 0.109651908 | 0 | 0 |
| 1302_N | 0.3693254133 | 0 | 0 | 0 | 0 | 0 | 0 | 0.1484766022 | 0.3068389767 | 0.0297598737 | 0.1455991341 | 0 | 0 |
| 1337_N | 0.4014048729 | 0 | 0 | 0 | 0.0078737021 | 0 | 0 | 0.0432714628 | 0.3850681507 | 0.0253463193 | 0.1370354921 | 0 | 0 |
| 1336_N | 0.2255540196 | 0 | 0 | 0 | 0.0169000686 | 0 | 0 | 0.0883850762 | 0.3659047932 | 0.0357963317 | 0.2674597107 | 0 | 0 |
| 1311_H | 0.0211963905 | 0 | 0 | 0 | 0.0126992799 | 0 | 0 | 0.0188990168 | 0.432851562 | 0.0162149662 | 0.4981387844 | 0 | 0 |
| 1311_N | 0.0362863676 | 0 | 0 | 0 | 0.0281374428 | 0 | 0 | 0.0271261430 | 0.4102161097 | 0.0231835581 | 0.4750503788 | 0 | 0 |
| 1311_H5 | 0 | 0 | 0 | 0 | 0.0275479189 | 0 | 0 | 0.0056553182 | 0.442649433 | 0.0191548716 | 0.5049924581 | 0 | 0 |
| 1313_H | 0.0044660730 | 0 | 0 | 0 | 0.0788396689 | 0 | 0 | 0.0920031186 | 0.2687782599 | 0.0080154655 | 0.5469860674 | 0 | 0.000911346445 |
| 1313_N | 0.0488658785 | 0 | 0 | 0.0118179089 | 0.0848669747 | 0 | 0 | 0.0898795688 | 0.2794466773 | 0.0155342914 | 0.4691847412 | 0 | 0.000403959112 |
| 1322_H | 0.0061487782 | 0 | 0 | 0.0355291222 | 0.0555516895 | 0 | 0 | 0.1042589862 | 0.3014836834 | 0.0117387600 | 0.4847655893 | 0 | 0.000523390969 |
| 1312_L | 0 | 0 | 0 | 0.0219895615 | 0.0108012811 | 0 | 0 | 0.0258781326 | 0.2440997277 | 0.0185003765 | 0.6777054851 | 0 | 0.001025435429 |
| 1312_H | 0.2056515498 | 0 | 0 | 0.0172584876 | 0.0725114521 | 0 | 0 | 0.1479519382 | 0.2745697455 | 0.0194466401 | 0.2619841356 | 7.47E-05 | 0.000551337476 |
| 1231_N | 0.0863408927 | 0 | 0 | 0.0137334530 | 0.0312243142 | 0 | 0.0022877912 | 0.1240643972 | 0.3216740704 | 0.0289267291 | 0.3917483521 | 0 | 0 |
| 1322_N | 0 | 0 | 0 | 0.0386401386 | 0.0607808773 | 0 | 0 | 0.0415264639 | 0.3216972245 | 0.0195644944 | 0.5171437923 | 0 | 0.000647008844 |
| 1313_L | 0 | 0 | 0 | 0.0127640117 | 0.0616926746 | 0 | 0 | 0.0476236437 | 0.2401411436 | 0.0123524193 | 0.623972125 | 0 | 0.001453981948 |
| 1337_L | 0.0119037820 | 0 | 0 | 0.0172872290 | 0.0252313620 | 0 | 0 | 0.0595112638 | 0.2584812727 | 0.0284768941 | 0.5991081962 | 0 | 0 |
| 1335_N | 0.0808129115 | 0 | 0 | 0 | 0.0652420775 | 0 | 0 | 0.0438185943 | 0.4478014814 | 0.0275619750 | 0.3347629601 | 0 | 0 |
| 1322_L | 0 | 0 | 0 | 0.0333885424 | 0.0340994285 | 0 | 0 | 0.0007754620 | 0.3046664044 | 0.0164220280 | 0.6106481345 | 0 | 0 |
| 1301_N | 0 | 0 | 0 | 4.55E-05 | 0.0638089181 | 0 | 0 | 0.0438778216 | 0.2811478509 | 0.0087790441 | 0.601526456 | 0 | 0.000814436311 |
| 1335_L | 0 | 0 | 0 | 0.0200691061 | 0.0251750880 | 4.07E-05 | 0 | 0.0345829553 | 0.2234685201 | 0.0208642244 | 0.6757994542 | 0 | 0 |
| 1231_L | 0 | 0 | 0 | 0.0310380859 | 0.0179789909 | 0 | 0 | 0.0064414421 | 0.2796288669 | 0.023691629 | 0.6412209851 | 0 | 0 |
| 1231_D | 0.2087479678 | 0 | 0 | 0 | 0.1478134837 | 0 | 0 | 0.1618252961 | 0.3137623738 | 0.0175869238 | 0.1502639547 | 0 | 0 |
| 1251_N | 0 | 0 | 0 | 0.0312848860 | 0.0870396801 | 0 | 0 | 0.0584096848 | 0.2975556962 | 0.0182574990 | 0.5067430602 | 0 | 0.000709493406 |
| 1314_N | 0 | 0 | 0 | 0.0361749417 | 0.0431423019 | 0 | 0 | 0 | 0.2482456336 | 0.0145447017 | 0.6575621058 | 0 | 0.000330315045 |
| 1251_L | 0 | 0 | 0 | 0.0369684136 | 0.0135858862 | 0 | 0 | 0 | 0.2553770286 | 0.0284540974 | 0.665505718 | 0 | 0.000108856120 |
| 1312_N | 0.0046862126 | 0 | 0 | 0.0345346724 | 0.0726986417 | 0 | 0 | 0.0879582237 | 0.3010804995 | 0.0113480927 | 0.4876936572 | 0 | 0 |
| 1306_L | 0 | 0 | 0 | 0 | 0 | 0 | 0 | 0 | 0.4360659335 | 0.0053806039 | 0.5585534625 | 0 | 0 |
| 1343_L | 0 | 0 | 0 | 0.0541678310 | 0.0328809791 | 0 | 0 | 0.0023538365 | 0.2784019775 | 0.0144294901 | 0.6171716513 | 0 | 0.000594234380 |
| 1313_D | 0 | 0 | 0 | 0 | 0.1439339852 | 0 | 0 | 0.0013454574 | 0.2889585263 | 0 | 0.5657620311 | 0 | 0 |
| 1312_D | 0 | 0 | 0 | 0.0126301737 | 0.4779222477 | 0 | 0 | 0.0257602694 | 0.1881065052 | 0.0024599937 | 0.2931208101 | 0 | 0 |
| 1343_H | 0 | 0 | 0 | 0.0706649558 | 0.0749871658 | 0 | 0 | 0.0362955224 | 0.2902850549 | 0.0139659996 | 0.5130962794 | 0 | 0.000705021855 |
| 1343_N | 0 | 0 | 0 | 0.0728820676 | 0.0627542503 | 0 | 0 | 0.0423480248 | 0.3017573476 | 0.0147779244 | 0.5046334778 | 0 | 0.000846907318 |
| 1343_D | 0 | 0 | 0 | 0.0255792033 | 0.4086456826 | 0 | 0 | 0.0015868571 | 0.1738488136 | 0.0006748011 | 0.3896646422 | 0 | 0 |
| 1244_D | 0.0003183245 | 0 | 0 | 0.0328511514 | 0.2170645794 | 0 | 0 | 0.0998567513 | 0.3285386979 | 0.0646305251 | 0.2567399702 | 0 | 0 |

**Supplementary Table 8.** In silico deconvolution scores of bulk RNAseq using two separate methods, ESTIMATE (Estimation of STromal and Immune cells in MAlignant Tumor tissues using Expression data, source https://www.aging-us.com/article/101415/text) and MuSiC (MUlti-Subject SIngle Cell deconvolution- source: https://www.nature.com/articles/s41467-018-08023-x).

| Sample ID | Genotype | Seq Type | Sample Type | Cluster | Batch | Total cells | Cells after QC | Genes before filter | Genes after filter | Doublets before ambient RNA correction | Doublets after ambient RNA correction | Reference |
|---|---|---|---|---|---|---|---|---|---|---|---|---|
| 0393 | TMEM127 | snRNA-seq | PT | C2 | Batch1 | 8397 | 6774 | 30713 | 23522 | 1064 | 1031 | https://pmc.ncbi.nlm.nih.gov/articles/PMC10637630/ |
| 1119 | TMEM127 | snRNA-seq | PT | C2 | Batch1 | 9702 | 7559 | 30333 | 22860 | 1114 | 1437 | https://pmc.ncbi.nlm.nih.gov/articles/PMC10637630/ |
| 1182 | RET | snRNA-seq | PT | C2 | Batch1 | 9573 | 7937 | 29763 | 21716 | 1360 | 1142 | https://pmc.ncbi.nlm.nih.gov/articles/PMC10637630/ |
| 1196 | RET | snRNA-seq | PT | C2 | Batch1 | 7130 | 5962 | 29137 | 20618 | 862 | 678 | https://pmc.ncbi.nlm.nih.gov/articles/PMC10637630/ |
| 0802 | TMEM127 | snRNA-seq | PT | C2 | Batch1 | 5843 | 4466 | 27923 | 18751 | 574 | 504 | https://pmc.ncbi.nlm.nih.gov/articles/PMC10637630/ |
| 1195 | EPAS1 | snRNA-seq | PT | C1B | Batch2 | 10460 | 8829 | 29868 | 22437 | 1058 | 879 | this manuscript |
| 1244 | VHL | snRNA-seq | PT | C1B | Batch2 | 8713 | 7531 | 30289 | 22633 | 1147 | 779 | this manuscript |
| 1259 | EPAS1 | snRNA-seq | PT | C1B | Batch2 | 7627 | 5274 | 28817 | 20467 | 977 | 852 | this manuscript |
| 1312_D | NF1 | snRNA-seq | PDT | C2 | Batch2 | 6549 | 4648 | 29104 | 20722 | 654 | 316 | this manuscript |
| 1313_D | Unknown | snRNA-seq | PDT | C1B | Batch2 | 7463 | 4117 | 29772 | 21995 | 791 | 493 | this manuscript |
| 1314 | SDHB | snRNA-seq | PT | C1A | Batch2 | 11666 | 9615 | 30800 | 23990 | 1896 | 1190 | this manuscript |
| 1312_L | NF1 | snRNA-seq | L | C2 | Batch2 | 2928 | 2095 | 28040 | 20021 | 192 | 179 | this manuscript |
| 1312_H | NF1 | scRNA-seq | H | C2 | Batch2 | 3633 | 1881 | 31969 | 24845 | 238 | 106 | this manuscript |
| 1313_H | Unknown | scRNA-seq | H | C1B | Batch2 | 9233 | 4792 | 31159 | 24139 | 1149 | 1142 | this manuscript |
| 1313_N | Unknown | snRNA-seq | N | C1B | Batch2 | 2090 | 1652 | 27107 | 18059 | 92 | 109 | this manuscript |
| 1313_L | Unknown | scRNA-seq | L | C1B | Batch3 | 43811 | 15443 | 31962 | 26236 | 8871 | 9636 | this manuscript |
| 1322_D | SDHB | scRNA-seq | PDT | C1A | Batch3 | 16467 | 11178 | 31816 | 25723 | 2013 | 936 | this manuscript |
| 1322_H | SDHB | scRNA-seq | H | C1A | Batch3 | 8153 | 5730 | 32080 | 25854 | 989 | 725 | this manuscript |
| 1322_N | SDHB | scRNA-seq | N | C1A | Batch3 | 7171 | 4458 | 32099 | 25960 | 723 | 631 | this manuscript |
| 1322_L | SDHB | scRNA-seq | L | C1A | Batch3 | 16602 | 1437 | 30744 | 24536 | 2326 | 3854 | this manuscript |
| 1322 | SDHB | snRNA-seq | PT | C1A | Batch3 | 17378 | 13485 | 30083 | 22745 | 2327 | 2141 | this manuscript |
| 1343_H | Unknown | scRNA-seq | H | C1B | Batch4 | 14357 | 12214 | 31490 | 24128 | 1040 | 1176 | this manuscript |
| 1343_N | Unknown | scRNA-seq | N | C1B | Batch4 | 27860 | 22283 | 32104 | 25570 | 4856 | 3870 | this manuscript |
| 1343_D | Unknown | scRNA-seq | PDT | C1B | Batch4 | 17065 | 10800 | 31626 | 25159 | 2550 | 1773 | this manuscript |
| 1415_N1 | Unknown | scRNA-seq | N(1) | C2 | Batch4 | 11908 | 7952 | 31397 | 24815 | 2046 | 541 | this manuscript |
| 1415_N2 | Unknown | scRNA-seq | N(2) | C2 | Batch4 | 7701 | 4050 | 32232 | 25864 | 795 | 485 | this manuscript |
| 1415_N3 | Unknown | scRNA-seq | N(3) | C2 | Batch4 | 7999 | 4173 | 31861 | 25499 | 796 | 581 | this manuscript |
| 1415_D1 | Unknown | scRNA-seq | PDT(1) | C2 | Batch4 | 7489 | 3790 | 31231 | 24514 | 838 | 576 | this manuscript |
| 1415_D2 | Unknown | scRNA-seq | PDT(2) | C2 | Batch4 | 10443 | 6278 | 31535 | 24959 | 1144 | 601 | this manuscript |
| 1415_D3 | Unknown | scRNA-seq | PDT(3) | C2 | Batch4 | 9326 | 5244 | 31734 | 25177 | 979 | 635 | this manuscript |
| **Total** | | | | | | **334737** | **211647** | | | | | |
| **Median** | | | | | | | | | **24133.5** | | | |

**Supplementary Table 9.** General metrics of the single cell/single nuclei RNAseq cohort. Abbreviations: PT - Primary Tumor, PDT - Primary Dissociated Tumor, N - Normoxia, H - Hypoxia1%, L - Long Term, N(1-3) and PDT(1-3) Ran three-times under different digestion conditions / C1A - Krebs Cycle Cluster, C1B - Pseudohypoxic Cluster, C2 - Kinase Signaling Cluster, U - Unknown

| Gene | p val | avg log2FC | pct.1 | pct.2 | p val adj | cluster | cluster annotation |
|---|---|---|---|---|---|---|---|
| MIAT | 0 | 3.911153639 | 0.889 | 0.191 | 0 | 0 | Tumor cells |
| DPP6 | 0 | 3.388400748 | 0.954 | 0.302 | 0 | 0 | Tumor cells |
| AGBL4 | 0 | 3.424296847 | 0.859 | 0.211 | 0 | 0 | Tumor cells |
| MYT1L | 0 | 3.50002487 | 0.839 | 0.191 | 0 | 0 | Tumor cells |
| HAND2-AS1 | 0 | 2.853068462 | 0.902 | 0.259 | 0 | 0 | Tumor cells |
| RALYL | 0 | 3.458142796 | 0.965 | 0.324 | 0 | 0 | Tumor cells |
| EML5 | 0 | 3.261360207 | 0.897 | 0.259 | 0 | 0 | Tumor cells |
| RIMS2 | 0 | 2.943156007 | 0.971 | 0.344 | 0 | 0 | Tumor cells |
| UNC79 | 0 | 3.253239116 | 0.856 | 0.231 | 0 | 0 | Tumor cells |
| HS6ST3 | 0 | 2.700433129 | 0.912 | 0.289 | 0 | 0 | Tumor cells |
| CTNNA2 | 0 | 2.808266213 | 0.876 | 0.255 | 0 | 0 | Tumor cells |
| KCNMA1 | 0 | 2.74337418 | 0.937 | 0.324 | 0 | 0 | Tumor cells |
| CACNA2D3 | 0 | 3.290482301 | 0.879 | 0.268 | 0 | 0 | Tumor cells |
| RIMBP2 | 0 | 2.700197936 | 0.836 | 0.229 | 0 | 0 | Tumor cells |
| DANT2 | 0 | 3.094549041 | 0.838 | 0.234 | 0 | 0 | Tumor cells |
| CADPS | 0 | 2.756608806 | 0.81 | 0.207 | 0 | 0 | Tumor cells |
| CACNA1D | 0 | 3.360284063 | 0.79 | 0.188 | 0 | 0 | Tumor cells |
| NEBL | 0 | 2.360406237 | 0.861 | 0.264 | 0 | 0 | Tumor cells |
| NLGN1 | 0 | 2.848324461 | 0.844 | 0.248 | 0 | 0 | Tumor cells |
| FMN2 | 0 | 3.697268418 | 0.76 | 0.166 | 0 | 0 | Tumor cells |
| COL1A2 | 0 | 2.433898085 | 0.881 | 0.194 | 0 | 1 | Fibroblasts |
| COL6A2 | 0 | 2.676140481 | 0.915 | 0.242 | 0 | 1 | Fibroblasts |
| COL3A1 | 0 | 2.65573856 | 0.856 | 0.197 | 0 | 1 | Fibroblasts |
| PRKG1 | 0 | 1.867556333 | 0.95 | 0.32 | 0 | 1 | Fibroblasts |
| TPM2 | 0 | 3.115575427 | 0.789 | 0.163 | 0 | 1 | Fibroblasts |
| BGN | 0 | 3.009089819 | 0.8 | 0.176 | 0 | 1 | Fibroblasts |
| PDGFRB | 0 | 2.648625489 | 0.787 | 0.164 | 0 | 1 | Fibroblasts |
| PLAC9 | 0 | 3.256475942 | 0.758 | 0.148 | 0 | 1 | Fibroblasts |
| COL6A1 | 0 | 2.617066528 | 0.859 | 0.249 | 0 | 1 | Fibroblasts |
| MT1E | 0 | 3.388047001 | 0.779 | 0.172 | 0 | 1 | Fibroblasts |
| FN1 | 0 | 2.693231078 | 0.95 | 0.352 | 0 | 1 | Fibroblasts |
| IGFBP7 | 0 | 2.372799691 | 0.979 | 0.384 | 0 | 1 | Fibroblasts |
| MGP | 0 | 2.698040124 | 0.804 | 0.215 | 0 | 1 | Fibroblasts |
| IGFBP4 | 0 | 2.279875282 | 0.847 | 0.263 | 0 | 1 | Fibroblasts |
| IFITM3 | 0 | 2.31903771 | 0.806 | 0.23 | 0 | 1 | Fibroblasts |
| MFGE8 | 0 | 2.786104435 | 0.743 | 0.176 | 0 | 1 | Fibroblasts |
| MT2A | 0 | 3.265554035 | 0.958 | 0.391 | 0 | 1 | Fibroblasts |
| LGALS1 | 0 | 2.184447056 | 0.928 | 0.362 | 0 | 1 | Fibroblasts |
| NR2F2 | 0 | 2.799973784 | 0.756 | 0.193 | 0 | 1 | Fibroblasts |
| SPARC | 0 | 1.637492039 | 0.872 | 0.314 | 0 | 1 | Fibroblasts |
| PLEKHG1 | 0 | 5.162807693 | 0.885 | 0.135 | 0 | 2 | Endothelial cells |
| FLT1 | 0 | 6.90800501 | 0.827 | 0.084 | 0 | 2 | Endothelial cells |
| LDB2 | 0 | 5.406904618 | 0.946 | 0.228 | 0 | 2 | Endothelial cells |
| ADGRL4 | 0 | 5.134498516 | 0.799 | 0.104 | 0 | 2 | Endothelial cells |
| PTPRB | 0 | 5.864196354 | 0.75 | 0.066 | 0 | 2 | Endothelial cells |
| ST6GALNAC3 | 0 | 5.040823843 | 0.835 | 0.153 | 0 | 2 | Endothelial cells |
| VWF | 0 | 6.861377097 | 0.751 | 0.069 | 0 | 2 | Endothelial cells |
| PLVAP | 0 | 6.573943747 | 0.741 | 0.061 | 0 | 2 | Endothelial cells |
| PRKCH | 0 | 3.797867609 | 0.826 | 0.15 | 0 | 2 | Endothelial cells |
| MECOM | 0 | 5.564797869 | 0.769 | 0.099 | 0 | 2 | Endothelial cells |
| EGFL7 | 0 | 4.743617161 | 0.789 | 0.121 | 0 | 2 | Endothelial cells |
| HSPG2 | 0 | 4.465367979 | 0.873 | 0.217 | 0 | 2 | Endothelial cells |
| SHANK3 | 0 | 4.897004595 | 0.74 | 0.101 | 0 | 2 | Endothelial cells |
| PECAM1 | 0 | 3.590532068 | 0.884 | 0.246 | 0 | 2 | Endothelial cells |
| NOTCH4 | 0 | 6.067559606 | 0.684 | 0.052 | 0 | 2 | Endothelial cells |
| KIAA1217 | 0 | 3.987135686 | 0.812 | 0.185 | 0 | 2 | Endothelial cells |
| CALCRL | 0 | 4.761666192 | 0.764 | 0.149 | 0 | 2 | Endothelial cells |
| ERG | 0 | 4.164671183 | 0.711 | 0.101 | 0 | 2 | Endothelial cells |
| ANO2 | 0 | 7.351358953 | 0.631 | 0.031 | 0 | 2 | Endothelial cells |
| WWTR1 | 0 | 3.84899204 | 0.882 | 0.292 | 0 | 2 | Endothelial cells |
| BEX1 | 0 | 5.163529806 | 0.898 | 0.112 | 0 | 3 | Tumor cells |
| PCSK1N | 0 | 3.863202444 | 0.915 | 0.181 | 0 | 3 | Tumor cells |
| SCG3 | 0 | 3.992472837 | 0.809 | 0.085 | 0 | 3 | Tumor cells |
| BEX2 | 0 | 4.485579172 | 0.792 | 0.081 | 0 | 3 | Tumor cells |
| L1CAM | 0 | 3.558368556 | 0.808 | 0.113 | 0 | 3 | Tumor cells |
| NSG1 | 0 | 4.326676537 | 0.773 | 0.082 | 0 | 3 | Tumor cells |
| SCG5 | 0 | 3.413976249 | 0.872 | 0.183 | 0 | 3 | Tumor cells |
| MLLT11 | 0 | 4.364240654 | 0.789 | 0.104 | 0 | 3 | Tumor cells |
| CLU | 0 | 3.669200608 | 0.865 | 0.182 | 0 | 3 | Tumor cells |
| CHRNA3 | 0 | 3.547835456 | 0.869 | 0.192 | 0 | 3 | Tumor cells |
| RGS4 | 0 | 3.645064487 | 0.848 | 0.172 | 0 | 3 | Tumor cells |
| STMN2 | 0 | 3.490229561 | 0.829 | 0.154 | 0 | 3 | Tumor cells |
| LINC00682 | 0 | 3.904783057 | 0.775 | 0.102 | 0 | 3 | Tumor cells |
| OCIAD2 | 0 | 4.099840899 | 0.77 | 0.098 | 0 | 3 | Tumor cells |
| APLP1 | 0 | 3.917008983 | 0.75 | 0.079 | 0 | 3 | Tumor cells |
| TAGLN3 | 0 | 4.457309835 | 0.727 | 0.056 | 0 | 3 | Tumor cells |
| SEC11C | 0 | 3.834897354 | 0.821 | 0.153 | 0 | 3 | Tumor cells |
| PRSS3 | 0 | 3.528181143 | 0.79 | 0.125 | 0 | 3 | Tumor cells |
| PTPRN | 0 | 2.948539672 | 0.852 | 0.194 | 0 | 3 | Tumor cells |
| QDPR | 0 | 3.335051353 | 0.869 | 0.213 | 0 | 3 | Tumor cells |
| PRKG1 | 0 | 2.669208344 | 0.922 | 0.441 | 0 | 4 | Fibroblasts |
| EPS8 | 0 | 2.263722866 | 0.8 | 0.432 | 0 | 4 | Fibroblasts |
| IGFBP7 | 0 | 0.7245265705 | 0.872 | 0.504 | 0 | 4 | Fibroblasts |
| NR2F2-AS1 | 0 | 2.472265226 | 0.65 | 0.289 | 0 | 4 | Fibroblasts |
| PDE1A | 0 | 2.173810135 | 0.591 | 0.273 | 0 | 4 | Fibroblasts |
| EBF2 | 0 | 2.521531793 | 0.554 | 0.237 | 0 | 4 | Fibroblasts |
| COL4A2 | 0 | 1.188491221 | 0.815 | 0.508 | 0 | 4 | Fibroblasts |
| COL4A1 | 0 | 1.210529612 | 0.8 | 0.502 | 0 | 4 | Fibroblasts |
| COL6A2 | 0 | 1.029628402 | 0.681 | 0.384 | 0 | 4 | Fibroblasts |
| ATP10A | 0 | 2.968926931 | 0.471 | 0.174 | 0 | 4 | Fibroblasts |
| CLMN | 0 | 2.84616679 | 0.578 | 0.292 | 0 | 4 | Fibroblasts |
| FN1 | 0 | 0.6542765674 | 0.76 | 0.478 | 0 | 4 | Fibroblasts |
| RBPMS | 0 | 1.953625325 | 0.707 | 0.435 | 0 | 4 | Fibroblasts |
| CALD1 | 0 | 1.528388921 | 0.895 | 0.628 | 0 | 4 | Fibroblasts |
| PTPRG | 0 | 1.419895063 | 0.858 | 0.595 | 0 | 4 | Fibroblasts |
| PDGFRB | 0 | 1.373479402 | 0.556 | 0.297 | 0 | 4 | Fibroblasts |
| COL1A2 | 0 | 0.9716586117 | 0.59 | 0.343 | 0 | 4 | Fibroblasts |
| MSC-AS1 | 0 | 1.913094304 | 0.488 | 0.249 | 0 | 4 | Fibroblasts |
| CRISPLD2 | 0 | 3.026182821 | 0.436 | 0.198 | 0 | 4 | Fibroblasts |
| CCDC102B | 0 | 2.230891239 | 0.528 | 0.3 | 0 | 4 | Fibroblasts |
| RBM47 | 0 | 4.938855411 | 0.857 | 0.042 | 0 | 5 | Monocytes/Macrophages |
| TBXAS1 | 0 | 6.040708377 | 0.807 | 0.062 | 0 | 5 | Monocytes/Macrophages |
| DOCK2 | 0 | 5.066445811 | 0.784 | 0.071 | 0 | 5 | Monocytes/Macrophages |
| PTPRC | 0 | 3.823850927 | 0.747 | 0.046 | 0 | 5 | Monocytes/Macrophages |
| AOAH | 0 | 4.579189712 | 0.75 | 0.05 | 0 | 5 | Monocytes/Macrophages |
| ARHGAP15 | 0 | 3.27628658 | 0.807 | 0.118 | 0 | 5 | Monocytes/Macrophages |
| CD74 | 0 | 7.170481125 | 0.764 | 0.075 | 0 | 5 | Monocytes/Macrophages |
| PLXDC2 | 0 | 4.173605002 | 0.94 | 0.256 | 0 | 5 | Monocytes/Macrophages |
| LAPTM5 | 0 | 5.611621494 | 0.737 | 0.057 | 0 | 5 | Monocytes/Macrophages |
| HLA-DRB1 | 0 | 7.508238159 | 0.72 | 0.053 | 0 | 5 | Monocytes/Macrophages |
| MS4A7 | 0 | 7.621758947 | 0.684 | 0.019 | 0 | 5 | Monocytes/Macrophages |
| PIK3R5 | 0 | 5.803352027 | 0.679 | 0.019 | 0 | 5 | Monocytes/Macrophages |
| HLA-DRA | 0 | 8.649218387 | 0.71 | 0.051 | 0 | 5 | Monocytes/Macrophages |
| CD83 | 0 | 6.285132985 | 0.72 | 0.062 | 0 | 5 | Monocytes/Macrophages |
| IQGAP2 | 0 | 3.689786954 | 0.733 | 0.083 | 0 | 5 | Monocytes/Macrophages |
| MCTP1 | 0 | 3.791935611 | 0.764 | 0.114 | 0 | 5 | Monocytes/Macrophages |
| RNF144B | 0 | 5.803344965 | 0.692 | 0.046 | 0 | 5 | Monocytes/Macrophages |
| FCGR2A | 0 | 8.251856351 | 0.641 | 0.011 | 0 | 5 | Monocytes/Macrophages |
| C5AR1 | 0 | 6.309823815 | 0.682 | 0.057 | 0 | 5 | Monocytes/Macrophages |
| TNFRSF1B | 0 | 4.393019054 | 0.697 | 0.076 | 0 | 5 | Monocytes/Macrophages |
| RIMS2 | 0 | 1.253757737 | 0.805 | 0.495 | 0 | 6 | Tumor cells |
| DPP6 | 0 | 1.160269868 | 0.76 | 0.46 | 0 | 6 | Tumor cells |
| KCNH8 | 0 | 2.974291795 | 0.469 | 0.187 | 0 | 6 | Tumor cells |
| PCLO | 0 | 2.059834917 | 0.742 | 0.465 | 0 | 6 | Tumor cells |
| MEG3 | 0 | 1.313425454 | 0.859 | 0.586 | 0 | 6 | Tumor cells |
| AC013652.1 | 0 | 3.311486452 | 0.415 | 0.142 | 0 | 6 | Tumor cells |
| VGF | 0 | 0.5018913236 | 0.464 | 0.192 | 0 | 6 | Tumor cells |
| CACNA2D3 | 0 | 1.500629789 | 0.684 | 0.417 | 0 | 6 | Tumor cells |
| BMPR1B | 0 | 1.411454323 | 0.603 | 0.337 | 0 | 6 | Tumor cells |
| FGF14 | 0 | 1.668981682 | 0.746 | 0.481 | 0 | 6 | Tumor cells |
| SLC35F3 | 0 | 1.809613876 | 0.505 | 0.243 | 0 | 6 | Tumor cells |
| HS6ST3 | 0 | 1.513644277 | 0.7 | 0.441 | 0 | 6 | Tumor cells |
| CCSER1 | 0 | 1.245771964 | 0.88 | 0.622 | 0 | 6 | Tumor cells |
| SYT1 | 0 | 1.406674534 | 0.804 | 0.546 | 0 | 6 | Tumor cells |
| RIMBP2 | 0 | 1.582814433 | 0.635 | 0.377 | 0 | 6 | Tumor cells |
| DLGAP1 | 0 | 2.535041083 | 0.598 | 0.34 | 0 | 6 | Tumor cells |
| RELN | 0 | 2.90024903 | 0.395 | 0.141 | 0 | 6 | Tumor cells |
| PRSS3 | 0 | 1.597397471 | 0.438 | 0.185 | 0 | 6 | Tumor cells |
| CTNNA2 | 0 | 1.713765177 | 0.658 | 0.406 | 0 | 6 | Tumor cells |
| PRSS12 | 0 | 3.678556385 | 0.314 | 0.062 | 0 | 6 | Tumor cells |
| THBS2 | 0 | 3.216878382 | 0.737 | 0.1 | 0 | 7 | Fibroblasts |
| PDGFRA | 0 | 2.96059831 | 0.755 | 0.12 | 0 | 7 | Fibroblasts |
| LTBP1 | 0 | 2.211807479 | 0.92 | 0.286 | 0 | 7 | Fibroblasts |
| MSC-AS1 | 0 | 1.743881136 | 0.869 | 0.242 | 0 | 7 | Fibroblasts |
| LTBP2 | 0 | 3.054804034 | 0.722 | 0.096 | 0 | 7 | Fibroblasts |
| COL1A2 | 0 | 2.541081166 | 0.95 | 0.337 | 0 | 7 | Fibroblasts |
| ITGA11 | 0 | 2.930933596 | 0.713 | 0.104 | 0 | 7 | Fibroblasts |
| COL1A1 | 0 | 2.945515385 | 0.853 | 0.244 | 0 | 7 | Fibroblasts |
| SRPX2 | 0 | 2.239977303 | 0.773 | 0.17 | 0 | 7 | Fibroblasts |
| COL3A1 | 0 | 2.174300829 | 0.93 | 0.335 | 0 | 7 | Fibroblasts |
| MRC2 | 0 | 2.125714838 | 0.737 | 0.142 | 0 | 7 | Fibroblasts |
| NNMT | 0 | 1.873392674 | 0.83 | 0.238 | 0 | 7 | Fibroblasts |
| COL6A3 | 0 | 2.330933619 | 0.849 | 0.258 | 0 | 7 | Fibroblasts |
| PLSCR4 | 0 | 2.470049109 | 0.793 | 0.203 | 0 | 7 | Fibroblasts |
| ANGPTL4 | 0 | 1.61085537 | 0.838 | 0.248 | 0 | 7 | Fibroblasts |
| YAP1 | 0 | 1.612561507 | 0.855 | 0.268 | 0 | 7 | Fibroblasts |
| PDGFRB | 0 | 1.612435593 | 0.878 | 0.293 | 0 | 7 | Fibroblasts |
| CDH11 | 0 | 1.786965643 | 0.806 | 0.222 | 0 | 7 | Fibroblasts |
| EDIL3 | 0 | 2.247672334 | 0.834 | 0.255 | 0 | 7 | Fibroblasts |
| DDR2 | 0 | 1.941303795 | 0.806 | 0.23 | 0 | 7 | Fibroblasts |
| PTPRC | 0 | 5.441675846 | 0.758 | 0.061 | 0 | 8 | Lymphocytes |
| SYTL3 | 0 | 6.445982508 | 0.724 | 0.075 | 0 | 8 | Lymphocytes |
| ARHGAP15 | 0 | 4.052127136 | 0.779 | 0.134 | 0 | 8 | Lymphocytes |
| SKAP1 | 0 | 5.85140839 | 0.667 | 0.049 | 0 | 8 | Lymphocytes |
| IKZF1 | 0 | 5.591978466 | 0.626 | 0.041 | 0 | 8 | Lymphocytes |
| CD247 | 0 | 9.263070387 | 0.592 | 0.013 | 0 | 8 | Lymphocytes |
| IQGAP2 | 0 | 4.081272206 | 0.671 | 0.099 | 0 | 8 | Lymphocytes |
| RHOH | 0 | 6.795340908 | 0.528 | 0.02 | 0 | 8 | Lymphocytes |
| CD69 | 0 | 8.172636488 | 0.516 | 0.015 | 0 | 8 | Lymphocytes |
| SAMSN1 | 0 | 3.34599723 | 0.566 | 0.07 | 0 | 8 | Lymphocytes |
| STK17B | 0 | 4.27997908 | 0.635 | 0.142 | 0 | 8 | Lymphocytes |
| STAT4 | 0 | 4.998415433 | 0.604 | 0.12 | 0 | 8 | Lymphocytes |
| CELF2 | 0 | 3.202462445 | 0.765 | 0.289 | 0 | 8 | Lymphocytes |
| CD53 | 0 | 4.801460159 | 0.513 | 0.04 | 0 | 8 | Lymphocytes |
| ITK | 0 | 8.840761085 | 0.463 | 0.011 | 0 | 8 | Lymphocytes |
| CYTIP | 0 | 5.232735499 | 0.477 | 0.03 | 0 | 8 | Lymphocytes |
| ADGRE5 | 0 | 4.294937462 | 0.604 | 0.162 | 0 | 8 | Lymphocytes |
| RNF125 | 0 | 5.961267588 | 0.482 | 0.055 | 0 | 8 | Lymphocytes |
| CD96 | 0 | 7.264802664 | 0.437 | 0.013 | 0 | 8 | Lymphocytes |
| PRKCB | 0 | 3.948446517 | 0.514 | 0.094 | 0 | 8 | Lymphocytes |
| CDH19 | 0 | 10.08042213 | 0.91 | 0.028 | 0 | 9 | Schwann cells |
| SLC35F1 | 0 | 7.045032834 | 0.897 | 0.071 | 0 | 9 | Schwann cells |
| PTPRZ1 | 0 | 9.561099286 | 0.735 | 0.011 | 0 | 9 | Schwann cells |
| NKAIN3 | 0 | 6.891383164 | 0.745 | 0.037 | 0 | 9 | Schwann cells |
| NRXN1 | 0 | 5.6712098 | 0.982 | 0.291 | 0 | 9 | Schwann cells |
| FRMD5 | 0 | 5.119430968 | 0.85 | 0.178 | 0 | 9 | Schwann cells |
| ADAM23 | 0 | 6.258410719 | 0.82 | 0.16 | 0 | 9 | Schwann cells |
| SOX6 | 0 | 5.004451325 | 0.773 | 0.12 | 0 | 9 | Schwann cells |
| IL1RAPL2 | 0 | 6.725410703 | 0.724 | 0.071 | 0 | 9 | Schwann cells |
| SEMA3C | 0 | 5.786582658 | 0.694 | 0.071 | 0 | 9 | Schwann cells |
| XKR4 | 0 | 5.142886048 | 0.709 | 0.098 | 0 | 9 | Schwann cells |
| CTNNA3 | 0 | 4.259082703 | 0.833 | 0.222 | 0 | 9 | Schwann cells |
| NEGR1 | 0 | 4.340410322 | 0.86 | 0.256 | 0 | 9 | Schwann cells |
| HMCN1 | 0 | 5.692534292 | 0.701 | 0.105 | 0 | 9 | Schwann cells |
| KIRREL3 | 0 | 5.502400014 | 0.702 | 0.107 | 0 | 9 | Schwann cells |
| PTPRJ | 0 | 4.13558299 | 0.797 | 0.24 | 0 | 9 | Schwann cells |
| ANK3 | 0 | 3.521737349 | 0.854 | 0.301 | 0 | 9 | Schwann cells |
| GINS3 | 0 | 6.831664673 | 0.604 | 0.058 | 0 | 9 | Schwann cells |
| CASC15 | 0 | 3.515573693 | 0.76 | 0.218 | 0 | 9 | Schwann cells |
| COL28A1 | 0 | 6.851968578 | 0.574 | 0.033 | 0 | 9 | Schwann cells |
| L1CAM | 0 | 2.329426934 | 0.864 | 0.171 | 0 | 10 | Tumor cells |
| APLP1 | 0 | 2.426417102 | 0.827 | 0.135 | 0 | 10 | Tumor cells |
| PCSK1N | 0 | 1.298704254 | 0.928 | 0.243 | 0 | 10 | Tumor cells |
| SEZ6L2 | 0 | 2.182546563 | 0.903 | 0.221 | 0 | 10 | Tumor cells |
| SCG3 | 0 | 2.232621474 | 0.829 | 0.147 | 0 | 10 | Tumor cells |
| BEX1 | 0 | 1.61400604 | 0.857 | 0.18 | 0 | 10 | Tumor cells |
| SYT4 | 0 | 2.186879867 | 0.883 | 0.207 | 0 | 10 | Tumor cells |
| PHOX2A | 0 | 2.231113155 | 0.852 | 0.189 | 0 | 10 | Tumor cells |
| SCN3B | 0 | 2.344550843 | 0.85 | 0.191 | 0 | 10 | Tumor cells |
| NSG1 | 0 | 2.202067063 | 0.797 | 0.14 | 0 | 10 | Tumor cells |
| SCG5 | 0 | 2.01394825 | 0.898 | 0.241 | 0 | 10 | Tumor cells |
| CHRNA3 | 0 | 2.022727659 | 0.906 | 0.249 | 0 | 10 | Tumor cells |
| FAIM2 | 0 | 1.920705816 | 0.824 | 0.168 | 0 | 10 | Tumor cells |
| CLU | 0 | 2.210743347 | 0.893 | 0.24 | 0 | 10 | Tumor cells |
| LINC00682 | 0 | 1.789133637 | 0.81 | 0.158 | 0 | 10 | Tumor cells |
| NEFL | 0 | 2.565270327 | 0.755 | 0.104 | 0 | 10 | Tumor cells |
| NSG2 | 0 | 2.72090418 | 0.856 | 0.207 | 0 | 10 | Tumor cells |
| BEX2 | 0 | 2.028484055 | 0.79 | 0.141 | 0 | 10 | Tumor cells |
| STMN2 | 0 | 1.991386599 | 0.857 | 0.211 | 0 | 10 | Tumor cells |
| ELAVL4 | 0 | 1.960332518 | 0.879 | 0.234 | 0 | 10 | Tumor cells |
| CYP11B1 | 0 | 9.549515168 | 0.933 | 0.009 | 0 | 11 | Adrenocortical cells |
| CYP11A1 | 0 | 10.78806316 | 0.904 | 0.011 | 0 | 11 | Adrenocortical cells |
| RBM47 | 0 | 5.890218285 | 0.939 | 0.074 | 0 | 11 | Adrenocortical cells |
| TNXB | 0 | 8.007418046 | 0.94 | 0.088 | 0 | 11 | Adrenocortical cells |
| SCARB1 | 0 | 7.188304057 | 0.973 | 0.259 | 0 | 11 | Adrenocortical cells |
| GRAMD1B | 0 | 6.612037777 | 0.964 | 0.267 | 0 | 11 | Adrenocortical cells |
| ABTB2 | 0 | 6.95126647 | 0.779 | 0.093 | 0 | 11 | Adrenocortical cells |
| KCNQ1 | 0 | 5.013304896 | 0.786 | 0.101 | 0 | 11 | Adrenocortical cells |
| STAR | 0 | 9.303080424 | 0.679 | 0.007 | 0 | 11 | Adrenocortical cells |
| NDC80 | 0 | 8.499816386 | 0.689 | 0.018 | 0 | 11 | Adrenocortical cells |
| SLC16A9 | 0 | 8.102017641 | 0.687 | 0.025 | 0 | 11 | Adrenocortical cells |
| MAPK4 | 0 | 7.942552942 | 0.682 | 0.023 | 0 | 11 | Adrenocortical cells |
| SH3BP5 | 0 | 5.401365174 | 0.892 | 0.24 | 0 | 11 | Adrenocortical cells |
| CCBE1 | 0 | 6.916717059 | 0.667 | 0.034 | 0 | 11 | Adrenocortical cells |
| ERN1 | 0 | 5.046229085 | 0.833 | 0.225 | 0 | 11 | Adrenocortical cells |
| CYP21A2 | 0 | 9.260117275 | 0.608 | 0.003 | 0 | 11 | Adrenocortical cells |
| ZNF331 | 0 | 4.33564642 | 0.904 | 0.324 | 0 | 11 | Adrenocortical cells |
| ALAS1 | 0 | 6.216845302 | 0.696 | 0.116 | 0 | 11 | Adrenocortical cells |
| CASP9 | 0 | 5.787160299 | 0.688 | 0.117 | 0 | 11 | Adrenocortical cells |
| SEMA6A | 0 | 4.713120127 | 0.771 | 0.212 | 0 | 11 | Adrenocortical cells |
| MKI67 | 0 | 7.842176922 | 0.697 | 0.005 | 0 | 12 | Endothelial cells |
| ASPM | 0 | 7.155651854 | 0.698 | 0.008 | 0 | 12 | Endothelial cells |
| DIAPH3 | 0 | 6.903475054 | 0.804 | 0.018 | 0 | 12 | Endothelial cells |
| TOP2A | 0 | 6.892537092 | 0.732 | 0.014 | 0 | 12 | Endothelial cells |
| ANLN | 0 | 6.841615013 | 0.717 | 0.009 | 0 | 12 | Endothelial cells |
| CENPF | 0 | 6.793178912 | 0.697 | 0.013 | 0 | 12 | Endothelial cells |
| RRM2 | 0 | 6.604640206 | 0.677 | 0.009 | 0 | 12 | Endothelial cells |
| NCAPG | 0 | 6.388699028 | 0.623 | 0.009 | 0 | 12 | Endothelial cells |
| BIRC5 | 0 | 6.376809848 | 0.603 | 0.007 | 0 | 12 | Endothelial cells |
| TPX2 | 0 | 6.35985028 | 0.697 | 0.018 | 0 | 12 | Endothelial cells |
| MELK | 0 | 6.241493265 | 0.654 | 0.012 | 0 | 12 | Endothelial cells |
| KNL1 | 0 | 6.075130242 | 0.64 | 0.014 | 0 | 12 | Endothelial cells |
| CENPE | 0 | 5.543557018 | 0.628 | 0.024 | 0 | 12 | Endothelial cells |
| PRC1 | 0 | 5.194407946 | 0.669 | 0.038 | 0 | 12 | Endothelial cells |
| NUSAP1 | 0 | 5.145394263 | 0.724 | 0.03 | 0 | 12 | Endothelial cells |
| TYMS | 0 | 5.132780773 | 0.63 | 0.028 | 0 | 12 | Endothelial cells |
| CENPK | 0 | 4.697694324 | 0.658 | 0.035 | 0 | 12 | Endothelial cells |
| BRCA2 | 0 | 4.051254943 | 0.652 | 0.047 | 0 | 12 | Endothelial cells |
| KIF20B | 0 | 3.866954059 | 0.663 | 0.064 | 0 | 12 | Endothelial cells |
| SMC4 | 0 | 2.843579594 | 0.804 | 0.221 | 0 | 12 | Endothelial cells |
| VWF | 0 | 2.38758214 | 0.841 | 0.164 | 0 | 13 | Endothelial cells |
| ANO2 | 0 | 2.670164965 | 0.782 | 0.115 | 0 | 13 | Endothelial cells |
| PLEKHG1 | 0 | 2.262997045 | 0.902 | 0.239 | 0 | 13 | Endothelial cells |
| ST6GALNAC3 | 0 | 2.509464895 | 0.904 | 0.248 | 0 | 13 | Endothelial cells |
| MECOM | 0 | 2.748377165 | 0.836 | 0.192 | 0 | 13 | Endothelial cells |
| LDB2 | 0 | 2.491369482 | 0.966 | 0.328 | 0 | 13 | Endothelial cells |
| PLVAP | 1.31E-307 | 1.924464912 | 0.748 | 0.156 | 4.00E-303 | 13 | Endothelial cells |
| PTPRB | 2.82E-305 | 2.112438975 | 0.755 | 0.161 | 8.56E-301 | 13 | Endothelial cells |
| NOTCH4 | 6.79E-300 | 2.543931652 | 0.685 | 0.14 | 2.06E-295 | 13 | Endothelial cells |
| PRKCH | 1.64E-292 | 2.189322042 | 0.88 | 0.245 | 4.99E-288 | 13 | Endothelial cells |
| NOSTRIN | 1.60E-290 | 2.584472267 | 0.509 | 0.079 | 4.86E-286 | 13 | Endothelial cells |
| DIPK2B | 9.98E-284 | 2.666205079 | 0.583 | 0.106 | 3.03E-279 | 13 | Endothelial cells |
| FLT1 | 1.61E-277 | 1.88681233 | 0.776 | 0.188 | 4.91E-273 | 13 | Endothelial cells |
| KIAA1217 | 4.56E-276 | 2.339577144 | 0.88 | 0.272 | 1.38E-271 | 13 | Endothelial cells |
| GRB10 | 2.46E-272 | 2.412596762 | 0.966 | 0.436 | 7.49E-268 | 13 | Endothelial cells |
| PTPRD | 8.86E-269 | 1.549780536 | 0.834 | 0.206 | 2.69E-264 | 13 | Endothelial cells |
| MAGI1 | 6.36E-266 | 2.382408075 | 0.983 | 0.463 | 1.93E-261 | 13 | Endothelial cells |
| EGFL7 | 2.77E-264 | 2.171564074 | 0.797 | 0.215 | 8.41E-260 | 13 | Endothelial cells |
| PREX2 | 1.13E-262 | 2.620334831 | 0.646 | 0.141 | 3.43E-258 | 13 | Endothelial cells |
| DOCK9 | 1.34E-260 | 2.314470383 | 0.945 | 0.401 | 4.09E-256 | 13 | Endothelial cells |
| ASIC2 | 0 | 4.848322683 | 0.868 | 0.092 | 0 | 14 | Tumor cells |
| DPP10 | 0 | 3.437557121 | 0.812 | 0.043 | 0 | 14 | Tumor cells |
| FAM153CP | 0 | 3.635119929 | 0.862 | 0.113 | 0 | 14 | Tumor cells |
| CNTNAP4 | 0 | 3.566368182 | 0.824 | 0.081 | 0 | 14 | Tumor cells |
| LINC01266 | 0 | 3.799641276 | 0.793 | 0.051 | 0 | 14 | Tumor cells |
| CDH4 | 0 | 4.194971254 | 0.834 | 0.092 | 0 | 14 | Tumor cells |
| GPR139 | 0 | 3.752134438 | 0.777 | 0.046 | 0 | 14 | Tumor cells |
| ADGRL3 | 0 | 2.511643912 | 0.831 | 0.102 | 0 | 14 | Tumor cells |
| XKR4 | 0 | 1.557528664 | 0.834 | 0.112 | 0 | 14 | Tumor cells |
| SNTG2 | 0 | 3.980207139 | 0.765 | 0.047 | 0 | 14 | Tumor cells |
| COL27A1 | 0 | 3.355542015 | 0.818 | 0.1 | 0 | 14 | Tumor cells |
| EYS | 0 | 3.310489464 | 0.837 | 0.125 | 0 | 14 | Tumor cells |
| TAFA5 | 0 | 4.122808688 | 0.759 | 0.053 | 0 | 14 | Tumor cells |
| LINC02641 | 0 | 3.102090645 | 0.824 | 0.119 | 0 | 14 | Tumor cells |
| TENM2 | 0 | 3.268176975 | 0.774 | 0.07 | 0 | 14 | Tumor cells |
| DIO3OS | 0 | 4.832641897 | 0.712 | 0.014 | 0 | 14 | Tumor cells |
| STK32B | 0 | 3.001181461 | 0.787 | 0.098 | 0 | 14 | Tumor cells |
| AF279873.3 | 0 | 2.98882409 | 0.781 | 0.097 | 0 | 14 | Tumor cells |
| BAIAP3 | 0 | 2.877512489 | 0.777 | 0.094 | 0 | 14 | Tumor cells |
| CACNG4 | 0 | 3.721796144 | 0.762 | 0.08 | 0 | 14 | Tumor cells |
| C5orf17 | 4.96E-80 | 3.859803903 | 0.769 | 0.04 | 1.51E-75 | 15 | Tumor cells |
| MMP1 | 8.29E-57 | 4.198691268 | 0.962 | 0.094 | 2.52E-52 | 15 | Tumor cells |
| CDH12 | 1.72E-52 | 5.296945363 | 0.692 | 0.053 | 5.22E-48 | 15 | Tumor cells |
| AC009315.1 | 3.35E-47 | 3.540831841 | 0.692 | 0.054 | 1.02E-42 | 15 | Tumor cells |
| DISP3 | 2.17E-43 | 4.918681552 | 0.423 | 0.023 | 6.60E-39 | 15 | Tumor cells |
| CLDN5 | 6.42E-34 | 2.945265627 | 0.808 | 0.102 | 1.95E-29 | 15 | Tumor cells |
| AC134312.4 | 1.83E-33 | 3.405368881 | 0.538 | 0.046 | 5.56E-29 | 15 | Tumor cells |
| AL121917.1 | 3.72E-32 | 4.378392646 | 0.5 | 0.043 | 1.13E-27 | 15 | Tumor cells |
| FABP5 | 7.26E-32 | 3.961327699 | 1 | 0.206 | 2.21E-27 | 15 | Tumor cells |
| GXYLT2 | 1.45E-31 | 4.572074989 | 0.615 | 0.068 | 4.39E-27 | 15 | Tumor cells |
| BMP2 | 1.76E-31 | 3.459120988 | 0.615 | 0.065 | 5.36E-27 | 15 | Tumor cells |
| S100A16 | 3.10E-31 | 3.621627615 | 0.731 | 0.092 | 9.42E-27 | 15 | Tumor cells |
| TREM1 | 1.40E-30 | 3.564907364 | 0.577 | 0.059 | 4.25E-26 | 15 | Tumor cells |
| PTHLH | 1.83E-30 | 4.300778163 | 0.462 | 0.039 | 5.56E-26 | 15 | Tumor cells |
| CARTPT | 4.57E-28 | 2.38110253 | 0.962 | 0.186 | 1.39E-23 | 15 | Tumor cells |
| SORCS3 | 7.08E-27 | 3.281329873 | 0.962 | 0.22 | 2.15E-22 | 15 | Tumor cells |
| FAM153CP | 1.14E-26 | 3.773429653 | 0.731 | 0.114 | 3.45E-22 | 15 | Tumor cells |
| LINC02810 | 1.19E-26 | 3.66191592 | 0.346 | 0.024 | 3.62E-22 | 15 | Tumor cells |
| TMEM45A | 2.77E-26 | 3.557023077 | 0.885 | 0.187 | 8.42E-22 | 15 | Tumor cells |
| FSIP2 | 3.09E-25 | 3.298894374 | 0.538 | 0.06 | 9.38E-21 | 15 | Tumor cells |

**Supplementary Table 10.** Top markers of the 16 cell clusters annotated in sc/snRNAseq cohort.

| Gene | p val | avg log2FC | pct.1 | pct.2 | p val adj | cluster | cell type reclustered |
|---|---|---|---|---|---|---|---|
| PWRN1 | 0 | 2.485974989 | 0.621 | 0.255 | 0 | 0 | Tumor cells |
| MT-ND4L | 0 | 0.2780189501 | 0.429 | 0.789 | 0 | 0 | Tumor cells |
| PCDH15 | 0 | 2.213799623 | 0.528 | 0.193 | 0 | 0 | Tumor cells |
| MIR646HG | 0 | 2.949634849 | 0.418 | 0.103 | 0 | 0 | Tumor cells |
| ADAMTS9 | 0 | 0.2835121038 | 0.331 | 0.633 | 0 | 0 | Tumor cells |
| SNTG1 | 0 | 2.32800584 | 0.604 | 0.304 | 0 | 0 | Tumor cells |
| ZBTB7A | 0 | 0.2578341671 | 0.284 | 0.574 | 0 | 0 | Tumor cells |
| UGCG | 0 | 0.2658170526 | 0.187 | 0.459 | 0 | 0 | Tumor cells |
| PTOV1 | 0 | 0.3256897212 | 0.273 | 0.544 | 0 | 0 | Tumor cells |
| SORCS3 | 0 | 1.925807148 | 0.541 | 0.273 | 0 | 0 | Tumor cells |
| SP3 | 0 | 0.2995791027 | 0.32 | 0.588 | 0 | 0 | Tumor cells |
| TANK | 0 | 0.2637986931 | 0.282 | 0.547 | 0 | 0 | Tumor cells |
| RIF1 | 0 | 0.2502611691 | 0.331 | 0.596 | 0 | 0 | Tumor cells |
| ZNF580 | 0 | 0.2603298533 | 0.149 | 0.413 | 0 | 0 | Tumor cells |
| SH3KBP1 | 0 | 0.2875417741 | 0.164 | 0.427 | 0 | 0 | Tumor cells |
| TPD52L2 | 0 | 0.2872193391 | 0.234 | 0.496 | 0 | 0 | Tumor cells |
| SGCD | 0 | 2.646497482 | 0.435 | 0.174 | 0 | 0 | Tumor cells |
| AK3 | 0 | 0.3462150941 | 0.308 | 0.568 | 0 | 0 | Tumor cells |
| EIF2AK2 | 0 | 0.3031902474 | 0.353 | 0.612 | 0 | 0 | Tumor cells |
| AP3D1 | 0 | 0.3070200476 | 0.332 | 0.59 | 0 | 0 | Tumor cells |
| OST4 | 0 | 5.08639482 | 0.91 | 0.077 | 0 | 1 | Tumor cells |
| RPS10 | 0 | 4.745578511 | 0.91 | 0.088 | 0 | 1 | Tumor cells |
| RPL27 | 0 | 4.133516672 | 0.93 | 0.109 | 0 | 1 | Tumor cells |
| UBL5 | 0 | 4.965324691 | 0.932 | 0.111 | 0 | 1 | Tumor cells |
| RPL35A | 0 | 4.360713691 | 0.963 | 0.144 | 0 | 1 | Tumor cells |
| RPS13 | 0 | 4.454446425 | 0.95 | 0.135 | 0 | 1 | Tumor cells |
| FAU | 0 | 4.597955023 | 0.972 | 0.158 | 0 | 1 | Tumor cells |
| RPL36AL | 0 | 4.346815941 | 0.914 | 0.103 | 0 | 1 | Tumor cells |
| RPL24 | 0 | 4.344297435 | 0.966 | 0.155 | 0 | 1 | Tumor cells |
| S100A11 | 0 | 5.47981983 | 0.906 | 0.096 | 0 | 1 | Tumor cells |
| COX7C | 0 | 4.461779665 | 0.94 | 0.131 | 0 | 1 | Tumor cells |
| TIMP1 | 0 | 3.063853402 | 0.941 | 0.132 | 0 | 1 | Tumor cells |
| SRP14 | 0 | 4.362159814 | 0.949 | 0.141 | 0 | 1 | Tumor cells |
| RPS8 | 0 | 4.165711549 | 0.976 | 0.169 | 0 | 1 | Tumor cells |
| RPS7 | 0 | 4.21043851 | 0.965 | 0.158 | 0 | 1 | Tumor cells |
| COX6A1 | 0 | 4.784560664 | 0.909 | 0.105 | 0 | 1 | Tumor cells |
| RPS25 | 0 | 4.345351979 | 0.967 | 0.163 | 0 | 1 | Tumor cells |
| RPS15A | 0 | 3.91493179 | 0.95 | 0.147 | 0 | 1 | Tumor cells |
| RPL26 | 0 | 4.453042995 | 0.975 | 0.173 | 0 | 1 | Tumor cells |
| ATP5MC3 | 0 | 4.699714055 | 0.874 | 0.076 | 0 | 1 | Tumor cells |
| FOSB | 0 | 2.583373132 | 0.478 | 0.145 | 0 | 2 | Tumor cells |
| NR4A3 | 0 | 2.504976459 | 0.415 | 0.134 | 0 | 2 | Tumor cells |
| NR4A2 | 0 | 2.01974495 | 0.475 | 0.224 | 0 | 2 | Tumor cells |
| NR4A1 | 0 | 1.463605895 | 0.592 | 0.344 | 0 | 2 | Tumor cells |
| BTG2 | 0 | 1.751468928 | 0.435 | 0.193 | 0 | 2 | Tumor cells |
| HOMER1 | 0 | 2.271702262 | 0.49 | 0.249 | 0 | 2 | Tumor cells |
| FOS | 0 | 1.23135561 | 0.561 | 0.324 | 0 | 2 | Tumor cells |
| EGR1 | 0 | 1.465416891 | 0.533 | 0.299 | 0 | 2 | Tumor cells |
| ARC | 0 | 1.558719145 | 0.461 | 0.238 | 0 | 2 | Tumor cells |
| TIPARP | 0 | 2.153756791 | 0.388 | 0.178 | 0 | 2 | Tumor cells |
| IFRD1 | 0 | 1.946253971 | 0.596 | 0.39 | 0 | 2 | Tumor cells |
| ZNF10 | 0 | 1.570952823 | 0.662 | 0.512 | 0 | 2 | Tumor cells |
| ARIH1 | 0 | 1.722375863 | 0.797 | 0.708 | 0 | 2 | Tumor cells |
| EIF4A3 | 7.55E-302 | 2.076773998 | 0.439 | 0.237 | 2.29E-297 | 2 | Tumor cells |
| USP36 | 1.07E-301 | 1.654557548 | 0.598 | 0.409 | 3.25E-297 | 2 | Tumor cells |
| RABGEF1 | 5.10E-299 | 1.477332885 | 0.699 | 0.575 | 1.55E-294 | 2 | Tumor cells |
| ZNF331 | 4.45E-297 | 1.824768046 | 0.602 | 0.437 | 1.35E-292 | 2 | Tumor cells |
| VPS37B | 1.96E-289 | 1.963897621 | 0.456 | 0.255 | 5.95E-285 | 2 | Tumor cells |
| ZNF571 | 2.79E-281 | 2.247099587 | 0.383 | 0.189 | 8.49E-277 | 2 | Tumor cells |
| BTAF1 | 3.73E-275 | 1.676491678 | 0.624 | 0.487 | 1.13E-270 | 2 | Tumor cells |
| FN1 | 0 | 6.400830523 | 0.83 | 0.2 | 0 | 3 | Tumor cells |
| IGFBP7 | 0 | 6.754631428 | 0.727 | 0.126 | 0 | 3 | Tumor cells |
| PRKG1 | 0 | 5.36697075 | 0.748 | 0.155 | 0 | 3 | Tumor cells |
| MT2A | 0 | 6.190909036 | 0.855 | 0.263 | 0 | 3 | Tumor cells |
| COL4A2 | 0 | 5.993230825 | 0.696 | 0.142 | 0 | 3 | Tumor cells |
| COL4A1 | 0 | 6.254736876 | 0.701 | 0.148 | 0 | 3 | Tumor cells |
| COL6A2 | 0 | 6.584350834 | 0.634 | 0.089 | 0 | 3 | Tumor cells |
| LGALS1 | 0 | 2.210865529 | 0.767 | 0.259 | 0 | 3 | Tumor cells |
| COL3A1 | 0 | 6.795141288 | 0.585 | 0.088 | 0 | 3 | Tumor cells |
| TMSB4X | 0 | 2.415770346 | 0.87 | 0.383 | 0 | 3 | Tumor cells |
| TIMP1 | 0 | 3.296498035 | 0.838 | 0.352 | 0 | 3 | Tumor cells |
| S100A6 | 0 | 0.9881840646 | 0.805 | 0.326 | 0 | 3 | Tumor cells |
| EPS8 | 0 | 3.712448337 | 0.625 | 0.164 | 0 | 3 | Tumor cells |
| FTL | 0 | 1.930539687 | 0.912 | 0.455 | 0 | 3 | Tumor cells |
| COL1A2 | 0 | 6.867426431 | 0.527 | 0.075 | 0 | 3 | Tumor cells |
| VIM | 0 | 2.05794913 | 0.793 | 0.342 | 0 | 3 | Tumor cells |
| TMSB10 | 0 | 1.602399075 | 0.844 | 0.393 | 0 | 3 | Tumor cells |
| IGFBP4 | 0 | 5.690224081 | 0.532 | 0.09 | 0 | 3 | Tumor cells |
| MT1E | 0 | 5.403383404 | 0.54 | 0.103 | 0 | 3 | Tumor cells |
| RPLP1 | 0 | 1.570724319 | 0.904 | 0.467 | 0 | 3 | Tumor cells |
| AKR1C1 | 0 | 1.690064625 | 0.901 | 0.369 | 0 | 0 | Fibroblast |
| PLIN2 | 0 | 1.707247601 | 0.895 | 0.367 | 0 | 0 | Fibroblast |
| NQO1 | 0 | 1.407410561 | 0.711 | 0.232 | 0 | 0 | Fibroblast |
| C1R | 0 | 1.42854038 | 0.797 | 0.319 | 0 | 0 | Fibroblast |
| AKR1C3 | 0 | 1.756624357 | 0.653 | 0.188 | 0 | 0 | Fibroblast |
| GLRX | 0 | 1.115388422 | 0.826 | 0.364 | 0 | 0 | Fibroblast |
| PECAM1 | 0 | 1.074984811 | 0.73 | 0.27 | 0 | 0 | Fibroblast |
| AKR1B1 | 0 | 1.798477616 | 0.644 | 0.197 | 0 | 0 | Fibroblast |
| RAB13 | 0 | 1.165930548 | 0.829 | 0.385 | 0 | 0 | Fibroblast |
| C1S | 0 | 1.448860305 | 0.701 | 0.26 | 0 | 0 | Fibroblast |
| MTSS1 | 0 | 1.411635516 | 0.692 | 0.252 | 0 | 0 | Fibroblast |
| RGS6 | 0 | 1.499024189 | 0.675 | 0.236 | 0 | 0 | Fibroblast |
| TWIST1 | 0 | 1.471178279 | 0.649 | 0.215 | 0 | 0 | Fibroblast |
| GYPC | 0 | 1.218922476 | 0.743 | 0.319 | 0 | 0 | Fibroblast |
| CTSD | 0 | 1.177509218 | 0.906 | 0.492 | 0 | 0 | Fibroblast |
| SOD2 | 0 | 1.822854081 | 0.933 | 0.529 | 0 | 0 | Fibroblast |
| SRPX2 | 0 | 0.9901249856 | 0.693 | 0.286 | 0 | 0 | Fibroblast |
| MT1X | 0 | 1.016039352 | 0.802 | 0.403 | 0 | 0 | Fibroblast |
| AKR1C2 | 0 | 2.005996974 | 0.519 | 0.125 | 0 | 0 | Fibroblast |
| DBI | 0 | 0.8652705699 | 0.77 | 0.382 | 0 | 0 | Fibroblast |
| FOSB | 0 | 4.759786428 | 0.618 | 0.117 | 0 | 1 | Fibroblast |
| SLC38A11 | 0 | 4.224259306 | 0.556 | 0.087 | 0 | 1 | Fibroblast |
| NR4A1 | 0 | 3.773072103 | 0.621 | 0.173 | 0 | 1 | Fibroblast |
| MYH11 | 0 | 3.123418944 | 0.619 | 0.171 | 0 | 1 | Fibroblast |
| FOS | 0 | 4.451525065 | 0.722 | 0.304 | 0 | 1 | Fibroblast |
| LAMA3 | 0 | 3.935687843 | 0.496 | 0.084 | 0 | 1 | Fibroblast |
| ATF3 | 0 | 4.053376183 | 0.524 | 0.122 | 0 | 1 | Fibroblast |
| ANO1 | 0 | 2.519016613 | 0.608 | 0.22 | 0 | 1 | Fibroblast |
| KLF2 | 0 | 4.529775045 | 0.453 | 0.07 | 0 | 1 | Fibroblast |
| RCAN2 | 0 | 2.788311444 | 0.762 | 0.385 | 0 | 1 | Fibroblast |
| JAG1 | 0 | 2.488989066 | 0.607 | 0.234 | 0 | 1 | Fibroblast |
| BTG2 | 0 | 4.176932821 | 0.457 | 0.104 | 0 | 1 | Fibroblast |
| HES4 | 0 | 2.94192984 | 0.465 | 0.128 | 0 | 1 | Fibroblast |
| CCDC3 | 0 | 3.006405956 | 0.553 | 0.224 | 0 | 1 | Fibroblast |
| PLCL1 | 0 | 2.882110305 | 0.769 | 0.443 | 0 | 1 | Fibroblast |
| CPM | 0 | 2.817848756 | 0.598 | 0.272 | 0 | 1 | Fibroblast |
| ATP10A | 0 | 2.205117337 | 0.677 | 0.354 | 0 | 1 | Fibroblast |
| CTNNA3 | 0 | 2.563344753 | 0.611 | 0.288 | 0 | 1 | Fibroblast |
| PMEPA1 | 0 | 2.340095426 | 0.6 | 0.286 | 0 | 1 | Fibroblast |
| HSPA1B | 0 | 4.018014464 | 0.417 | 0.106 | 0 | 1 | Fibroblast |
| TFPI2 | 0 | 4.016292903 | 0.875 | 0.192 | 0 | 2 | Fibroblast |
| NPY | 0 | 6.372461054 | 0.88 | 0.211 | 0 | 2 | Fibroblast |
| POSTN | 0 | 2.828725111 | 0.678 | 0.121 | 0 | 2 | Fibroblast |
| PENK | 0 | 5.056034304 | 0.897 | 0.356 | 0 | 2 | Fibroblast |
| SERPINE2 | 0 | 2.902085392 | 0.83 | 0.297 | 0 | 2 | Fibroblast |
| PTX3 | 0 | 3.151875408 | 0.564 | 0.09 | 0 | 2 | Fibroblast |
| AC105402.3 | 0 | 2.592050683 | 0.603 | 0.178 | 0 | 2 | Fibroblast |
| ODAM | 0 | 6.285269724 | 0.331 | 0.014 | 0 | 2 | Fibroblast |
| SORCS1 | 0 | 4.743763959 | 0.327 | 0.062 | 0 | 2 | Fibroblast |
| MGST1 | 0 | 2.113408229 | 0.553 | 0.291 | 0 | 2 | Fibroblast |
| SCG2 | 0 | 2.994163928 | 0.567 | 0.31 | 0 | 2 | Fibroblast |
| CCSER1 | 0 | 2.061919214 | 0.476 | 0.237 | 0 | 2 | Fibroblast |
| NDUFA4 | 0 | 2.581499203 | 0.6 | 0.363 | 0 | 2 | Fibroblast |
| COL1A1 | 0 | 1.733499896 | 0.838 | 0.606 | 0 | 2 | Fibroblast |
| GNG11 | 0 | 2.336358076 | 0.841 | 0.619 | 0 | 2 | Fibroblast |
| FGF14 | 0 | 2.469590017 | 0.401 | 0.187 | 0 | 2 | Fibroblast |
| PCLO | 0 | 2.708188335 | 0.353 | 0.14 | 0 | 2 | Fibroblast |
| CHGB | 0 | 3.176829098 | 0.564 | 0.352 | 0 | 2 | Fibroblast |
| SRSF6 | 0 | 0.2961482626 | 0.118 | 0.328 | 0 | 2 | Fibroblast |
| VCAN | 0 | 1.131918295 | 0.369 | 0.165 | 0 | 2 | Fibroblast |
| MIR100HG | 0 | 2.795639048 | 0.857 | 0.323 | 0 | 3 | Fibroblast |
| BICC1 | 0 | 2.721430402 | 0.745 | 0.225 | 0 | 3 | Fibroblast |
| FAP | 0 | 2.649913372 | 0.643 | 0.125 | 0 | 3 | Fibroblast |
| VCAN | 0 | 3.099410824 | 0.659 | 0.151 | 0 | 3 | Fibroblast |
| BNC2 | 0 | 4.077860893 | 0.571 | 0.073 | 0 | 3 | Fibroblast |
| MTHFD1L | 0 | 2.193603639 | 0.759 | 0.271 | 0 | 3 | Fibroblast |
| PVT1 | 0 | 2.019542944 | 0.744 | 0.257 | 0 | 3 | Fibroblast |
| COL8A1 | 0 | 3.550338598 | 0.601 | 0.117 | 0 | 3 | Fibroblast |
| DDAH1 | 0 | 2.794426647 | 0.674 | 0.198 | 0 | 3 | Fibroblast |
| ITGA11 | 0 | 1.798514721 | 0.715 | 0.242 | 0 | 3 | Fibroblast |
| ADAMTS2 | 0 | 1.728846457 | 0.761 | 0.29 | 0 | 3 | Fibroblast |
| THBS2 | 0 | 1.834691215 | 0.738 | 0.267 | 0 | 3 | Fibroblast |
| ABCA9 | 0 | 3.03345179 | 0.562 | 0.098 | 0 | 3 | Fibroblast |
| NAALADL2 | 0 | 1.662699705 | 0.729 | 0.284 | 0 | 3 | Fibroblast |
| LAMA2 | 0 | 2.47503929 | 0.605 | 0.16 | 0 | 3 | Fibroblast |
| PAPPA | 0 | 3.349580504 | 0.669 | 0.225 | 0 | 3 | Fibroblast |
| ADAMTS12 | 0 | 1.927957301 | 0.741 | 0.297 | 0 | 3 | Fibroblast |
| SUGCT | 0 | 2.206495498 | 0.594 | 0.152 | 0 | 3 | Fibroblast |
| LMO7 | 0 | 2.526417764 | 0.575 | 0.134 | 0 | 3 | Fibroblast |
| AFAP1 | 0 | 1.924979495 | 0.649 | 0.212 | 0 | 3 | Fibroblast |
| MEG3 | 0 | 4.047182706 | 0.873 | 0.243 | 0 | 4 | Fibroblast |
| MEG8 | 0 | 3.767672282 | 0.783 | 0.166 | 0 | 4 | Fibroblast |
| RALYL | 0 | 4.667244048 | 0.683 | 0.082 | 0 | 4 | Fibroblast |
| DLK1 | 0 | 3.261316501 | 0.711 | 0.143 | 0 | 4 | Fibroblast |
| FAM155A | 0 | 2.983993169 | 0.822 | 0.265 | 0 | 4 | Fibroblast |
| RIMS2 | 0 | 3.906710545 | 0.655 | 0.105 | 0 | 4 | Fibroblast |
| STC1 | 0 | 1.979057952 | 0.817 | 0.296 | 0 | 4 | Fibroblast |
| CCSER1 | 0 | 2.879400005 | 0.756 | 0.248 | 0 | 4 | Fibroblast |
| EPB41L3 | 0 | 3.997912178 | 0.59 | 0.091 | 0 | 4 | Fibroblast |
| CXCL1 | 0 | 1.030276267 | 0.746 | 0.274 | 0 | 4 | Fibroblast |
| RYR2 | 0 | 3.117044105 | 0.585 | 0.115 | 0 | 4 | Fibroblast |
| LSAMP | 0 | 3.238759047 | 0.592 | 0.122 | 0 | 4 | Fibroblast |
| SYT1 | 0 | 2.86869225 | 0.671 | 0.206 | 0 | 4 | Fibroblast |
| NRG1 | 0 | 3.393219432 | 0.564 | 0.102 | 0 | 4 | Fibroblast |
| CHGA | 0 | 2.64681786 | 0.672 | 0.213 | 0 | 4 | Fibroblast |
| LRRTM4 | 0 | 4.533787449 | 0.517 | 0.061 | 0 | 4 | Fibroblast |
| TENM3 | 0 | 4.343503851 | 0.503 | 0.054 | 0 | 4 | Fibroblast |
| KCNMB2 | 0 | 4.346214664 | 0.499 | 0.054 | 0 | 4 | Fibroblast |
| DPP6 | 0 | 3.731366315 | 0.515 | 0.088 | 0 | 4 | Fibroblast |
| CTNND2 | 0 | 3.853069277 | 0.466 | 0.058 | 0 | 4 | Fibroblast |
| NCAM1 | 0 | 5.647982669 | 0.502 | 0.111 | 0 | 4 | Endothelial cells |
| NOTCH3 | 0 | 5.608079508 | 0.683 | 0.06 | 0 | 2 | Endothelial cells |
| TMEM178B | 0 | 5.563676729 | 0.489 | 0.091 | 0 | 4 | Endothelial cells |
| ADAMTS18 | 0 | 5.510217367 | 0.432 | 0.021 | 0 | 3 | Endothelial cells |
| S1PR3 | 0 | 5.496691376 | 0.691 | 0.036 | 0 | 2 | Endothelial cells |
| EDNRA | 0 | 5.464592379 | 0.704 | 0.043 | 0 | 2 | Endothelial cells |
| PDGFRB | 0 | 5.461844517 | 0.814 | 0.057 | 0 | 2 | Endothelial cells |
| NDUFA4L2 | 0 | 5.435685077 | 0.824 | 0.067 | 0 | 2 | Endothelial cells |
| ADGRG6 | 0 | 5.422674105 | 0.345 | 0.015 | 0 | 3 | Endothelial cells |
| LINC02147 | 0 | 5.263367262 | 0.544 | 0.037 | 0 | 3 | Endothelial cells |
| KCNQ5 | 0 | 5.048343792 | 0.493 | 0.098 | 0 | 4 | Endothelial cells |
| PLAC9 | 0 | 4.990207512 | 0.838 | 0.085 | 0 | 2 | Endothelial cells |
| PDE1A | 0 | 4.923149218 | 0.753 | 0.064 | 0 | 2 | Endothelial cells |
| KCNIP4 | 0 | 4.618132618 | 0.526 | 0.195 | 0 | 3 | Endothelial cells |
| CDH23 | 0 | 4.496021454 | 0.302 | 0.033 | 0 | 3 | Endothelial cells |
| PRUNE2 | 0 | 4.480997078 | 0.571 | 0.13 | 0 | 4 | Endothelial cells |
| CCDC102B | 0 | 4.45992751 | 0.76 | 0.097 | 0 | 2 | Endothelial cells |
| NEGR1 | 0 | 4.202377583 | 0.472 | 0.071 | 0 | 4 | Endothelial cells |
| COL6A3 | 0 | 3.94858893 | 0.71 | 0.058 | 0 | 2 | Endothelial cells |
| PRKG1 | 0 | 3.936825646 | 0.965 | 0.215 | 0 | 2 | Endothelial cells |
| CACNB2 | 0 | 3.907309065 | 0.679 | 0.281 | 0 | 4 | Endothelial cells |
| COL6A1 | 0 | 3.869309547 | 0.877 | 0.14 | 0 | 2 | Endothelial cells |
| COL3A1 | 0 | 3.862133706 | 0.882 | 0.101 | 0 | 2 | Endothelial cells |
| COL1A2 | 0 | 3.833819396 | 0.841 | 0.077 | 0 | 2 | Endothelial cells |
| IL1R1 | 0 | 3.768902443 | 0.722 | 0.089 | 0 | 2 | Endothelial cells |
| UNC5C | 0 | 3.643678967 | 0.628 | 0.231 | 0 | 4 | Endothelial cells |
| GPM6A | 0 | 3.623238157 | 0.581 | 0.094 | 0 | 3 | Endothelial cells |
| RAMP2 | 0 | 3.60308521 | 0.856 | 0.172 | 0 | 1 | Endothelial cells |
| ANK2 | 0 | 3.588977925 | 0.635 | 0.222 | 0 | 4 | Endothelial cells |
| MMP1 | 0 | 3.495191426 | 0.808 | 0.062 | 0 | 2 | Endothelial cells |
| COL6A2 | 0 | 3.467811139 | 0.919 | 0.256 | 0 | 2 | Endothelial cells |
| IFI27 | 0 | 3.319570983 | 0.84 | 0.21 | 0 | 1 | Endothelial cells |
| IL33 | 0 | 3.28314666 | 0.352 | 0.044 | 0 | 3 | Endothelial cells |
| VWA1 | 0 | 3.246644887 | 0.79 | 0.194 | 0 | 1 | Endothelial cells |
| TIMP1 | 0 | 3.212150335 | 0.975 | 0.344 | 0 | 2 | Endothelial cells |
| RBP7 | 0 | 3.195138111 | 0.679 | 0.135 | 0 | 1 | Endothelial cells |
| ITM2A | 0 | 3.174540658 | 0.651 | 0.092 | 0 | 1 | Endothelial cells |
| ZEB2 | 0 | 3.165786936 | 0.81 | 0.177 | 0 | 2 | Endothelial cells |
| STK32B | 0 | 3.136606098 | 0.773 | 0.196 | 0 | 5 | Endothelial cells |
| ANKS1B | 0 | 3.107140575 | 0.559 | 0.139 | 0 | 4 | Endothelial cells |
| FIGN | 0 | 3.08204209 | 0.473 | 0.088 | 0 | 4 | Endothelial cells |
| FLRT2 | 0 | 3.013420049 | 0.589 | 0.113 | 0 | 3 | Endothelial cells |
| CLEC14A | 0 | 2.960461531 | 0.818 | 0.19 | 0 | 1 | Endothelial cells |
| KLF2 | 0 | 2.935572226 | 0.812 | 0.257 | 0 | 1 | Endothelial cells |
| LPAR1 | 0 | 2.868448517 | 0.485 | 0.084 | 0 | 4 | Endothelial cells |
| MT2A | 0 | 2.834084762 | 0.997 | 0.357 | 0 | 2 | Endothelial cells |
| PCBP3 | 0 | 2.826138624 | 0.573 | 0.157 | 0 | 4 | Endothelial cells |
| SLC9A3R2 | 0 | 2.727366297 | 0.865 | 0.298 | 0 | 1 | Endothelial cells |
| SOX18 | 0 | 2.700902772 | 0.776 | 0.209 | 0 | 1 | Endothelial cells |
| PGM5 | 0 | 2.683688806 | 0.526 | 0.185 | 0 | 3 | Endothelial cells |
| COL27A1 | 0 | 2.653460186 | 0.59 | 0.154 | 0 | 5 | Endothelial cells |
| PCNX2 | 0 | 2.635892059 | 0.546 | 0.134 | 0 | 4 | Endothelial cells |
| VAMP5 | 0 | 2.625897349 | 0.74 | 0.18 | 0 | 1 | Endothelial cells |
| MARCKSL1 | 0 | 2.606053844 | 0.735 | 0.187 | 0 | 1 | Endothelial cells |
| EPB41L3 | 0 | 2.602341575 | 0.656 | 0.205 | 0 | 3 | Endothelial cells |
| P2RY8 | 0 | 2.551527795 | 0.4 | 0.26 | 0 | 0 | Endothelial cells |
| NEK10 | 0 | 2.543727065 | 0.675 | 0.24 | 0 | 5 | Endothelial cells |
| RNASE1 | 0 | 2.538655985 | 0.826 | 0.199 | 0 | 1 | Endothelial cells |
| ADAMTSL1 | 0 | 2.537437492 | 0.453 | 0.147 | 0 | 3 | Endothelial cells |
| LGALS1 | 0 | 2.48350148 | 0.967 | 0.331 | 0 | 2 | Endothelial cells |
| HYAL2 | 0 | 2.478647317 | 0.795 | 0.231 | 0 | 1 | Endothelial cells |
| DOCK8 | 0 | 2.471716596 | 0.715 | 0.262 | 0 | 5 | Endothelial cells |
| MIR99AHG | 0 | 2.461608134 | 0.905 | 0.441 | 0 | 3 | Endothelial cells |
| BNC2 | 0 | 2.437261074 | 0.666 | 0.226 | 0 | 3 | Endothelial cells |
| PID1 | 0 | 2.388574215 | 0.605 | 0.149 | 0 | 4 | Endothelial cells |
| LRRC1 | 0 | 2.379033626 | 0.473 | 0.144 | 0 | 3 | Endothelial cells |
| MYL12B | 0 | 2.372273709 | 0.873 | 0.315 | 0 | 1 | Endothelial cells |
| PLSCR4 | 0 | 2.369515116 | 0.456 | 0.16 | 0 | 3 | Endothelial cells |
| BCAM | 0 | 2.343245252 | 0.724 | 0.174 | 0 | 1 | Endothelial cells |
| KIT | 1.92E-283 | 2.30411901 | 0.555 | 0.183 | 5.82E-279 | 5 | Endothelial cells |
| GNG11 | 0 | 2.279783416 | 0.89 | 0.347 | 0 | 1 | Endothelial cells |
| SRP14 | 0 | 2.270122294 | 0.889 | 0.336 | 0 | 1 | Endothelial cells |
| S100A10 | 0 | 2.239541209 | 0.834 | 0.276 | 0 | 1 | Endothelial cells |
| BMP6 | 0 | 2.229593974 | 0.467 | 0.171 | 0 | 3 | Endothelial cells |
| SLC38A1 | 3.19E-252 | 2.22591055 | 0.546 | 0.191 | 9.68E-248 | 5 | Endothelial cells |
| ADAM19 | 0 | 2.201665246 | 0.766 | 0.294 | 0 | 5 | Endothelial cells |
| DNASE1L3 | 0 | 2.163449397 | 0.405 | 0.131 | 0 | 3 | Endothelial cells |
| IFITM2 | 0 | 2.151279327 | 0.836 | 0.27 | 0 | 1 | Endothelial cells |
| MAPK10 | 0 | 2.133596632 | 0.602 | 0.216 | 0 | 4 | Endothelial cells |
| ST3GAL6 | 0 | 2.042825488 | 0.867 | 0.459 | 0 | 5 | Endothelial cells |
| NSD2 | 0 | 2.027840643 | 0.607 | 0.212 | 0 | 4 | Endothelial cells |
| ARHGEF28 | 1.52E-301 | 2.012371284 | 0.825 | 0.438 | 4.61E-297 | 5 | Endothelial cells |
| UBB | 0 | 1.993934233 | 0.838 | 0.285 | 0 | 1 | Endothelial cells |
| VEGFA | 0 | 1.963445097 | 0.557 | 0.171 | 0 | 4 | Endothelial cells |
| AL356124.1 | 3.57E-263 | 1.944661637 | 0.743 | 0.344 | 1.08E-258 | 5 | Endothelial cells |
| MYOF | 0 | 1.932252316 | 0.685 | 0.335 | 0 | 3 | Endothelial cells |
| SLC4A7 | 0 | 1.905570018 | 0.918 | 0.537 | 0 | 5 | Endothelial cells |
| PDGFD | 0 | 1.890799752 | 0.641 | 0.49 | 0 | 0 | Endothelial cells |
| PVT1 | 0 | 1.880717861 | 0.602 | 0.21 | 0 | 4 | Endothelial cells |
| SMAD1 | 0 | 1.872509727 | 0.869 | 0.448 | 0 | 5 | Endothelial cells |
| PARD3 | 0 | 1.872402261 | 0.765 | 0.366 | 0 | 4 | Endothelial cells |
| INPP5D | 0 | 1.862920693 | 0.596 | 0.467 | 0 | 0 | Endothelial cells |
| PGF | 0 | 1.818982552 | 0.711 | 0.261 | 0 | 5 | Endothelial cells |
| CPNE5 | 0 | 1.7807257 | 0.497 | 0.38 | 0 | 0 | Endothelial cells |
| OSBPL10 | 0 | 1.76245203 | 0.857 | 0.443 | 0 | 5 | Endothelial cells |
| GRB10 | 0 | 1.759716538 | 0.943 | 0.811 | 0 | 0 | Endothelial cells |
| PIK3R3 | 0 | 1.741206745 | 0.791 | 0.637 | 0 | 0 | Endothelial cells |
| STOX2 | 0 | 1.722232784 | 0.727 | 0.396 | 0 | 3 | Endothelial cells |
| PLXDC2 | 0 | 1.707804123 | 0.822 | 0.446 | 0 | 3 | Endothelial cells |
| PRR16 | 0 | 1.651064301 | 0.6 | 0.208 | 0 | 4 | Endothelial cells |
| PLXNA2 | 0 | 1.626533615 | 0.733 | 0.62 | 0 | 0 | Endothelial cells |
| RASGRF2 | 0 | 1.602737735 | 0.741 | 0.62 | 0 | 0 | Endothelial cells |
| ELMO1 | 0 | 1.589946434 | 0.937 | 0.805 | 0 | 0 | Endothelial cells |
| ST6GALNAC3 | 0 | 1.587819095 | 0.899 | 0.766 | 0 | 0 | Endothelial cells |
| UTY | 0 | 1.533151232 | 0.41 | 0.248 | 0 | 0 | Endothelial cells |
| ANO2 | 0 | 1.52748786 | 0.743 | 0.532 | 0 | 0 | Endothelial cells |
| FLT1 | 0 | 1.527485272 | 0.879 | 0.762 | 0 | 0 | Endothelial cells |
| LDLRAD4 | 0 | 1.503809885 | 0.79 | 0.64 | 0 | 0 | Endothelial cells |
| SVIL | 2.70E-266 | 1.393211305 | 0.91 | 0.649 | 8.20E-262 | 5 | Endothelial cells |
| CSGALNACT1 | 1.22E-264 | 1.390810771 | 0.921 | 0.661 | 3.69E-260 | 5 | Endothelial cells |
| LAMA4 | 2.08E-306 | 1.37728265 | 0.942 | 0.668 | 6.32E-302 | 5 | Endothelial cells |
| DOCK4 | 0 | 1.351521443 | 0.993 | 0.921 | 0 | 5 | Endothelial cells |
| BTNL9 | 0 | 1.338695226 | 0.521 | 0.372 | 0 | 0 | Endothelial cells |
| GALNT18 | 0 | 1.260522115 | 0.781 | 0.663 | 0 | 0 | Endothelial cells |
| TSHZ2 | 0 | 1.195569647 | 0.712 | 0.596 | 0 | 0 | Endothelial cells |
| DIAPH2 | 2.11E-269 | 1.122136362 | 0.968 | 0.821 | 6.42E-265 | 5 | Endothelial cells |
| MAGI1 | 1.28E-277 | 1.002955202 | 0.985 | 0.912 | 3.88E-273 | 5 | Endothelial cells |
| MEG8 | 0 | 0.7181919367 | 0.512 | 0.228 | 0 | 0 | Endothelial cells |
| RIMS2 | 0 | 0.2825754388 | 0.409 | 0.19 | 0 | 0 | Endothelial cells |
| CCSER1 | 0 | 0.2663755521 | 0.686 | 0.467 | 0 | 0 | Endothelial cells |
| CD247 | 0 | 6.296154097 | 0.806 | 0.031 | 0 | 0 | Immune clusters |
| SKAP1 | 0 | 4.473454197 | 0.746 | 0.087 | 0 | 0 | Immune clusters |
| ETS1 | 0 | 4.805296549 | 0.764 | 0.135 | 0 | 0 | Immune clusters |
| STAT4 | 0 | 3.578681039 | 0.809 | 0.207 | 0 | 0 | Immune clusters |
| SLC38A1 | 0 | 4.723296926 | 0.692 | 0.091 | 0 | 0 | Immune clusters |
| CD96 | 0 | 7.011385723 | 0.602 | 0.013 | 0 | 0 | Immune clusters |
| SYNE2 | 0 | 4.591363444 | 0.73 | 0.144 | 0 | 0 | Immune clusters |
| ITK | 0 | 5.865184231 | 0.589 | 0.032 | 0 | 0 | Immune clusters |
| PPP1R16B | 0 | 5.400533275 | 0.624 | 0.078 | 0 | 0 | Immune clusters |
| RNF125 | 0 | 5.206502117 | 0.637 | 0.104 | 0 | 0 | Immune clusters |
| FYN | 0 | 3.603358597 | 0.898 | 0.415 | 0 | 0 | Immune clusters |
| CNOT6L | 0 | 4.284116638 | 0.815 | 0.338 | 0 | 0 | Immune clusters |
| BCL11B | 0 | 6.074345206 | 0.484 | 0.014 | 0 | 0 | Immune clusters |
| PARP8 | 0 | 3.41776448 | 0.834 | 0.373 | 0 | 0 | Immune clusters |
| RORA | 0 | 2.973009785 | 0.711 | 0.257 | 0 | 0 | Immune clusters |
| SYTL3 | 0 | 3.13491488 | 0.792 | 0.351 | 0 | 0 | Immune clusters |
| CARD11 | 0 | 3.185192337 | 0.568 | 0.13 | 0 | 0 | Immune clusters |
| BACH2 | 0 | 5.627574368 | 0.523 | 0.086 | 0 | 0 | Immune clusters |
| SLFN12L | 0 | 5.753416632 | 0.455 | 0.018 | 0 | 0 | Immune clusters |
| CD69 | 0 | 2.692315045 | 0.575 | 0.145 | 0 | 0 | Immune clusters |
| FCN1 | 0 | 4.870062339 | 0.924 | 0.049 | 0 | 1 | Immune clusters |
| S100A9 | 0 | 5.583725231 | 0.902 | 0.104 | 0 | 1 | Immune clusters |
| SERPINA1 | 0 | 4.169554356 | 0.883 | 0.105 | 0 | 1 | Immune clusters |
| BCL2A1 | 0 | 3.740784401 | 0.896 | 0.136 | 0 | 1 | Immune clusters |
| VCAN | 0 | 2.850157016 | 0.822 | 0.075 | 0 | 1 | Immune clusters |
| CSTA | 0 | 4.763102299 | 0.785 | 0.055 | 0 | 1 | Immune clusters |
| FGR | 0 | 2.41701419 | 0.889 | 0.16 | 0 | 1 | Immune clusters |
| AQP9 | 0 | 5.201837538 | 0.776 | 0.047 | 0 | 1 | Immune clusters |
| S100A8 | 0 | 6.04475445 | 0.801 | 0.073 | 0 | 1 | Immune clusters |
| LST1 | 0 | 3.342059728 | 0.916 | 0.193 | 0 | 1 | Immune clusters |
| LYZ | 0 | 3.905317705 | 0.943 | 0.224 | 0 | 1 | Immune clusters |
| AIF1 | 0 | 2.60434484 | 0.923 | 0.21 | 0 | 1 | Immune clusters |
| IFI30 | 0 | 2.949418338 | 0.953 | 0.24 | 0 | 1 | Immune clusters |
| SLC11A1 | 0 | 1.499724055 | 0.907 | 0.197 | 0 | 1 | Immune clusters |
| CFD | 0 | 2.987468036 | 0.845 | 0.138 | 0 | 1 | Immune clusters |
| KYNU | 0 | 2.767676748 | 0.967 | 0.265 | 0 | 1 | Immune clusters |
| IL1B | 0 | 3.630458265 | 0.875 | 0.177 | 0 | 1 | Immune clusters |
| SMIM25 | 0 | 4.650533129 | 0.756 | 0.06 | 0 | 1 | Immune clusters |
| KLF4 | 0 | 2.188249315 | 0.871 | 0.19 | 0 | 1 | Immune clusters |
| NCF2 | 0 | 2.509921951 | 0.809 | 0.13 | 0 | 1 | Immune clusters |
| C1QA | 0 | 4.948799381 | 0.782 | 0.098 | 0 | 2 | Immune clusters |
| C1QC | 0 | 5.977650104 | 0.728 | 0.045 | 0 | 2 | Immune clusters |
| C1QB | 0 | 5.04622534 | 0.733 | 0.077 | 0 | 2 | Immune clusters |
| LGMN | 0 | 2.591424584 | 0.777 | 0.149 | 0 | 2 | Immune clusters |
| DAB2 | 0 | 3.098677799 | 0.723 | 0.109 | 0 | 2 | Immune clusters |
| MRC1 | 0 | 1.871738854 | 0.758 | 0.155 | 0 | 2 | Immune clusters |
| SELENOP | 0 | 4.583097787 | 0.719 | 0.132 | 0 | 2 | Immune clusters |
| CD14 | 0 | 3.155120266 | 0.748 | 0.192 | 0 | 2 | Immune clusters |
| SLC1A3 | 0 | 1.300690645 | 0.849 | 0.297 | 0 | 2 | Immune clusters |
| SLCO2B1 | 0 | 0.8290591158 | 0.752 | 0.2 | 0 | 2 | Immune clusters |
| FPR3 | 0 | 1.871621225 | 0.649 | 0.117 | 0 | 2 | Immune clusters |
| STAB1 | 0 | 0.9732754403 | 0.752 | 0.224 | 0 | 2 | Immune clusters |
| CD163 | 0 | 1.805563454 | 0.811 | 0.284 | 0 | 2 | Immune clusters |
| HLA-DRA | 0 | 3.054559717 | 0.91 | 0.391 | 0 | 2 | Immune clusters |
| MS4A4A | 0 | 2.491380295 | 0.67 | 0.155 | 0 | 2 | Immune clusters |
| MS4A7 | 0 | 1.927708875 | 0.853 | 0.338 | 0 | 2 | Immune clusters |
| MERTK | 0 | 0.7840930855 | 0.737 | 0.224 | 0 | 2 | Immune clusters |
| APOE | 0 | 3.9696107 | 0.615 | 0.106 | 0 | 2 | Immune clusters |
| MAFB | 0 | 2.13402217 | 0.778 | 0.27 | 0 | 2 | Immune clusters |
| MARCKS | 0 | 2.241930695 | 0.742 | 0.235 | 0 | 2 | Immune clusters |
| MEG8 | 0 | 1.960907689 | 0.583 | 0.13 | 0 | 3 | Immune clusters |
| FAM155A | 0 | 1.602667271 | 0.635 | 0.206 | 0 | 3 | Immune clusters |
| FRMD4A | 0 | 2.973497616 | 0.778 | 0.358 | 0 | 3 | Immune clusters |
| CCSER1 | 0 | 1.942936705 | 0.749 | 0.353 | 0 | 3 | Immune clusters |
| RALYL | 0 | 1.784240677 | 0.486 | 0.124 | 0 | 3 | Immune clusters |
| PLXDC2 | 0 | 2.524306932 | 0.948 | 0.589 | 0 | 3 | Immune clusters |
| SLC1A3 | 0 | 2.257480436 | 0.692 | 0.345 | 0 | 3 | Immune clusters |
| DOCK4 | 0 | 2.760363734 | 0.84 | 0.496 | 0 | 3 | Immune clusters |
| KCNMA1 | 0 | 2.648801071 | 0.627 | 0.289 | 0 | 3 | Immune clusters |
| FMNL2 | 0 | 2.689132953 | 0.718 | 0.385 | 0 | 3 | Immune clusters |
| RBM47 | 0 | 2.21918267 | 0.834 | 0.501 | 0 | 3 | Immune clusters |
| KCNQ3 | 0 | 3.9427693 | 0.453 | 0.121 | 0 | 3 | Immune clusters |
| LRMDA | 0 | 2.471457576 | 0.947 | 0.615 | 0 | 3 | Immune clusters |
| MEG3 | 0 | 0.6131170372 | 0.539 | 0.211 | 0 | 3 | Immune clusters |
| SLC8A1 | 0 | 2.303358837 | 0.821 | 0.505 | 0 | 3 | Immune clusters |
| RIMS2 | 0 | 1.935964932 | 0.435 | 0.128 | 0 | 3 | Immune clusters |
| MERTK | 0 | 2.767866439 | 0.575 | 0.273 | 0 | 3 | Immune clusters |
| SYT1 | 0 | 1.720747969 | 0.481 | 0.187 | 0 | 3 | Immune clusters |
| CSGALNACT1 | 0 | 2.496597444 | 0.588 | 0.297 | 0 | 3 | Immune clusters |
| ARHGAP24 | 0 | 2.503354091 | 0.732 | 0.442 | 0 | 3 | Immune clusters |
| FN1 | 0 | 5.907243765 | 0.88 | 0.143 | 0 | 4 | Immune clusters |
| PTPRG | 0 | 5.503779846 | 0.861 | 0.148 | 0 | 4 | Immune clusters |
| PRKG1 | 0 | 4.879490638 | 0.885 | 0.194 | 0 | 4 | Immune clusters |
| COL4A1 | 0 | 5.689148636 | 0.814 | 0.14 | 0 | 4 | Immune clusters |
| EBF1 | 0 | 5.681876877 | 0.782 | 0.118 | 0 | 4 | Immune clusters |
| COL4A2 | 0 | 5.496712507 | 0.819 | 0.176 | 0 | 4 | Immune clusters |
| CALD1 | 0 | 5.872603528 | 0.773 | 0.145 | 0 | 4 | Immune clusters |
| DLC1 | 0 | 4.468438222 | 0.865 | 0.24 | 0 | 4 | Immune clusters |
| COL3A1 | 0 | 6.163366381 | 0.662 | 0.061 | 0 | 4 | Immune clusters |
| SOX5 | 0 | 4.378878377 | 0.757 | 0.163 | 0 | 4 | Immune clusters |
| COL6A2 | 0 | 5.403523874 | 0.686 | 0.098 | 0 | 4 | Immune clusters |
| COL6A1 | 0 | 6.048798011 | 0.655 | 0.077 | 0 | 4 | Immune clusters |
| TPSB2 | 0 | 0.9172572567 | 0.644 | 0.073 | 0 | 4 | Immune clusters |
| PALM2-AKAP2 | 0 | 3.553785778 | 0.768 | 0.201 | 0 | 4 | Immune clusters |
| CPA3 | 0 | 1.746127786 | 0.613 | 0.059 | 0 | 4 | Immune clusters |
| TPSAB1 | 0 | 1.681361106 | 0.606 | 0.063 | 0 | 4 | Immune clusters |
| COL1A2 | 0 | 6.134803781 | 0.587 | 0.067 | 0 | 4 | Immune clusters |
| TIMP3 | 0 | 4.625231754 | 0.68 | 0.161 | 0 | 4 | Immune clusters |
| COL6A3 | 0 | 6.199121043 | 0.56 | 0.042 | 0 | 4 | Immune clusters |
| IGFBP7 | 0 | 5.530845972 | 0.829 | 0.323 | 0 | 4 | Immune clusters |
| SLC24A3 | 0 | 6.337401702 | 0.962 | 0.05 | 0 | 5 | Immune clusters |
| KIT | 0 | 5.667537711 | 0.952 | 0.058 | 0 | 5 | Immune clusters |
| TPSB2 | 0 | 7.113834721 | 0.965 | 0.075 | 0 | 5 | Immune clusters |
| MS4A2 | 0 | 5.702234885 | 0.9 | 0.043 | 0 | 5 | Immune clusters |
| CPA3 | 0 | 6.330190297 | 0.912 | 0.061 | 0 | 5 | Immune clusters |
| TPSAB1 | 0 | 6.377604493 | 0.906 | 0.065 | 0 | 5 | Immune clusters |
| GATA2 | 0 | 5.605237323 | 0.868 | 0.047 | 0 | 5 | Immune clusters |
| IL1RL1 | 0 | 6.034304397 | 0.815 | 0.024 | 0 | 5 | Immune clusters |
| HPGDS | 0 | 5.073223525 | 0.873 | 0.084 | 0 | 5 | Immune clusters |
| VWA5A | 0 | 5.546974569 | 0.806 | 0.047 | 0 | 5 | Immune clusters |
| SLC18A2 | 0 | 4.866002058 | 0.858 | 0.107 | 0 | 5 | Immune clusters |
| NTM | 0 | 5.78124147 | 0.799 | 0.05 | 0 | 5 | Immune clusters |
| RHEX | 0 | 5.181167949 | 0.777 | 0.03 | 0 | 5 | Immune clusters |
| CLU | 0 | 5.344480466 | 0.842 | 0.095 | 0 | 5 | Immune clusters |
| IL18R1 | 0 | 4.5207653 | 0.85 | 0.107 | 0 | 5 | Immune clusters |
| RGS13 | 0 | 4.858687214 | 0.795 | 0.068 | 0 | 5 | Immune clusters |
| ADAM12 | 0 | 5.290270243 | 0.736 | 0.043 | 0 | 5 | Immune clusters |
| MAST4 | 0 | 3.519453621 | 0.807 | 0.13 | 0 | 5 | Immune clusters |
| BACE2 | 0 | 4.703030145 | 0.726 | 0.053 | 0 | 5 | Immune clusters |
| CADPS | 0 | 4.749164934 | 0.747 | 0.087 | 0 | 5 | Immune clusters |
| RALYL | 0 | 4.254658089 | 0.885 | 0.176 | 0 | 6 | Immune clusters |
| LSAMP | 0 | 4.824823266 | 0.749 | 0.112 | 0 | 6 | Immune clusters |
| RYR2 | 0 | 4.897339665 | 0.654 | 0.081 | 0 | 6 | Immune clusters |
| AGBL4 | 0 | 3.971726452 | 0.649 | 0.09 | 0 | 6 | Immune clusters |
| CTNND2 | 0 | 4.297008605 | 0.605 | 0.074 | 0 | 6 | Immune clusters |
| TENM3 | 0 | 4.45163783 | 0.59 | 0.073 | 0 | 6 | Immune clusters |
| SORCS3 | 0 | 5.545972219 | 0.527 | 0.033 | 0 | 6 | Immune clusters |
| RIMBP2 | 0 | 5.517583628 | 0.537 | 0.058 | 0 | 6 | Immune clusters |
| HCN1 | 0 | 4.401031053 | 0.527 | 0.056 | 0 | 6 | Immune clusters |
| TMEM178B | 0 | 5.297085706 | 0.505 | 0.051 | 0 | 6 | Immune clusters |
| SRRM3 | 0 | 5.144706051 | 0.38 | 0.028 | 0 | 6 | Immune clusters |
| MIR137HG | 2.63E-307 | 4.605779429 | 0.432 | 0.037 | 8.00E-303 | 6 | Immune clusters |
| MEG8 | 5.30E-302 | 3.983047584 | 0.861 | 0.2 | 1.61E-297 | 6 | Immune clusters |
| DPP6 | 5.80E-300 | 4.969952152 | 0.717 | 0.133 | 1.76E-295 | 6 | Immune clusters |
| FAM163A | 1.75E-297 | 6.473303802 | 0.354 | 0.026 | 5.31E-293 | 6 | Immune clusters |
| FAM155A | 9.74E-297 | 4.384845299 | 0.912 | 0.272 | 2.96E-292 | 6 | Immune clusters |
| GRIK2 | 1.31E-289 | 3.919891894 | 0.6 | 0.084 | 3.98E-285 | 6 | Immune clusters |
| UNC5C | 9.03E-289 | 4.384794036 | 0.749 | 0.149 | 2.74E-284 | 6 | Immune clusters |
| RIMS2 | 2.07E-288 | 3.934859577 | 0.788 | 0.172 | 6.28E-284 | 6 | Immune clusters |
| FAM135B | 2.42E-284 | 5.245102299 | 0.38 | 0.032 | 7.37E-280 | 6 | Immune clusters |

**Supplementary Table 11.** Top markers of reclustered tumor cells.

| Gene | Selected for AU | Signature type |
|---|---|---|
| ALDO1* | Yes | HIF1α |
| ALDOC* | Yes | HIF1α |
| AMIGO2* | Yes | HIF1α |
| ANG* | Yes | HIF1α |
| ENO1* | Yes | HIF1α |
| FABP3* | Yes | HIF1α |
| FGF11* | Yes | HIF1α |
| HK2* | Yes | HIF1α |
| ICAM1* | Yes | HIF1α |
| LDHA* | Yes | HIF1α |
| MAPK7* | Yes | HIF1α |
| PGAM1* | Yes | HIF1α |
| TIMP3* | Yes | HIF1α |
| CLDN3* | Yes | HIF2α |
| CXCL2* | Yes | HIF2α |
| ECM1* | Yes | HIF2α |
| FOS* | Yes | HIF2α |
| FOXD1* | Yes | HIF2α |
| HAND1* | Yes | HIF2α |
| HOXA10* | Yes | HIF2α |
| HOXA11* | Yes | HIF2α |
| JUNB* | Yes | HIF2α |
| KLF4* | Yes | HIF2α |
| NDNF* | Yes | HIF2α |
| SLC2A1* | Yes | HIF2α |
| STC1* | Yes | HIF2α |
| VIP* | Yes | HIF2α |

**Supplementary Table 12.** AUCell signatures used to define HIF1A and HIF2A signatures (indicate the source references for curating these signatures)**.** *included in AUCell signature plots. HIF1A=13/63 and HIF2A=14/102 References: Downes, Nicholas L et al. "Differential but Complementary HIF1α and HIF2α Transcriptional Regulation." Molecular therapy : the journal of the American Society of Gene Therapy vol. 26,7 (2018): 1735-1745. doi:10.1016/j.ymthe.2018.05.004; Cho, Hyejin et al. "On-target efficacy of a HIF-2α antagonist in preclinical kidney cancer models." Nature vol. 539,7627 (2016): 107-111. doi:10.1038/nature19795; López-Jiménez, Elena et al. "Research resource: Transcriptional profiling reveals different pseudohypoxic signatures in SDHB and VHL-related pheochromocytomas." Molecular endocrinology (Baltimore, Md.) vol. 24,12 (2010): 2382-91. doi:10.1210/me.2010-0256

| Gene | Signature type |
|---|---|
| SOX10 | SCP |
| PLP1 | SCP |
| SOX2 | SCP |
| MPZ | SCP |
| ERBB3 | SCP |
| FOXD3 | SCP |
| CDH19 | SCP |
| PTPRZ1 | SCP |
| ASCL1 | Bridge |
| HTR3A | Bridge |
| THBD | Bridge |
| DLL3 | Bridge |
| SOX11 | Bridge |
| SCG3 | Early Chromaffin2 |
| SLC35D3 | Early Chromaffin2 |
| LGR5 | Early Chromaffin2 |
| C1QL1 | Early Chromaffin2 |
| PNMT | Mature Chromaffin |
| PENK | Mature Chromaffin |
| EGR1 | Mature Chromaffin |
| JUNB | Mature Chromaffin |
| GATA2 | Mature Chromaffin |
| NPY | Mature Chromaffin |
| GAP43 | Sympathoblast |
| STMN4 | Sympathoblast |
| ELAVL3 | Sympathoblast |
| PRPH | Sympathoblast |
| STMN2 | Sympathoblast |

**Supplementary Table 13.** AUCell signatures used to define the chromaffin cell developmental lineages, encompassing unique genes to each cell subtypes.

**Supplementary Table 14.** Monocle 3 heat map markers

Table provided in synapse.org/PPGLorganoids.

| Transcription Factors | PDT | ST_N | ST_H | LT_N |
|---|---|---|---|---|
| JAZF1 | 0.48575622 | 0.340436419 | 0.279923057 | 0.279046048 |
| CREB5 | 0.357688742 | 0.232730373 | 0.155625865 | 0.176205283 |
| PBX3 | 0.15644946 | 0.141239692 | 0.140097395 | 0.146799665 |
| TRPS1 | 0.064181562 | 0.078012329 | 0.101865793 | 0.119545637 |
| FOXN3 | 0.163420845 | 0.105659788 | 0.12165717 | 0.106169311 |
| ENO1 | 0.09948651 | 0.1533218 | 0.195682322 | 0.10171365 |
| EBF2 | 0.019612499 | 0.038847471 | 0.041168871 | 0.083197005 |
| ID1 | 0.01257666 | 0.135094669 | 0.108387106 | 0.081973775 |
| ELF2 | 0.037387473 | 0.037317436 | 0.095194041 | 0.076818535 |
| PHLDA2 | 0.000917451 | 0.003286267 | 0.021099116 | 0.071672842 |
| SOX9 | 0.061873551 | 0.102210972 | 0.04763263 | 0.069147957 |
| CEBPD | 0.081617998 | 0.078303422 | 0.100674811 | 0.060342588 |
| KLF6 | 0.014000451 | 0.023176162 | 0.025608704 | 0.058847583 |
| JDP2 | 0.074092846 | 0.076543315 | 0.026980033 | 0.055774814 |
| SOX4 | 0.047717499 | 0.072467075 | 0.057879371 | 0.050705569 |
| SMAD1 | 0.006650993 | 0.017872734 | 0.030937281 | 0.048894067 |
| ZNF189 | 0.272648336 | 0.038328292 | 0.021446932 | 0.048229687 |
| JUNB | 0.094020162 | 0.05662962 | 0.061609633 | 0.04128937 |
| GATA3 | 0.073792871 | 0.078557483 | 0.045415513 | 0.041258004 |
| MEIS2 | 0.026097642 | 0.024221885 | 0.025188018 | 0.039727944 |
| HOXA9 | 0.021020649 | 0.021073962 | 0.060053455 | 0.039651980 |
| SAP30 | 0.033032505 | 0.038627015 | 0.080050461 | 0.039264485 |
| CREM | 0.033036625 | 0.041406633 | 0.046834799 | 0.038856408 |
| PPARG | 0.009773953 | 0.018266538 | 0.025389484 | 0.038175924 |
| JUN | 0.092762326 | 0.05139779 | 0.06165114 | 0.037634808 |
| GATA2 | 0.040154097 | 0.083682444 | 0.034270898 | 0.037464162 |
| HLF | 0.030215783 | 0.016975972 | 0.029849574 | 0.036732198 |
| PHOX2A | 0.038895133 | 0.074170152 | 0.048831417 | 0.036436138 |
| POU2F1 | 0.059715353 | 0.035516703 | 0.03701731 | 0.036220795 |
| HMGA2 | 0.003979437 | 0.010127406 | 0.014723678 | 0.034791633 |
| STAT3 | 0.012818495 | 0.020467234 | 0.049458887 | 0.034307039 |
| PHOX2B | 0.04317274 | 0.069811283 | 0.054818025 | 0.032878373 |
| TLX2 | 0.035278598 | 0.059338533 | 0.036595337 | 0.03142804 |
| NFKB1 | 0.003021166 | 0.006654174 | 0.01263642 | 0.031121299 |
| FOS | 0.245252208 | 0.041898056 | 0.034188116 | 0.027227529 |
| ASCL1 | 0.01420628 | 0.024491741 | 0.051328671 | 0.026637945 |
| FOSL2 | 0.034204623 | 0.022864165 | 0.034154099 | 0.026594375 |
| ELF1 | 0.032973077 | 0.026694078 | 0.034177473 | 0.025512002 |
| DDIT3 | 0.038723282 | 0.025449689 | 0.026704581 | 0.024709677 |
| EPAS1 | 0.011526087 | 0.016881247 | 0.032072581 | 0.02456736 |
| SREBF1 | 0.031293339 | 0.0675101 | 0.021118701 | 0.023950788 |
| PRRX1 | 0.006268088 | 0.009369072 | 0.031102144 | 0.019352783 |
| EGR1 | 0.140613566 | 0.012535155 | 0.014305973 | 0.01654038 |
| KDM5B | 0.014482987 | 0.007587987 | 0.017897419 | 0.015680749 |
| BCL11A | 0.012525687 | 0.002081175 | 0.010120355 | 0.015666164 |
| FOSB | 0.075266843 | 0.009990141 | 0.011635322 | 0.013129519 |
| KLF9 | 0.108768137 | 0.01616354 | 0.009799142 | 0.01123164 |
| NPAS2 | 0.000759692 | 0.005716735 | 0.003784513 | 0.010958937 |
| SHOX2 | 0.022394332 | 0.021714368 | 0.020734244 | 0.007943011 |
| ISL1 | 0.009229774 | 0.013109818 | 0.00652103 | 0.007876383 |
| EGR4 | 0.151548385 | 0.00521165 | 0.008794117 | 0.007366917 |
| RELB | 0.001888317 | 0.002453756 | 0.005547285 | 0.006915762 |
| HES4 | 0.000548992 | 0.001334537 | 0.001397501 | 0.006574550 |
| ATF3 | 0.042195442 | 0.004119535 | 0.007270905 | 0.006327960 |
| EZH2 | 0.003216554 | 0.00227355 | 0.004909891 | 0.006024548 |
| POLR2A | 0.002328275 | 0.004840196 | 0.008775286 | 0.005311195 |
| SOX7 | 0.075302204 | 0.0211367 | 0.011226667 | 0.005138265 |
| TFAP2C | 0.004047389 | 0.012193121 | 0.009195814 | 0.004040123 |
| REL | 0.001268841 | 0.001086197 | 0.002420765 | 0.003451798 |
| NR4A2 | 0.009148277 | 0.004952025 | 0.007647596 | 0.00202345 |
| HIST1H2BN | 0.020414116 | 0.002888924 | 0.004178299 | 0.001842185 |

**Supplementary Table 15.** Regulons associated with PPGLs and PDOs at distinct conditions analyzed by pySCENIC

| Cohort ID | SampleID_study labels | NOR_gsva | MES_gsva | MES-NOR Score | Sample Source Reference |
|---|---|---|---|---|---|
| TCGA | TCGA-P7-A5NY-01A | 0.211211157 | 0.166927193 | -0.044283965 | Fishbein et al ( PMID28162975) |
| TCGA | TCGA-P7-A5NY-05A | 0.406414727 | 0.177915123 | -0.228499603 | Fishbein et al ( PMID28162975) |
| TCGA | TCGA-P8-A5KC-01A | 0.13144194 | 0.19838261 | 0.066940669 | Fishbein et al ( PMID28162975) |
| TCGA | TCGA-P8-A6RX-01A | 0.123373997 | 0.127742675 | 0.004368678 | Fishbein et al ( PMID28162975) |
| TCGA | TCGA-P8-A5KD-01A | 0.269666841 | -0.023423795 | -0.293090636 | Fishbein et al ( PMID28162975) |
| TCGA | TCGA-PR-A5PF-01A | -0.129933566 | -0.235983489 | -0.106049923 | Fishbein et al ( PMID28162975) |
| TCGA | TCGA-PR-A5PG-01A | 0.274731734 | -0.014099995 | -0.288831728 | Fishbein et al ( PMID28162975) |
| TCGA | TCGA-PR-A5PH-01A | 0.217380969 | 0.034813159 | -0.18256781 | Fishbein et al ( PMID28162975) |
| TCGA | TCGA-QR-A6GO-01A | 0.10156372 | 0.016074155 | -0.085489565 | Fishbein et al ( PMID28162975) |
| TCGA | TCGA-QR-A6GR-01A | 0.045252953 | 0.215369253 | 0.1701163 | Fishbein et al ( PMID28162975) |
| TCGA | TCGA-QR-A6GS-01A | 0.288059609 | 0.28574503 | -0.002314579 | Fishbein et al ( PMID28162975) |
| TCGA | TCGA-QR-A6GT-01A | 0.137441069 | 0.199753162 | 0.062312094 | Fishbein et al ( PMID28162975) |
| TCGA | TCGA-QR-A6GU-01A | -0.135377095 | 0.205339676 | 0.340716771 | Fishbein et al ( PMID28162975) |
| TCGA | TCGA-QR-A6GW-01A | 0.350640968 | 0.082421599 | -0.268219369 | Fishbein et al ( PMID28162975) |
| TCGA | TCGA-QR-A6GX-01A | 0.061585165 | 0.306468702 | 0.244883537 | Fishbein et al ( PMID28162975) |
| TCGA | TCGA-QR-A6GY-01A | 0.146819692 | -0.018724286 | -0.165543978 | Fishbein et al ( PMID28162975) |
| TCGA | TCGA-P8-A6RY-01A | -0.478608299 | -0.277218754 | 0.201389544 | Fishbein et al ( PMID28162975) |
| TCGA | TCGA-QR-A6H0-01A | -0.510216558 | 0.227506597 | 0.737723154 | Fishbein et al ( PMID28162975) |
| TCGA | TCGA-QR-A6H1-01A | 0.487784655 | -0.240383024 | -0.728167679 | Fishbein et al ( PMID28162975) |
| TCGA | TCGA-QR-A6H2-01A | -0.20220381 | -0.15880371 | 0.0434001 | Fishbein et al ( PMID28162975) |
| TCGA | TCGA-QR-A6H3-01A | -0.471323464 | -0.291213716 | 0.180109748 | Fishbein et al ( PMID28162975) |
| TCGA | TCGA-QR-A6H5-01A | 0.208341517 | 0.12687235 | -0.081469167 | Fishbein et al ( PMID28162975) |
| TCGA | TCGA-QR-A6H6-01A | 0.081758582 | 0.123656137 | 0.041897555 | Fishbein et al ( PMID28162975) |
| TCGA | TCGA-QR-A6ZZ-01A | 0.387697571 | -0.288974806 | -0.676672377 | Fishbein et al ( PMID28162975) |
| TCGA | TCGA-QR-A700-01A | -0.47509003 | -0.229672361 | 0.245417668 | Fishbein et al ( PMID28162975) |
| TCGA | TCGA-QR-A702-01A | -0.288781572 | 0.035239866 | 0.324021438 | Fishbein et al ( PMID28162975) |
| TCGA | TCGA-QR-A703-01A | 0.309208189 | -0.391642808 | -0.700850996 | Fishbein et al ( PMID28162975) |
| TCGA | TCGA-QR-A705-01A | -0.28444123 | 0.203646095 | 0.488087325 | Fishbein et al ( PMID28162975) |
| TCGA | TCGA-QR-A706-01A | 0.29706885 | -0.049427119 | -0.346495969 | Fishbein et al ( PMID28162975) |
| TCGA | TCGA-QR-A707-01A | 0.1903567 | 0.152693244 | -0.037663456 | Fishbein et al ( PMID28162975) |
| TCGA | TCGA-QR-A708-01A | -0.256916497 | -0.439940803 | -0.183024306 | Fishbein et al ( PMID28162975) |
| TCGA | TCGA-QR-A70A-01A | 0.090273223 | 0.034014784 | -0.056258439 | Fishbein et al ( PMID28162975) |
| TCGA | TCGA-QR-A70C-01A | 0.20115974 | 0.145802552 | -0.055357188 | Fishbein et al ( PMID28162975) |
| TCGA | TCGA-QR-A70D-01A | -0.308299354 | 0.216400558 | 0.524699912 | Fishbein et al ( PMID28162975) |
| TCGA | TCGA-QR-A70E-01A | -0.463137084 | -0.494874922 | -0.031737838 | Fishbein et al ( PMID28162975) |
| TCGA | TCGA-QR-A70G-01B | 0.135860085 | 0.059552969 | -0.076307116 | Fishbein et al ( PMID28162975) |
| TCGA | TCGA-QR-A70H-01A | 0.096517706 | -0.117603739 | -0.214121445 | Fishbein et al ( PMID28162975) |
| TCGA | TCGA-QR-A70I-01A | 0.031797376 | 0.382364258 | 0.350566882 | Fishbein et al ( PMID28162975) |
| TCGA | TCGA-QR-A70J-01A | 0.440135018 | 0.294984188 | -0.14515083 | Fishbein et al ( PMID28162975) |
| TCGA | TCGA-QR-A70K-01A | -0.157222174 | 0.35941647 | 0.516638644 | Fishbein et al ( PMID28162975) |
| TCGA | TCGA-QR-A70M-01A | 0.410176392 | -0.447767843 | -0.857944236 | Fishbein et al ( PMID28162975) |
| TCGA | TCGA-QR-A70N-01A | 0.047989453 | -0.209680138 | -0.257669591 | Fishbein et al ( PMID28162975) |
| TCGA | TCGA-QR-A70O-01A | 0.086341092 | -0.064330682 | -0.150671774 | Fishbein et al ( PMID28162975) |
| TCGA | TCGA-QR-A70P-01A | -0.021547422 | 0.160624656 | 0.182172078 | Fishbein et al ( PMID28162975) |
| TCGA | TCGA-QR-A70R-01A | -0.291131582 | 0.129587792 | 0.420719374 | Fishbein et al ( PMID28162975) |
| TCGA | TCGA-QR-A70T-01A | -0.17952441 | 0.029471121 | 0.208995531 | Fishbein et al ( PMID28162975) |
| TCGA | TCGA-QR-A70U-01A | 0.087911382 | -0.045743612 | -0.133654995 | Fishbein et al ( PMID28162975) |
| TCGA | TCGA-QR-A70V-01A | -0.001927646 | -0.134412912 | -0.132485267 | Fishbein et al ( PMID28162975) |
| TCGA | TCGA-QR-A70W-01A | -0.058033019 | 0.013047049 | 0.071080068 | Fishbein et al ( PMID28162975) |
| TCGA | TCGA-QR-A70X-01A | -0.073329068 | 0.155742523 | 0.229071591 | Fishbein et al ( PMID28162975) |
| TCGA | TCGA-QR-A7IN-01A | 0.299527427 | -0.231181947 | -0.530709374 | Fishbein et al ( PMID28162975) |
| TCGA | TCGA-QR-A7IP-01A | -0.207037554 | -0.141172413 | 0.065865141 | Fishbein et al ( PMID28162975) |
| TCGA | TCGA-QT-A5XJ-01A | -0.629323495 | -0.499791303 | 0.129532193 | Fishbein et al ( PMID28162975) |
| TCGA | TCGA-QT-A5XK-01A | 0.20558272 | 0.205977876 | 0.000395156 | Fishbein et al ( PMID28162975) |
| TCGA | TCGA-QT-A5XL-01A | 0.358324193 | -0.112822621 | -0.471146813 | Fishbein et al ( PMID28162975) |
| TCGA | TCGA-QT-A5XM-01A | -0.356679708 | -0.268705955 | 0.087973753 | Fishbein et al ( PMID28162975) |
| TCGA | TCGA-QT-A5XN-01A | 0.059419247 | 0.395295534 | 0.335876287 | Fishbein et al ( PMID28162975) |
| TCGA | TCGA-QT-A5XO-01A | -0.027428292 | -0.177303282 | -0.149874991 | Fishbein et al ( PMID28162975) |
| TCGA | TCGA-QT-A5XP-01A | -0.052834184 | 0.159222982 | 0.212057165 | Fishbein et al ( PMID28162975) |
| TCGA | TCGA-QT-A69Q-01A | 0.009157781 | -0.33309434 | -0.342252121 | Fishbein et al ( PMID28162975) |
| TCGA | TCGA-QT-A7U0-01A | 0.216213565 | -0.153982403 | -0.370195969 | Fishbein et al ( PMID28162975) |
| TCGA | TCGA-RT-A6Y9-01A | -0.124335845 | 0.108222535 | 0.23255838 | Fishbein et al ( PMID28162975) |
| TCGA | TCGA-RT-A6YA-01A | 0.152455661 | 0.240315209 | 0.087859548 | Fishbein et al ( PMID28162975) |
| TCGA | TCGA-RT-A6YC-01A | -0.150668084 | 0.482199353 | 0.632867437 | Fishbein et al ( PMID28162975) |
| TCGA | TCGA-RW-A67V-01A | -0.270991549 | -0.349240592 | -0.078249043 | Fishbein et al ( PMID28162975) |
| TCGA | TCGA-RW-A67W-01A | 0.29784154 | 0.018547194 | -0.279294346 | Fishbein et al ( PMID28162975) |
| TCGA | TCGA-RW-A67X-01A | -0.138533917 | 0.081787151 | 0.220321068 | Fishbein et al ( PMID28162975) |
| TCGA | TCGA-RW-A67Y-01A | 0.324469925 | -0.28170388 | -0.606173805 | Fishbein et al ( PMID28162975) |
| TCGA | TCGA-RW-A680-01A | -0.172928657 | 0.0983149 | 0.271243557 | Fishbein et al ( PMID28162975) |
| TCGA | TCGA-RW-A681-01A | -0.326406327 | -0.36051866 | -0.034112334 | Fishbein et al ( PMID28162975) |
| TCGA | TCGA-RW-A684-01A | 0.445608325 | -0.108657853 | -0.554266178 | Fishbein et al ( PMID28162975) |
| TCGA | TCGA-RW-A685-01A | -0.397644493 | -0.163781473 | 0.23386302 | Fishbein et al ( PMID28162975) |
| TCGA | TCGA-RW-A686-01A | 0.02311883 | 0.189928739 | 0.166809909 | Fishbein et al ( PMID28162975) |
| TCGA | TCGA-QR-A6GZ-05A | 0.011636219 | 0.199110058 | 0.187473839 | Fishbein et al ( PMID28162975) |
| TCGA | TCGA-RW-A688-01A | -0.044569768 | -0.326599535 | -0.282029768 | Fishbein et al ( PMID28162975) |
| TCGA | TCGA-RW-A689-01A | -0.078448388 | 0.111647982 | 0.190096369 | Fishbein et al ( PMID28162975) |
| TCGA | TCGA-RW-A68A-01A | 0.417158585 | -0.219401854 | -0.636560438 | Fishbein et al ( PMID28162975) |
| TCGA | TCGA-RW-A68B-01A | 0.380182556 | -0.401432155 | -0.781614712 | Fishbein et al ( PMID28162975) |
| TCGA | TCGA-RW-A68C-01A | -0.012902012 | -0.065695299 | -0.052793287 | Fishbein et al ( PMID28162975) |
| TCGA | TCGA-RW-A68D-01A | -0.215914812 | -0.013570122 | 0.20234469 | Fishbein et al ( PMID28162975) |
| TCGA | TCGA-RW-A68F-01A | -0.141826115 | -0.103910187 | 0.037915929 | Fishbein et al ( PMID28162975) |
| TCGA | TCGA-RW-A68G-01A | 0.164264164 | 0.297087016 | 0.132822852 | Fishbein et al ( PMID28162975) |
| TCGA | TCGA-RW-A7CZ-01A | -0.416562636 | 0.235983456 | 0.652546092 | Fishbein et al ( PMID28162975) |
| TCGA | TCGA-RW-A7D0-01A | 0.471643277 | -0.161481743 | -0.633125019 | Fishbein et al ( PMID28162975) |
| TCGA | TCGA-RW-A8AZ-01A | -0.076059088 | 0.151658795 | 0.227717883 | Fishbein et al ( PMID28162975) |
| TCGA | TCGA-RX-A8JQ-01A | 0.507122445 | 0.397921862 | -0.109200583 | Fishbein et al ( PMID28162975) |
| TCGA | TCGA-S7-A7WL-01A | 0.038892551 | 0.275133455 | 0.236240904 | Fishbein et al ( PMID28162975) |
| TCGA | TCGA-S7-A7WM-01A | -0.317266619 | 0.423231241 | 0.74049786 | Fishbein et al ( PMID28162975) |
| TCGA | TCGA-S7-A7WN-01A | -0.041229612 | 0.082694739 | 0.123924351 | Fishbein et al ( PMID28162975) |
| TCGA | TCGA-S7-A7WO-01A | -0.511042794 | 0.257278148 | 0.768320942 | Fishbein et al ( PMID28162975) |
| TCGA | TCGA-S7-A7WP-01A | 0.114491941 | -0.203868083 | -0.318360024 | Fishbein et al ( PMID28162975) |
| TCGA | TCGA-S7-A7WQ-01A | -0.048266981 | 0.062426269 | 0.11069325 | Fishbein et al ( PMID28162975) |
| TCGA | TCGA-S7-A7WR-01A | -0.173566249 | 0.163243174 | 0.336809423 | Fishbein et al ( PMID28162975) |
| TCGA | TCGA-S7-A7WT-01A | 0.01736313 | 0.103875112 | 0.086511982 | Fishbein et al ( PMID28162975) |
| TCGA | TCGA-S7-A7WU-01A | -0.100792595 | 0.222990073 | 0.323782668 | Fishbein et al ( PMID28162975) |
| TCGA | TCGA-S7-A7WV-01A | -0.044270574 | 0.060410596 | 0.104681169 | Fishbein et al ( PMID28162975) |
| TCGA | TCGA-S7-A7WW-01A | -0.369600691 | 0.002864046 | 0.372464737 | Fishbein et al ( PMID28162975) |
| TCGA | TCGA-S7-A7WX-01A | 0.208412296 | 0.179738641 | -0.028673655 | Fishbein et al ( PMID28162975) |
| TCGA | TCGA-S7-A7X0-01A | 0.048386848 | 0.438345433 | 0.389958585 | Fishbein et al ( PMID28162975) |
| TCGA | TCGA-S7-A7X1-01A | 0.085325051 | -0.251164079 | -0.33648913 | Fishbein et al ( PMID28162975) |
| TCGA | TCGA-S7-A7X2-01A | 0.245745713 | 0.022711634 | -0.22303408 | Fishbein et al ( PMID28162975) |
| TCGA | TCGA-SA-A6C2-01A | -0.02332539 | 0.088012891 | 0.111338281 | Fishbein et al ( PMID28162975) |
| TCGA | TCGA-SP-A6QC-01A | -0.053986874 | 0.189506626 | 0.2434935 | Fishbein et al ( PMID28162975) |
| TCGA | TCGA-SP-A6QD-01A | -0.020638777 | 0.07709037 | 0.097729147 | Fishbein et al ( PMID28162975) |
| TCGA | TCGA-SP-A6QF-01A | -0.083135671 | 0.285434559 | 0.368570231 | Fishbein et al ( PMID28162975) |
| TCGA | TCGA-SP-A6QG-01A | -0.290948438 | -0.333604331 | -0.042655893 | Fishbein et al ( PMID28162975) |
| TCGA | TCGA-SP-A6QH-01A | -0.147679291 | 0.134316182 | 0.281995473 | Fishbein et al ( PMID28162975) |
| TCGA | TCGA-SP-A6QI-01A | -0.205751064 | 0.032258611 | 0.238009675 | Fishbein et al ( PMID28162975) |
| TCGA | TCGA-SP-A6QJ-01A | 0.163362664 | 0.205519716 | 0.042157052 | Fishbein et al ( PMID28162975) |
| TCGA | TCGA-SP-A6QK-01A | 0.211687069 | 0.362118418 | 0.150431349 | Fishbein et al ( PMID28162975) |
| TCGA | TCGA-SQ-A6I4-01A | 0.023872874 | 0.120920824 | 0.09704795 | Fishbein et al ( PMID28162975) |
| TCGA | TCGA-SQ-A6I6-01A | 0.30748948 | 0.149929325 | -0.157560155 | Fishbein et al ( PMID28162975) |
| TCGA | TCGA-SR-A6MP-01A | -0.021668955 | 0.006452786 | 0.028121741 | Fishbein et al ( PMID28162975) |
| TCGA | TCGA-SR-A6MQ-01A | 0.274406026 | 0.08911775 | -0.185288275 | Fishbein et al ( PMID28162975) |
| TCGA | TCGA-SR-A6MR-01A | -0.359108892 | 0.263110096 | 0.622218988 | Fishbein et al ( PMID28162975) |
| TCGA | TCGA-SR-A6MS-01A | -0.172175774 | -0.356612066 | -0.184436293 | Fishbein et al ( PMID28162975) |
| TCGA | TCGA-SR-A6MT-01A | -0.246698764 | -0.311382388 | -0.064683624 | Fishbein et al ( PMID28162975) |
| TCGA | TCGA-SR-A6MU-01A | -0.176942188 | -0.080635227 | 0.096306962 | Fishbein et al ( PMID28162975) |
| TCGA | TCGA-SR-A6MV-01A | 0.398753962 | 0.242403684 | -0.156350278 | Fishbein et al ( PMID28162975) |
| TCGA | TCGA-SR-A6MX-01A | -0.340003707 | -0.049609532 | 0.290394175 | Fishbein et al ( PMID28162975) |
| TCGA | TCGA-RW-A686-06A | -0.023998639 | -0.077248491 | -0.053249852 | Fishbein et al ( PMID28162975) |
| TCGA | TCGA-SR-A6MX-05A | -0.291578334 | -0.15096127 | 0.140617064 | Fishbein et al ( PMID28162975) |
| TCGA | TCGA-SR-A6MY-01A | 0.479627657 | 0.104209938 | -0.375417719 | Fishbein et al ( PMID28162975) |
| TCGA | TCGA-SR-A6MZ-01A | 0.227721258 | 0.139221211 | -0.088500048 | Fishbein et al ( PMID28162975) |
| TCGA | TCGA-SR-A6N0-01A | -0.478053566 | 0.296051221 | 0.774104787 | Fishbein et al ( PMID28162975) |
| TCGA | TCGA-TT-A6YJ-01A | -0.016840721 | -0.073842806 | -0.057002086 | Fishbein et al ( PMID28162975) |
| TCGA | TCGA-TT-A6YK-01A | 0.269568542 | 0.351096392 | 0.08152785 | Fishbein et al ( PMID28162975) |
| TCGA | TCGA-TT-A6YN-01A | 0.177260265 | -0.057669616 | -0.234929882 | Fishbein et al ( PMID28162975) |
| TCGA | TCGA-TT-A6YO-01A | 0.018672216 | -0.001692331 | -0.020364547 | Fishbein et al ( PMID28162975) |
| TCGA | TCGA-TT-A6YP-01A | 0.030723339 | -0.243805647 | -0.274528986 | Fishbein et al ( PMID28162975) |
| TCGA | TCGA-W2-A7H5-01B | 0.206612375 | -0.400357275 | -0.606969651 | Fishbein et al ( PMID28162975) |
| TCGA | TCGA-W2-A7H7-01A | -0.006222388 | -0.234295523 | -0.228073134 | Fishbein et al ( PMID28162975) |
| TCGA | TCGA-W2-A7HA-01B | 0.427318036 | 0.019954964 | -0.407363073 | Fishbein et al ( PMID28162975) |
| TCGA | TCGA-W2-A7HB-01A | -0.122545231 | -0.324110821 | -0.20156559 | Fishbein et al ( PMID28162975) |
| TCGA | TCGA-W2-A7HC-01A | -0.116730387 | 0.025925593 | 0.14265598 | Fishbein et al ( PMID28162975) |
| TCGA | TCGA-W2-A7HD-01A | 0.427335839 | -0.136448188 | -0.563784027 | Fishbein et al ( PMID28162975) |
| TCGA | TCGA-W2-A7HE-01A | -0.206416707 | -0.109405592 | 0.097011115 | Fishbein et al ( PMID28162975) |
| TCGA | TCGA-W2-A7HF-01A | -0.029762036 | -0.301465458 | -0.271703422 | Fishbein et al ( PMID28162975) |
| TCGA | TCGA-W2-A7HH-01A | 0.231222863 | -0.266421191 | -0.497644054 | Fishbein et al ( PMID28162975) |
| TCGA | TCGA-SR-A6MX-06A | -0.284746186 | -0.21462316 | 0.070123026 | Fishbein et al ( PMID28162975) |
| TCGA | TCGA-WB-A80K-01A | -0.315001268 | -0.24230185 | 0.072699417 | Fishbein et al ( PMID28162975) |
| TCGA | TCGA-WB-A80L-01A | 0.162000974 | -0.016145406 | -0.178146379 | Fishbein et al ( PMID28162975) |
| TCGA | TCGA-WB-A80M-01A | 0.474104095 | -0.36042052 | -0.834524615 | Fishbein et al ( PMID28162975) |
| TCGA | TCGA-WB-A80N-01A | -0.019946687 | 0.085990862 | 0.105937549 | Fishbein et al ( PMID28162975) |
| TCGA | TCGA-WB-A80O-01A | 0.25314121 | 0.35437307 | 0.10123186 | Fishbein et al ( PMID28162975) |
| TCGA | TCGA-WB-A80P-01A | -0.134788847 | -0.00440316 | 0.130385687 | Fishbein et al ( PMID28162975) |
| TCGA | TCGA-WB-A80Q-01A | 0.037203629 | 0.093400438 | 0.056196809 | Fishbein et al ( PMID28162975) |
| TCGA | TCGA-WB-A80V-01A | 0.141722313 | -0.061261972 | -0.202984286 | Fishbein et al ( PMID28162975) |
| TCGA | TCGA-WB-A80Y-01A | -0.010997042 | -0.105437909 | -0.094440867 | Fishbein et al ( PMID28162975) |
| TCGA | TCGA-WB-A814-01A | 0.28533935 | 0.09993601 | -0.18540334 | Fishbein et al ( PMID28162975) |
| TCGA | TCGA-WB-A815-01A | 0.032921486 | 0.285280661 | 0.252359175 | Fishbein et al ( PMID28162975) |
| TCGA | TCGA-WB-A816-01A | -0.087074112 | -0.002869261 | 0.084204851 | Fishbein et al ( PMID28162975) |
| TCGA | TCGA-WB-A817-01A | -0.358602159 | -0.1752167 | 0.183385459 | Fishbein et al ( PMID28162975) |
| TCGA | TCGA-WB-A818-01A | 0.212342017 | -0.082440896 | -0.294782912 | Fishbein et al ( PMID28162975) |
| TCGA | TCGA-WB-A819-01A | -0.28315447 | -0.122244953 | 0.160909517 | Fishbein et al ( PMID28162975) |
| TCGA | TCGA-WB-A81A-01A | -0.32559079 | -0.383665356 | -0.058074566 | Fishbein et al ( PMID28162975) |
| TCGA | TCGA-WB-A81D-01A | 0.074948349 | -0.139530803 | -0.214479152 | Fishbein et al ( PMID28162975) |
| TCGA | TCGA-WB-A81E-01A | -0.545464021 | -0.28346916 | 0.261994861 | Fishbein et al ( PMID28162975) |
| TCGA | TCGA-WB-A81F-01A | 0.233455553 | -0.374489512 | -0.607945065 | Fishbein et al ( PMID28162975) |
| TCGA | TCGA-WB-A81G-01A | -0.454735948 | 0.180800053 | 0.635536001 | Fishbein et al ( PMID28162975) |
| TCGA | TCGA-WB-A81H-01A | 0.030415043 | 0.433157634 | 0.402742591 | Fishbein et al ( PMID28162975) |
| TCGA | TCGA-WB-A81I-01A | 0.169241874 | -0.064387217 | -0.233629091 | Fishbein et al ( PMID28162975) |
| TCGA | TCGA-WB-A81J-01A | -0.105759154 | -0.24945714 | -0.143697986 | Fishbein et al ( PMID28162975) |
| TCGA | TCGA-WB-A81K-01A | 0.332522046 | -0.194694001 | -0.527216047 | Fishbein et al ( PMID28162975) |
| TCGA | TCGA-WB-A81M-01A | 0.269345121 | -0.349478343 | -0.618823464 | Fishbein et al ( PMID28162975) |
| TCGA | TCGA-WB-A81N-01A | 0.402890279 | -0.186198964 | -0.589089243 | Fishbein et al ( PMID28162975) |
| TCGA | TCGA-WB-A81P-01A | -0.17359755 | 0.252906318 | 0.426503868 | Fishbein et al ( PMID28162975) |
| TCGA | TCGA-WB-A81Q-01A | 0.570431508 | -0.179494503 | -0.749926011 | Fishbein et al ( PMID28162975) |
| TCGA | TCGA-WB-A81R-01A | -0.225588387 | -0.082260678 | 0.143327709 | Fishbein et al ( PMID28162975) |
| TCGA | TCGA-WB-A81S-01A | 0.334897199 | 0.384821472 | 0.049924273 | Fishbein et al ( PMID28162975) |
| TCGA | TCGA-WB-A81T-01A | 0.038964277 | -0.00697191 | -0.045936187 | Fishbein et al ( PMID28162975) |
| TCGA | TCGA-WB-A81V-01A | -0.180287488 | 0.081883753 | 0.262171241 | Fishbein et al ( PMID28162975) |
| TCGA | TCGA-WB-A81W-01A | 0.063995547 | 0.203155724 | 0.139160177 | Fishbein et al ( PMID28162975) |
| TCGA | TCGA-WB-A821-01A | 0.189783841 | -0.163985082 | -0.353768923 | Fishbein et al ( PMID28162975) |
| TCGA | TCGA-WB-A822-01A | -0.205243966 | 0.432650734 | 0.6378947 | Fishbein et al ( PMID28162975) |
| TCGA | TCGA-XG-A823-01A | -0.275237581 | -0.121510322 | 0.153727258 | Fishbein et al ( PMID28162975) |
| TCGA | TCGA-W2-A7UY-01A | 0.098332465 | -0.200070688 | -0.298403153 | Fishbein et al ( PMID28162975) |
| sc/snRNA UT | 802 | -0.209547525 | 0.129951112 | 0.339498638 | This study |
| sc/snRNA UT | 1119 | -0.001436776 | -0.509217674 | -0.507780898 | This study |
| sc/snRNA UT | 1182 | 0.457429809 | -0.321157967 | -0.778587776 | This study |
| sc/snRNA UT | 1244 | -0.280876858 | -0.206933294 | 0.073943565 | This study |
| sc/snRNA UT | 1259 | -0.23041154 | -0.181987771 | 0.048423769 | This study |
| sc/snRNA UT | 1312 | -0.341329155 | 0.08735574 | 0.428684895 | This study |
| sc/snRNA UT | 1313 | -0.097589395 | 0.061379113 | 0.158968507 | This study |
| sc/snRNA UT | 1314 | -0.67800293 | -0.308308436 | 0.369694494 | This study |
| sc/snRNA UT | 1322 | 0.511418811 | -0.320345762 | -0.831764574 | This study |
| sc/snRNA UT | 1322 | -0.1911704 | -0.247522898 | -0.056352498 | This study |
| sc/snRNA UT | 1343 | 0.227608502 | 0.339574755 | 0.111966253 | This study |
| sc/snRNA UT | 1415 | -0.201132606 | -0.182194911 | 0.018937695 | This study |
| sc/snRNA UT | 1415 | -0.044789969 | -0.450600406 | -0.405810437 | This study |
| sc/snRNA UT | 1415 | -0.202365731 | -0.229736072 | -0.027370341 | This study |
| sc/snRNA UT | 393 | 0.473214034 | -0.003343561 | -0.476557596 | This study |
| sc/snRNA UT | 1195 | -0.141392139 | -0.395163226 | -0.253771087 | This study |
| sc/snRNA UT | 1196 | -0.120354495 | -0.196625196 | -0.076270701 | This study |
| UT bulk | 1241 | -0.281394916 | -0.021319835 | 0.260075082 | This study |
| UT bulk | 1234 | -0.215699204 | -0.415351258 | -0.199652054 | This study |
| UT bulk | 1236 | -0.036667794 | 0.214963208 | 0.251631002 | This study |
| UT bulk | 1262 | -0.568375475 | 0.151058936 | 0.719434411 | This study |
| UT bulk | 1263 | -0.199563786 | 0.190113257 | 0.389677044 | This study |
| UT bulk | 1301 | -0.168526249 | 0.033658818 | 0.202185066 | This study |
| UT bulk | 1306 | -0.101122029 | -0.09511261 | 0.006009419 | This study |
| UT bulk | 1311 | -0.265809681 | -0.145869975 | 0.119939706 | This study |
| UT bulk | 1289 | -0.24590046 | -0.326089395 | -0.080188935 | This study |
| UT bulk | 1325 | 0.038626193 | -0.373343412 | -0.411969605 | This study |
| UT bulk | 1295 | 0.017966918 | 0.175994487 | 0.158027569 | This study |
| UT bulk | 1328 | -0.4881004 | 0.370035113 | 0.858135513 | This study |
| UT bulk | 1317 | 0.079067781 | -0.110755804 | -0.189823585 | This study |
| UT bulk | 1327 | 0.37438468 | 0.215191572 | -0.159193108 | This study |
| UT bulk | 1359 | -0.084398092 | 0.276850047 | 0.361248139 | This study |
| UT bulk | 1361 | 0.251092276 | -0.149875152 | -0.400967428 | This study |
| UT bulk | 1415 | 0.517390689 | 0.211875072 | -0.305515617 | This study |
| UT bulk | 1373 | 0.393502623 | -0.163911661 | -0.557414284 | This study |
| UT bulk | 1405 | 0.259891025 | 0.109716422 | -0.150174603 | This study |
| UT bulk | 1098 | -0.511518427 | 0.315356013 | 0.82687444 | This study |
| UT bulk | 1100 | -0.215150279 | 0.224762164 | 0.439912444 | This study |
| UT bulk | 1104 | 0.415617779 | 0.066335537 | -0.349282241 | This study |
| UT bulk | 1108 | -0.249749277 | -0.507889518 | -0.25814024 | This study |
| UT bulk | 1111 | 0.52250385 | -0.279452806 | -0.801956656 | This study |
| UT bulk | 1119 | -0.059707325 | -0.012563186 | 0.047144138 | This study |
| UT bulk | 1125 | 0.359585112 | -0.102617143 | -0.462202255 | This study |
| UT bulk | 1130 | 0.023813863 | -0.2332201 | -0.257033963 | This study |
| UT bulk | 1143 | 0.246293345 | 0.392599891 | 0.146306547 | This study |
| UT bulk | 1145 | -0.51047378 | 0.330362092 | 0.840835872 | This study |
| UT bulk | 1147 | -0.141154961 | 0.159893677 | 0.301048638 | This study |
| UT bulk | 1157 | 0.423866971 | 0.00992455 | -0.413942421 | This study |
| UT bulk | 1164 | -0.121305614 | 0.348560975 | 0.469866589 | This study |
| UT bulk | 1169 | -0.007491 | -0.205137036 | -0.197646037 | This study |
| UT bulk | 1183 | 0.048874774 | -0.302004007 | -0.350878781 | This study |
| UT bulk | 1190 | 0.283846745 | -0.248212583 | -0.532059328 | This study |
| UT bulk | 1194 | -0.428544393 | -0.032690269 | 0.395854124 | This study |
| UT bulk | 1195 | 0.021679234 | -0.255752141 | -0.277431375 | This study |
| UT bulk | 1204 | 0.033103848 | -0.0068378 | -0.039941647 | This study |
| UT bulk | 1212 | 0.136875153 | -0.047421943 | -0.184297097 | This study |
| UT bulk | 1219 | -0.361466503 | 0.128012132 | 0.489478635 | This study |
| UT bulk | 1226 | -0.151920784 | -0.18699409 | -0.035073306 | This study |
| UT bulk | 1228 | -0.006642388 | 0.156411635 | 0.163054023 | This study |
| UT bulk | 1231 | -0.339255404 | -0.097000022 | 0.242255382 | This study |
| UT bulk | 128 | -0.120509765 | 0.188079448 | 0.308589212 | This study |
| UT bulk | 166 | 0.106343875 | 0.37430797 | 0.267964095 | This study |
| UT bulk | 493 | 0.259954355 | -0.332840771 | -0.592795126 | This study |
| UT bulk | 563 | 0.131297206 | 0.227796408 | 0.096499201 | This study |
| snRNA Zethoven | E240 | -0.129772248 | -0.098882908 | 0.030889339 | Zethoven et al (PMID30946460) |
| snRNA Zethoven | E243 | -0.072629605 | -0.200679104 | -0.128049499 | Zethoven et al (PMID30946460) |
| snRNA Zethoven | E007 | 0.425210766 | 0.227658047 | -0.197552718 | Zethoven et al (PMID30946460) |
| snRNA Zethoven | E008 | 0.209246105 | 0.204706224 | -0.004539881 | Zethoven et al (PMID30946460) |
| snRNA Zethoven | E017 | -0.389709976 | -0.201657832 | 0.188052144 | Zethoven et al (PMID30946460) |
| snRNA Zethoven | E018 | 0.452208682 | 0.03393066 | -0.418278022 | Zethoven et al (PMID30946460) |
| snRNA Zethoven | E019 | -0.153061065 | -0.280617664 | -0.127556599 | Zethoven et al (PMID30946460) |
| snRNA Zethoven | E024 | 0.051916351 | -0.384306433 | -0.436222784 | Zethoven et al (PMID30946460) |
| snRNA Zethoven | E025 | -0.238966764 | -0.262657242 | -0.023690477 | Zethoven et al (PMID30946460) |
| snRNA Zethoven | E027 | 0.28749481 | 0.041905442 | -0.245589368 | Zethoven et al (PMID30946460) |
| snRNA Zethoven | E033 | -0.289666393 | -0.310945178 | -0.021278785 | Zethoven et al (PMID30946460) |
| snRNA Zethoven | E035 | 0.013033417 | -0.085151535 | -0.098184952 | Zethoven et al (PMID30946460) |
| snRNA Zethoven | E041 | -0.277152485 | 0.226495114 | 0.503647599 | Zethoven et al (PMID30946460) |
| snRNA Zethoven | E042 | 0.149556069 | -0.076064345 | -0.225620414 | Zethoven et al (PMID30946460) |
| snRNA Zethoven | E174 | 0.028575459 | 0.294435773 | 0.265860314 | Zethoven et al (PMID30946460) |
| snRNA Zethoven | E196 | -0.548553717 | -0.412910988 | 0.135642729 | Zethoven et al (PMID30946460) |
| snRNA Zethoven | E205 | 0.522213629 | -0.217724004 | -0.739937633 | Zethoven et al (PMID30946460) |
| snRNA Zethoven | E206 | 0.209310216 | 0.316507414 | 0.107197198 | Zethoven et al (PMID30946460) |
| snRNA Zethoven | E208 | -0.533444505 | -0.53342865 | 1.59E-05 | Zethoven et al (PMID30946460) |
| snRNA Zethoven | E209 | -0.005010364 | -0.349602483 | -0.344592119 | Zethoven et al (PMID30946460) |
| snRNA Zethoven | E210 | 0.3014667 | 0.380983723 | 0.079517023 | Zethoven et al (PMID30946460) |
| snRNA Zethoven | E211 | -0.566626766 | -0.601243897 | -0.034617131 | Zethoven et al (PMID30946460) |
| snRNA Zethoven | E213 | -0.584857928 | -0.019880162 | 0.564977766 | Zethoven et al (PMID30946460) |
| snRNA Zethoven | E214 | -0.585778889 | 0.331571761 | 0.91735065 | Zethoven et al (PMID30946460) |
| snRNA Zethoven | E216 | -0.526186323 | 0.324079137 | 0.85026546 | Zethoven et al (PMID30946460) |
| snRNA Zethoven | E235 | 0.220985531 | 0.098801516 | -0.122184015 | Zethoven et al (PMID30946460) |
| snRNA Zethoven | E237 | 0.187270795 | 0.118395322 | -0.068875473 | Zethoven et al (PMID30946460) |
| snRNA Zethoven | E215 | -0.235194596 | -0.092417715 | 0.142776882 | Zethoven et al (PMID30946460) |
| snRNA Zethoven | E326 | -0.097995126 | -0.586562822 | -0.488567696 | Zethoven et al (PMID30946460) |
| snRNA Zethoven | P010 | 0.221653719 | -0.100512791 | -0.32216651 | Zethoven et al (PMID30946460) |
| snRNA Zethoven | P018.PGL1 | -0.147537361 | -0.126439403 | 0.021097958 | Zethoven et al (PMID30946460) |
| snRNA Zethoven | P018.PGL3 | 0.03882159 | -0.111359679 | -0.150181269 | Zethoven et al (PMID30946460) |

**Supplementary Table 16.** NOR and MES gsva scores and differential MES-NOR score in primary pheochromocytomas and paragangliomas calculated from tumor matrix expression data (bulk RNAseq) or tumor cell expression expression data (single nuclei data) in the designated cohorts (see Methods for details)

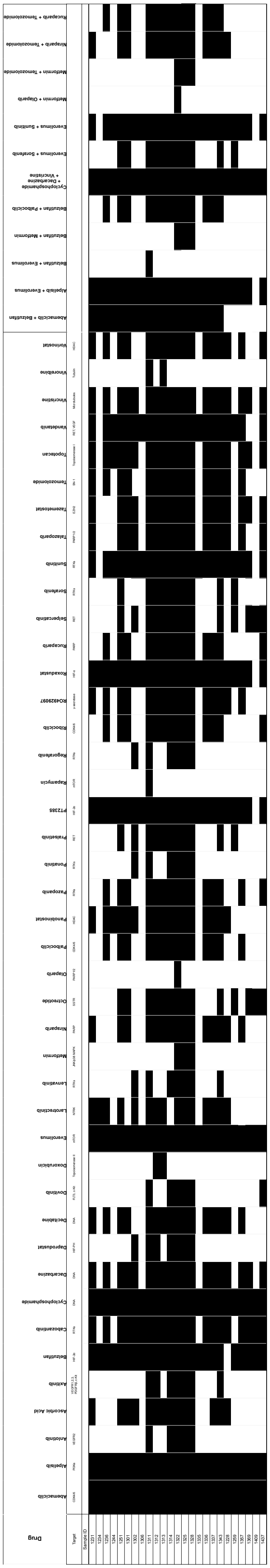

**Supplementary Table 17.** Drugs and drug combinations used for PPGL organoid screen.

| Sample ID | PDO condition | Drugs.Tested | Drugs inducing 25 % Cell Death | Drugs inducing 50 % Cell Death | Drugs inducing 75 % Cell Death |
|---|---|---|---|---|---|
| 1228 | N | 26 | 19.2% (5/26) | 3.8% (1/26) | 3.8% (1/26) |
| 1231 | L | 13 | 30.8% (4/13) | 0% (0/13) | 0% (0/13) |
| 1231 | N | 26 | 15.4% (4/26) | 3.8% (1/26) | 0% (0/26) |
| 1234 | N | 10 | 10% (1/10) | 0% (0/10) | 0% (0/10) |
| 1236 | N | 28 | 32.1% (9/28) | 7.1% (2/28) | 3.6% (1/28) |
| 1244 | L | 13 | 7.7% (1/13) | 0% (0/13) | 0% (0/13) |
| 1244 | N | 13 | 7.7% (1/13) | 0% (0/13) | 0% (0/13) |
| 1251 | L | 38 | 44.7% (17/38) | 0% (0/38) | 0% (0/38) |
| 1251 | N | 39 | 2.6% (1/39) | 0% (0/39) | 0% (0/39) |
| 1259 | N | 16 | 50% (8/16) | 6.2% (1/16) | 0% (0/16) |
| 1301 | N | 35 | 14.3% (5/35) | 0% (0/35) | 0% (0/35) |
| 1302 | N | 28 | 7.1% (2/28) | 0% (0/28) | 0% (0/28) |
| 1306 | L | 13 | 53.8% (7/13) | 23.1% (3/13) | 0% (0/13) |
| 1306 | N | 13 | 38.5% (5/13) | 7.7% (1/13) | 0% (0/13) |
| 1311 | N | 50 | 12% (6/50) | 0% (0/50) | 0% (0/50) |
| 1311 | H | 50 | 12% (6/50) | 0% (0/50) | 0% (0/50) |
| 1312 | L | 39 | 61.5% (24/39) | 43.6% (17/39) | 17.9% (7/39) |
| 1312 | L | 24 | 20.8% (5/24) | 8.3% (2/24) | 0% (0/24) |
| 1312 | N | 39 | 2.6% (1/39) | 0% (0/39) | 0% (0/39) |
| 1312 | H | 39 | 28.2% (11/39) | 0% (0/39) | 0% (0/39) |
| 1312 | H | 26 | 0% (0/26) | 0% (0/26) | 0% (0/26) |
| 1313 | L | 40 | 27.5% (11/40) | 5% (2/40) | 0% (0/40) |
| 1313 | L | 14 | 21.4% (3/14) | 0% (0/14) | 0% (0/14) |
| 1313 | N | 40 | 7.5% (3/40) | 0% (0/40) | 0% (0/40) |
| 1313 | H | 40 | 12.5% (5/40) | 0% (0/40) | 0% (0/40) |
| 1314 | N | 44 | 29.5% (13/44) | 4.5% (2/44) | 0% (0/44) |
| 1314 | H | 45 | 17.8% (8/45) | 8.9% (4/45) | 4.4% (2/45) |
| 1322 | L | 50 | 64% (32/50) | 46% (23/50) | 18% (9/50) |
| 1322 | N | 51 | 49% (25/51) | 3.9% (2/51) | 0% (0/51) |
| 1322 | H | 51 | 23.5% (12/51) | 0% (0/51) | 0% (0/51) |
| 1328 | N | 49 | 53.1% (26/49) | 22.4% (11/49) | 6.1% (3/49) |
| 1335 | N | 12 | 8.3% (1/12) | 0% (0/12) | 0% (0/12) |
| 1336 | N | 33 | 54.5% (18/33) | 0% (0/33) | 0% (0/33) |
| 1337 | L | 34 | 70.6% (24/34) | 29.4% (10/34) | 2.9% (1/34) |
| 1337 | N | 34 | 5.9% (2/34) | 0% (0/34) | 0% (0/34) |
| 1343 | N | 41 | 0% (0/41) | 0% (0/41) | 0% (0/41) |
| 1343 | H | 41 | 24.4% (10/41) | 4.9% (2/41) | 0% (0/41) |
| 1357 | N | 25 | 32% (8/25) | 12% (3/25) | 0% (0/25) |
| 1369 | H | 18 | 11.1% (2/18) | 0% (0/18) | 0% (0/18) |
| 1369 | N | 18 | 44.4% (8/18) | 5.6% (1/18) | 0% (0/18) |
| 1409 | H | 8 | 0% (0/8) | 0% (0/8) | 0% (0/8) |
| 1409 | N | 8 | 12.5% (1/8) | 0% (0/8) | 0% (0/8) |
| 1437 | H | 24 | 4.2% (1/24) | 0% (0/24) | 0% (0/24) |
| 1437 | N | 25 | 4% (1/25) | 0% (0/25) | 0% (0/25) |
| | | 30.07 | | | |

**Supplementary Table 18.** Response to drug screen based on viability assay by individual organoid sample; N=normoxia, short term; H=hypoxia, short term; L=normoxia, long term

**Supplementary Table 19.** Differentially expressed genes between top Abemaciclib responders vs non responders. UP: (log2FC > 2 & Pvalue < 0.05) upregulated genes in responders, DOWN: (log2FC < 2 & Pvalue < 0.05) downregulated genes in responders, NO: not differentially expressed.

Table provided in synapse.org/PPGLorganoids.

**Supplementary Table 20.** GSEA of Abemaciclib responders vs non responders

Table provided in synapse.org/PPGLorganoids.

#### Supplementary Figures

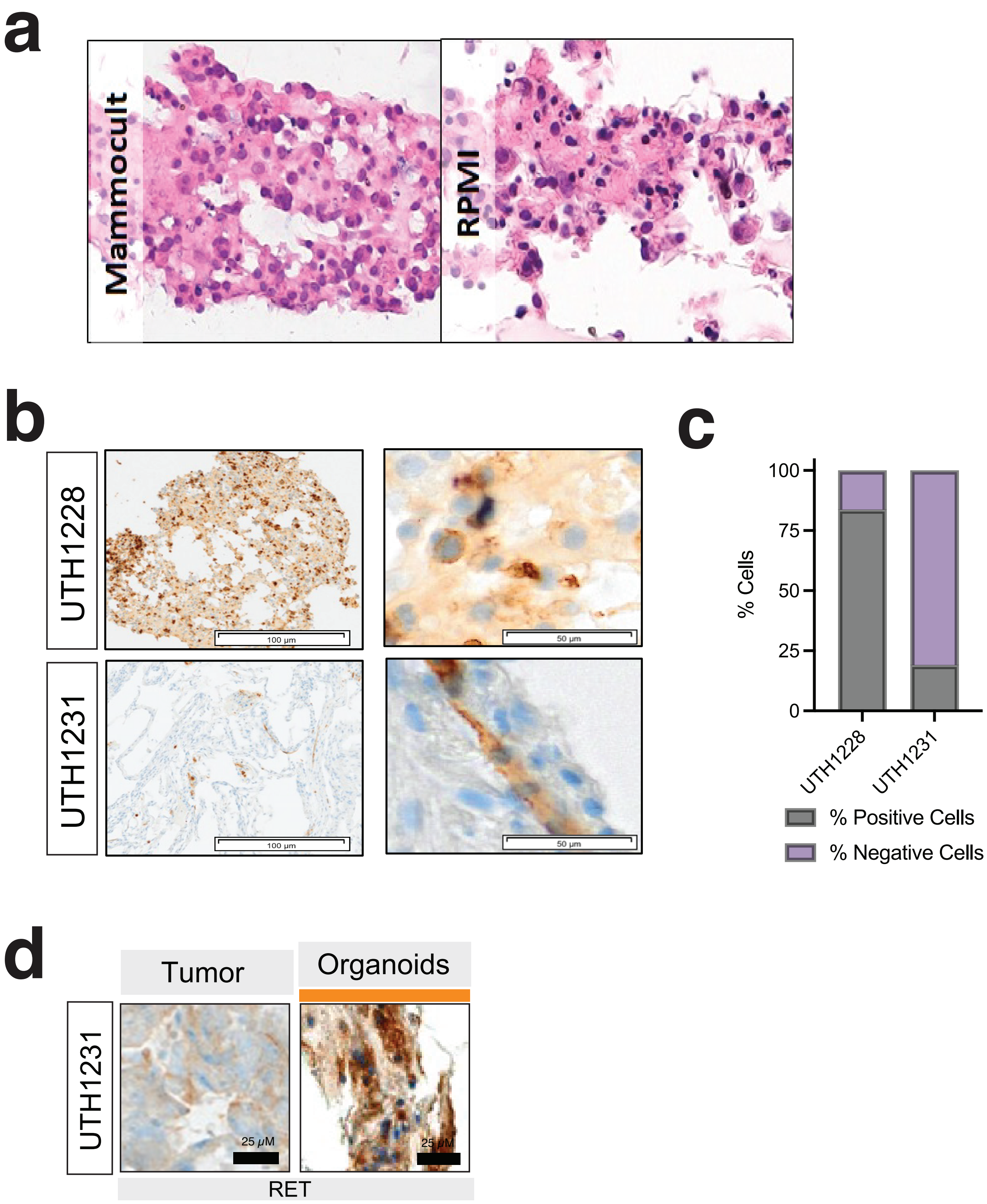

**Supplementary Figure 1.** a) H&E staining of one PDO cultured under two different culture media conditions, Mammocult (left) and RPMI (right); b) Immunohistochemistry (IHC) staining of endothelial marker CD34 in two PDOs at low (right) and high (left) magnification; c) the quantification of positive CD34 cells in the entire PDO section (the percentage of positive and negative cells is displayed); d) IHC of RET of the primary tumor and respective long term PDO (UTH1231), scale bar is as indicated (Related to Fig 2c).

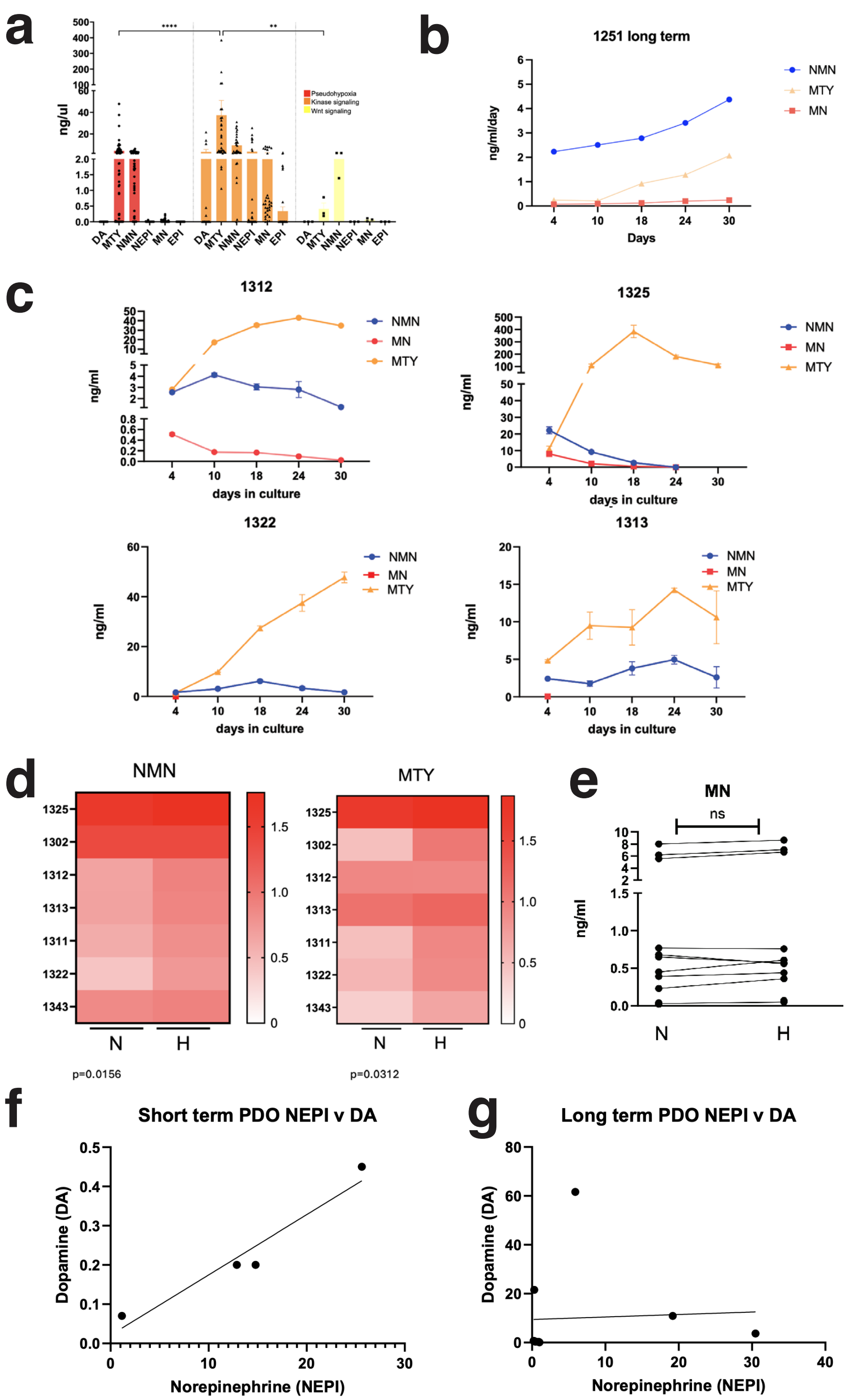

| Pearson r | |
|---|---|
| r | 0.9671 |
| 95% confidence interval | 0.08576 to 0.9993 |
| R squared | 0.9353 |
| P value | |
| P (two-tailed) | 0.0329 |

| Pearson r | |
|---|---|
| r | 0.05492 |
| 95% confidence interval | -0.5953 to 0.6617 |
| R squared | 0.003016 |
| P value | |
| P (two-tailed) | 0.8802 |

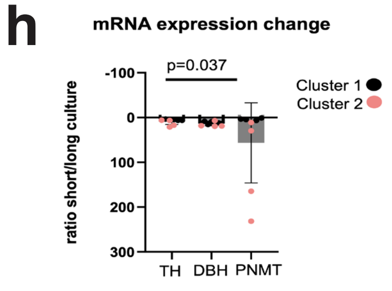

**Supplementary Figure 2.** a) Aggregate values of NEPI, EPI, DA, NMN, MN and MTY levels in supernatant in samples split by the tumor molecular cluster; b) Longitudinal profile of NMN, MN and MTY supernatant levels of a long term PDO culture derived from an EPAS1-derived pheochromocytoma that shows detectable NMN and trace amounts of MN; c) Longitudinal levels of NMN, MN and MTY measured in four distinct long term PDO samples;) d) Paired normoxia (N) and hypoxia (H) supernatant levels of NMN (left) and MTY (right), values were log transformed, p values NMN p=0.0156, and MTY p=0.0312, two-tailed p value, Wilcoxon matched-pairs signed rank test; e) Supernatant levels of MN in paired N and H; f) Correlation between NEPI and DA levels in short-term cultures r, CI, r2, p value are shown; g) Correlation between NEPI and DA levels in long-term cultures, r, CI, r2, p value are shown; h) Ratio of mRNA expression levels of TH, DBH and PNMT, encoding for enzymes of the biosynthetic cascade of catecholamines collected at day 6 or day 30 of culture, ratio of long/short term mRNA levels are shown.

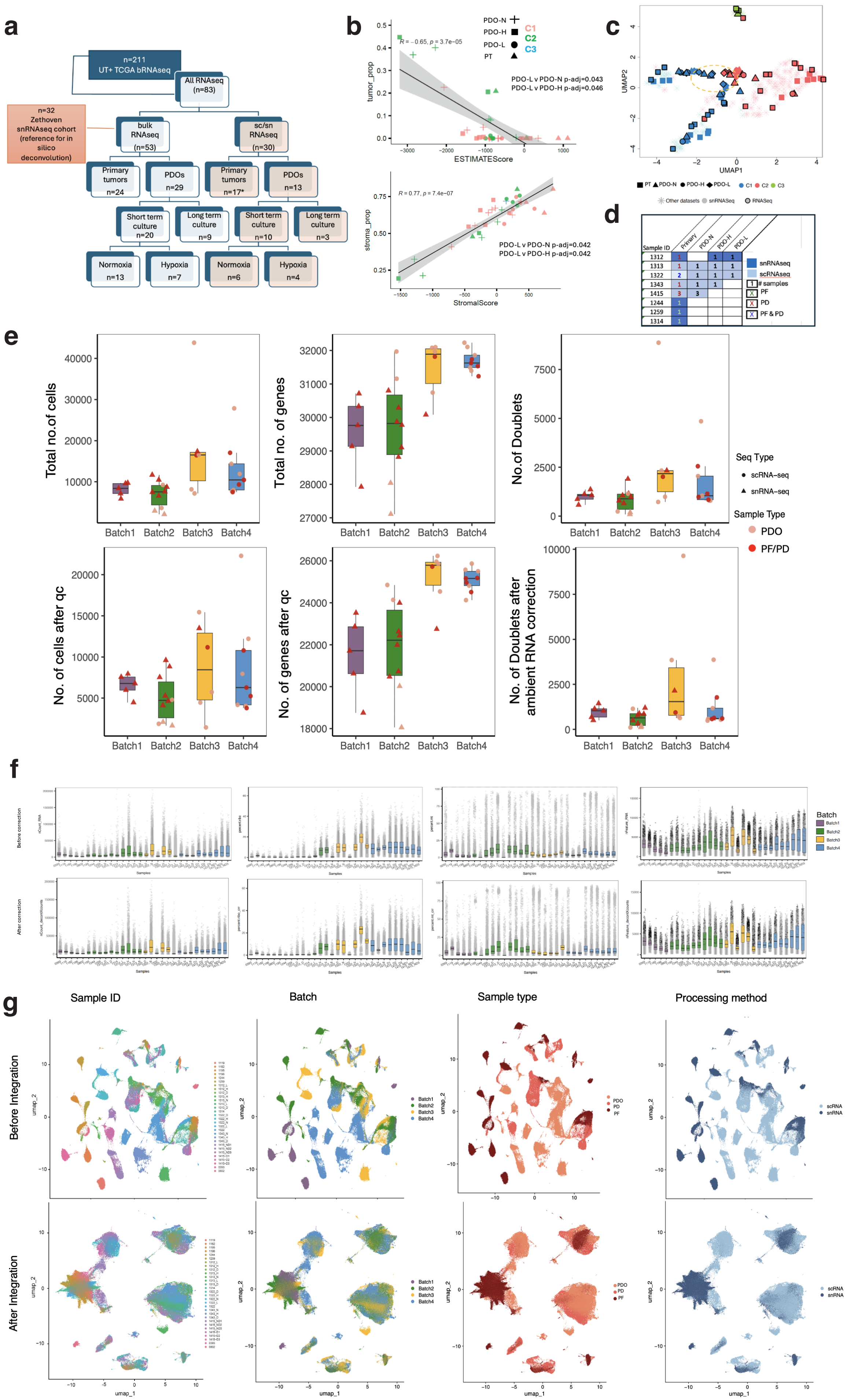
a
n=211
UT+ TCGA bRNAseq
All RNAseq
(n=83)
n=32
Zethoven
snRNAseq cohort
(reference for in
silico
deconvolution)
bulk
RNAseq
(n=53)
sc/sn
RNAseq
(n=30)
Primary
tumors
n=24
PDOs
n=29
Primary
tumors
n=17*
PDOs
n=13
Short term
culture
n=20
Long term
culture
n=9
Short term
culture
n=10
Long term
culture
n=3
Normoxia
n=13
Hypoxia
n=7
Normoxia
n=6
Hypoxia
n=4
b
PDO-N
PDO-H
PDO-L
PT
C1
C2
C3
tumor_prop
ESTIMATEScore
PDO-L v PDO-N p-adj=0.043
PDO-L v PDO-H p-adj=0.046
stroma_prop
StromalScore
PDO-L v PDO-N p-adj=0.042
PDO-L v PDO-H p-adj=0.042
c
UMAP1
UMAP2
d
Sample ID
Primary
PDO-N
PDO-H
PDO-L
1312
1313
1322
1343
1415
1244
1259
1314
snRNAseq
scRNAseq
### samples
PF
PD
PF & PD
e
Total no.of cells
Total no. of genes
No.of Doublets
No. of cells after qc
No. of genes after qc
No. of Doublets after ambient RNA correction
Batch1
Batch2
Batch3
Batch4
Seq Type
scRNA−seq
snRNA−seq
Sample Type
PDO
PF/PD
f
Before correction
After correction
Samples
Batch
Batch1
Batch2
Batch3
Batch4
g
Sample ID
Batch
Sample type
Processing method
Before Integration
After Integration
umap_1
umap_2
PDO
PD
PF
scRNA
snRNA

**Supplementary Figure 3.** a) Number of samples undergoing RNAseq analysis, including bulk and single cell/single nuclei (sc/snRNA-Seq) as well as external data added for the analyses, some samples were analyzed by both sc/sn and bulk RNAseq, *samples added to increase cell clustering fidelity; b) Top: estimated score of tumor cells (x-axis: ESTIMATE) and proportion of tumor cells (y-axis: MUSiC-Multi-subject Single-cell Deconvolution) from bulk RNAseq of PDO-N, PDO-H, PDO-L and respective primary tumors (PT), indicated by symbol and molecular cluster colors as indicated, correlation score and adjusted p-value are shown, bottom: estimated stromal cell score (x-axis: ESTIMATE) and proportion of stromal cells (y-axis: MUSiC) from bulk RNAseq of PDO-N, PDO-H, PDO-L and respective primary tumors (PT), depicted by symbol and molecular cluster colors as indicated, correlation score and adjusted p-value are shown; c) UMAP of PDOs and PTs (83 samples shown in Fig 4a) based on deconvoluted or tumor cell only data (sample types include PDO-N,H,L and primary tumors-PT, molecular classification into pseudohypoxic (C1), kinase (C2), Wnt signaling (C3), type of sample process bRNAseq-bulk (thick outline), sn/scRNaseq: single nuclei/single cell RNAseq); d) condition, number and processing mode of PDOs and parental tumors by sc/snRNAseq, related to Fig 4e; e) Plots showing number of cells, number of genes and number of doublets after quality control filtering or ambient RNA correction by sequencing batch; f) Metrics of sc/snRNAseq (ribosomal, mitochondria, RNA levels an expressed genes) before (top) and after (bottom) ambient RNA correction; g) sc/snRNAseq samples before (top) and after (bottom) data integration displayed by batch, sample type and processing method (PDO-organoid, PD-dissociated tumor cells, PF, frozen tumor, indicating the respective processing batches (n=4);

**a**

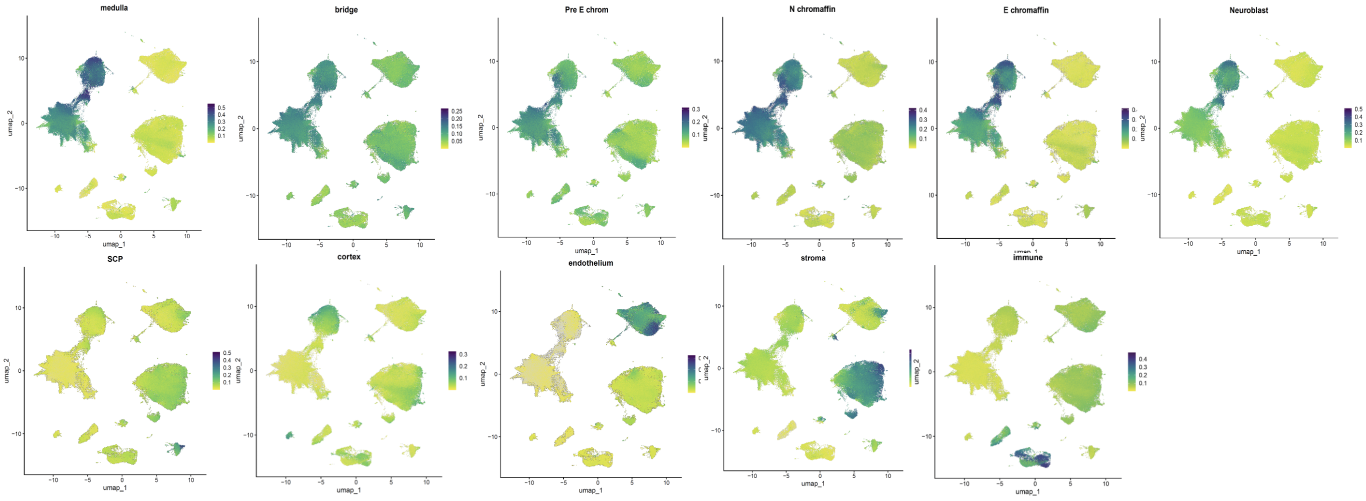

**b**

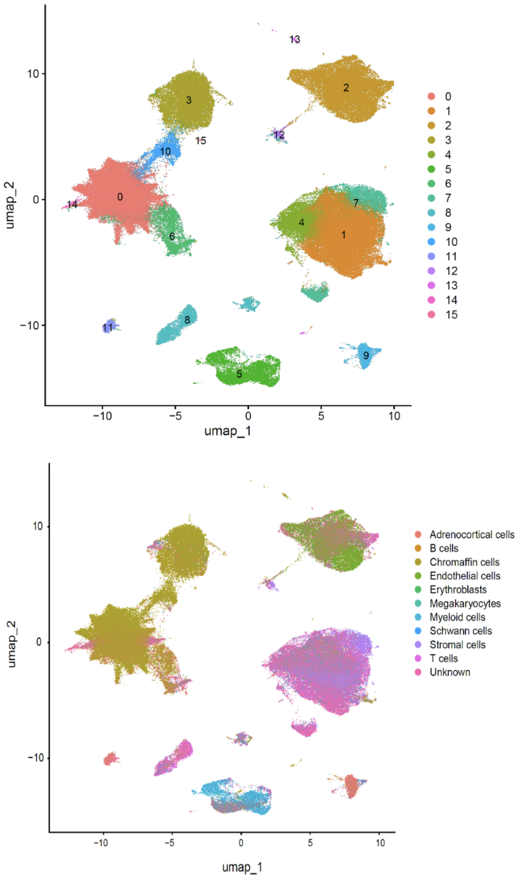

**Supplementary Figure 4.** a) AUCell score cluster cell annotation based on Hannjemaaijer et al, 2021; b) cluster annotation based on Garnett (cluster numbers on top, Garnett cell annotation (bottom);

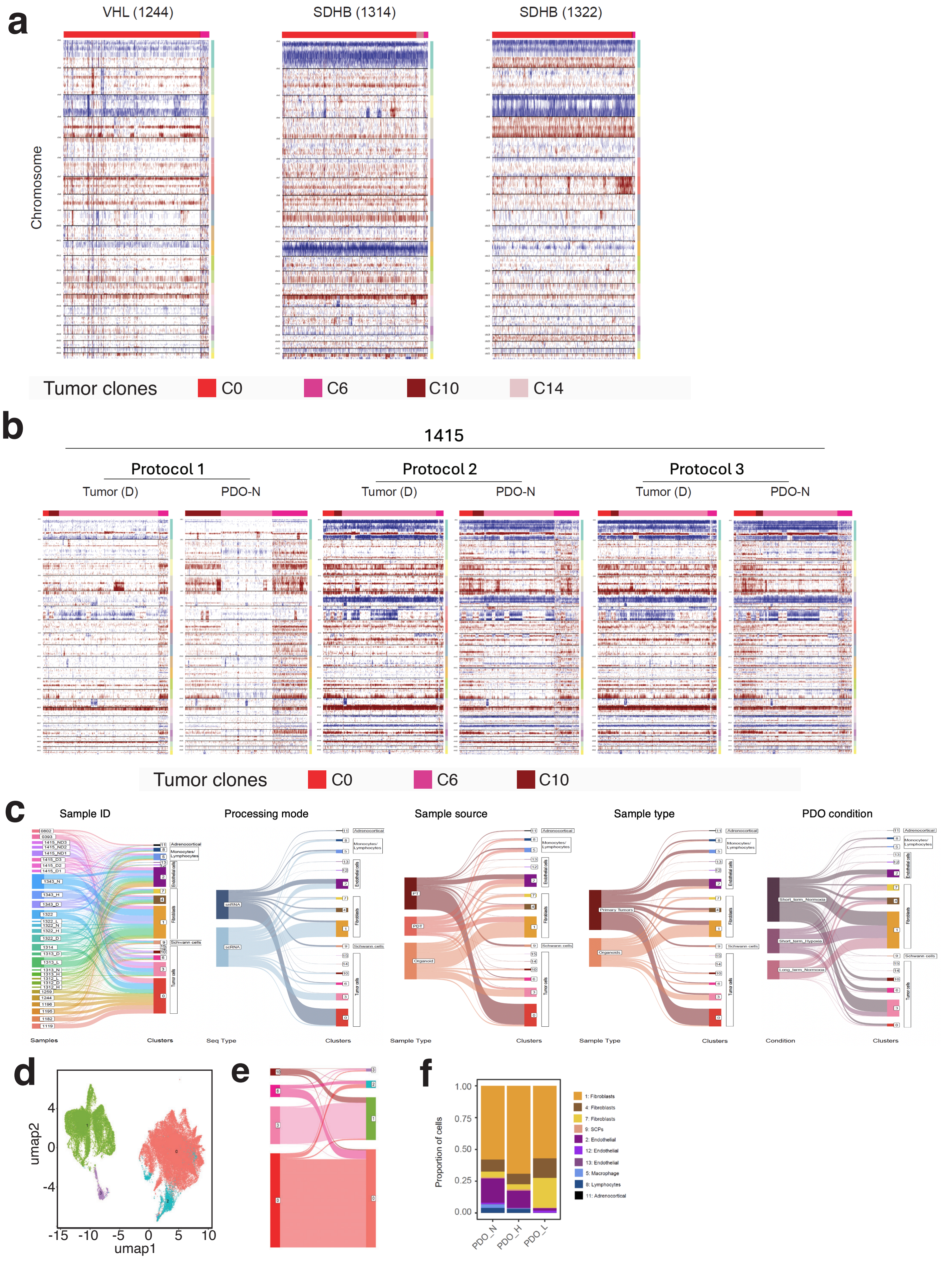
a
VHL (1244)
SDHB (1314)
SDHB (1322)
Chromosome
Tumor clones
C0
C6
C10
C14
b
1415
Protocol 1
Protocol 2
Protocol 3
Tumor (D)
PDO-N
Tumor clones
C0
C6
C10
c
Sample ID
Processing mode
Sample source
Sample type
PDO condition
Samples
Clusters
Seq Type
Sample Type
Condition
snRNA
scRNA
PT
PDT
Organoid
Primary Tumors
Organoids
Short_term_Normoxia
Short_term_Hypoxia
Long_term_Normoxia
Adrenocortical
Monocytes/ Lymphocytes
Endothelial cells
Fibroblasts
Schwann cells
Tumor cells
d
umap2
umap1
e
f
Proportion of cells
PDO_N
PDO_H
PDO_L
1: Fibroblasts
4: Fibroblasts
7: Fibroblasts
9: SCPs
2: Endothelial
12: Endothelial
13: Endothelial
5: Macrophage
8: Lymphocytes
11: Adrenocortical

**Supplementary Figure 5.** a) inferCNV of three representative primary PPGLs with known genotype and chromosomal copy changes; b) inferCNV of the same sample processed with slight modification of the protocol applied to primary tumor; representative primary PPGLs with known genotype and chromosomal copy changes; c) Sankey plots displaying (from left to right) the impact of clusters by sample, processing mode, sample source, sample type and PDO condition; d) UMAP of reclustered tumor clusters generating 4 new clusters 0-3, e) Sankey plot showing the distribution of the new cluster composition relative to the original tumor clusters (d and e are related to Fig 4h); f) Bar plot showing the proportion of nontumor cell clusters across the three culture conditions.

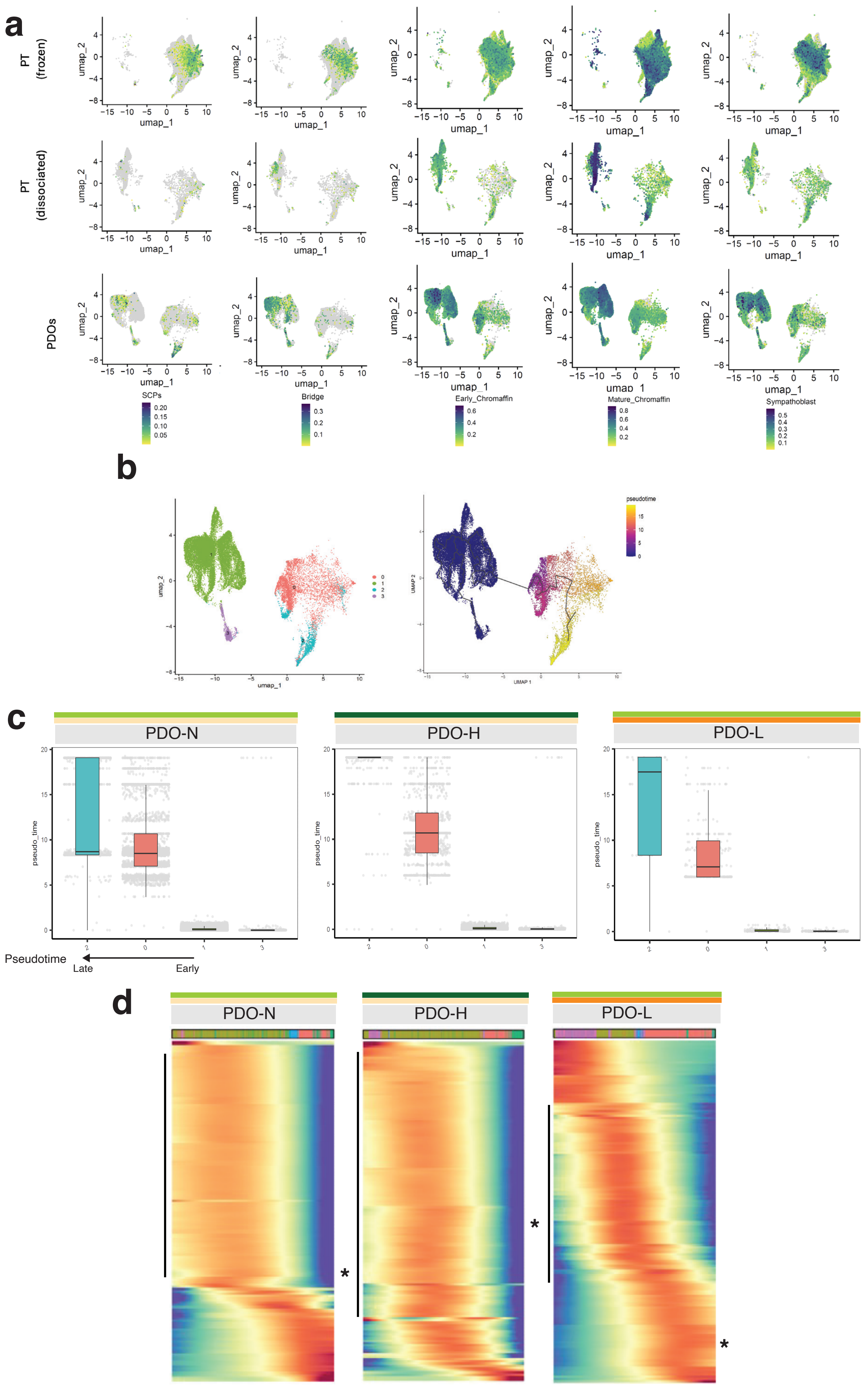
a
PT (frozen)
PT (dissociated)
PDOs
umap_1
umap_2
SCPs
Bridge
Early_Chromaffin
Mature_Chromaffin
Sympathoblast
b
umap_1
umap_2
UMAP 1
UMAP 2
pseudotime
c
PDO-N
PDO-H
PDO-L
pseudo_time
Pseudotime
Late
Early
d
PDO-N
PDO-H
PDO-L
*

**Supplementary Figure 6.** a)UMAP plot of the AUCell score of five cell lineage type signatures in tumor cells, Schwann cell precursor (SCP) cells, bridge cells, early chromaffin cells, mature chromaffin cells and sympathoblasts in primary tumors (PT-frozen, PT-dissociated, and PDOs, the expression scale is depicted (related to Fig 5e); b) UMAP plot of reclustered tumors cells depicting 4 tumor clusters as indicated (left) and corresponding trajectory analysis of primary tumor and PDOs using Monocle 3 (right); c) Derived plots for each PDO condition, ordered from latest to earliest estimated cluster along the pseudotime trajectory; d) Heatmap of Monocle3 data for all three PDO conditions: PDO-N, PDO-H and PDO-L, a dominant cluster (vertical line) of genes that were differentially expressed along the trajectory continuum involves cell respiration, oxidative phosphorylation, the NTRK1 gene (asterisk) shows variable expression timing for each PDO condition (related to Suppl Table 14).

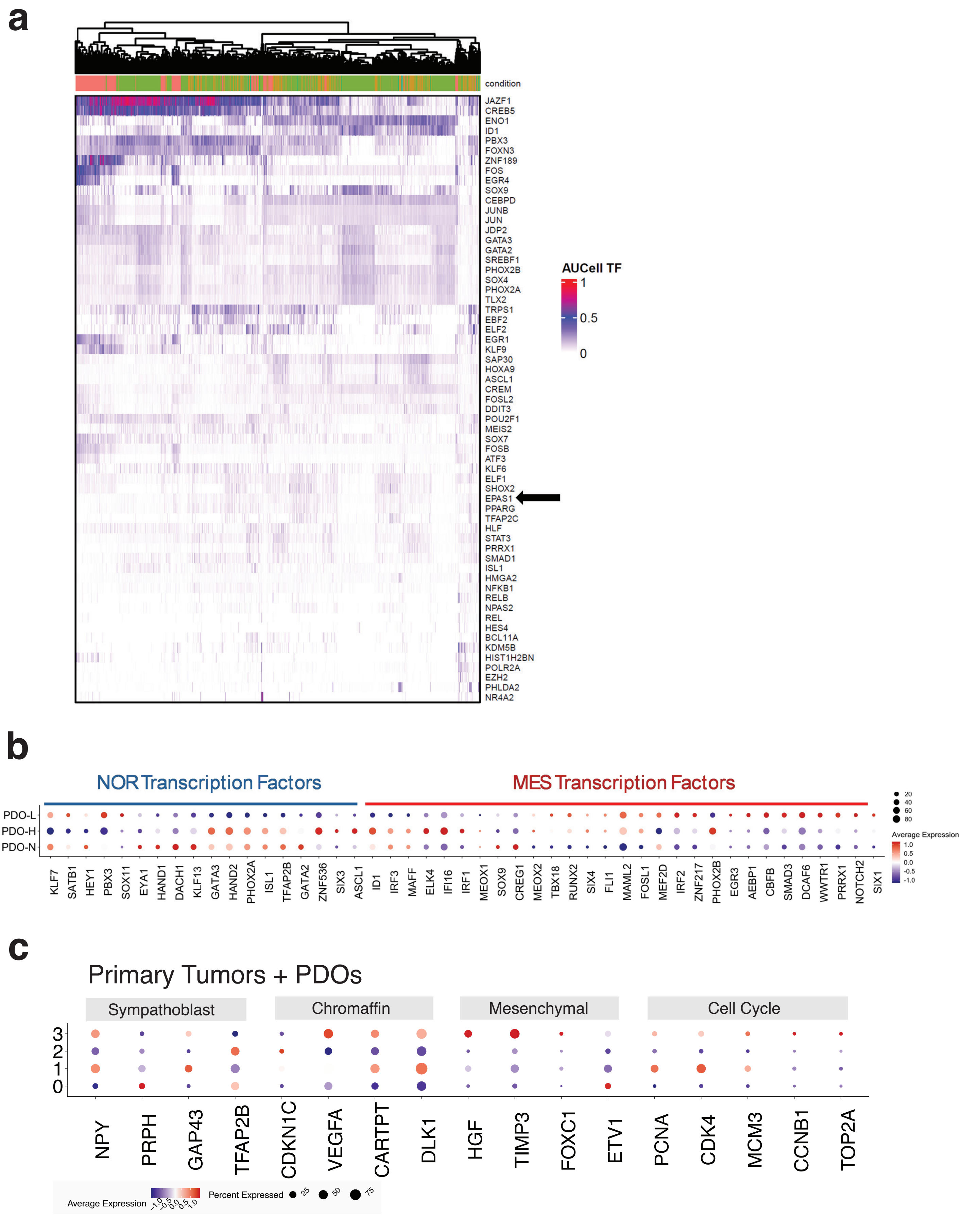
a
condition
JAZF1
CREB5
ENO1
ID1
PBX3
FOXN3
ZNF189
FOS
EGR4
SOX9
CEBPD
JUNB
JUN
JDP2
GATA3
GATA2
SREBF1
PHOX2B
SOX4
PHOX2A
TLX2
TRPS1
EBF2
ELF2
EGR1
KLF9
SAP30
HOXA9
ASCL1
CREM
FOSL2
DDIT3
POU2F1
MEIS2
SOX7
FOSB
ATF3
KLF6
ELF1
SHOX2
EPAS1
PPARG
TFAP2C
HLF
STAT3
PRRX1
SMAD1
ISL1
HMGA2
NFKB1
RELB
NPAS2
REL
HES4
BCL11A
KDM5B
HIST1H2BN
POLR2A
EZH2
PHLDA2
NR4A2
AUCell TF
1
0.5
0
b
NOR Transcription Factors
MES Transcription Factors
PDO-L
PDO-H
PDO-N
KLF7
SATB1
HEY1
PBX3
SOX11
EYA1
HAND1
DACH1
KLF13
GATA3
HAND2
PHOX2A
ISL1
TFAP2B
GATA2
ZNF536
SIX3
ASCL1
ID1
IRF3
MAFF
ELK4
IFI16
IRF1
MEOX1
SOX9
CREG1
MEOX2
TBX18
RUNX2
SIX4
FLI1
MAML2
FOSL1
MEF2D
IRF2
ZNF217
PHOX2B
EGR3
AEBP1
CBFB
SMAD3
DCAF6
WWTR1
PRRX1
NOTCH2
SIX1
20
40
60
80
Average Expression
1.0
0.5
0.0
-0.5
-1.0
c
Primary Tumors + PDOs
Sympathoblast
Chromaffin
Mesenchymal
Cell Cycle
3
2
1
0
NPY
PRPH
GAP43
TFAP2B
CDKN1C
VEGFA
CARTPT
DLK1
HGF
TIMP3
FOXC1
ETV1
PCNA
CDK4
MCM3
CCNB1
TOP2A
Average Expression
-1.0
-0.5
0.0
0.5
1.0
Percent Expressed
25
50
75

**Supplementary Figure 7.** a) py-SCENIC heat map analysis showing resulting 61 significant transcription factor modules in primary and PDO of the indicated condition, the EPAS1 gene is indicated with an arrow; b) Dot plot of transcription factors that are associated with superenhancers defining NOR and MES distribution in neuroblastomas (Related to Fig 6f); c) Dot plot of the expression of selected markers of sympathoblast, chromaffin, mesenchymal and cell cycle in primary tumors (frozen PT and dissociated PDT) and PDOs across individual tumor cell clusters (Related to Fig 6g).

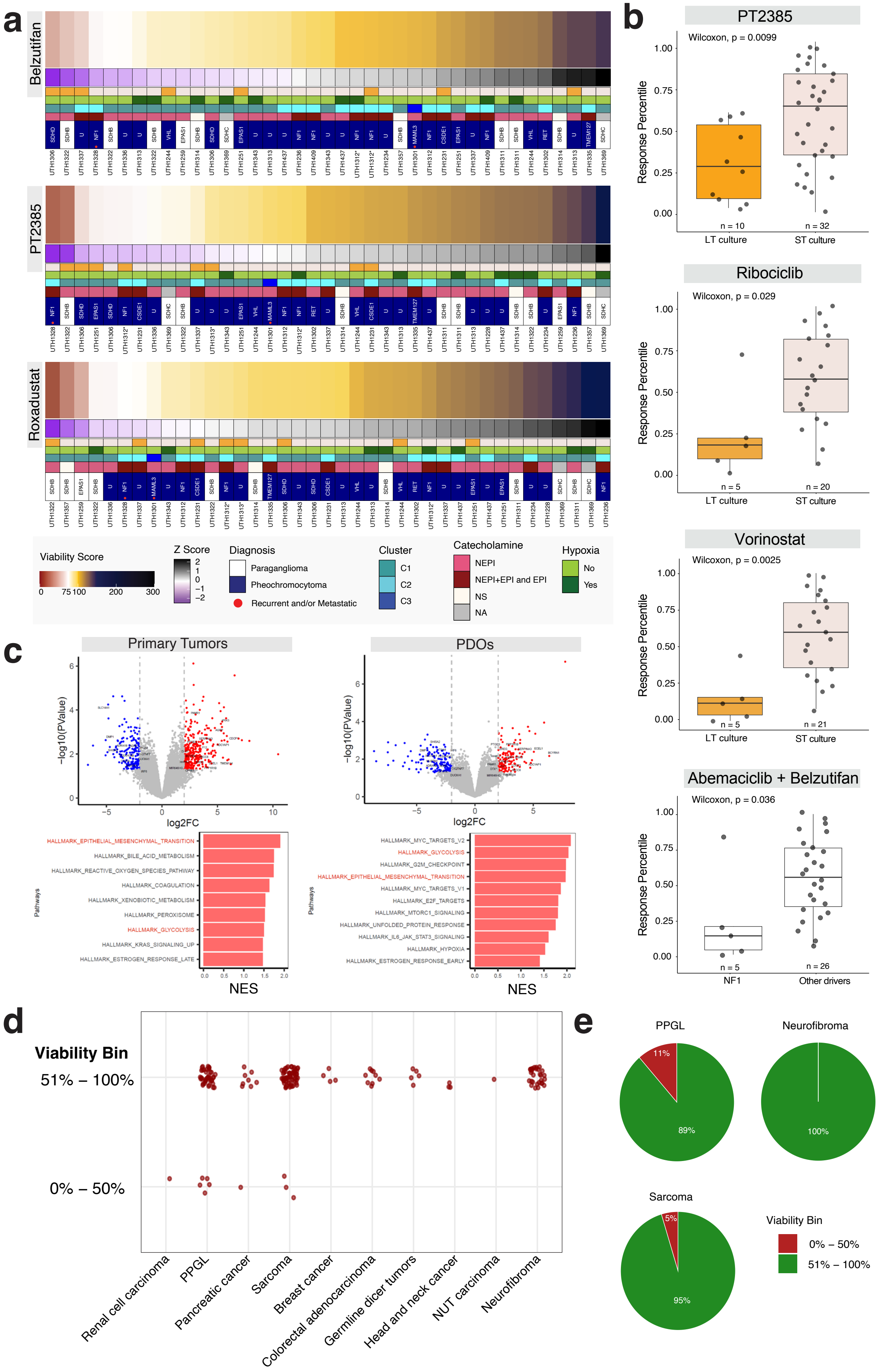

a
Belzutifan
PT2385
Roxadustat
Viability Score
0
75
100
200
300
Z Score
2
1
0
−1
−2
Diagnosis
Paraganglioma
Pheochromocytoma
Recurrent and/or Metastatic
Cluster
C1
C2
C3
Catecholamine
NEPI
NEPI+EPI and EPI
NS
NA
Hypoxia
No
Yes
b
PT2385
Wilcoxon, p = 0.0099
Response Percentile
n = 10
n = 32
LT culture
ST culture
Ribociclib
Wilcoxon, p = 0.029
n = 5
n = 20
Vorinostat
Wilcoxon, p = 0.0025
n = 5
n = 21
Abemaciclib + Belzutifan
Wilcoxon, p = 0.036
n = 5
n = 26
NF1
Other drivers
c
Primary Tumors
PDOs
−log10(PValue)
log2FC
Pathways
NES
HALLMARK_EPITHELIAL_MESENCHYMAL_TRANSITION
HALLMARK_BILE_ACID_METABOLISM
HALLMARK_REACTIVE_OXYGEN_SPECIES_PATHWAY
HALLMARK_COAGULATION
HALLMARK_XENOBIOTIC_METABOLISM
HALLMARK_PEROXISOME
HALLMARK_GLYCOLYSIS
HALLMARK_KRAS_SIGNALING_UP
HALLMARK_ESTROGEN_RESPONSE_LATE
HALLMARK_MYC_TARGETS_V2
HALLMARK_GLYCOLYSIS
HALLMARK_G2M_CHECKPOINT
HALLMARK_EPITHELIAL_MESENCHYMAL_TRANSITION
HALLMARK_MYC_TARGETS_V1
HALLMARK_E2F_TARGETS
HALLMARK_MTORC1_SIGNALING
HALLMARK_UNFOLDED_PROTEIN_RESPONSE
HALLMARK_IL6_JAK_STAT3_SIGNALING
HALLMARK_HYPOXIA
HALLMARK_ESTROGEN_RESPONSE_EARLY
d
Viability Bin
51% – 100%
0% – 50%
Renal cell carcinoma
PPGL
Pancreatic cancer
Sarcoma
Breast cancer
Colorectal adenocarcinoma
Germline dicer tumors
Head and neck cancer
NUT carcinoma
Neurofibroma
e
PPGL
11%
89%
Neurofibroma
100%
Sarcoma
5%
95%
Viability Bin
0% – 50%
51% – 100%

**Supplementary Figure 8.** a) Heatmaps of PDO sensitivity to selected drugs including viability score ranges from 0 to 300% (scores above 100% indicate growth in response to drug). The score is normalized to the mean response to treatment across all samples. Each column is a unique PDO sample, red indicates higher sensitivity to treatment than average, and blue indicates higher viability than the average: Belzutifan, PT-2385 and Roxadustat; b) Sensitivity rank plots comparing the response of the specified PDO groups; c)Volcano plots showing differentially expressed genes between Abemaciclib responders and non-responders in primary tumor (left) and PDOs (right), significantly upregulated and downregulated are indicated in red and blue, respectively, and common genes between the two sets of samples are labeled, bar plots indicate GSEA NES scores of significantly enriched Hallmark pathways in responder samples of each data set, with common pathways highlighted in red (Related to Suppl Table 19); d) Viability to 1$\mu$M Abemaciclib was assessed across 139 PDOs of 10 distinct tumor lineages including epithelial, mesodermal and neural crest tumors, as indicated. For each sample, viability was categorized as either above or below 50%; e) For all tumor types with at least 10 samples tested, Abemaciclib responses were summarized in a pie chart, showing the percentage of samples within each tumor type that exhibited ≤50% viability.
